# Selection maintains genetic coupling of functional trait trade-offs despite gene flow in an oak hybrid zone

**DOI:** 10.64898/2026.05.18.725696

**Authors:** Ryosuke K. Ito, Kaede Konrai, Kurita Omura, Seiya Sunayama, Yusuke Onoda, Yuji Isagi

## Abstract

Resource allocation trade-offs during seedling establishment can generate coordinated interspecific variation in functional traits that reflects contrasting growth–survival strategies. Yet whether and how natural selection maintains these trade-offs and their multilocus genetic architecture remain poorly understood. Using a natural hybrid zone between two East Asian oaks, we evaluated whether loci associated with these trade-offs showed signatures of selection and genetic coupling despite ongoing hybridisation. Acorns collected across the zone were cultivated in a common garden, where we quantified 26 seedling growth and functional traits, and analysed whole-genome variation. We identified coordinated trade-offs within leaf function and plant architecture. Trait-associated genomic regions demonstrated excess inter-chromosomal linkage disequilibrium in the hybrids, consistent with genetic coupling, and exhibited elevated interspecific differentiation and steeper local genomic clines than background loci. Together, these results suggest that selection contributes to maintaining resource-allocation trade-offs and genetic coupling among their underlying loci despite ongoing gene flow.

## Introduction

Tree species diversity and functional diversity in forests underpin ecosystem functions (Loreau et al., 2001; Cardinale et al., 2012; Slik et al., 2015). Functional trait variation among tree species is structured along coordinated axes of resource acquisition and use, growth, and tissue maintenance, collectively described as functional trait spectra (Wright et al., 2004; Reich, 2014; Díaz et al., 2016). In forest trees, organ-level trait strategies governing resource acquisition and tissue persistence in leaves, roots, and stems are thought to correspond to whole-plant strategies of growth and survival (Reich, 2014; Rüger et al., 2018). Resource-allocation constraints underlying these strategies can generate growth–survival trade-offs, in which rapid resource acquisition and growth, potentially enhancing competitive performance, come at the cost of tissue persistence, survival, and tolerance to environmental stress (Wright et al., 2010; Rüger et al., 2018, 2020). Such trade-offs may be particularly consequential during seedling establishment, when resource limitation and high mortality make variation in growth, survival, and stress tolerance particularly important for successful recruitment (Sterck et al., 2011; Rüger et al., 2018).

Although extensive ecological evidence documents coordinated variation between functional traits and demographic rates, as well as patterns consistent with resource-allocation trade-offs, the evolutionary and genetic bases of these patterns remain poorly understood (Wang et al., 2024). Molecular studies of specific functional trade-offs, particularly that between growth and defence, have shown that alternative functions can be coordinated through shared regulatory mechanisms, including phytohormone signalling (Züst & Agrawal, 2017; He et al., 2022). Such coordination can, in principle, have a relatively simple genetic basis when one or a few regulatory loci exert pleiotropic effects on multiple functions. In contrast, organism-level strategies involving growth, survival, defence, and resource use are likely to emerge from coordinated variation across multiple traits and physiological pathways. Their genetic basis may therefore involve not only pleiotropic effects of individual loci but also non-random associations among alleles at multiple loci—that is, linkage disequilibrium—which can couple the selective effects of those loci (Barton & Hewitt, 1985; Butlin & Smadja, 2018). Because recombination erodes such multilocus associations, the evolution and maintenance of integrated strategies involving multiple loci may be more difficult than those arising from pleiotropic effects at individual loci. This challenge may be particularly pronounced under ongoing gene flow, which reduces allele-frequency differentiation between populations or species. However, few studies have tested whether natural selection can maintain integrated multivariate strategies and their underlying genetic associations despite the effects of gene flow and recombination (Wang et al., 2024).

Hybrid zones between closely related species provide natural systems in which the interactions among natural selection, gene flow, and recombination can be examined simultaneously. Gene flow tends to reduce interspecific differences in allele frequencies, whereas recombination erodes associations among alleles at different loci, particularly when those loci are unlinked. The joint persistence of species-associated multivariate strategies and excess associations among their underlying loci under these conditions could therefore indicate that selection counteracts the loss of allele-frequency differentiation and multilocus association caused by gene flow and recombination. However, population structure and recent admixture can generate linkage disequilibrium even without selection maintaining particular allele combinations, whereas variation in recombination rate and linked selection can produce heterogeneous patterns of genomic differentiation (Wolf & Ellegren, 2017; Schield et al., 2024). Distinguishing among these processes therefore requires the integration of phenotypic differentiation, genotype–phenotype associations, genomic signatures of selection, and patterns of ancestry and linkage disequilibrium. Hybrid zones thus provide an opportunity to test whether natural selection contributes to maintaining the genetic architecture of alternative resource-allocation strategies despite ongoing gene flow (Haenel et al., 2021; Schield et al., 2024).

The elevational hybrid zone between *Quercus mongolica* var. *crispula* (Qmon) and *Q. serrata* subsp. *serrata* (Qser) provides a suitable model system for testing whether natural selection maintains divergent multivariate seedling strategies despite ongoing gene flow (Aizawa et al., 2021). Phylogenetic relationships and demographic modelling indicate that the Japanese hybrid zone between Qmon and Qser most likely formed through secondary contact (Hipp et al., 2020; Tamaki et al., 2021). Both species are widely distributed across East Asia but show contrasting distributions along latitudinal and elevational gradients (Figure 1A). In Japan, they form narrow hybrid zones (2.5–4.9 km; Ito et al., unpublished data), across which environmental conditions vary with temperature and maximum snow depth. *Quercus* species are wind-pollinated, with pollen dispersal ranging from several hundred metres to several kilometres (Ashley, 2021), which facilitates interspecific gene flow and syngameon formation (Cannon & Petit, 2020). Because seedling establishment is a strong demographic filter, divergence in resource allocation and early growth during this stage may strongly influence recruitment and thereby contribute to functional and ecological differentiation between the species. Together, the sharp environmental transition, ongoing gene flow, and demographic importance of early establishment make this system well suited for integrating cline modelling, common-garden experiments, and genome-wide analyses to test whether natural selection contributes to maintaining divergent seedling strategies and their multilocus genetic basis.

**Figure 1.**
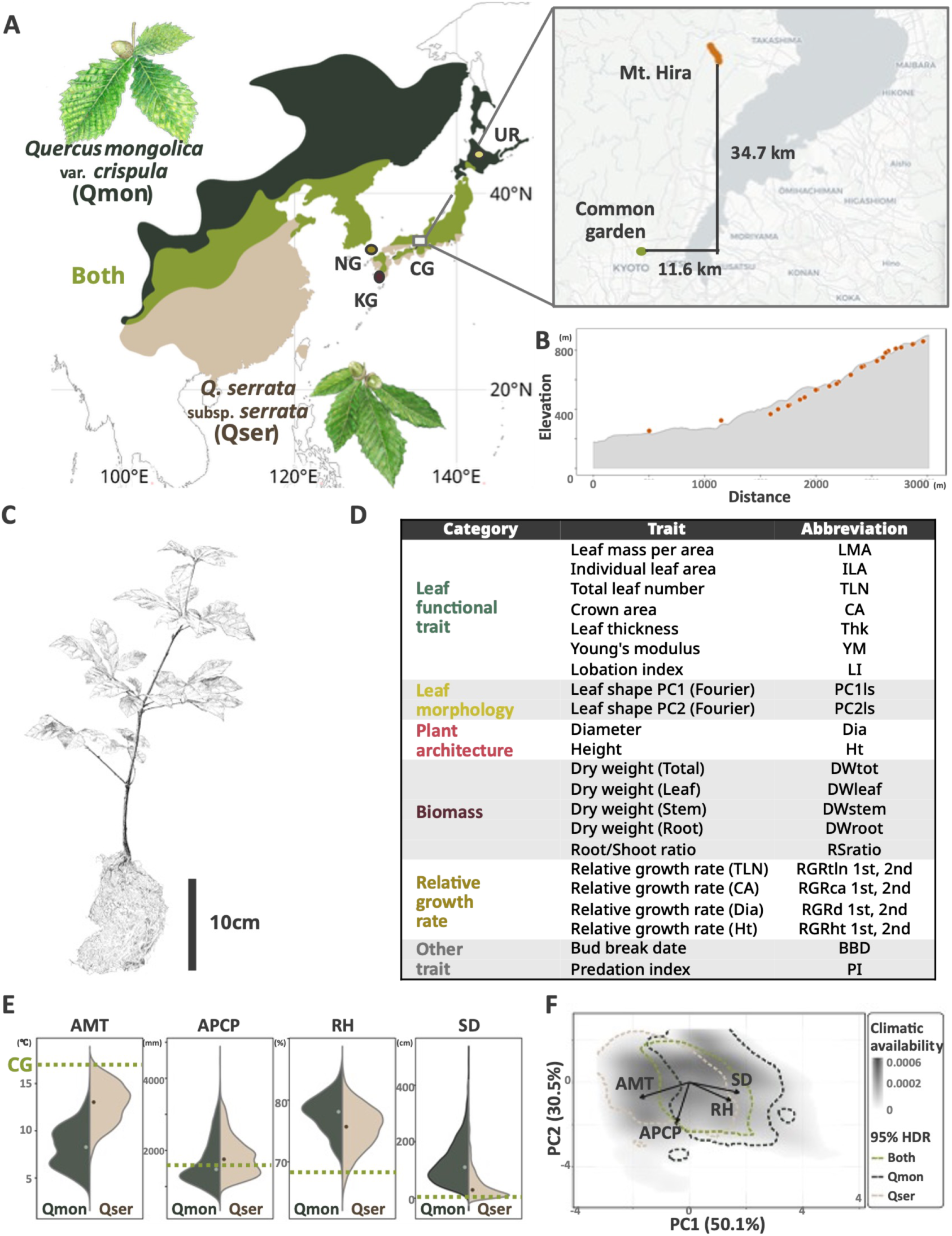
Study system, measured traits, and environmental context. (A) Geographic ranges of *Quercus mongolica* var. *crispula* (Qmon) and *Q. serrata* subsp. *serrata* (Qser) with sampling localities. Dark green, beige, and light green indicate Qmon-only, Qser-only, and overlap, respectively. Circles (UR, NG, and KG) represent pure-line sampling sites; the rectangle indicates the acorn sampling area in the Hira Mountains (hybrid zone) and the location of the common garden (CG). (B) Acorn sampling sites along the elevational transect at Mt. Hira. (C) Representative seedlings photographed after two growing seasons. (D) The 26 measured or derived seedling traits with abbreviations and categories (see Table S2 for details). (E) Comparison of environmental variables between Qmon and Qser occurrence sites in Japan and the CG conditions (dashed line): annual mean temperature (AMT), annual precipitation (APCP), maximum snow depth (SD), and annual mean relative humidity (RH). Occurrence data are from the Forest Agency’s Biodiversity Baseline Survey; climate data are from the agro-meteorological grid square data, NARO. (F) Realised climatic niches of Qmon and Qser based on PCA-env using the four climatic variables in (E). Dashed lines indicate the climatic niche space occupied by each species and their overlap, while “Both” indicates co-occurrence sites representing 0.1-ha plots in which both species were recorded. Arrows indicate the loadings of the climatic variables on the PCA axes.

In this study, we integrated phenotypic and genomic evidence to evaluate the hypothesis that natural selection contributes to maintaining divergent multivariate seedling strategies and genetic coupling among trait-associated loci in the Qmon–Qser hybrid zone. Testing this hypothesis required evidence that (i) ongoing interspecific gene flow occurs across the hybrid zone; (ii) seedling traits exhibit coordinated variation consistent with a growth–survival trade-off and are associated with genomic ancestry; (iii) trait-associated loci exhibit signatures of natural selection and greater interspecific differentiation at these loci than at background loci; and (iv) associations among these loci are stronger than expected from population structure and admixture alone.

## Materials and methods

### Climatic niche differentiation and co-occurrence between *Quercus mongolica* and *Q. serrata*

To evaluate climatic niche differentiation and local co-occurrence between *Quercus mongolica* var. *crispula* (Qmon) and *Q. serrata* subsp. *serrata* (Qser), we examined species-specific climatic responses, overlap in their realized climatic niches, and the climatic conditions associated local co-occurrence. Species occurrence records were obtained from 0.1-ha plots systematically distributed across Japan on an approximately 4-km grid as part of the National Forest Inventory conducted by the Forestry Agency of Japan. Only trees with a diameter at breast height (DBH) ≥18 cm were included. We defined a species as present when at least one qualifying tree was recorded within a plot and defined local co-occurrence as the presence of both species within the same plot. After excluding sites with incomplete data, we assigned 1-km-resolution climate data for 2016–2025, obtained from the Agro-Meteorological Grid Square Data System of the National Agriculture and Food Research Organization to 16,169 plots. Annual mean temperature (AMT), annual precipitation (APCP), maximum snow depth (SD), and relative humidity (RH) were selected after screening for collinearity and were standardized before analysis.

We first evaluated climatic differentiation while accounting for spatial autocorrelation using a spatial-factor joint species distribution model implemented with spOccupancy (sfJSDM; Doser et al. 2022). Linear and quadratic terms were included for each climatic variable, and interspecific differences were assessed based on whether the 95% Bayesian credible intervals for the differences between species-specific coefficients excluded zero. We then compared the realized climatic niches of the two species using PCA-env, with all surveyed plots defining the common available environmental background (Broennimann et al. 2012). Niche overlap was quantified using Schoener’s *D*. Finally, to describe where local co-occurrence occurred within the shared climatic space, we identified the intersection of the 95% highest-density regions of the two species and calculated the proportion of its area in PCA-env space that was contained within the 95% highest-density region of the co-occurrence sites. Details of data processing, variable selection, model formulation, prior distributions, MCMC settings, and PCA-env procedures are provided in Supplementary Methods 1.

### Plant material and growth conditions

We collected 316 acorns from the forest floor at 48 sites along an elevational gradient in the natural hybrid zone of Qmon and Qser in the Hira Mountains, central Japan (Shiga Prefecture) in April 2022. At each site, acorns were collected from spatially dispersed points within a radius of approximately 50 m to reduce repeated sampling from the same maternal tree, although maternal identities could not be determined. Acorns were germinated outdoors in a common garden at Kyoto University (Sakyo-ku, Kyoto City, Kyoto Prefecture; Figure 1A, B), and each seedling was transplanted into an individual 15-cm pot approximately one week after germination. The individual pots were arranged in 21 spatial blocks, with seedlings allocated so that collection elevations were distributed as evenly as possible among blocks and positions randomised within each block. All acorns and seedlings received standardised fungicide, pesticide, irrigation, and fertiliser treatments. Full details of cultivation conditions, experimental layout, and phenotypic measurements are provided in Supplementary Methods 2. At the end of the second growing season, seedling traits were measured in 240 surviving individuals, of which 223 were included in the whole-genome analyses. To assess whether putatively pure parental individuals of Qmon and Qser were present in the hybrid zone samples, we collected parental species reference samples showing typical leaf morphology from three allopatric sites where only one species occurred. These reference samples comprised three Qmon individuals from northern Hokkaido (UR); two Qser individuals from Tsushima Island, Nagasaki Prefecture (NG); and two individuals from western Kagoshima Prefecture (KG; Figure 1A).

### Whole genome sequencing, read mapping, and genotype calling

Genomic DNA was extracted using a Qiagen DNeasy Plant Mini Kit (Qiagen, Germany) and a modified CTAB method (Porebski et al., 1997). Libraries were prepared using Illumina DNA Prep (Illumina, USA) and Nextera-compatible indexing primers (IDT, USA) following a modified protocol (Gaio et al., 2022). Shotgun sequencing was performed on a DNBSEQ-G400 platform (MGI, China) and conducted by BGI (China) and Azenta (USA).

Read preprocessing was performed using fastp v1.0 (Chen, 2025) to remove adapters and trim low-quality bases and reads. The reads were mapped to the *Q. mongolica* genome assembly (GenBank accession GCA_011696235.1; Ai et al., 2022) using BWA-MEM2 (Md et al., 2019). Duplicate reads were marked using Picard v3.4 (Broad Institute, 2025), and local realignment around indels was performed using GATK v3.8 (Van der Auwera & O’Connor, 2020). Variant calling was performed using ANGSD v0.94 (Korneliussen et al., 2014) based on genotype likelihood (filtering options: -GL 1, -minMapQ 30 -minQ 20 -minInd 200 - setMinDepth 3 -SNP_pval 1e-6 -doMajorMinor 1 -doMaf 1). For genotype imputation and haplotype calling, we applied a two-step procedure: we initially performed pre-imputation on genotype likelihoods using Beagle v3.3 (Browning & Browning, 2011), masked sites with a genotype probability (GP) of < 0.9, and subsequently conducted post-imputation using Beagle v5.5 (Browning et al., 2021). To improve the accuracy, we jointly analysed these data with whole-genome data from 784 adult individuals of Qmon and Qser generated by Ito et al. (in preparation), resulting in a dataset of 1,007 individuals. To evaluate whether mapping performance varied systematically with genomic ancestry, we examined the relationships between HI and several mapping quality-control metrics. The proportions of properly paired reads, singleton reads, supplementary alignments, and reads whose mates mapped to different chromosomes were calculated from samtools flagstat output. Pearson’s correlation coefficients between hybrid index and each mapping quality-control metric were calculated in R. All correlations were weak (|*r*| ≤ 0.156), providing no evidence of ancestry-associated differences in these mapping metrics (Table S1).

Two SNP datasets were generated for downstream analyses. SNP set A (3.3 million SNPs) comprised the post-imputation genotypes and was used for genome-wide analyses, including admixture mapping and selection scans, without additional filtering. SNP set B (0.2 million SNPs) was constructed as a putatively neutral dataset for population-genetic analyses: we removed sites within gene regions and their ±10-kb flanks using bedtools v2.18 (Quinlan, 2014) and bcftools v1.22, and then performed LD pruning in PLINK v1.9 (Gaunt et al., 2007; Chang et al., 2015) at r² = 0.2. The analysis workflow and code are available in the GitHub repository.

### Population structure and hybridisation analysis

For genetic characterisation of the hybrid zone, we assessed the evidence of hybridisation, classified individuals into hybrid classes (F1, F2, and both backcrosses), and fitted genetic clines. Using SNP set B, we inferred the population genetic structure using genetic PCA (PLINK v2.0) and ADMIXTURE v1.3 (Alexander & Lange, 2011). In ADMIXTURE, we evaluated the values of K (number of ancestral clusters) from 1 to 8 and selected K = 2, which resulted in the lowest cross-validation (CV) error. Next, to test whether hybridisation is ongoing and long-standing, we inferred hybrid class membership from the relationship between hybrid index (HI) and interspecific heterozygosity (Het) using triangulaR v1.14 (Wiens et al., 2025). To generate triangulaR inputs, we thinned SNP set A to one SNP per 10 kb using vcftools v0.1.17 (Danecek et al., 2011). Based on the PCA and ADMIXTURE results, we selected five parental-species reference individuals per species and defined 627 ancestry-informative markers (AIMs) as loci with an allele-frequency difference (Δp) ≥ 0.9 between the two reference groups. To evaluate the spatial segregation within the hybrid zone and the effects of environmental factors, we fitted genetic clines using both a sampling site-based approach (hzar v0.2; Derryberry, 2014) and an individual-based approach (PyMC v5.26; Abril-Pla et al., 2023). To compare the relative contributions of dispersal and environment to cline shape, we fitted models using either elevation (environmental distance) or geographic distance as the cline axis and compared the models using AICc (hzar) and LOOIC (PyMC).

### Phenotypic measurements and trait derivation

At the end of the second growing season, we measured 26 phenotypic traits and grouped them into six categories: “leaf functional trait”, “leaf morphology”, “plant architecture”, “biomass”, “relative growth rate”, and “other trait” (Figure 1C, D; Table S2). The detailed measurement and calculation procedures are provided in Supplementary Methods 1. The crown area (CA) and leaf image data were archived in Dryad.

### Seedling performance versus hybrid index

We evaluated the relationship between seedling vigor/depression under common garden conditions and the ancestry proportion between Qmon and Qser using the germination rate, survival over two growing seasons, and total dry weight after two seasons (DWtot). Germination and survival rates were calculated for each collection site, and their relationships with the site-mean hybrid index (HI) were estimated using generalised additive models (GAMs; mgcv v.1.9; Wood, 2011; Figure 1B). To stabilise the inference, models for germination and survival were weighted by the number of individuals per site, whereas DWtot was analysed at the individual level.

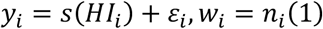

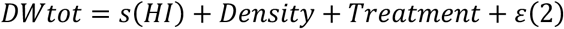

where *y_i_* is the germination or survival rate at site *i*, *HI_i_* is the mean HI at site *i*, and *w_i_* is the weight given by the site sample size *n_i_*. For the DWtot model, we included common-garden covariates describing local growing conditions (Density: the number of neighbouring seedlings within the garden; Treatment: heat-stress experimental treatment category [stress/control/not included]; Ito et al., in preparation).

Because the HI could not be determined for non-germinated acorns, the analyses may be affected by statistical bias. To evaluate the potential effect of this missingness, we conducted a supplementary analysis treating elevation as a proxy for HI, initially estimating the positive relationship between elevation and HI using a GAM with the same model specification, and subsequently dividing the elevational range into eight bins (83 m per bin). We calculated the germination and survival rates within each bin and compared their trends to assess whether HI missingness owing to non-germination affected the main conclusions.

To assess interspecific differences in photosynthetic capacity per unit leaf area and their relationship with HI, we measured leaf gas exchange traits. First, stomatal conductance (gsw) was measured using LI-600 (LI-COR, USA). For the 93 individuals in the garden, we selected three healthy upper leaves per individual and conducted two replicate measurements. In addition, we measured the photosynthetic rate on a leaf area basis (*A_area_*) of 15 individuals using an LI-6800 (LI-COR, USA). Measurements were conducted during September of the first growing season (07:30–12:45) using one healthy upper leaf per individual. Chamber conditions were set to a reference *CO*_2_ concentration of 420 ppm, leaf temperature of 30 °C, and photosynthetic photon flux density of 1000 μmol *m*^−2^ *s*^−1^. During the measurements, the leaf vapour pressure deficit was maintained at approximately 1 kPa, indicating that the plants were not exposed to strong atmospheric drought stress. The relationships between HI and gas-exchange traits were estimated using GAMs, while accounting for environmental covariates (leaf temperature, leaf vapour pressure deficit, and incident light):

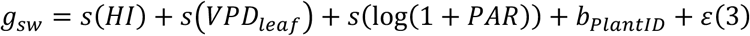

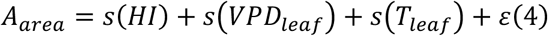

where *VPD_leaf_* is the leaf vapour pressure deficit, *T_leaf_* is the leaf temperature, and *b_PlantID_* represents an individual-level random effect (PlantID).

### Phenotypic data analysis and structural equation modelling

To characterise resource allocation-based trait covariation and how the ancestry proportion (HI) in the Qmon–Qser hybrid zone influences this structure, we implemented a four-step analytical workflow: (i) imputation of missing trait values, (ii) estimation of trait–trait correlations, (iii) identification of putative trade-offs (negative trait associations) and quantification of the effects of HI and total dry weight on these patterns, and (iv) inference of a comprehensive trade-off architecture using structural equation modelling (SEM).

i. Missing trait values were imputed using the random-forest algorithm implemented in missForest v1.6 (Stekhoven & Bühlmann, 2012). To assess the imputation accuracy, we randomly masked 10% of the observed values and compared the imputed versus observed values to obtain high concordance (Pearson’s *r* = 0.99; NRMSE = 0.16).
ii. Trait–trait associations were summarised using Spearman’s rank correlations (Hmisc v5.2; Harrell, 2025) and visualised with corrplot v0.95 (Wei et al., 2024).
iii. We performed PCA on the 24 traits, excluding total dry weight (centred and scaled), to obtain trait loadings and individual scores. We then fitted the GAMs to quantify the associations between each PC and the HI.

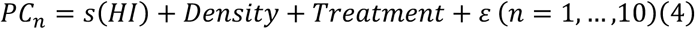

To evaluate how DWtot varies with multivariate trait variation captured by PCs, we fitted a GAM including PC1–PC10 simultaneously as smoothing terms:

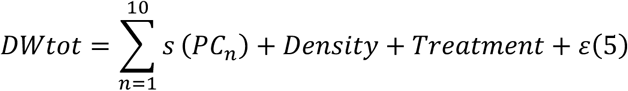

For the PC–HI analyses, P-values across PCs were adjusted using the Benjamini–Hochberg procedure (Benjamini, 1995).
(iv) We used piecewiseSEM v2.3 (Lefcheck, 2016) to construct SEMs in which HI affected multiple traits (Figure S1). Model fit was assessed using Fisher’s C and Akaike information criterion (AIC) from d-separation tests, and the path structure was refined by iteratively adding or removing paths based on conditional-independence violations (P < 0.05) and weak effects (|β| < 0.10 and P > 0.05). The details are provided in Supplementary Methods 2.

### Admixture mapping for genome-wide association analysis of seedling traits

To infer genomic loci associated with seedling trait variation, we performed admixture mapping using GEMMA v0.98 (Zhou & Stephens, 2012). To mitigate confounding from shared causal and correlational structures across traits, we employed a multivariate linear mixed model (mvLMM) and jointly analysed PC1–4 as traits (cumulative proportion of variance explained > 0.5). We used SNP set A as the genotype dataset and converted it to the GEMMA input format using plink2. Following McFarlane & Pemberton (2021), we included, as additional fixed effects, the individual-level Q-score matrix representing genomic background as well as the number of neighbouring individuals within the common garden and the heat-stress treatment group (stress/control/not included) to represent cultivation conditions (“-not-snp”). As a random effect, we used a relatedness (kinship) matrix estimated using the GEMMA. For outlier detection, we used P-values from Wald tests in the mvLMM and applied a Benjamini–Hochberg correction using adjP < 0.05.

To define the genomic interval for identifying genes associated with significant SNPs, we estimated linkage disequilibrium (LD) decay separately in *Q. serrata* parental-type (HI < 0.07) and *Q. mongolica* parental-type (HI > 0.93) individuals using PopLDdecay v3.4 (Zhang et al., 2019). In both parental-type groups, pairwise LD declined to (*r*^2^ = 0.1) at approximately 25 kb. We therefore defined significant SNPs as trait loci and extracted genes located within 25 kb upstream or downstream of each locus using bedtools for subsequent analyses.

### Selection scans

To identify genomic regions influenced by natural selection, we conducted outlier detection based on five metrics (XP-EHH, *F_st_*, *dXY*, π, and Tajima’s *D*). To minimise the effects of introgression, we classified common-garden individuals into three groups based on HI: Qser parental-type (HI < 0.07), Hybrid (0.07 ≤ HI ≤ 0.93), and Qmon parental-type (HI > 0.93), and calculated each metric using only the parental-type subsets.

First, we scanned for recent strong positive selection using XP-EHH, computed between the parental species using rehh v3.2 (Gautier & Vitalis, 2017). Outliers were defined as the upper 1% and lower 1% tails of the genome-wide XP-EHH distribution (corresponding to the extended haplotypes in Qser and Qmon, respectively). Genes within ±25 kb of outlier sites were extracted using bedtools. Next, we screened for highly differentiated genomic regions in the parental species using *F_st_* and *dXY*. We computed both statistics in 10-kb windows using GenoPop v0.9 (Gurke & Mayer, 2025), designated the upper 1% of windows genome-wide as outliers, and extracted genes overlapping these windows. Finally, to identify within-species signatures consistent with reduced diversity and/or departures from neutrality, we calculated nucleotide diversity (π) and Tajima’s *D* in 10-kb windows for each parental species using GenoPop. We defined outliers as the lower 1% tail of the genome-wide window distribution for each statistic and extracted genes that overlapped the outlier windows.

### Enrichment analysis

We conducted a gene set enrichment analysis in ShinyGO v0.85 (Ge et al., 2020) for genes that were significant in admixture mapping (adjP < 0.05) and identified them as outliers in at least one selection-scan metric. Gene annotations were assigned through BLASTP (Camacho et al., 2009) searches against two reference species supported by ShinyGO: *Arabidopsis thaliana* (TAIR10) and *Quercus lobata* (ValleyOak3.0; Ensembl Plants). For each gene, we retained the best hit with an e-value threshold of 1.0 × 10⁻⁴. Enrichment analysis was performed separately for each reference using the Gene Ontology biological process, cellular component, and molecular function categories available in ShinyGO. For each reference, the background set comprised all genes that received an annotation, and ShinyGO’s default settings were used for statistical tests. P-values were adjusted using the Benjamini–Hochberg procedure, with adjP < 0.05 considered significant.

### Evaluation of genetic coupling and divergence among trait loci

To test for gene coupling among trait-associated regions (trait loci), we quantified whether inter-chromosomal linkage disequilibrium (ICLD) among trait loci exceeded the genomic background expected under admixture. Because recent admixture can generate LD among loci that differ in allele frequency between parental populations, high absolute ICLD among trait loci with divergent allele frequencies between parental populations, high absolute ICLD among trait loci alone was not considered evidence of genetic coupling. We initially removed loci with minor allele frequency (MAF) < 0.10 from SNP set A and, as in the selection-scan analyses, classified common-garden individuals into three HI-based groups: hybrids (0.07 ≤ HI ≤ 0.93), Qser parental-type (HI < 0.07), and Qmon parental-type (HI > 0.93). Because ICLD depends on interspecific allele-frequency differences, we calculated Δp for each SNP between the Qser and Qmon parental-type groups using vcftools, retained SNPs with 0.20 < Δp < 0.65, and binned them into 0.05-wide Δp intervals. Within each Δp bin, we calculated the ICLD between pairs of trait loci whose trait association differed in the PC with the largest absolute effect (PCmax), defined from the admixture-mapping effect-size vector β, and calculated the ICLD for an equal number of randomly sampled background (non-associated) loci. ICLD was estimated as inter-chromosomal haplotype r² using vcftools --interchrom-hap-r2. We compared r² distributions between trait loci and background loci within each HI group (hybrids, Qser parental type, and Qmon parental type), and additionally compared ICLD among trait loci across the three groups. Statistical significance was evaluated using the Wilcoxon rank-sum test.

Additionally, to test whether trait loci showed signatures of divergent selection, we compared parental-type genetic differentiation and SNP-wise local cline slopes between the trait loci and the genomic background. For the differentiation-based comparison, we used F_34_ and dXY estimated in 10-kb windows in the selection scan analyses, contrasted the distributions of windows overlapping trait loci with those of non-overlapping background windows, and evaluated significance using Wilcoxon rank-sum tests. For local-cline comparison, we used SNP set A and extracted ancestry-informative markers (AIMs; 42k SNPs) under the same criteria as in the cline analysis, except that no SNP thinning was applied. For each SNP, we estimated the median and 95% credible intervals (lower and upper bounds) of the slope parameter *v* using bgc-hm (Gompert et al., 2024). We then evaluated whether these slope summaries differed between the trait loci and background SNPs using Wilcoxon rank-sum tests, similar to the differentiation-based comparisons. Finally, to quantify the magnitude of the difference non-parametrically, we performed a bootstrap-style resampling: we drew background SNP sets (matched in size to the trait-locus set) with replacement 5,000 times, and for each replicate computed the probability that the trait loci exceeded the background in each metric, Pr(diff > 0).

## Results

### Climatic niche differentiation and co-occurrence of the two *Quercus* species

Clear differentiation in climatic responses was detected between *Q. mongolica* var. *crispula* (Qmon) and *Q. serrata* subsp. *serrata* (Qser), although the two species co-occurred locally at a subset of sites. Of the 16,169 sites included in the analysis, Qmon occurred at 2,544 sites (15.7%), whereas Qser occurred at 2,971 sites (18.4%). The two species co-occurred within the same 0.1-ha plot at 460 sites, corresponding to 9.1% of the 5,055 sites at which at least one species was present. The spatial joint species distribution model showed generally satisfactory convergence (Figure S2; maximum *R*^^^<1.014; minimum ESS >357.8).

The climatic response coefficients of Qmon and Qser differed for at least one of the linear or quadratic terms for all four climatic variables: annual mean temperature (AMT), annual precipitation (APCP), maximum snow depth (SD), and relative humidity (RH) (Figures 1E, S3; Table S3). The 95% Bayesian credible intervals for the differences between the species-specific coefficients excluded zero for the linear (Δ*β*=-4.86±0.14) and quadratic (Δ*β*=0.28±0.12) terms of AMT, the linear (Δ*β*=0.62±0.07) and quadratic (Δ*β*=0.14±0.02) terms of APCP, the linear term of SD (Δ*β*=0.67±0.06), and the quadratic term of RH (Δ*β*=-0.15±0.01). In contrast, the 95% credible intervals for the quadratic SD term and the linear RH term included zero. Qmon showed higher occurrence probabilities under cooler climatic conditions and at greater snow depths than Qser.

The major climatic range represented by the co-occurrence sites encompassed most of the climatic niche space shared by the two species, despite their overall climatic differentiation. Specifically, 96.5% of the shared 95% climatic niche region of Qmon and Qser was contained within the 95% highest-density region of the co-occurrence sites. PCA-env analysis indicated partial overlap between their realized climatic niches, with a Schoener’s *D* of 0.31 (Figure 1F). The distance between their niche centroids was 1.61 PCA-score units, indicating displacement of the climatic niche centres of the two species.

### Genetic population structures and hybridisation dynamics

Genetic PCA and ADMIXTURE revealed apparent genetic differentiation between Qmon and *Q. serrata*subsp. *serrata* (Qser), as well as the presence of hybrids (Figures 2A, C). In the PCA, PC1 (17.6% of the variance explained) separated the Qmon populations (including UR) on the positive axis from the Qser populations, including NG and KG, on the negative axis, with a limited number of individuals occupying intermediate positions, consistent with the admixture (Figures 2A, S4; Table S4). ADMIXTURE supported the same pattern with K = 2, resulting in the lowest cross-validation error (Figure S5). Based on the assignments of the allopatric parental-reference populations, Cluster 1 corresponded to Qser and Cluster 2 corresponded to Qmon (Figure 2C).

**Figure 2.**
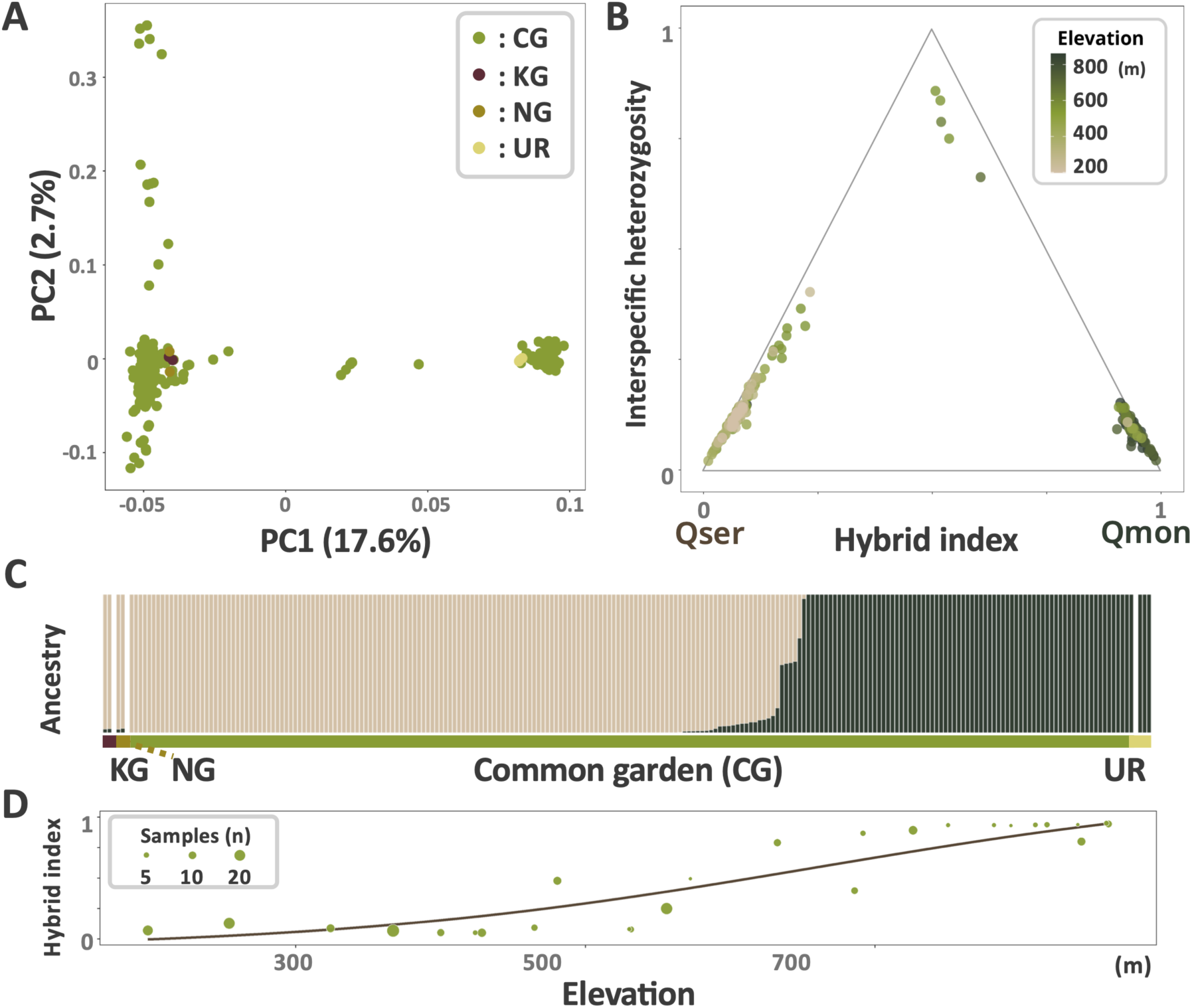
Genetic structure of individuals from common-garden and pure-line populations. (A) Genomic PCA (PC1–PC2). Point colors indicate sampling origin (CG, KG, NG, and UR). (B) Relationship between hybrid index (HI) and interspecific heterozygosity (Het) for CG individuals; point color represents acorn sampling elevation. (C) ADMIXTURE results at the optimal K = 2. (D) Fitted genomic clines (brown) with 95% confidence intervals (light gray); point size scales with the number of samples per site.

Hybrid-class inference indicated that hybridisation persisted across generations (Figure 2B). Using triangulaR, we assigned individuals to hybrid classes based on the relationship between the hybrid index (HI) and interspecific heterozygosity (Het), based on the criteria of Milne and Abbott (2008). Specifically, we classified two individuals as F1 (0.25 ≤ HI ≤ 0.75 and Het ≥ 0.85), three as F2 or later-generation hybrids (0.25 ≤ HI ≤ 0.75 and Het < 0.85), 147 as backcrosses toward Qser (0.025 ≤ HI < 0.25), 71 as backcrosses toward Qmon (0.75 < HI ≤ 0.975), four as Qser parental-type (HI < 0.025), and seven as Qmon parental-type (HI > 0.975).

Cline analyses supported an elevational genetic cline across sampling sites (Figures 2D, S6, and S7). In both the site- and individual-based analyses, models parameterised by elevation were more strongly supported than those based on horizontal distance (site-based: ΔAICc = 919; individual-based: ΔLOOIC = 56.6; Figure S6; Table S5). Estimates of cline centre and width differed between analytical units: the site-based analysis resulted in a centre of 652.9–668.3 m and a width of 646.2–729.0 m, whereas the individual-based analysis produced a centre of 555.4–605.1 m and a width of 154.9–209.8 m.

### Relationship between seedling vigor and hybrid index

Germination rate, two-season survival, and total dry weight showed no significant relationships with HI (all P-values > 0.21, Deviance explained < 0.04; Figures 3, S8; Table S6). Although genotypes could not be obtained for acorns that did not germinate, auxiliary analysis using elevation as a proxy for HI revealed no apparent relationships, indicating that the effect of HI missingness on our main conclusions was limited. Specifically, although elevation and HI were strongly correlated (Deviance explained = 0.66 or 0.88, P < 0.01; Figures S8A, B), germination and survival rates across eight elevational bins varied within ranges of 0.67–0.87 and 0.67–0.86, respectively, and did not support a monotonic trend (max difference ≈ 0.20; Figure S8C). Similarly, the association between total dry weight and HI was very weak (Deviance explained = 0.11, P = 0.82; Figure 3B; Table S6), providing no statistical evidence for a causal relationship between HI and biomass under common garden conditions. In contrast, leaf gas exchange measurements indicated a small positive effect of HI on stomatal conductance (*g_s_*_w_) as estimated using LI-600 (P = 0.042; Figure S8G; Table S6). Photosynthetic rate (*A_Ca_*_420_) measured with the LI-6800 also showed a positive estimated effect of HI, but this effect was not statistically supported (P = 0.33; Figure S8H).

**Figure 3.**
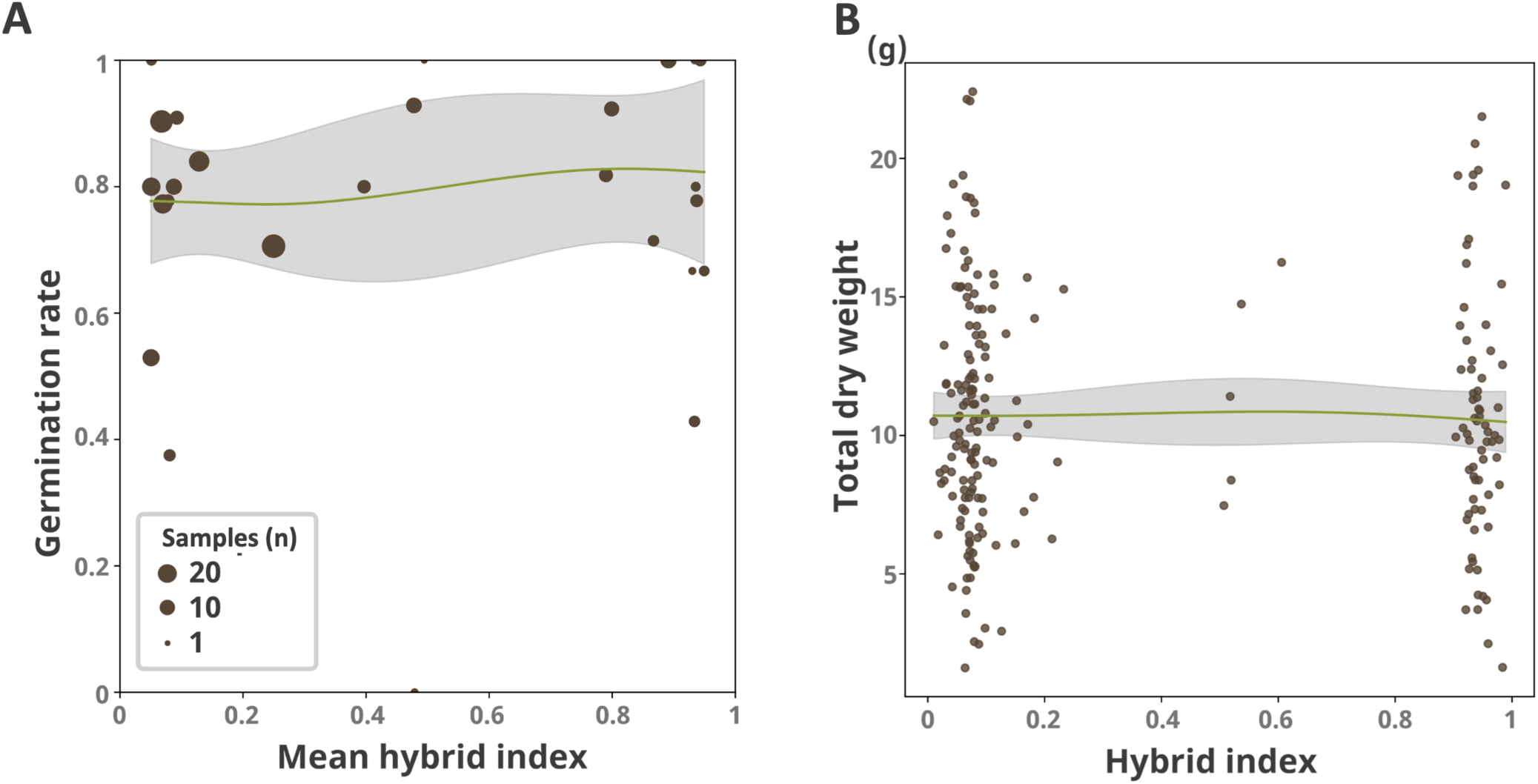
Relationships between the hybrid index and germination, survival, and biomass. (A) Site-level mean HI vs. germination rate. The light-green curve shows the GAM fit with 95% CI in light gray; point size indicates sample size per site. (B) Individual-level HI vs. total dry weight (DWtot). The light-green curve shows the GAM fit with 95% CI in light gray.

### Trait covariance structure and resource allocation trade-offs

Multiple analyses supported substantial trait covariation and the presence of trade-offs. In the correlation matrix, 53.8% (175/325) of all pairwise correlations among the 26 traits were significant (Figure S9). In the PCA, the cumulative variance explained exceeded 50% by PC4 and 80% by PC10 (Table S7). Based on trait loadings, PC1 primarily captured variation in leaf functional traits, plant architecture, and biomass (TLN, CA, Ht, Dia, DWleaf, DWstem, DWroot), whereas PC2 and PC3 reflected covariation between leaf functional traits and relative growth rate (PC2: LI, RGRtln1st, RGRca1st, RGRd2nd, RGRht1st; PC3: ILA, LI, RGRca2nd, RGRd1st, RGRd2nd, RGRht2nd). PC4 was dominated by plant architecture and relative growth rate (Dia, Ht, RGRtln1st, RGRtln2nd, RGRca1st, RGRht1st; Figures 4A, S10; Table S8). In GAMs linking PC scores to covariates, HI was significantly associated with PC2 and PC3 (adjP < 0.01), whereas total dry weight was significantly associated with PC1–PC9 (P < 0.01; Table S7). From the opposing signs of loadings on the HI-associated PCs (PC2 and PC3), we identified candidate trade-off pairs within trait categories. Within leaf functional traits, ILA and LI contrasted with TLN and YM (opposite-signed loadings), and within plant architecture, Dia contrasted with Ht. Moreover, the loading structures of PC1 and PC4 suggested a seasonal shift in allocation, as first-season relative growth rates (RGRtln1st, RGRca1st, RGRht1st) contrasted with second-season relative growth rates (RGRtln2nd, RGRca2nd, RGRht2nd).

**Figure 4.**
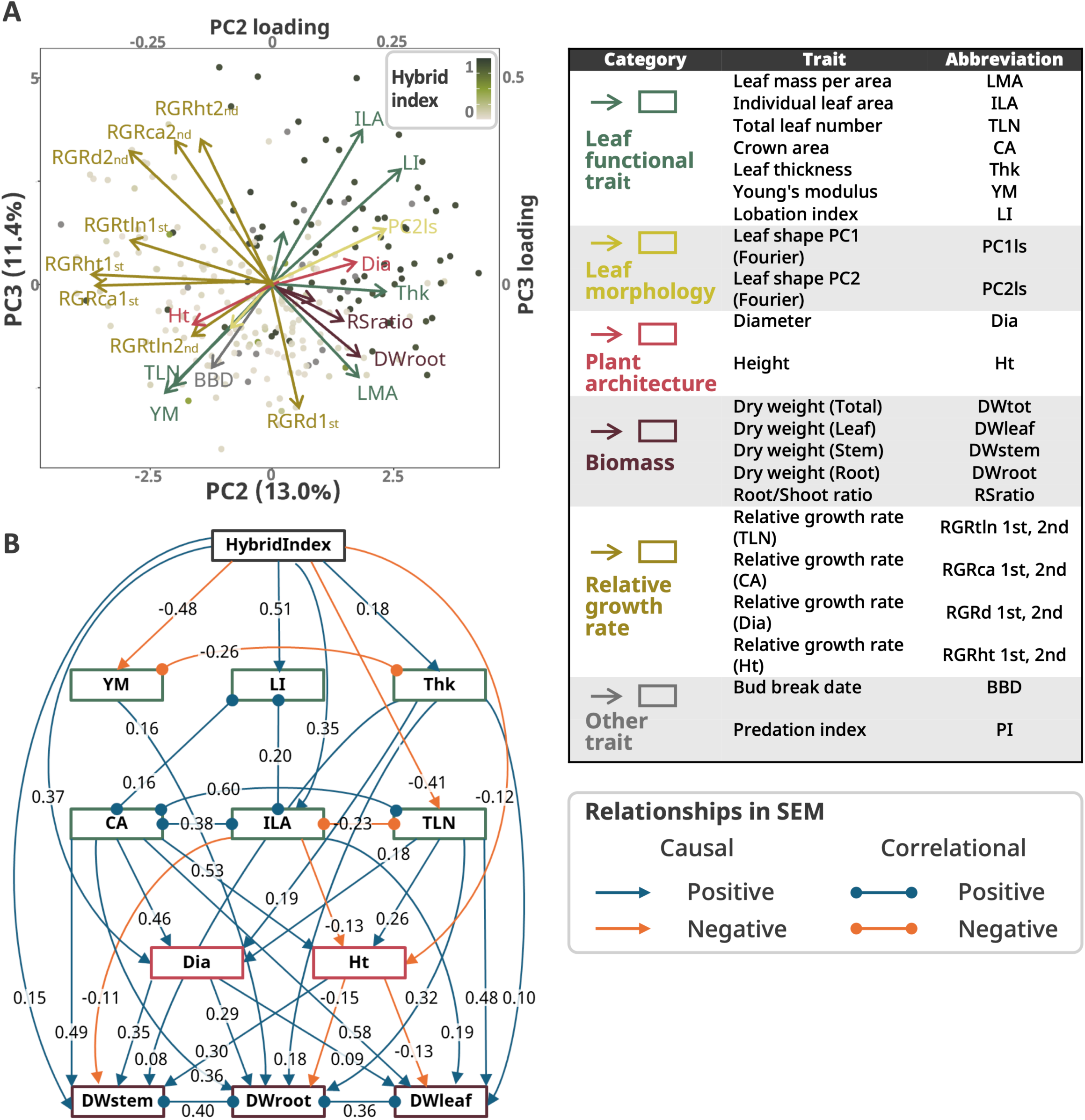
Covariance structure and trait trade-offs in seedlings. (A) PCA scores and loadings for the 26 traits. PCs significantly associated with HI (PC2 and PC3) are shown. Point color indicates individual HI; loading colors indicate pre-defined trait categories (see legend table at right). (B) Structural equation model (SEM) based on selected traits. Arrows indicate putative causal effects; undirected links represent correlations. Blue indicates positive effects, and orange indicates negative effects. Numbers on paths are standardized estimates (partial effects). Tile borders indicate trait categories.

Furthermore, SEM supported two within-category trade-offs [leaf thickness (Thk)–Young’s modulus (YM) and individual leaf area (ILA)–total leaf number (TLN); Figure 4B; Table S9]. In both pairs, HI had opposite-signed effects on the two traits, and the traits were negatively correlated. Specifically, HI had a positive effect on Thk (+0.18) but a negative effect on YM (−0.48), and Thk and YM were negatively correlated (−0.26). Likewise, HI positively affected ILA (+0.35) but negatively affected TLN (−0.41), with a negative correlation between ILA and TLN (−0.23). For stem diameter (Dia) and height (Ht), a negative correlation was not detected; nonetheless, HI showed a positive effect on Dia (+0.37) and a negative effect on Ht (−0.12), and both traits contributed positively to stem dry weight (Dia → DWstem: +0.08; Ht → DWstem: +0.30). This pattern is consistent with an allocation-mediated trade-off and aligns with the PCA loading structure. Collectively, these results suggest that allocation strategies shift along the HI gradient, including leaf mechanical investment versus leaf quantity and stem thickening versus height growth. The final SEM fit the data well (Fisher’s C = 53.3, P = 0.43; χ² = 25.9, P = 0.47), and all d-separation tests were non-significant (P > 0.07), supporting model consistency (Figure 4D; Table S9).

### Trait association genes and their functional implication

Admixture mapping identified 1,795 SNPs significantly associated with seedling phenotypic trait PCs (PC1–4; adjP < 0.05; Figure 5A; Table S10). We defined these SNPs as trait loci for subsequent gene-coupling analyses. Based on LD decay analyses, in which r² remained above 0.1 up to approximately 29.0 kb in Qmon parental-type individuals and 25.2 kb in Qser parental-type individuals, we conservatively used a 25-kb window and extracted 626 candidate genes located within ± 25 kb of the loci (nearest gene per SNP; Figure S11; Table S11). In parallel, genome-wide selection scans based on five metrics (XP-EHH, *F_st_*, dXY, π, and Tajima’s *D*) identified genes near outlier regions, yielding 2,821 genes for XP-EHH, 203 for *F_st_*, 119 for dXY, 152 for π, and 302 for Tajima’s *D* (Figure 5A). Intersecting the admixture-mapping candidates with the selection-scan outliers yielded 148 genes located near trait loci and flagged as outliers by at least one metric (Figure 5B). Of these, 101 and 111 had annotated homologs in *Q. lobata* and *A. thaliana*, respectively. Gene set enrichment analyses conducted separately using each reference annotation, revealed four significant pathways in *Q. lobata* and five in *A. thaliana* (adjP < 0.05; Figure 5C; Table S12). Notably, GO:0035251 (“UDP-glucosyltransferase activity”) and GO:0046527 (“Glucosyltransferase activity”) were significantly enriched in both analyses, suggesting glycosylation-related secondary metabolism as a potential contributor to resource-allocation variation.

**Figure 5.**
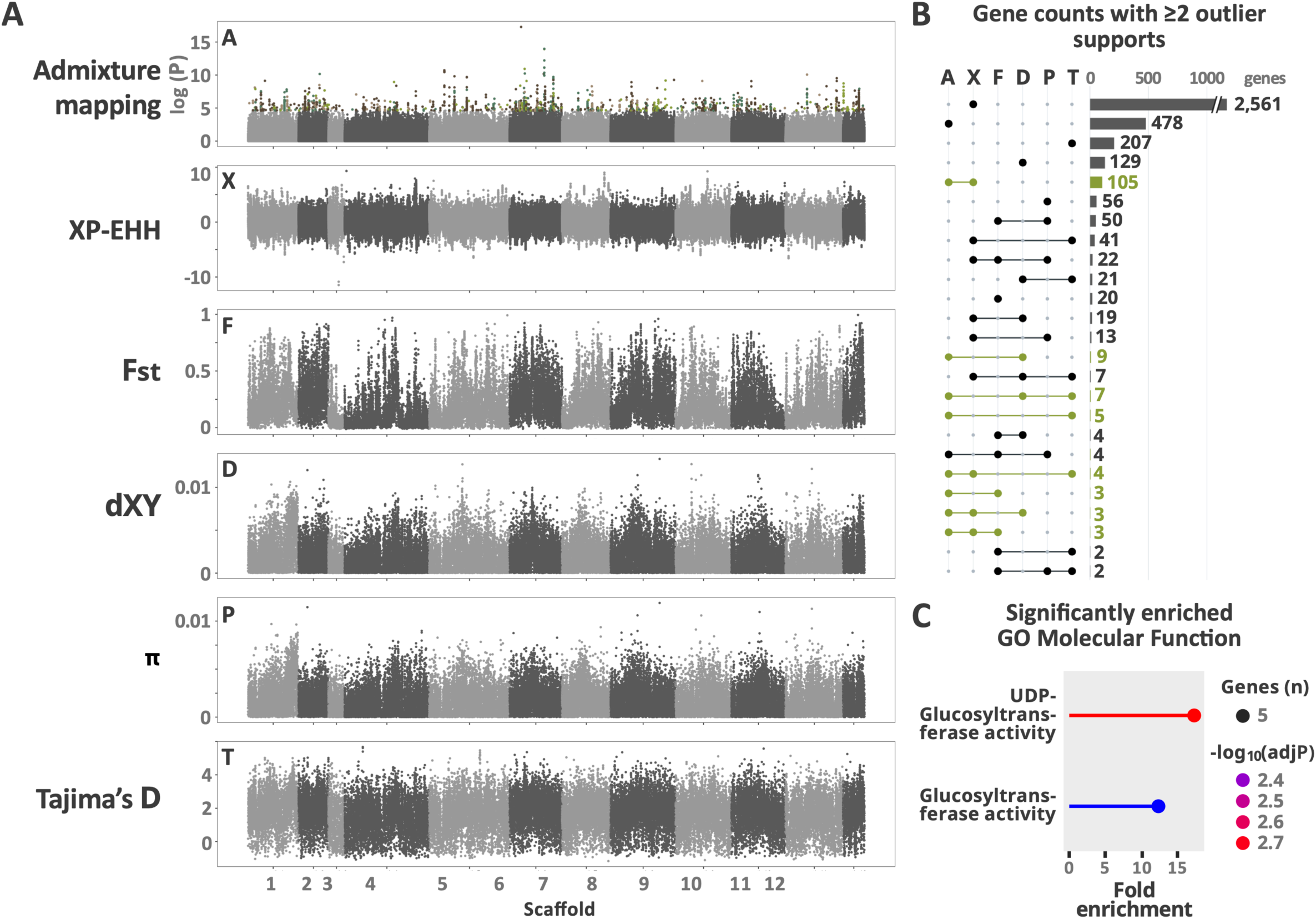
Trait-associated loci and genome scans for selection. (A) Manhattan plots. Top: admixture mapping for PC1–PC4 using multivariate tests (significance threshold adjP < 0.05, Benjamini–Hochberg). Bottom: XP-EHH, F_ST_, dXY, π, and Tajima’s D scans. The x-axis shows genomic coordinates. (B) Upset plot showing counts of genes identified as outliers by two or more metrics. Labels in light green indicate combinations that include admixture mapping and at least one selection metric. (C) Lollipop plot of gene-set enrichment results based on *Q. lobata* annotation (adjP < 0.05).

### Genetic coupling among trait loci and divergent selection

Among trait-associated regions (trait loci), hybrids (0.07 ≤ HI ≤ 0.93) showed pronounced gene coupling relative to the background. Except for the lowest interspecific allele-frequency difference bin (Δp = 0.20–0.25), inter-chromosomal r² between trait loci in hybrids consistently exceeded background levels (Wilcoxon’s P < 0.05; Figure 6; Table S13), with a maximum median difference of 0.11 in the Δp = 0.45–0.50 bin. In the parental groups, significant differences were observed in certain bins; however, the effect sizes were negligible (median difference < 0.006 for all cases). Hybrids also exhibited higher r² between trait loci than either parental group; for Δp > 0.35, all median differences exceeded 0.05 and were significant (Wilcoxon’s P < 0.05; Table S14). Together, these results indicate a hybrid-specific elevation in inter-chromosomal linkage disequilibrium (ICLD), which is consistent with genetic coupling among trait loci.

**Figure 6.**
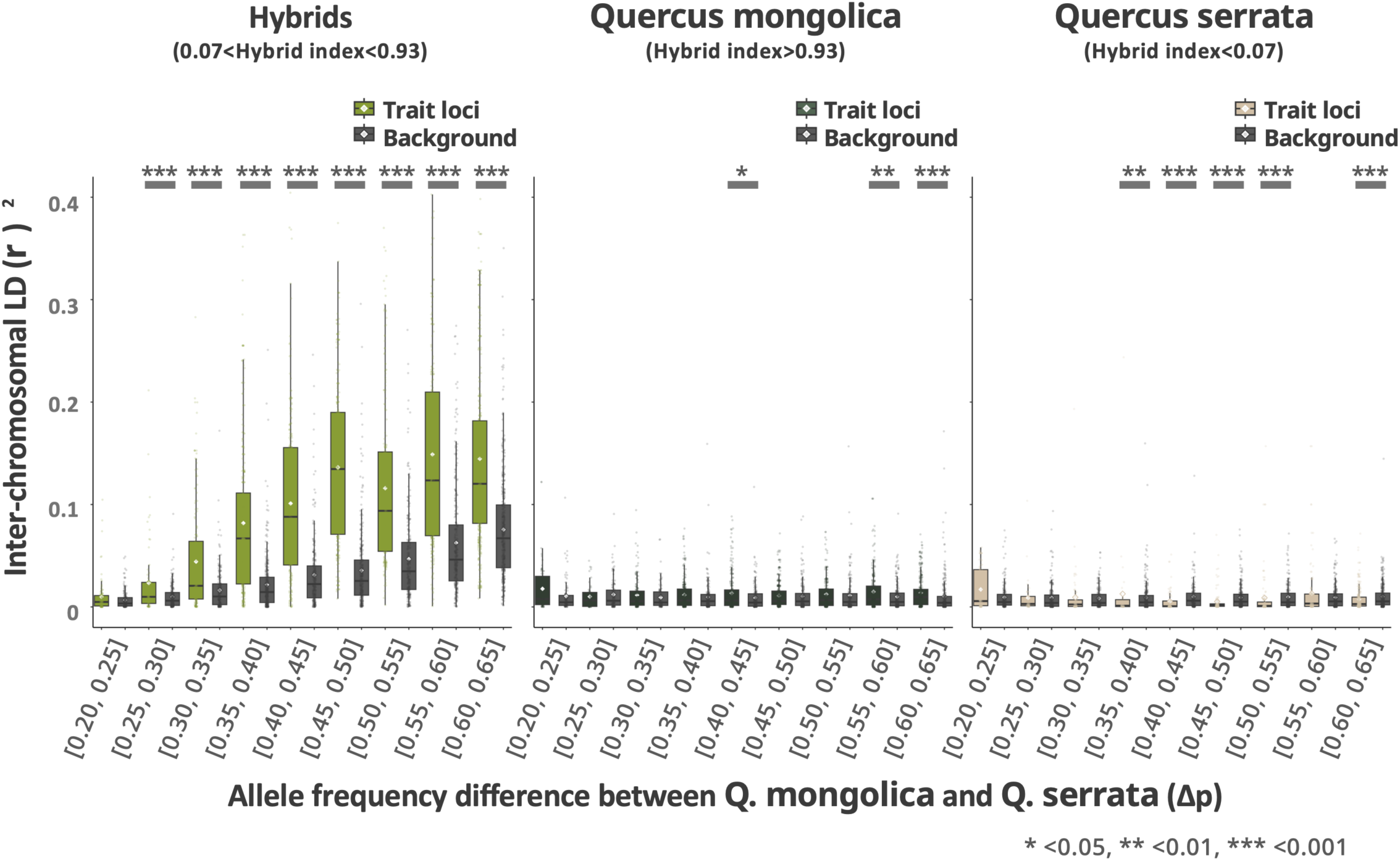
Inter-chromosomal linkage disequilibrium (ICLD) among trait loci. Comparison of ICLD (haplotype r², vcftools --interchrom-hap-r2) between trait loci and background in Hybrid (0.07 < HI < 0.93), Qmon parental (HI ≥ 0.93), and Qser parental (HI ≤ 0.07) subsets. The x-axis shows bins of parental allele frequency difference Δp (width 0.05); the y-axis shows r². Distributions are shown for each Δp bin; statistical tests are reported in the main text.

Comparisons of genetic differentiation between parental types and local cline slopes in hybrids consistently indicated that trait loci exceeded background levels, supporting divergent selection. *F_ST_* and dXY, estimated in 10-kb windows, were significantly higher for windows containing trait loci than for background windows (Wilcoxon’s P < 0.05; Figure S12). In the local cline analysis with bgc-hm, trait loci also exhibited larger slope estimates (v) than those for background loci, as indicated in the median and lower and upper bounds of the 95% credible interval (Figure S13; Table S15). Although the median difference was not apparently significant, the probabilities that the trait loci exceeded the background were high for both credit interval bounds [Pr (diff > 0) = 0.99 and 1.00]. Collectively, these results suggest that divergent selection at trait loci contributes to the formation and maintenance of gene coupling.

## Discussion

This study demonstrated that natural selection can maintain a growth-survival trade-off and its underlying genomic architecture despite ongoing gene flow in a hybrid zone between *Quercus mongolica* var. *crispula* (Qmon) and *Q. serrata* subsp. *serrata* (Qser). Our findings provide an evolutionary framework for understanding how functional differentiation can be maintained between species despite genetic exchange, with important implications for the maintenance of functional diversity and local species coexistence in tree communities.

### Trait trade-offs and their adaptive evolutionary context

Analyses of the 26 traits revealed a structured pattern of covariation among the seedling traits, consistent with multiple trade-offs. In the trait–trait correlation matrix, significant correlations were identified for more than half of all pairwise combinations, indicating pervasive covariation among the traits (Figure S9). Trait PCA combined with GAM-based association analyses further suggested that genetic background, summarised by the hybrid index (HI), was associated with multivariate phenotypic variation (Figures 4A, S10). Using structural equation modelling (SEM), we identified two trade-offs within leaf functional traits, represented by negative associations between leaf thickness (Thk) and Young’s modulus (YM) and between individual leaf area (ILA) and total leaf number (TLN). We also identified an intrinsic trade-off in plant architecture between stem diameter (Dia) and height (Ht), mediated by their shared positive effects on stem dry weight (DWstem). Together, these results support three coordinated trait trade-offs (Figure 4A, B).

The trade-off relationships inferred from SEM are broadly consistent with previous comparative studies across tree species and likely indicate the ecology and habitats of both Qmon and Qser. For leaf thickness (Thk) and Young’s modulus (YM), interspecific analyses have reported positive associations between tearing resistance and Thk, whereas YM is primarily determined by lamina density and cellulose density and is mainly independent of Thk (Onoda et al., 2011; Méndez-Alonzo et al., 2013). Therefore, investment in thickness and stiffness can enhance leaf mechanical performance; however, within species and among closely related taxa under common garden conditions, allocation constraints may generate an apparent trade-off between these two strategies.

In addition, trade-offs between ILA and TLN have been documented across and within species at individual and branch scales (Yang et al., 2008; Dombroskie & Aarssen, 2012). As ILA are linked to numerous environmental pressures, including herbivory, photosynthetic efficiency, leaf temperature regulation, and plant water status (Moles & Westoby, 2000), shifts along the ILA–TLN axis can indicate broader ecological strategies. In Qser, which occupies warmer habitats than Qmon and frequently experiences strong summer water stress, an increased investment in TLN coupled with a reduced ILA is consistent with the contrasting environments of the two species (Figure 1C). Along the same physiological axis, we also observed a weak positive association between HI and stomatal conductance (gsw; Figure S8). However, we considered this result as ancillary and did not provide further interpretation.

The Dia–Ht trade-off observed in this study has also been reported across broad taxonomic groups (Trouvé et al., 2015). Initial increases in seedling height can result in long-term advantages in light competition (Van Couwenberghe et al., 2013), whereas increased stem diameter enhances mechanical stability under wind and snow loads, while increased height may compromise this structural integrity (Peltola et al., 1999). In our system, Qser invested substantially in height growth, consistent with its broader overlap with evergreen oaks such as *Q. glauca* and *Q. myrsinifolia,* and the likelihood of stronger light competition. In contrast, Qmon’s increased investment in stem diameter likely indicates the snowy and wind conditions that characterise higher elevations and latitudes, positioning this axis as a competition–survival trade-off.

In the SEM, we also identified positive paths linking traits that belonged to different trade-offs (Thk → Dia = 0.19; TLN → Ht = 0.26). Seedlings with thicker and heavier leaves may require an increased investment in stem diameter to maintain mechanical support (Yang et al., 2008), whereas seedlings with more leaves may require increased height to acquire space and light (Van Couwenberghe et al., 2013). These cross-links indicate integrated trait syndromes and are consistent with studies on coordinated leaf-trait variation along an acquisitive–conservative resource-allocation continuum (Gorné et al., 2022). The directions of these negative and positive trait relationships were consistent with the corresponding PCA loadings, indicating that the PC axes captured multivariate trait covariation, including the trade-offs identified by SEM. Thus, loci associated with these PCs can be interpreted as genomic regions associated with coordinated trait variation rather than with individual traits in isolation. More broadly, the trade-offs and covariation patterns identified in the present study are consistent with the physiological and ecological expectations of seedlings facing contrasting environments.

### Detection of natural selection on resource allocation trade-offs

Within the framework of selection–gene flow antagonism, we evaluated whether natural selection maintains resource allocation trade-offs through gene coupling using three complementary lines of evidence: (i) evidence that the hybrid-zone cline is at or near equilibrium, (ii) robust identification of trait-associated regions, and (iii) demonstration of elevated inter-chromosomal linkage disequilibrium (ICLD) among these regions.

Genomic analyses of individuals sampled across the hybrid zone suggested that under continuous gene flow, the zone is close to equilibrium following secondary contact. The joint distribution of HI and interspecific heterozygosity indicated a continuous representation of hybrid classes, from F1 individuals to multigenerational backcrosses (Figure 2B). The zone covers only approximately 2.5 km horizontally, and oaks (*Quercus*) are wind-pollinated, with pollen dispersing across distances ranging from hundreds of meters to several kilometres (Ashley, 2021). Independent phylogenetic and population genetic evidence further supports secondary contact between Qmon and Qser (Hipp et al., 2020; Aizawa et al., 2021). These observations indicate that hybridisation persists sufficiently for the cline to approach equilibrium.

In cline analyses, the elevation model was consistently a better fit than the distance model, suggesting habitat segregation driven by elevation-associated environmental gradients such as temperature, snow depth, wind exposure, and vegetation (Figures 2D, S6, and S7). In contrast, in the common garden, which was warm, snow-free, and environmentally homogeneous, we observed no apparent differences in germination, two-year survival, or dry weight among Qmon, Qser, and the hybrids. This suggests that pronounced differences in the fitness are unlikely under these conditions. Accordingly, we observed no evidence of lethality or sterility at the seedling stage attributable to late-generation hybrid breakdown or Bateson–Dobzhansky–Muller incompatibilities (BDMIs), at least in the common garden environment. The low frequency of hybrids indicates that stronger barriers act before fertilisation, whereas incompatibilities during germination and early seedling establishment appear weak.

Next, by integrating admixture mapping with selection scans, we identified a set of trait-associated loci (trait loci) that were plausible in both functional and evolutionary terms (Figure 5A,B). GO enrichment analyses using both *Q. lobata* and *Arabidopsis thaliana* annotations consistently identified UDP-glucosyltransferase activity and glucosyltransferase activity, involving five candidate genes (Figure 5C). These functions point to UDP-glucose-related metabolic processes, in which glycosylation regulates the activity, stability, and transport of diverse metabolites and can thereby influence growth, defence, and resource allocation (Yonekura-Sakakibara et al., 2019; Pastorczyk-Szlenkier & Bednarek, 2021). Notably, one of the five candidate genes associated with these enriched functions was TPS1, which encodes trehalose-6-phosphate synthase 1. TPS1 catalyses the synthesis of trehalose-6-phosphate (T6P), a central sucrose-signalling metabolite that integrates carbon availability with plant growth and physiology. T6P interacts with major hormonal pathways, including auxin, strigolactone, and abscisic acid signalling, and this regulatory network has been proposed to mediate a productivity–survival trade-off that evolved under natural environmental constraints (Fichtner et al., 2020; Paul et al., 2026). Other candidates included CalS7 (AT1G06490), which is required for normal phloem transport and growth, and CSLG1 (AT4G24010), a cellulose synthase-like glycosyltransferase associated with polysaccharide biosynthesis. The recovery of functionally coherent candidates provides biological support for our trait-locus set, indicating that even for plastic growth-related seedling traits, common-garden homogenisation and reduced covariate noise in admixture mapping can result in robust genotype–phenotype signals (Donovan et al., 2011).

Additionally, the hybrid-specific elevation of ICLD provided key support for gene coupling among trait loci (Figure 6). Under selection–gene flow antagonism, interspecific hybridisation is expected to generate a transient pulse of linkage disequilibrium (LD), which is eroded through recombination and repeated backcrossing after secondary contact. If selection promotes particular multilocus allele combinations and is strongly related to recombination, LD decay can be slowed or prevented, enabling elevated LD to persist over extended timescales (Schield et al., 2024). Importantly, this coupling does not rely on physical linkages within chromosomes; it can also be expressed as LD among loci on different chromosomes.

These patterns cannot be explained by a simple habitat filtering alone, in which individual traits are independently targeted by selection. Such a scenario does not readily explain the excess inter-chromosomal LD observed among trait loci relative to the genomic background within hybrid individuals. Nor is the excess ICLD readily explained by reproductive-ecological biases alone. *Quercus* is wind-pollinated, making assortative mating through mate choice or pollinator-mediated preferences unlikely. In addition, there appears to be ample opportunity for local competitive interactions between the two species. Although Qmon and Qser showed clearly differentiated responses to climatic conditions, their realised climatic niches overlapped broadly. Moreover, the two species repeatedly co-occurred within the same 0.1-ha plots across much of this overlapping niche space. Thus, although climate-driven habitat filtering contributes to their broad-scale distributional differences, it does not result in complete spatial segregation at the local scale. Considering the crown area occupied by mature individuals of both species, such local co-occurrence indicates spatial conditions in which interspecific competition for resources, including light, could readily occur. Therefore, we interpreted the excess ICLD among trait loci as evidence consistent with gene coupling driven by natural selection on growth-survival trade-offs and, more broadly, on integrated structural and functional trait syndromes in seedlings. Notably, the magnitude of ICLD in our data was comparable to that reported in systems inferred to experience strong selection, such as sexual selection in a swallow species complex (Schield et al., 2024), suggesting that resource limitation and trait integration at the seedling stage can impose substantial selective pressures.

### Conclusions and perspectives

Our study showed that, even between closely related species experiencing ongoing genetic exchange, natural selection acting on integrated trait syndromes can contribute to maintaining functional trait differentiation, particularly the growth–survival trade-off, and its underlying genomic architecture. Such functional trade-offs and trait syndromes have long been considered to promote niche differentiation through differences in resource use and environmental responses, thereby contributing to species coexistence in plant communities. Indeed, although Qmon and Qser showed clearly differentiated climatic responses, their realised climatic niches overlapped broadly, and local co-occurrence within the same 0.1-ha plots was widely observed across this shared climatic space. Thus, natural selection that maintains functional differentiation despite gene flow may preserve ecological differences that contribute to local coexistence and, more broadly, may represent one evolutionary process supporting the high functional and species diversity of forest communities.

However, practical constraints prevented us from testing whether the trait associations and selective pressures identified at the seedling stage persisted into later ontogenetic stages and mature trees. We also did not directly quantify survival or growth responses under alternative stress conditions such as snow loading or low-light conditions. In addition, although our modelling framework reduced the influence of fine-scale genetic structures within the hybrid zone and environmental covariates, these sources of variation cannot be entirely eliminated. Moreover, although local co-occurrence demonstrates opportunities for interspecific interactions, we did not directly quantify competitive interactions or demographic processes required to establish stable coexistence. These limitations could be addressed by common garden experiments spanning contrasting environments, and by extension and replication across additional lineages and regions.

Our results suggest that the ability of natural selection to maintain integrated functional trait syndromes, including the growth–survival trade-off and their underlying genomic architecture, despite ongoing gene flow has implications for several evolutionary questions. First, it suggests that long-term, syngameon-like hybridisation in *Quercus*, and more broadly the high incidence of interspecific hybridisation in trees, may persist without eliminating species differentiation. Local selection may counteract homogenisation under ongoing gene flow, allowing differentiation to be maintained without relying solely on geographic isolation followed by secondary contact. Thus, selection on integrated structural and functional trait syndromes can act as an evolutionary barrier to homogenisation. Second, our results indicate that mechanisms maintaining diversity in tree communities should be considered in conventional ecological terms, such as habitat filtering and niche differentiation through resource partitioning, as well as be understood as molecular-evolutionary dynamics influenced by selection and gene coupling. Overall, this study provides testable molecular hypotheses that link the maintenance of coexistence to the mechanisms of adaptive evolution in tree communities.

## Acknowledgement

We are grateful to Victoria Sork, Michimasa Yamasaki and Keita Ido for advice on study design and data analyses. We thank Yumeko Tarusawa, Chikashi Hata, Mayu Katafuchi, Keigo Sakano, Anju Furuta and Yoshiyuki Suzue for assistance with trait measurements, and members of the Tropical Forest Resources and Environment Lab for help with sample collection. We thank the Uryu Experimental Forest, Field Science Center for Northern Biosphere, Hokkaido University, for permitting partial sample collection. This work was supported by JSPS KAKENHI (grant number 21H05314). We thank Editage for editorial assistance.

## Data availability statement

Raw sequence data have been deposited in the DNA Data Bank of Japan (DDBJ) under BioProject accession PRJDB40283. The code and scripts used for analyses are available in the GitHub: https://github.com/ryosuke110/Quercus_comgarden. Raw trait data and genetic data have been deposited in Dryad (DOI: 10.5061/dryad.4f4qrfjtd). Leaf scan images and crown area images are available from the corresponding author upon reasonable request due to their large file sizes.

## Competing interests statement

The authors declare no competing interests.

## Author contributions

Y.I., R.K.I., and Y.O. conceived and designed the study. R.K.I., K.K., K.O., and S.S. collected the data. R.K.I. and K.K. designed the analytical workflow and performed data analyses. R.K.I. wrote the manuscript. all authors revised the manuscript and approved the final version.

---

**Figure S1. Initial model of the SEM analysis.**

**Figure S2. Trace plots in spOccupancy analysis.** Trace plots of posterior MCMC samples for species-specific climatic-response coefficients estimated for Qmon and Qser.

**Figure S3. Results of the spOccupancy analysis.** (A) Comparison of standardized climatic variables, including annual mean temperature (AMT), annual precipitation (APCP), maximum snow depth (SD), and relative humidity (RH), between Qmon and Qser occurrence sites. (B) Geographic distribution of Qmon (dark green) and Qser (beige) occurrence sites. Sites where both species co-occurred are shown as light-green circles.

**Figure S4. Details of genomic PCA. (A) PC1–PC2, (B) PC1–PC3, (C) PC1–PC4, (D) PC1–PC5.**

**Figure S5. Details of ADMIXTURE analyses. (A) K = 2–4 bar plots, (B) cross-validation error for K = 1–12.**

**Figure S6. Site-based clines using hzar. (A) Elevation and (B) distance models.**

**Figure S7. Individual-based clines using PyMC. (A) Elevation model, (B) Distance model, and (C) null model with a constant mean HI.** Model comparisons are shown.

**Figure S8. Elevation, hybrid index (HI), early performance, and leaf gas exchange.** (A) Individual elevation versus HI. (B) Site elevation versus site-mean HI. (C) Germination and two-season survival across eight equal-elevation bins (83 m each). (D) Site-mean HI versus germination. (E) Site-mean HI versus survival. (F) Individual HI versus total dry weight (DWtot). (G) Stomatal conductance to water vapour, *g*_sw_ (mol *m*^−2^ *s*^−1^), versus HI. (H) Net *CO*_2_ assimilation at *C_a_* = 420 ppm, *A_area_* (μmol *m*^−2^ *s*^−1^), versus HI.

**Figure S9. Pairwise correlation matrix of the 26 traits with hierarchical clustering dendrogram.**

**Figure S10. PCA details. Significant (A) PC2 and (B) PC3 vs. HI, and non-significant (C) PC1 and (D) PC4. Point colors: (A, C) HI; (B, D) DWtot. Loading colors indicate trait categories.**

**Figure S11. Linkage disequilibrium decay in the parental-type groups (Qmon and Qser).**

**Figure S12. Parental genetic divergence (F_ST_, dXY) for 10 kb windows containing trait loci (light green) vs. background windows (gray).** Asterisks indicate significance (* P < 0.05, ** P < 0.01, *** P < 0.001).

**Figure S13. Local cline slope (v) comparison: median and 95% CI (lower, upper) for trait loci (light green) vs. background (gray).** (A) Violin plots; (B) bootstrap distributions of differences. Asterisk notation as in S12.

**Table S1. Summary of read-mapping quality-control statistics used to assess potential mapping bias associated with hybrid index (HI).** Values are presented as mean ± SD across individuals. Pearson’s R represents the correlation between each mapping metric and HI. Slope and P values were obtained from linear regressions of each mapping metric against HI.

**Table S2. Summary of seedling traits. Categories, sample sizes (n), measurement timing, and definitions.**

**Table S3. Posterior estimates of climatic effects on the occurrence of *Quercus mongolica* and *Q. serrata*.** Values represent posterior means with 95% credible intervals (CRL). Δ represents the posterior difference in regression coefficients between *Q. mongolica* and *Q. serrata* (*Q. mongolica* − *Q. serrata*). AMT, annual mean temperature; APCP, annual precipitation; SD, solar radiation; RH, relative humidity. A 95% CrI excluding zero indicates strong posterior support for a non-zero effect or species difference.

**Table S4. Proportion of variance explained by principal components from the genomic PCA.** Includes individual and cumulative variance (%).

**Table S5. Estimated parameters from the cline analyses and model comparison between the Elevation and Distance models.** Site-based (hzar) results present Centre and Width (mean, 95% HDI), AICc, and model weight. Individual-based (PyMC) results present Centre, Width, κ (mean, 95% HDI), LOOIC, and model weight.

**Table S6. Summary of generalised additive models (GAMs) relating hybrid index (HI), environmental covariates, and seedling fitness– and gas-exchange–related traits.** For each model, we present the response equation, sample size (n), error distribution and link function, deviance explained, total effective degrees of freedom (EDF), P value for the overall significance of smooth terms, and the Akaike information criterion (AIC) used for smoothing-parameter selection.

**Table S7. Variance explained by phenotypic principal components and linear GAM results linking PCs to hybrid index and total dry weight.** For each PC: variance explained, EDF, F-statistics, raw and FDR-adjusted P-values for models of the form PCₙ ∼ s(HI) + f(Block) + f(Treatment) and DWtot ∼ s(PCₙ) + f(Block) + f(Treatment). (Software and versions are specified in Methods.)

**Table S8. Loadings of morphological and physiological traits on the first four phenotypic principal components.** Loadings with |value| ≥ 0.25 are highlighted.

**Table S9. Standardized path coefficients and significance for pairwise relationships among traits in the final piecewise structural equation model.** Standardized coefficients (β), 95% CI, P-values, Fisher’s C, and model fit indices are presented.

**Table S10. Admixture-mapping outliers and their effects on phenotypic principal components.** For each SNP: chromosome, position, nearest gene and distance, effect sizes (β_PC1–β_PC4), SE, raw and adjusted P-values, and AIM/Δp metadata.

**Table S11. Within-chromosomal linkage disequilibrium at candidate maximum distances used for gene assignment.**

**Table S12. Enriched biological pathways from ShinyGO analyses.** Shown separately for *Quercus lobata* and *Arabidopsis thaliana* annotations: term ID/name, gene count, gene ratio, background size, and adjusted P-values (Benjamini–Hochberg FDR, as specified in the Methods).

**Table S13. Inter-chromosomal linkage disequilibrium (ICLD) across Δp bins: summary statistics and comparisons between trait-associated loci and genomic background.** For each Δp bin: mean, median, 95% CI of r², Wilcoxon test P-values, and effect sizes such as Cliff’s delta.

**Table S14. Among-group comparisons of ICLD at trait-associated loci (Hybrid, *Q. mongolica*, *Q. serrata*).** Δp-binned pairwise tests with summary statistics, P-values, effect sizes, and sample sizes.

**Table S15. Between-group comparisons of local cline slope parameters.** Median and 95% CI of v from bgc-hm, with trait-loci vs background contrasts and bootstrap estimates of Pr(diff > 0) based on 5,000 resamples.

## Notes

https://github.com/ryosuke110/Quercus-commongarden

### Competing Interest Statement

The authors have declared no competing interest.

### Summary of Updates

This version substantially revises the manuscript to focus on how natural selection maintains the genomic architecture of divergent resource-allocation strategies despite gene flow. We added nationwide analyses of climatic niche differentiation and local cooccurrence, revised the genotype phenotype association analysis using multivariate mixed models, and expanded genomic analyses of linkage disequilibrium decay, selection, genetic clines, and excess interchromosomal linkage disequilibrium at trait associated loci. We also added assessments of hybrid classification and ancestry associated mapping performance and extensively updated the text, figures, and Supporting Information.

