## Supplemental Figures, Tables, and Supplementary Methods. for "Selection maintains genetic coupling of functional trait trade-offs despite gene flow in an oak hybrid zone"

##### Supplementary Methods 1. Climatic niche differentiation and local co-occurrence between the two *Quercus* species

To characterize climatic niche differentiation and the conditions associated with local co-occurrence between *Q. mongolica* var. *crispula* (Qmon) and *Q. serrata* subsp. *serrata* (Qser), we analyzed species-specific climatic responses, overlap between their realized climatic niches, and the climatic space occupied by sites where the two species co-occurred.

Species occurrence records were obtained from the National Forest Inventory conducted by the Forestry Agency of Japan. In this inventory, tree species occurring within 0.1-ha plots systematically distributed across Japan on an approximately 4-km grid were recorded. A focal species was classified as present (1) when at least one individual was recorded within a plot and as absent (0) when no individual was recorded. Co-occurrence was defined as the presence of both focal species within the same inventory plot. Thus, co-occurrence in this study indicates that the two species occurred within the same 0.1-ha area.

Climatic conditions at each inventory plot were obtained from the 1-km-resolution Agro-Meteorological Grid Square Data System provided by the National Agriculture and Food Research Organization. Daily climate data for 2016–2025 were retrieved using Python, summarized for each year, and then averaged across the 10-year period. The climatic variables obtained were annual mean temperature (AMT), annual maximum temperature (Tmax), annual minimum temperature (Tmin), annual precipitation (APCP), maximum snow depth (SD), annual sunshine duration (SSD), and relative humidity (RH). Eight plots with missing climatic variables or spatial coordinates were excluded, leaving 16,169 plots for analysis.

Pearson correlation coefficients and variance inflation factors (VIFs) were examined to reduce multicollinearity among the explanatory variables. Relatively strong correlations were detected between AMT and Tmin, AMT and Tmax, and SD and SSD. AMT was retained as the representative temperature variable, whereas Tmax and Tmin were excluded. SD was retained in preference to SSD because snow depth has previously been associated with the density of Q.mon (Watanabe et al. 2021). The final set of explanatory variables—AMT, APCP, SD, and RH—satisfied both $|r|<0.7$ and VIF <2.5. All climatic variables were standardized to a mean of zero and a standard deviation of one before analysis.

Spatial joint species distribution model

To evaluate climatic differentiation between the two species while accounting for spatial autocorrelation, we fitted a spatial-factor joint species distribution model (sfJSDM) using the R package spOccupancy (Doser et al. 2022). Occurrence of species *s* at site *j*, denoted by $Y_{sj}$, was modeled as a Bernoulli random variable with occurrence probability $\psi_{sj}$:

$$Y_{sj} \sim \text{Bernoulli}\left(\psi_{sj}\right).$$

Occurrence probability was related to the standardized climatic variables using the following logit-linear predictor:

$$\begin{aligned}\text{logit}\left(\psi_{sj}\right) = \beta_{0s} &+ \beta_{1s}\text{AMT}_j + \beta_{2s}\text{AMT}_j^2 \\ &+\beta_{3s}\text{APCP}_j + \beta_{4s}\text{APCP}_j^2 \\ &+\beta_{5s}\text{SD}_j + \beta_{6s}\text{SD}_j^2 \\ &+\beta_{7s}\text{RH}_j + \beta_{8s}\text{RH}_j^2 + w_s\left(\boldsymbol{s}_j\right).\end{aligned}$$

Here, $\beta_{0s}$ is the species-specific intercept, and $\beta_{1s}, \ldots, \beta_{8s}$ are the species-specific climatic-response coefficients. All climatic variables in the equation represent standardized values. The term $w_s\left(\boldsymbol{s}_j\right)$ represents the latent spatial effect for species *s* at the spatial coordinates $\boldsymbol{s}_j$ of site *j*.

The spatial effect was modeled using an exponential covariance function and a nearest-neighbor Gaussian process (NNGP), with 15 nearest neighbors and one latent spatial factor. Geographic coordinates were projected using a Lambert conformal conic projection covering Japan, and spatial distances were expressed in kilometers. Species-specific climatic coefficients were estimated hierarchically as arising from community-level normal distributions. A normal prior with a mean of 0 and a variance of 2.72 was assigned to each community-level mean coefficient. Inverse-gamma priors with both shape and scale parameters equal to 0.1 were assigned to the corresponding among-species variances.

Four MCMC chains were run for 100,000 iterations each. The first 50,000 iterations of each chain were discarded as burn-in, and every tenth sample was retained thereafter. This yielded 5,000 retained posterior samples per chain and 20,000 posterior samples in total. Convergence was assessed using trace plots, the Gelman–Rubin statistic (\($\hat{R}$\)), and effective sample size (ESS). Values of $\hat{R}$<1.1 and ESS>200 were used as convergence criteria.

To directly compare the climatic responses of the two species, differences between species-specific coefficients were calculated for every MCMC sample as

$$\Delta\beta_k = \beta_{k,\mathrm{Qmon}} - \beta_{k,\mathrm{Qser}}.$$

A climatic response was considered to differ clearly between the species when the 95% Bayesian credible interval of ($\Delta\beta_k$) excluded zero. We also calculated the posterior probability $\Pr(\Delta\beta_k > 0)$.

The realized climatic niches of the two species and the climatic conditions represented by their co-occurrence sites were evaluated using the PCA-env approach (Broennimann et al. 2012; Gaytan-Legaria et al. 2025). Principal component analysis was performed using AMT, APCP, SD, and RH across all 16,169 inventory plots, which represented the common available environmental background. Occurrence sites of Qmon and Qser and sites where the two species co-occurred were then projected onto the first two principal-component axes.

The resulting environmental space was divided into a $100 \times 100$-cell grid. Kernel density estimation implemented in the R package ecospat was used to estimate the occurrence densities of Qmon and Qser and the density of co-occurrence sites in PCA environmental space. Occurrence densities were corrected for the availability of environmental conditions. Overlap between the realized climatic niches of Qmon and Qser was quantified using Schoener's *D* (Schoener, 1968) and the modified Hellinger index *I* (Warren et al., 2008).

To characterize the climatic conditions associated with local co-occurrence, we identified the intersection of the 95% highest-density regions (HDRs) of Qmon and Qser. We then calculated the proportion of this shared area in PCA environmental space that was contained within the 95% HDR of the co-occurrence sites.

#### Supplementary Methods 2. Cultivation conditions in the common garden and phenotypic measurements

Acorns of *Quercus mongolica* var. *crispula* (Qmon) and *Q. serrata* subsp. *serrata* (Qser) were collected from the forest floor at 48 sites in a natural hybrid zone in the Hira Mountains, Shiga Prefecture, Japan in early April 2022. At each site, acorns were collected from points distributed as widely as practicable within an area of approximately 50 m radius to reduce the probability of repeatedly sampling offspring from the same maternal tree. Because the acorns were collected from the forest floor, maternal identities could not be determined. Acorns were sown in 15-cm nursery pots filled with a 1:1 mixture of commercial seedling substrate (TAKII & CO., LTD; Japan) and akadama soil, and grown outdoors at the Kitashirakawa Experimental Farm, Kyoto University (Figure 1C). Prior to sowing, acorns were treated for 1 h with a Benlate wettable powder containing benomyl at a 1:2000 dilution (KINCHO Garden Products Co., Ltd., Japan). At sowing, acorns were initially planted in groups of four per collection site per pot and were transplanted into an individual 15-cm pots approximately one week after germination. After no new germination had been observed for seven consecutive days (8 June 2022), individual pots were arranged using a randomized block design. The garden was divided into 21 spatial blocks, and seedlings were allocated among blocks so that individuals originating from different collection elevations were distributed as evenly as possible. Positions were then randomized within each block. All pots received the same measured amount of pesticide (apploximately 1 mL per pot; Benica X Spray, KINCHO Garden Products Co., Ltd., Japan) in May. Seedlings were grown without artificial shading and irrigated daily until drainage from the bottom of each pot. Liquid fertiliser (Hyponex Undiluted Solution Liquid Fertilizer, NPK 6–10–5; HYPONeX JAPAN) was applied weekly at a 1:1000 dilution, with 50 mL administered to each pot per application.

We measured or derived 25 growth-related traits that could plausibly differ between Qmon and Qser. Total leaf number (TLN), crown area (CA), stem diameter at 1 cm above the ground (Dia), and height from the ground to the apical leaf tip (Ht) were recorded at approximately two months after sowing (19, 22, and 24 June 2022), four months after sowing (18–19 August 2022), and at the end of the second growing season (CA: 20–23 August 2023; other traits: 4–7 September 2023). Crown area was quantified from top-view images of the canopy projection: iPhone SE2 images (72 dpi) were used for the two- and four-month time points, and a Nikon Z50 with a 17–50-mm zoom lens (300 dpi) was used in the second season. For each image, resolution and size were calibrated based on a reference background sheet (A4 or 530 × 550 mm) using the measurement tool in Photoshop 2023 (Adobe). For each trait ($X$), we computed relative growth rates (RGRs) for the first season (Jun–Aug 2022: RGRtln1st, RGRca1st, RGRd1st, RGRht1st) and for the second season (Aug 2022–

Aug/Sep 2023: RGRtln2nd, RGRca2nd, RGRd2nd, RGRht2nd), defined as $(X_2 - X_1)/\Delta\mathrm{day}$. Raw image files used for CA estimation are available from the corresponding author upon reasonable request.

At the end of the second season (4–7 September 2023), all remaining individuals were harvested. After recording TLN, Dia, and Ht, we collected six representative leaves per seedling (or leaves totaling ≥2 g) to quantify individual leaf area (ILA), lobation index (LI), leaf dry mass per leaf area (LMA), and leaf-shape principal components based on elliptic Fourier analysis (PC1ls, PC2ls). Leaves were scanned with a flatbed scanner (GT-S650, EPSON). ILA, LI, and LMA were quantified from scanned images using custom scripts based on OpenCV (Bradski, 2000), and elliptic Fourier analyses were performed using Momocs (Bonhomme et al., 2014). All scripts used for these measurements are available in our GitHub repository. Leaf thickness and tensile properties (Young's modulus) were measured from a strip section excised from the central lamina of one representative leaf per individual; stress was calculated from the force–extension data and thickness.

For biomass measurements, tissues were oven-dried at 70°C for ≥24 h (drying oven: Yamato Scientific Co. Ltd., Japan) and weighed. In addition to total dry weight (DWtot), we quantified dry weights of roots (DWroot), stems (DWstem), and leaves (DWleaf). Budburst date (BBD) was recorded by visual inspection in the common garden from 21 March to 16 May 2023. Herbivory damage was quantified as a predation index based on visual assessment of feeding scars on scanned leaves and non-scanned leaves, categorized by the proportion of damaged area relative to total leaf area (category 1: >50%; category 2: 25–50%; category 3: 1–25%; category 4: <1%).

Separately from the main aims of this study, a subset of 66 seedlings (selected to span the elevation range evenly) was subjected to a late-summer heat-stress experiment to evaluate effects on mRNA expression (Ito et al., in prep). From 25 September 2022, seedlings were transferred indoors for 10 days and assigned either to a heat-stress treatment (n = 33; 35°C; irrigation 50 mL $day^{-1}$) or to a control treatment (n = 33; 22°C; irrigation 50 mL $day^{-1}$); after treatment, seedlings were returned to the common garden and maintained under the same conditions as other individuals. Potential confounding by the heat-stress treatment was addressed by including treatment category (heat stress, control, or not included) as an additional covariate or random effect, as appropriate, in downstream analyses.

**Supplimentary Methods 3. Structural equation modeling**

To test resource-allocation trade-offs and evaluate the effects of genetic background (hybrid index, HI), we constructed structural equation models (SEMs) using piecewiseSEM (Lefcheck, 2016). Because incorporating both trait–trait relationships and explicit temporal structure can lead to over-parameterization, we excluded relative growth rate (RGR) traits from SEM construction. We further excluded total dry weight (DWtot), which is hierarchically dependent on organ-level biomass traits, and leaf-related traits that either showed low loadings in the trait PCA or were judged to have relatively weak functional links to allocation processes that are central to this study (budburst date, BBD; predation index, Pred; and leaf-shape PCs from elliptic Fourier analysis, PC1ls and PC2ls). The final SEM therefore included HI and 11 phenotypic traits (Figure S1).

We organized these variables into four conceptual tiers (Figure S1): (i) genetic ancestry proportion (HI); (ii) leaf functional traits [individual leaf area (ILA), lobation index (LI), Young's modulus (YM), total leaf number (TLN), leaf thickness (Thk), and crown area (CA)]; (iii) plant architecture [stem diameter at 1 cm above ground (Dia) and plant height (Ht)]; and (iv) biomass allocation [root dry weight (DWroot), stem dry weight (DWstem), and leaf dry weight (DWleaf)].

The initial causal structure assumed three main components: (1) direct genetic effects of HI on traits across tiers (HI → leaf functional traits, architecture, and biomass allocation); (2) effects of leaf functional traits on architecture and biomass allocation, reflecting differences in carbon acquisition and associated downstream allocation (leaf functional traits → architecture and biomass allocation); and (3) effects of architectural constraints on biomass allocation (architecture → biomass allocation). In addition, within-tier covariances were included to capture structural constraints and potential investment trade-offs among traits. This initial model structure follows recent trait-based frameworks linking leaf traits to whole-plant allocation (Kleyer et al., 2019; Yin et al., 2019; Zhou et al., 2023; Figure S1).

**Supplimentary References**

Bonhomme, Vincent, Sandrine Picq, Cédric Gaucherel, and Julien Claude. 2014. “Momocs: Outline Analysis UsingR.” Journal of Statistical Software 56 (13): 1–24.

Bradski, G. 2000. “The OpenCV Library.” Dr. Dobb’s Journal of Software Tools.

Gaytan-Legaria, Ricardo, Ken Oyama, Octavio Rojas-Soto, and Antonio González-Rodríguez. 2025. “Dissimilar Climatic Niche Is Predictive of Contrasting Historical Demographic Changes and Altitudinal Shifts in Related Oak Species (*Quercus*).” American Journal of Botany 112 (11): e70122.

Kleyer, Michael, Juliane Trinogga, Miguel A. Cebrián-Piqueras, et al. 2019. “Trait Correlation Network Analysis Identifies Biomass Allocation Traits and Stem Specific Length as Hub Traits in Herbaceous Perennial Plants.” The Journal of Ecology 107 (2): 829–42.

Schoener, Thomas W. 1968. “The Anolis Lizards of Bimini: Resource Partitioning in a Complex Fauna.” Ecology 49 (4): 704–26.

Warren, Dan L., Richard E. Glor, and Michael Turelli. 2008. “Environmental Niche Equivalency versus Conservatism: Quantitative Approaches to Niche Evolution.” Evolution; International Journal of Organic Evolution 62 (11): 2868–83.

Watanabe, Yohei, Chihiro Shirahama, and Kiyoshi Ishida. 2021. “Habitat Segregation between *Fagus crenata* and *Quercus crispula*: Factors Causing Their Distributions on a Snowy Mountain.” Journal of the Japanese Forest Society 103: 379–90.

Yin, Qiulong, Tingting Tian, Xiaohui Han, et al. 2019. “The Relationships between Biomass Allocation and Plant Functional Trait.” Ecological Indicators 102 (July): 302–8.

Zhou, Tianyang, Wentao Du, Jinniu Wang, et al. 2023. “Divergent Responses of Plant Functional Traits and Biomass Allocation to Slope Aspects in Four Perennial Herbs of the Alpine Meadow Ecosystem.” Frontiers in Plant Science 14 (March): 1092821.

#### Figure S1

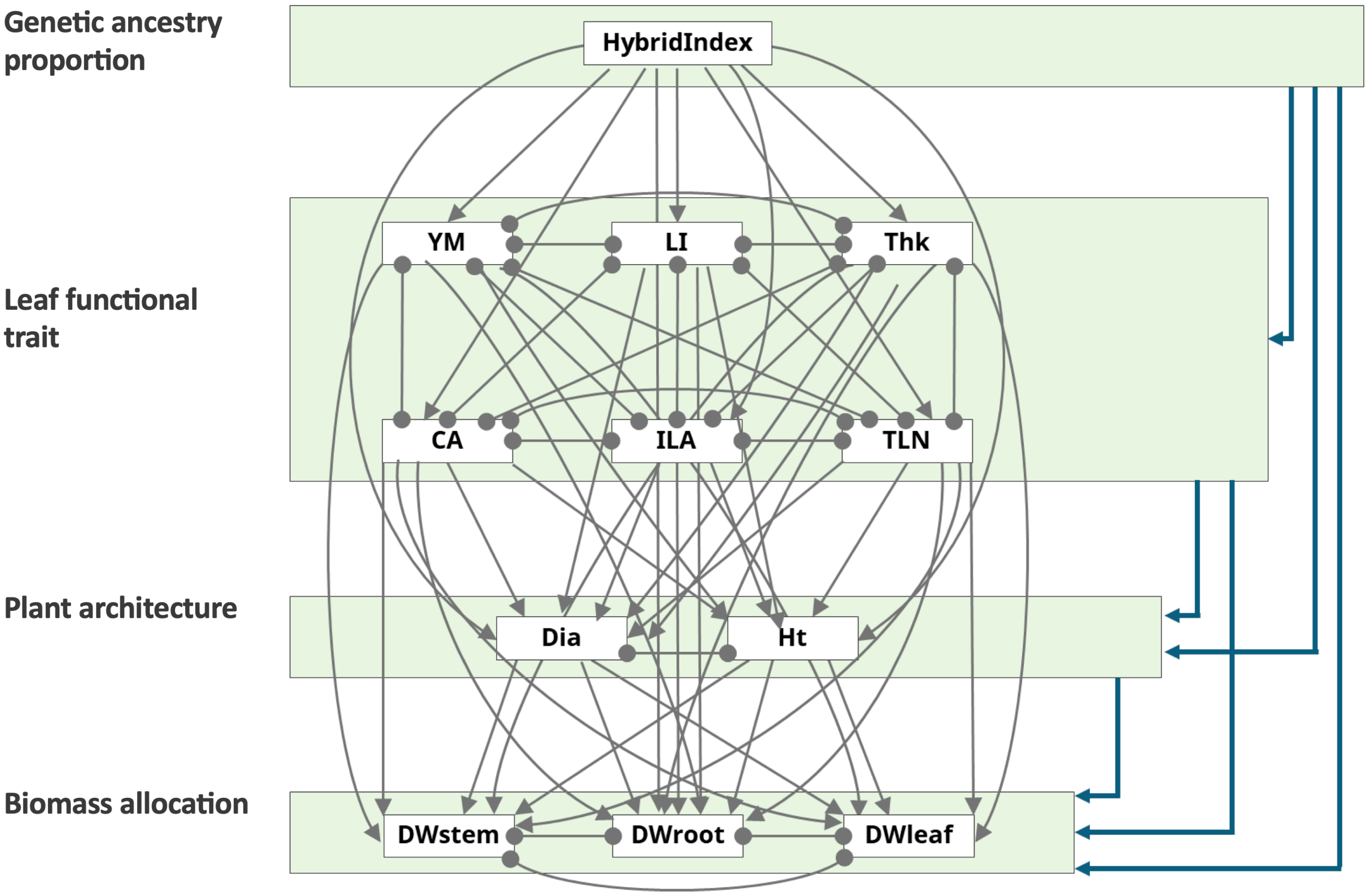


**Figure S1. Initial model of the SEM analysis.**

### Figure S2

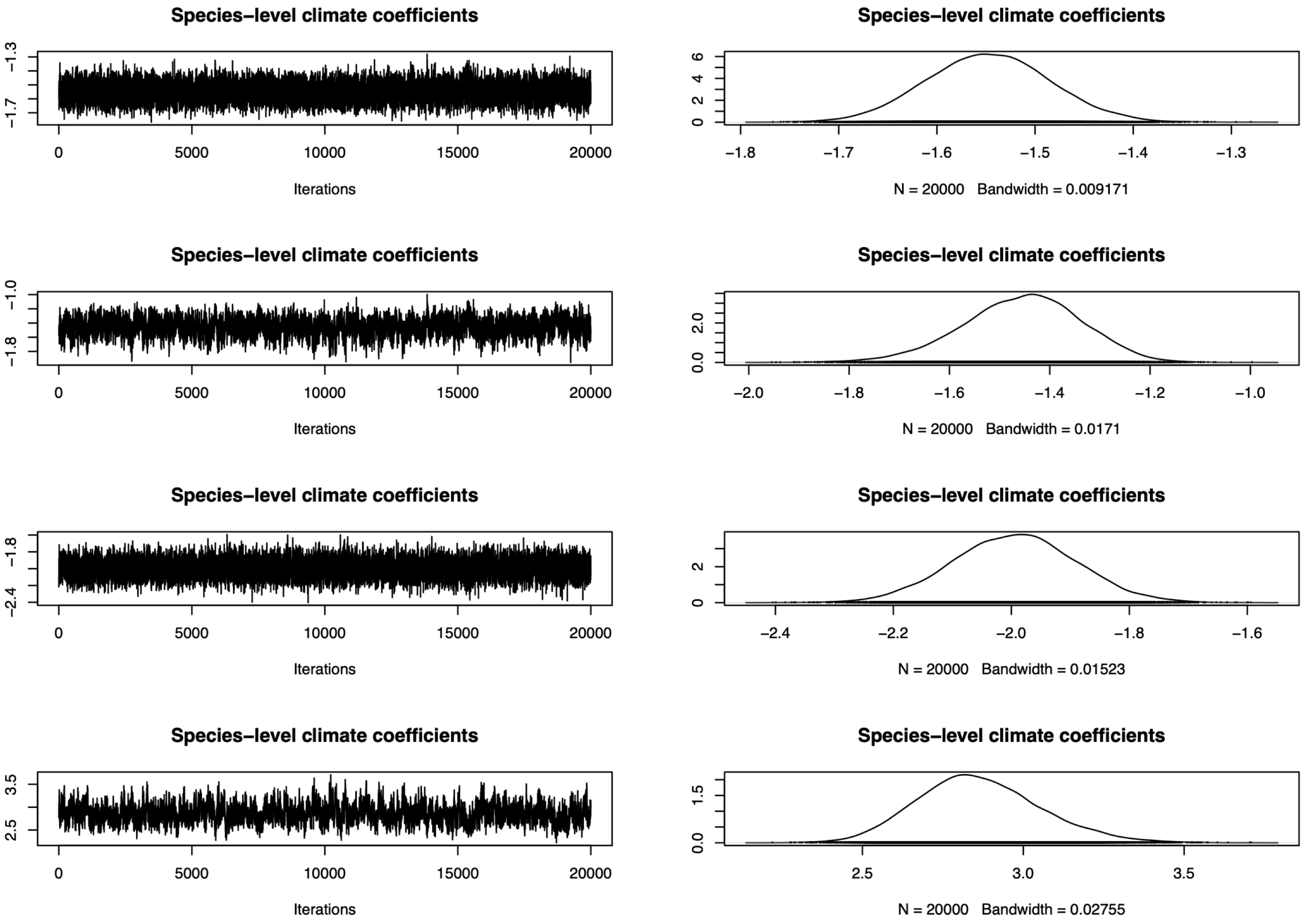


**Figure S2. Trace plots in spOccupancy analysis.** Trace plots of posterior MCMC samples for species-specific climatic-response coefficients estimated for Qmon and Qser.

#### Figure S3

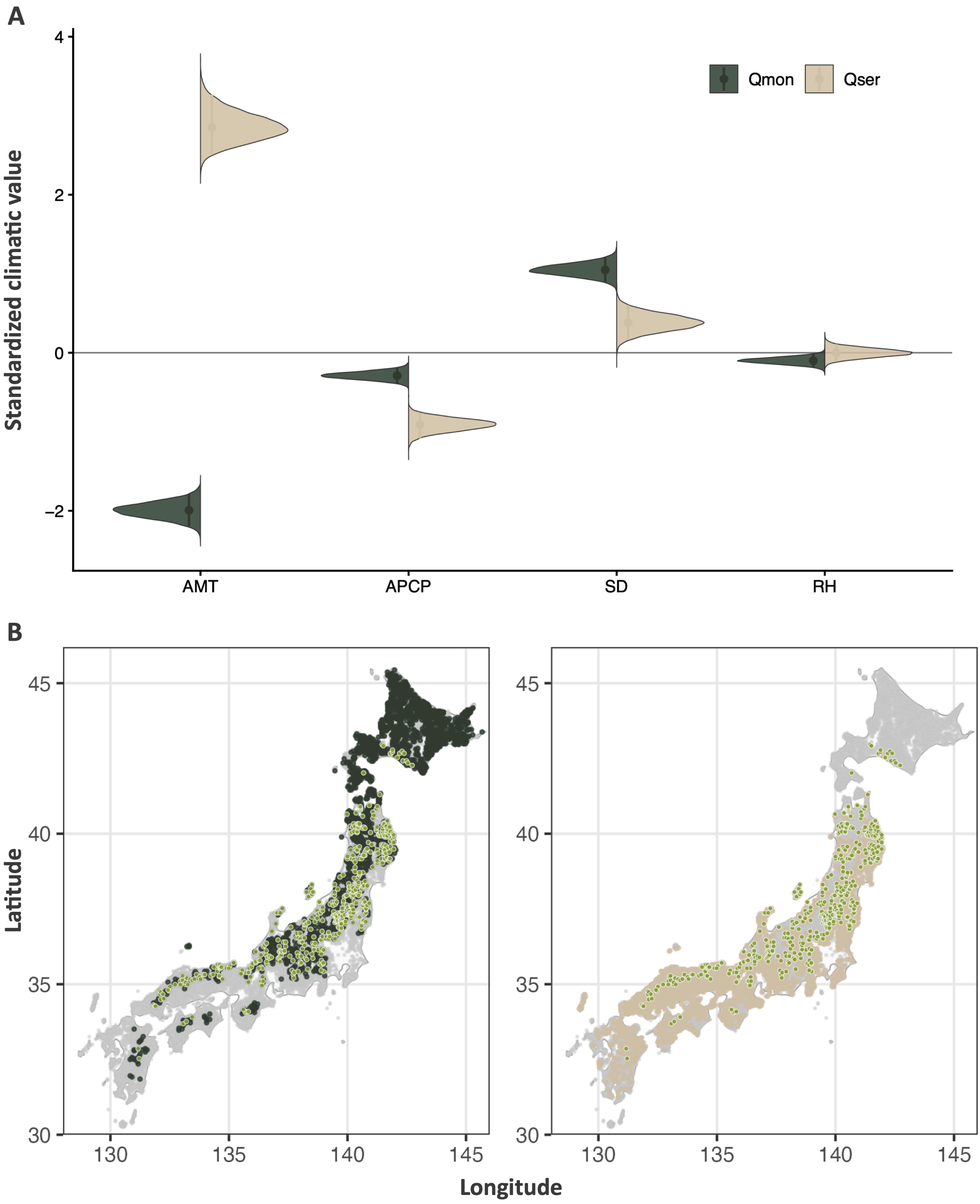


**Figure S3. Results of the spOccupancy analysis.** (A) Comparison of standardized climatic variables, including annual mean temperature (AMT), annual precipitation (APCP), maximum snow depth (SD), and relative humidity (RH), between Qmon and Qser occurrence sites. (B) Geographic distribution of Qmon (dark green) and Qser (beige) occurrence sites. Sites where both species co-occurred are shown as light-green circles.

### Figure S4

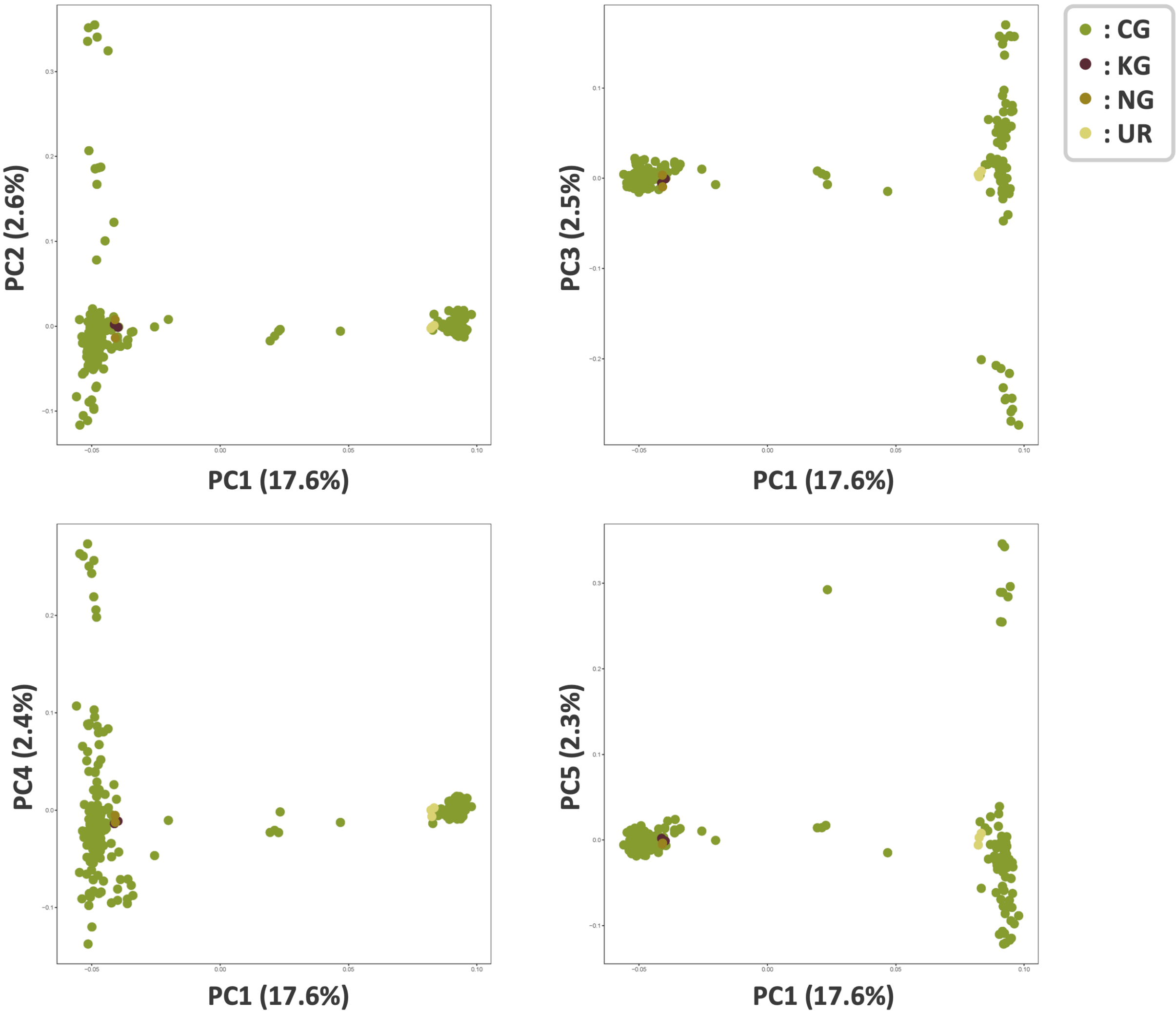


**Figure S4. Details of genomic PCA. (A) PC1–PC2, (B) PC1–PC3, (C) PC1–PC4, (D) PC1–PC5.**

Figure S5

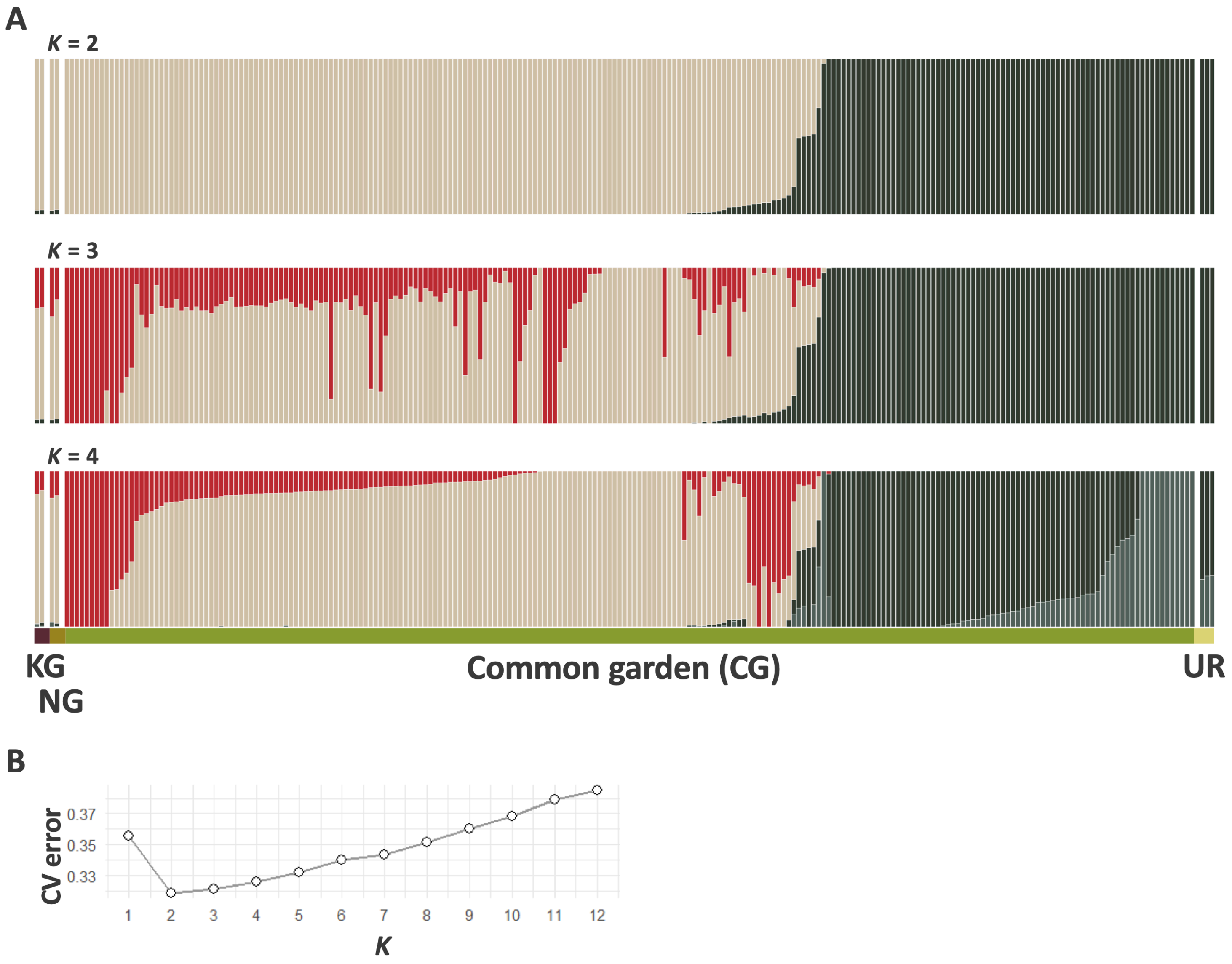


**Figure S5. Details of ADMIXTURE analyses. (A) K = 2–4 bar plots, (B) cross-validation error for K = 1–12.**

### Figure S6

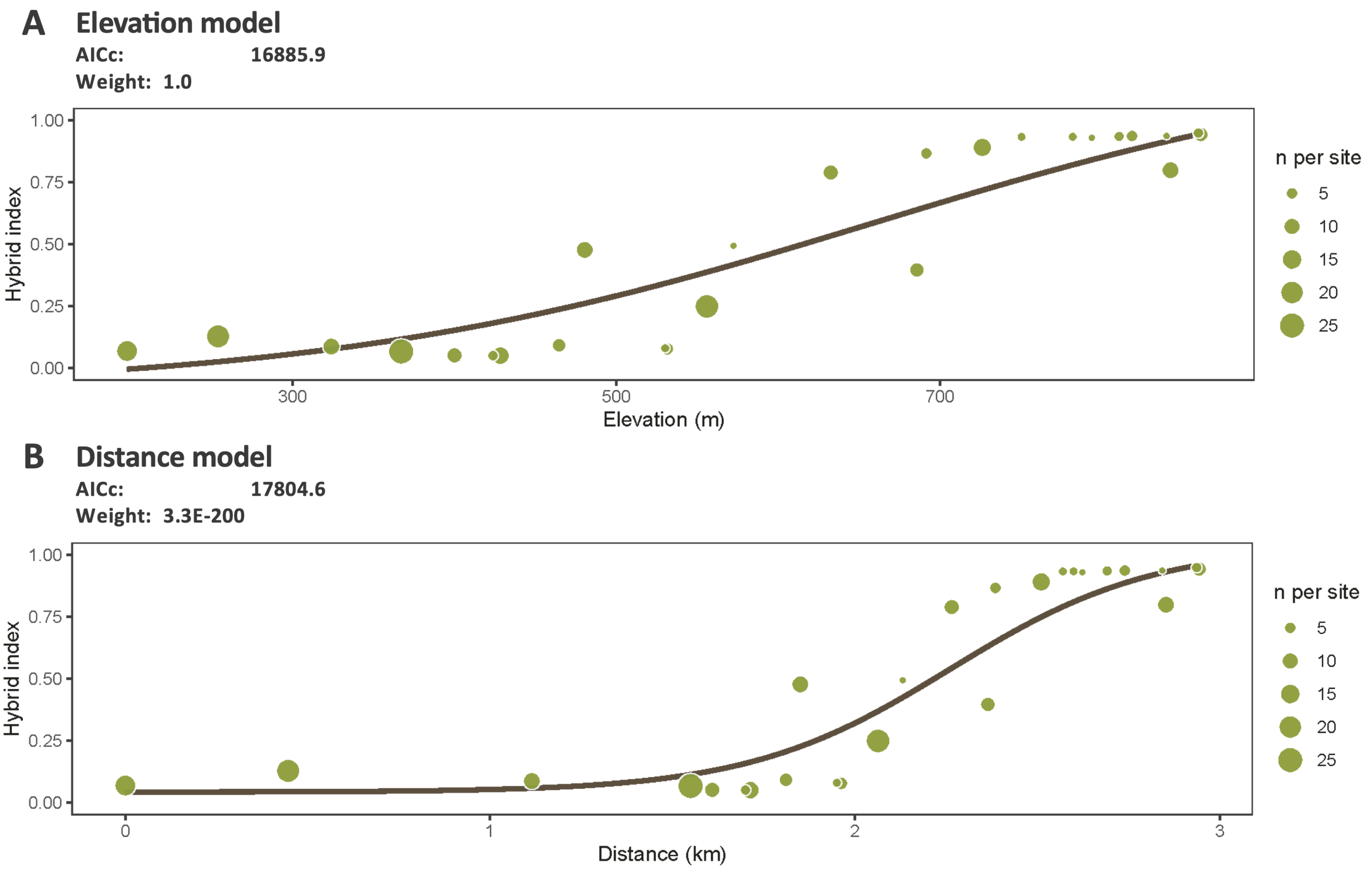


**Figure S6. Site-based clines using hzar. (A) Elevation and (B) distance models.**

### Figure S7

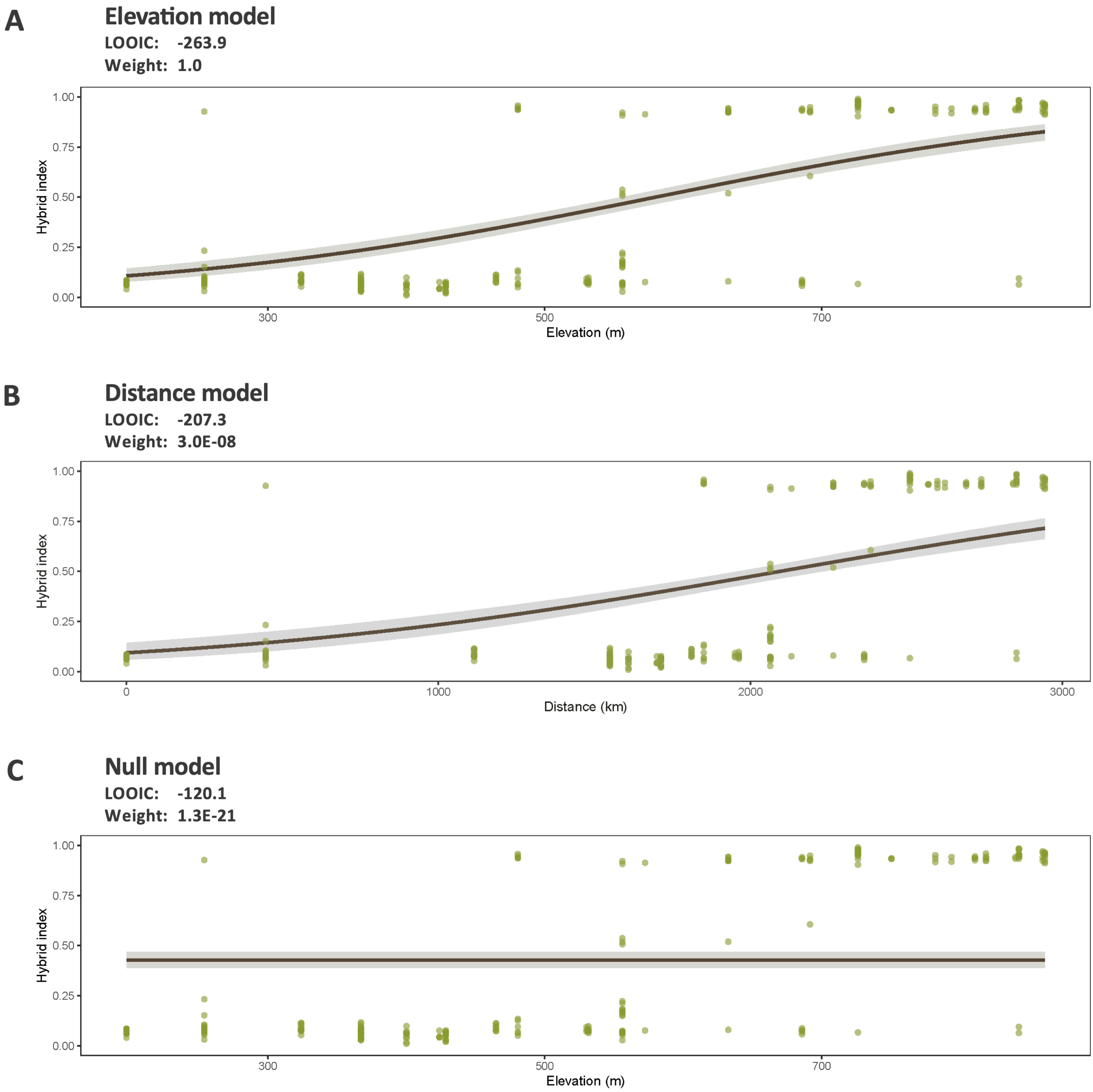


**Figure S7. Individual-based clines using PyMC. (A) Elevation model, (B) Distance model, and (C) null model with a constant mean HI.** Model comparisons are shown.

### Figure S8

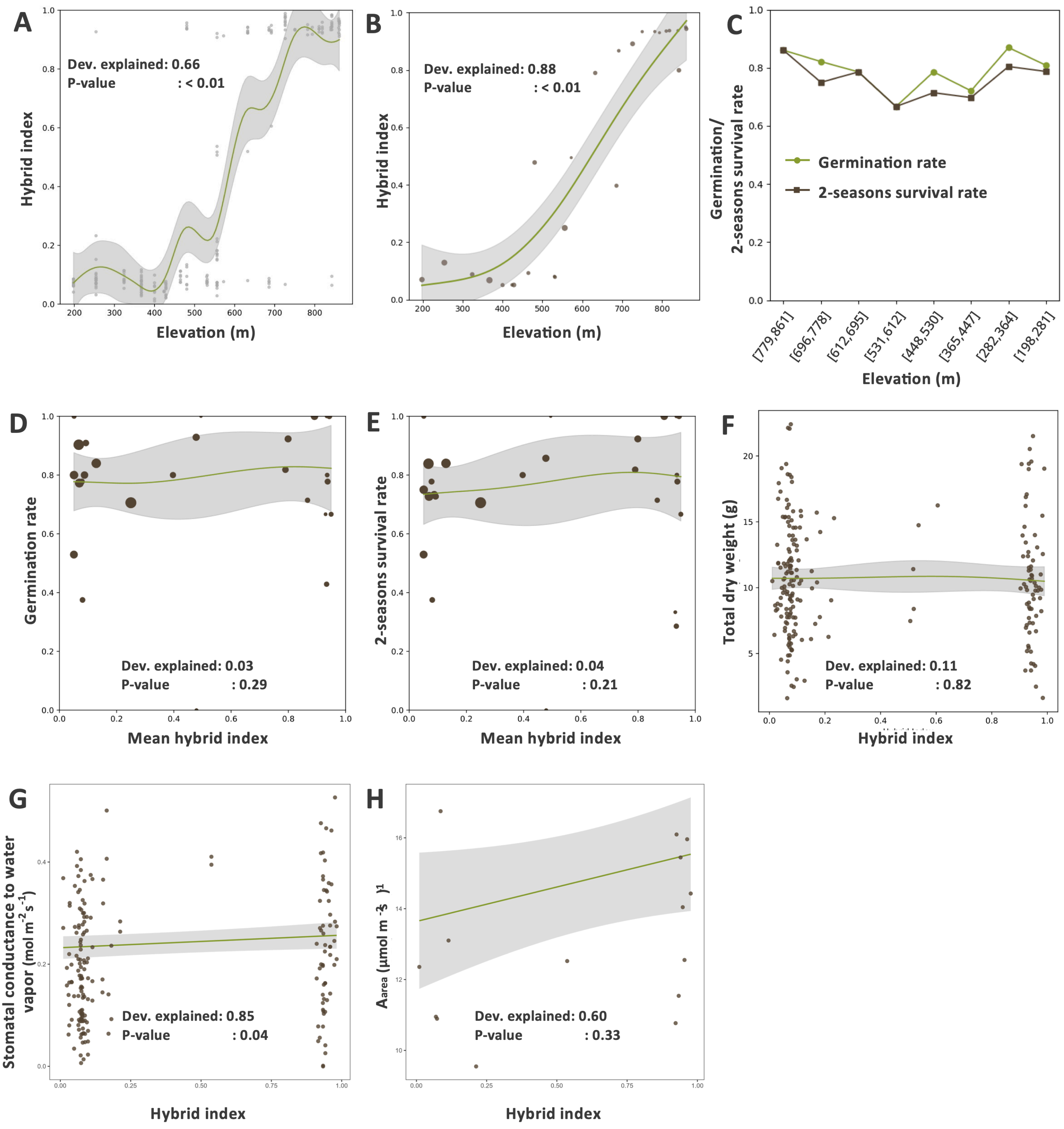


**Figure S8. Elevation, hybrid index (HI), early performance, and leaf gas exchange.** (A) Individual elevation versus HI. (B) Site elevation versus site-mean HI. (C) Germination and two-season survival across eight equal-elevation bins (83 m each). (D) Site-mean HI versus germination. (E) Site-mean HI versus survival. (F) Individual HI versus total dry weight (DWtot). (G) Stomatal conductance to water vapour, $g_{SW}$ (mol $m^{-2}$ $s^{-1}$), versus HI. (H) Net $CO_2$ assimilation at $C_a$ = 420 ppm, $A_{area}$ (μmol $m^{-2}$ $s^{-1}$), versus HI.

### Figure S9

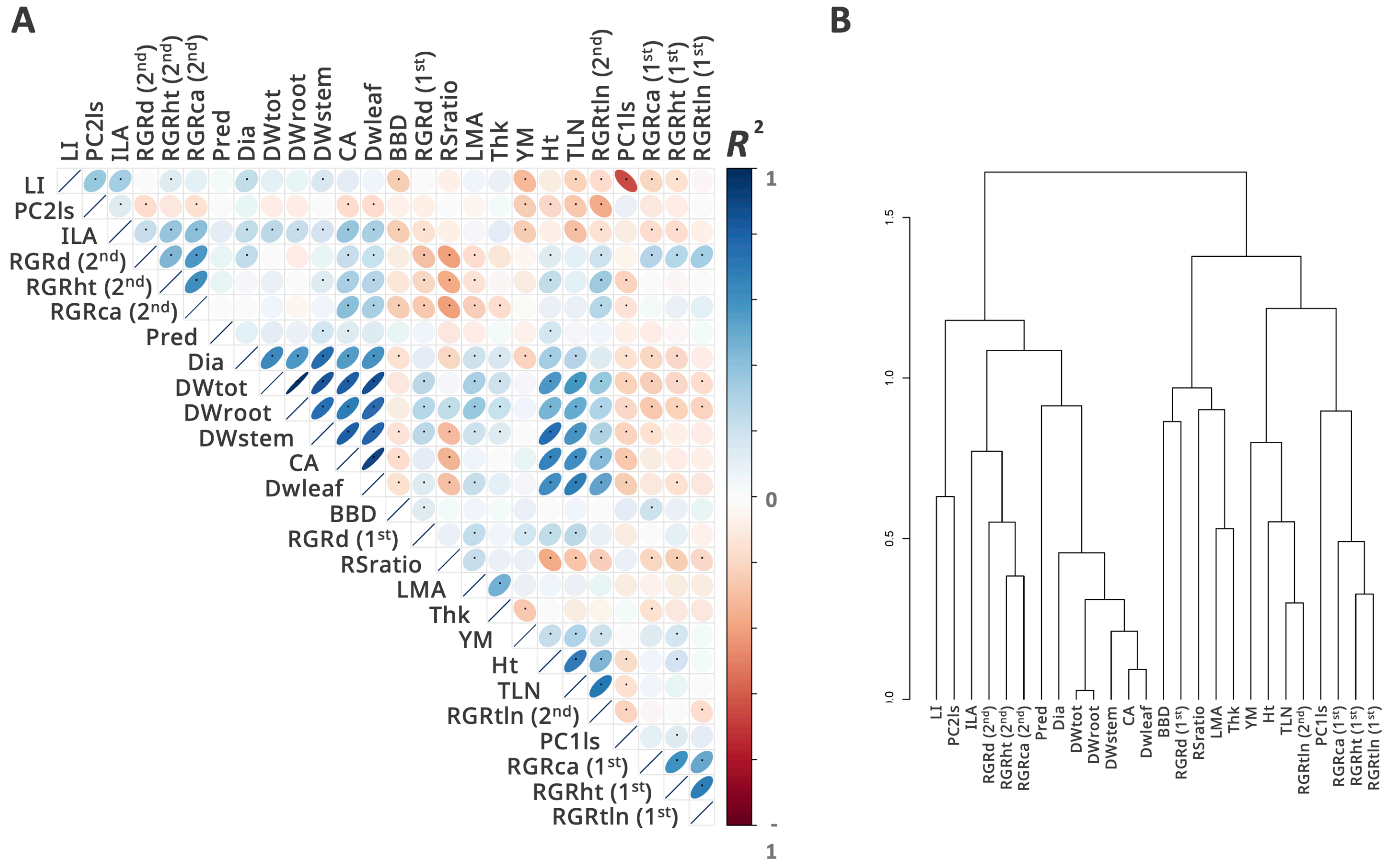


**Figure S9. Pairwise correlation matrix of the 26 traits with hierarchical clustering dendrogram.**

#### Figure S10

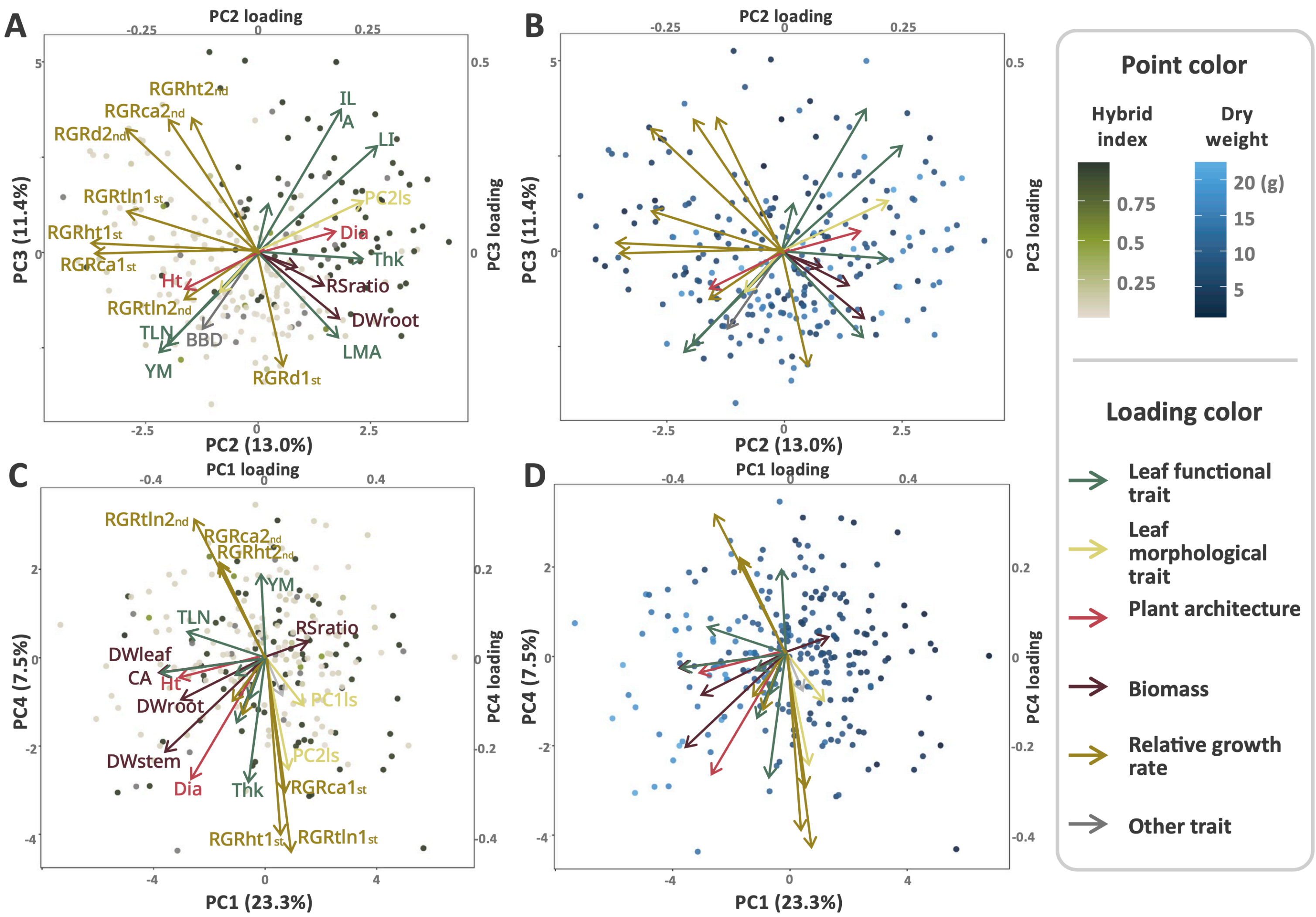


**Figure S10. PCA details. Significant (A) PC2 and (B) PC3 vs. HI, and non-significant (C) PC1 and (D) PC4. Point colors: (A, C) HI; (B, D) DWtot. Loading colors indicate trait categories.**

**Figure S11**

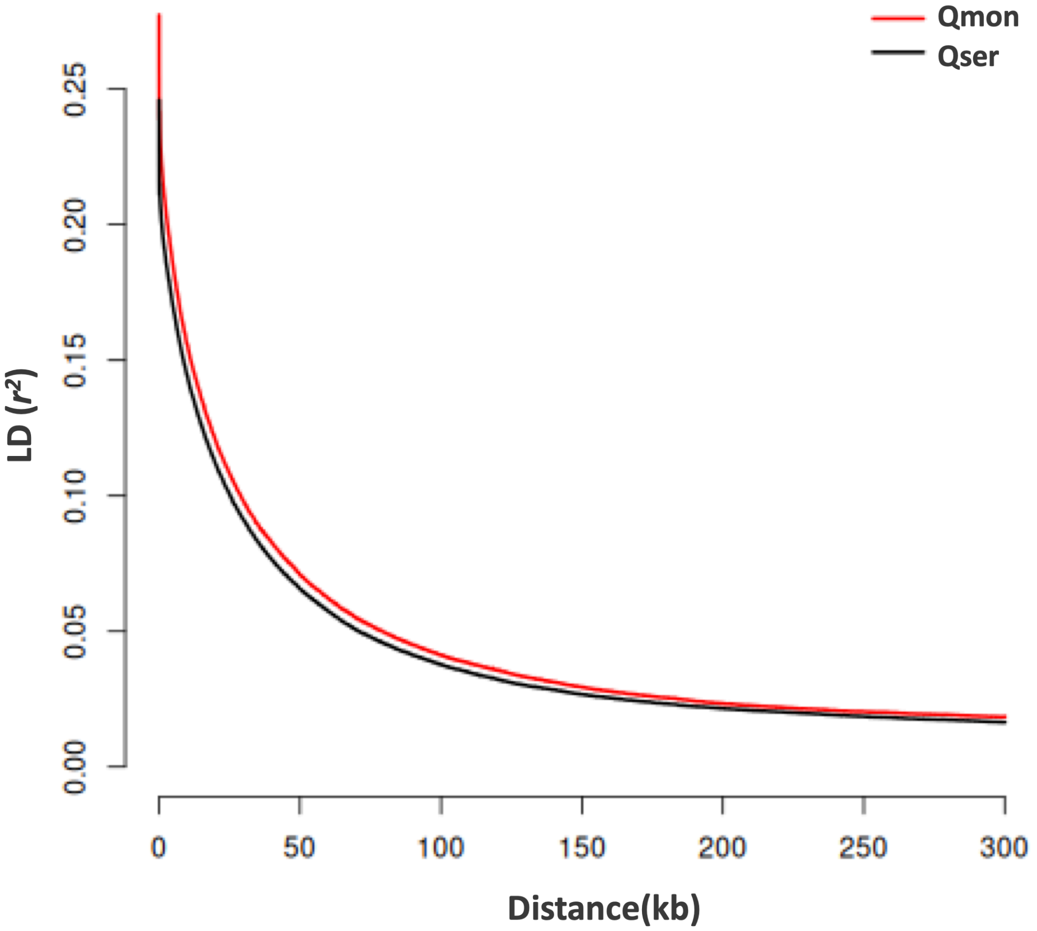


**Figure S11. Linkage disequilibrium decay in the parental-type groups (Qmon and Qser).**

### Figure S12

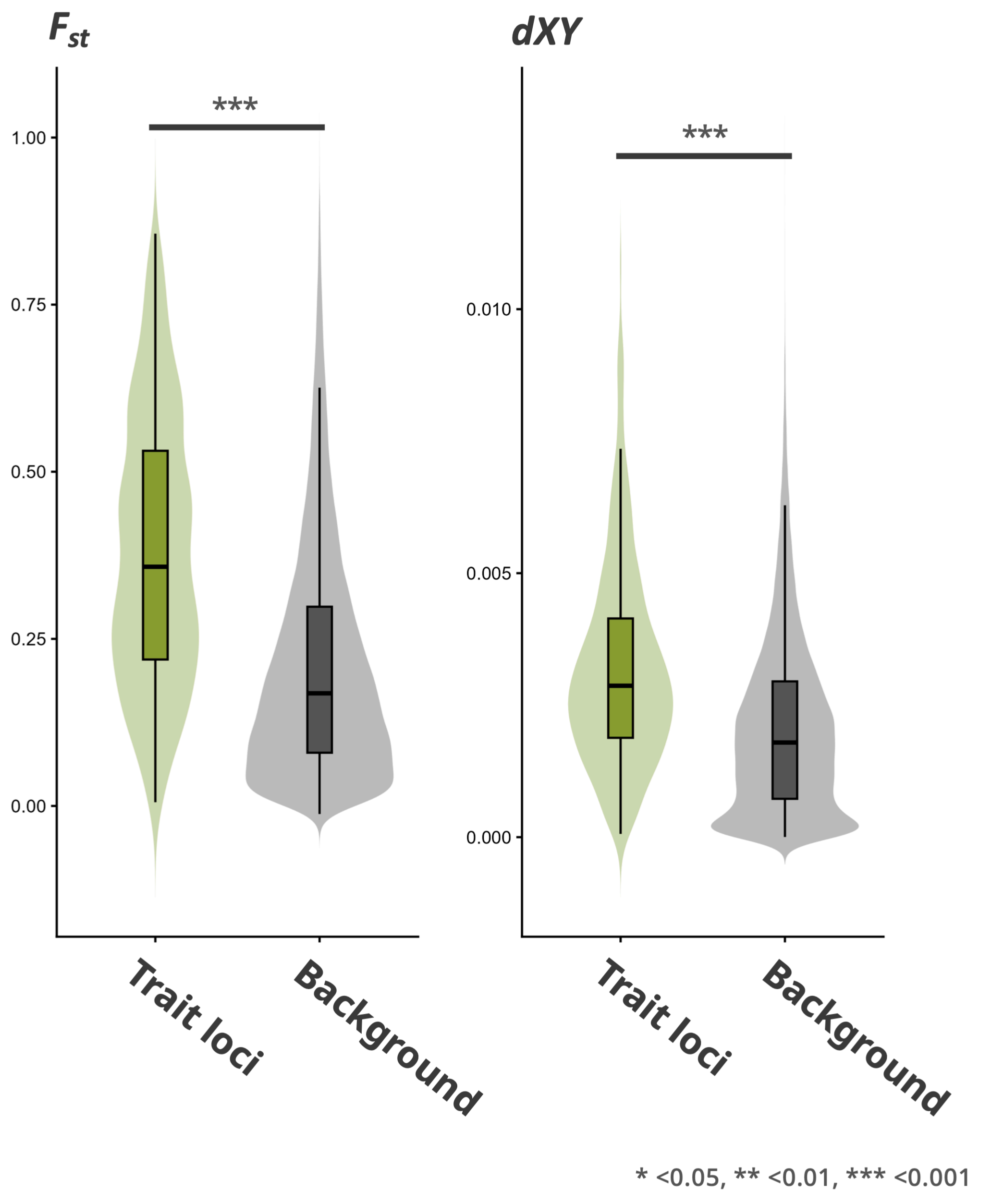


**Figure S12. Parental genetic divergence ($F_{ST}$, dXY) for 10 kb windows containing trait loci (light green) vs. background windows (gray).** Asterisks indicate significance (* $P < 0.05$, ** $P < 0.01$, *** $P < 0.001$).

### Figure S13

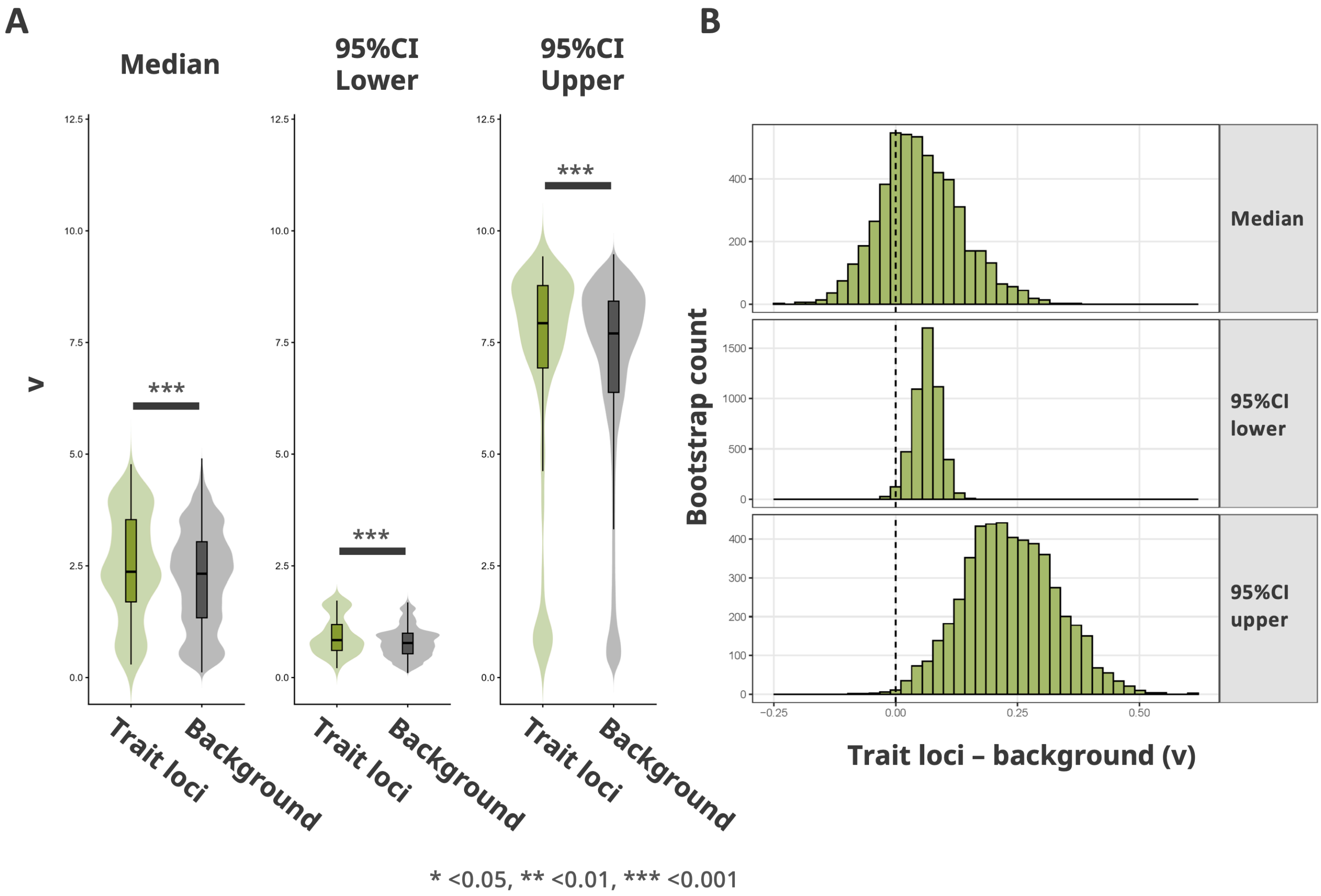


**Figure S13. Local cline slope (v) comparison: median and 95% CI (lower, upper) for trait loci (light green) vs. background (gray).** (A) Violin plots; (B) bootstrap distributions of differences. Asterisk notation as in S12.

**Table S1. Summary of read-mapping quality-control statistics used to assess potential mapping bias associated with hybrid index (HI).** Values are presented as mean ± SD across individuals. Pearson's R represents the correlation between each mapping metric and HI. Slope and P values were obtained from linear regressions of each mapping metric against HI.

| **Mapping quality-control metric** | **Mean ± SD (%)** | **Pearson's *R*** | **Slope** | ***P*** |
|---|---|---|---|---|
| Properly paired reads | 96.4 ± 0.38 | 0.15 | 0.14 | **0.02** |
| Singleton reads | 0.47 ± 0.17 | −0.10 | −0.04 | 0.15 |
| Supplementary alignments | 0.71 ± 0.06 | −0.16 | −0.02 | **0.02** |
| Mates mapped to different chromosomes | 2.40 ± 0.26 | −0.07 | −0.05 | 0.26 |

**Table S2. Summary of seedling traits. Categories, sample sizes (n), measurement timing, and definitions.**

| Trait | Abbreviation | Category | n | Measurement timing | Description |
|---|---|---|---|---|---|
| Leaf mass per area | LMA | Leaf functional trait | 236 | Sep. 2023 (2nd season) | Leaf dry mass divided by area ($g/cm^2$). |
| Individual leaf area | ILA | Leaf functional trait | 238 | Sep. 2023 (2nd season) | Average leaf area calculated from several representative leaves per individual. |
| Total leaf number | TLN | Leaf functional trait | 239 | Jun, Aug. 2022 (1st season); Sep. 2023 (2nd season) | Total number of leaves per individual. |
| Crown area | CA | Leaf functional trait | 234 | Jun, Aug. 2022 (1st season); Aug. 2023 (2nd season) | Estimated leaf crown area calculated from overhead photographs. |
| Leaf thickness | Thk | Leaf functional trait | 230 | Sep. 2023 (2nd season) | Leaf thickness. |
| Young's modulus | YM | Leaf functional trait | 230 | Sep. 2023 (2nd season) | Elastic modulus of the leaf indicating stiffness. |
| Lobation index | LI | Leaf functional trait | 238 | Sep. 2023 (2nd season) | Ratio of leaf perimeter to the square root of leaf area, used to quantify the degree of lobing. |
| Leaf shape PC1 (Fourier) | PC1ls | Morphological trait | 238 | Sep. 2023 (2nd season) | Principal component 1 of leaf contour shape from elliptical Fourier analysis. |
| Leaf shape PC2 (Fourier) | PC2ls | Morphological trait | 238 | Sep. 2023 (2nd season) | Principal component 2 of leaf contour shape from elliptical Fourier analysis. |
| Height | Ht | Plant architecture | 239 | Jun, Aug. 2022 (1st season); Sep. 2023 (2nd season) | Vertical length from the soil surface to the apex of the plant. |
| Diameter | Dia | Plant architecture | 239 | Jun, Aug. 2022 (1st season); Sep. 2023 (2nd season) | Stem diameter measured at the base. |
| Dry weight (Total) | DWtot | Biomass | 232 | Sep. 2023 (2nd season) | Total dry biomass after oven-drying all organs. |
| Dry weight (Root) | DWroot | Biomass | 236 | Sep. 2023 (2nd season) | Dry weight of the root system. |
| Dry weight (Stem) | DWstem | Biomass | 239 | Sep. 2023 (2nd season) | Dry weight of the stem. |
| Dry weight (Leaf) | DWleaf | Biomass | 237 | Sep. 2023 (2nd season) | Dry weight of all leaves. |
| Root/Shoot ratio | Rsratio | Biomass | 232 | Sep. 2023 (2nd season) | Root/Shoot ratio |
| Relative growth rate (Diameter) 1st section | RGRd1st | Relative growth rate | 234 | Jun to Aug. 2022 (1st season) | Relative increase in stem diameter over time. |
| Relative growth rate (Diameter) 2nd section | RGRd2nd | Relative growth rate | 237 | Aug. 2022 to Sep. 2023 (1st to 2nd season) | Relative increase in stem diameter over time. |
| Relative growth rate (Height) 1st section | RGRht1st | Relative growth rate | 234 | Jun to Aug. 2022 (1st season) | Relative increase in plant height over time. |
| Relative growth rate (Height) 2nd section | RGRht2nd | Relative growth rate | 238 | Aug. 2022 to Sep. 2023 (1st to 2nd season) | Relative increase in plant height over time. |
| Relative growth rate (Total leaf number) 1st section | RGRtln1st | Relative growth rate | 232 | Jun to Aug. 2022 (1st season) | Relative increase in total leaf number over time. |
| Relative growth rate (Total leaf number) 2nd section | RGRtln2nd | Relative growth rate | 237 | Aug. 2022 to Sep. 2023 (1st season) | Relative increase in total leaf number over time. |
| Relative growth rate (Crown area) 1st section | RGRca1st | Relative growth rate | 225 | Jun to Aug. 2022 (1st season) | Relative increase in crown area over time. |
| Relative growth rate (Crown area) 2nd section | RGRca2nd | Relative growth rate | 233 | Aug. 2022 to Aug. 2023 (1st to 2nd season) | Relative increase in crown area over time. |
| Bud break date | BBD | Other trait | 238 | Apr. 2023 (2nd season) | Number of days from April 1st of the second season to bud break. |
| Predation index | PI | Other trait | 238 | Sep. 2023 (2nd season) | Proportion of leaf area damaged by herbivores. |

**Table S3. Posterior estimates of climatic effects on the occurrence of Quercus mongolica and Q. serrata. Values represent posterior means with 95% credible intervals (CRL).** Δ represents the posterior difference in regression coefficients between Q. mongolica and Q. serrata (Q. mongolica − Q. serrata). AMT, annual mean temperature; APCP, annual precipitation; SD, maximum snow depth; RH, relative humidity. A 95% CrI excluding zero indicates strong posterior support for a non-zero effect or species difference.

| Predictor | Q. mongolica, posterior mean [95% CRL] | Q. serrata, posterior mean [95% CRL] | Difference (Δ), posterior mean [95% CRL] |
|---|---|---|---|
| Intercept | −1.547 [−1.669, −1.423] | −1.456 [−1.701, −1.240] | −0.091 [−0.183, 0.032] |
| AMT | −1.995 [−2.198, −1.788] | 2.863 [2.521, 3.267] | −4.858 [−5.055, −4.578] |
| $AMT^2$ | −1.530 [−1.664, −1.397] | −1.810 [−2.063, −1.598] | 0.281 [0.201, 0.399] |
| APCP | −0.289 [−0.388, −0.189] | −0.912 [−1.082, −0.759] | 0.623 [0.570, 0.693] |
| $APCP^2$ | 0.118 [0.051, 0.185] | −0.017 [−0.109, 0.072] | 0.136 [0.113, 0.160] |
| SD | 1.049 [0.891, 1.210] | 0.382 [0.163, 0.604] | 0.667 [0.606, 0.728] |
| $SD^2$ | −0.422 [−0.486, −0.360] | −0.366 [−0.499, −0.243] | −0.055 [−0.117, 0.013] |
| RH | −0.101 [−0.187, −0.016] | −0.002 [−0.115, 0.111] | −0.099 [−0.072, −0.127] |
| $RH^2$ | −0.073 [−0.129, −0.018] | 0.077 [0.015, 0.139] | −0.150 [−0.144, −0.157] |

**Table S4. Proportion of variance explained by principal components from the genomic PCA.** Includes individual and cumulative variance (%)

| PC | Contribution rate (%) | Cumulative variance (%) |
|---|---|---|
| PC1 | 17.56 | 17.56 |
| PC2 | 2.72 | 20.28 |
| PC3 | 2.51 | 22.79 |
| PC4 | 2.45 | 25.24 |
| PC5 | 2.31 | 27.55 |
| PC6 | 2.21 | 29.76 |
| PC7 | 2.12 | 31.88 |
| PC8 | 2.01 | 33.89 |
| PC9 | 1.97 | 35.86 |
| PC10 | 1.94 | 37.80 |
| PC11 | 1.91 | 39.71 |
| PC12 | 1.89 | 41.60 |

**Table S5. Estimated parameters from the cline analyses and model comparison between the Elevation and Distance models.** Site-based (hzar) results present Centre and Width (mean, 95% HDI), AICc, and model weight. Individual-based (PyMC) results present Centre, Width, κ (mean, 95% HDI), LOOIC, and model weight.

**Site-based cline**

| Model | Center | | Width | | | | AICc | Weight |
|---|---|---|---|---|---|---|---|---|
| | Mean | 95%HDI | Mean | 95%HDI | | | | |
| Elevation | 660.4 | [652.9, 668.3] | 685.6 | [646.2, 729.0] | - | - | 16885.9 | 1.00 |
| Distance | 2263 | [2251, 2274] | 1110 | [1075, 1146] | - | - | 17804.6 | 3.30E-200 |

**Individual-based cline**

| Model | Center | | Width | | kappa | | LOOIC | Weight |
|---|---|---|---|---|---|---|---|---|
| | Mean | 95%HDI | Mean | 95%HDI | Mean | 95%HDI | | |
| Elevation | 580.3 | [555.4, 605.1] | 181.4 | [154.9, 209.8] | 2.2 | [1.8, 2.6] | -263.9 | 1.00 |
| Distance | 2095.7 | [1960.6, 2231.5] | 6.3 | [0.6, 12.4] | 1.7 | [1.4, 2.0] | -207.3 | 3.00E-08 |
| Null | - | - | - | - | 1.1 | [0.9,1.2] | -120.1 | 1.30E-21 |

**Table S6. Summary of generalised additive models (GAMs) relating hybrid index (HI), environmental covariates, and seedling fitness- and gas-exchange-related traits.** For each model, we present the response equation, sample size (n), error distribution and link function, pseudo-$R^2$ (deviance explained), total effective degrees of freedom (EDF; summed across smooth terms), P-value for the focal smooth term, and the Akaike information criterion (AIC).

| Model | Eq. | n | Family (link) | Deviance explained | EDF (total) | P (focal smooth term) | AIC |
|---|---|---|---|---|---|---|---|
| Germination rate ~ s(mean HI) | 1 | 26 | Binomial (logit) | 0.03 | 1.0 | 0.29 | 97.1 |
| 2-season survival rate ~ s(mean HI) | 1 | 26 | Binomial (logit) | 0.04 | 1.0 | 0.21 | 100.2 |
| HI ~ s(Elevation) + Density + Treatment | – | 223 | Gaussian (identity) | 0.66 | 4.6 | < 0.01 | -0.3 |
| mean HI ~ s(Elevation) | – | 26 | Gaussian (identity) | 0.88 | 3.3 | < 0.01 | -15.3 |
| Total dry weight (DWtot) ~ s(HI) + Density + Treatment | 2 | 214 | Gaussian (identity) | 0.11 | 1 | 0.82 | 1223.7 |
| gsw ~ s(HI) + s(VPD_leaf) + s(log(1+PAR)) + PlantID (random) | 3 | 93 individuals × 2 measurements | Gaussian (identity) | 0.85 | 79.6 | 0.04 | -560.6 |
| A_area ~ s(HI) + s(VPD_leaf) + s(T_leaf) | 4 | 15 | Gaussian (identity) | 0.60 | 4.0 | 0.33 | 64.7 |

**Table S7. Variance explained by phenotypic principal components and linear GAM results linking PCs to hybrid index and total dry weight.** For each PC: variance explained, EDF, F-statistics, raw and FDR-adjusted P-values for models of the form $PC_n$ ~ s(HI) + f(Density) + f(Treatment) and DWtot ~ s($PC_n$) + f(Density) + f(Treatment). (Software and versions are specified in Methods.)

| PC | Hybrid index | | | Total dry weight | |
|---|---|---|---|---|---|
| | $R^2$ | adjP | EDF (total) | P-value | EDF (total) |
| PC1 | 0.08 | 0.12 | 1.0 | <0.01 | 3.5 |
| PC2 | 0.30 | <0.01 | 1.0 | <0.01 | 2.1 |
| PC3 | 0.20 | <0.01 | 1.2 | <0.01 | 1.0 |
| PC4 | 0.01 | 0.05 | 1.4 | <0.01 | 2.4 |
| PC5 | 0.02 | 0.82 | 1.0 | <0.01 | 1.0 |
| PC6 | 0.08 | 0.06 | 1.0 | <0.01 | 1.0 |
| PC7 | 0 | 0.12 | 2.1 | <0.01 | 2.8 |
| PC8 | -0.01 | 0.12 | 1.0 | <0.01 | 1.1 |
| PC9 | -0.02 | 0.57 | 1.0 | <0.01 | 5.8 |
| PC10 | 0.04 | 0.82 | 1.0 | <0.01 | 1.0 |

**Table S8. Loadings of morphological and physiological traits on the first four phenotypic principal components.** Loadings with |value| ≥ 0.25 are highlighted.

| Trait abbreviation | Category | PC1 | PC2 | PC3 | PC4 |
|---|---|---|---|---|---|
| LMA | Leaf functional trait | -0.11 | 0.18 | -0.22 | -0.15 |
| ILA | Leaf functional trait | -0.11 | 0.19 | **0.37** | -0.04 |
| TLN | Leaf functional trait | **-0.28** | -0.20 | -0.25 | 0.06 |
| CA | Leaf functional trait | **-0.38** | -0.03 | 0.05 | -0.04 |
| Thk | Leaf functional trait | -0.06 | 0.23 | -0.02 | **-0.28** |
| YM | Leaf functional trait | -0.02 | -0.22 | **-0.26** | 0.18 |
| LI | Leaf functional trait | -0.05 | **0.27** | **0.28** | -0.09 |
| PC1ls | Morphological trait | 0.13 | -0.08 | -0.11 | -0.11 |
| PC2ls | Morphological trait | 0.08 | 0.23 | 0.13 | -0.25 |
| Ht | Plant architecture | **-0.31** | -0.16 | -0.10 | -0.05 |
| Dia | Plant architecture | **-0.27** | 0.18 | 0.05 | **-0.27** |
| DWroot | Biomass | **-0.30** | 0.18 | -0.17 | -0.10 |
| DWstem | Biomass | **-0.36** | 0.09 | -0.04 | -0.21 |
| DWleaf | Biomass | **-0.38** | 0.00 | -0.02 | -0.04 |
| Rsratio | Biomass | 0.15 | 0.15 | -0.09 | 0.03 |
| RGRd1st | Relative growth rate | -0.08 | 0.06 | **-0.30** | -0.13 |
| RGRd2nd | Relative growth rate | -0.12 | **-0.29** | **0.32** | -0.10 |
| RGRht1st | Relative growth rate | 0.06 | **-0.37** | 0.02 | **-0.40** |
| RGRht2nd | Relative growth rate | -0.16 | -0.14 | **0.35** | 0.20 |
| RGRtln1st | Relative growth rate | 0.09 | **-0.29** | 0.11 | **-0.43** |
| RGRtln2nd | Relative growth rate | **-0.26** | -0.16 | -0.12 | **0.31** |
| RGRca1st | Relative growth rate | 0.07 | **-0.36** | 0.00 | **-0.30** |
| RGRca2nd | Relative growth rate | -0.16 | -0.20 | **0.35** | 0.21 |
| BBD | Other trait | 0.06 | -0.12 | -0.20 | -0.09 |
| PI | Other trait | -0.03 | 0.02 | 0.13 | -0.01 |

**Table S9. Standardized path coefficients and significance for pairwise relationships among traits in the final piecewise structural equation model.** Standardized coefficients (β), 95% CI, P-values, Fisher's C, and model fit indices are presented.

| Response | Predictor | Estimate | Std.Error | DF | Crit.Value | P-Value | Std.Estimate | |
|---|---|---|---|---|---|---|---|---|
| ILA | HybridIndex | 6022.77 | 1080.37 | 221 | 5.57 | < 0.001 | 0.35 | *** |
| LI | HybridIndex | 0.13 | 0.01 | 221 | 8.78 | < 0.001 | 0.51 | *** |
| YM | HybridIndex | -12.06 | 1.50 | 221 | -8.05 | < 0.001 | -0.48 | *** |
| TLN | HybridIndex | -18.42 | 2.73 | 221 | -6.75 | < 0.001 | -0.41 | *** |
| Dia | CA | 0.00 | 0.00 | 217 | 7.49 | < 0.001 | 0.48 | *** |
| Dia | LMA | 0.04 | 0.01 | 217 | 3.39 | 0.001 | 0.17 | *** |
| Dia | YM | -0.01 | 0.01 | 217 | -2.29 | 0.023 | -0.13 | * |
| Dia | TLN | 0.01 | 0.00 | 217 | 2.17 | 0.031 | 0.15 | * |
| Dia | HybridIndex | 0.84 | 0.15 | 217 | 5.45 | < 0.001 | 0.34 | *** |
| Ht | CA | 0.00 | 0.00 | 218 | 7.17 | < 0.001 | 0.53 | *** |
| Ht | TLN | 0.14 | 0.04 | 218 | 3.44 | 0.001 | 0.26 | *** |
| Ht | ILA | 0.00 | 0.00 | 218 | -2.00 | 0.047 | -0.13 | * |
| Ht | HybridIndex | -2.86 | 1.16 | 218 | -2.47 | 0.014 | -0.12 | * |
| Dwroot | LMA | 0.20 | 0.03 | 217 | 7.10 | < 0.001 | 0.31 | *** |
| Dwroot | CA | 0.00 | 0.00 | 217 | 7.45 | < 0.001 | 0.49 | *** |
| Dwroot | TLN | 0.03 | 0.01 | 217 | 3.00 | 0.003 | 0.18 | ** |
| Dwroot | Dia | 0.71 | 0.16 | 217 | 4.49 | < 0.001 | 0.23 | *** |
| Dwroot | Ht | -0.04 | 0.02 | 217 | -2.05 | 0.041 | -0.13 | * |
| Dwstem | ILA | 0.00 | 0.00 | 216 | -3.79 | < 0.001 | -0.11 | *** |
| Dwstem | LMA | 0.01 | 0.00 | 216 | 3.23 | 0.001 | 0.08 | ** |
| Dwstem | CA | 0.00 | 0.00 | 216 | 12.25 | < 0.001 | 0.49 | *** |
| Dwstem | Dia | 0.26 | 0.02 | 216 | 11.25 | < 0.001 | 0.35 | *** |
| Dwstem | Ht | 0.02 | 0.00 | 216 | 8.55 | < 0.001 | 0.30 | *** |
| Dwstem | HybridIndex | 0.30 | 0.06 | 216 | 5.54 | < 0.001 | 0.16 | *** |
| Dwleaf | LMA | 0.03 | 0.00 | 216 | 7.59 | < 0.001 | 0.17 | *** |
| Dwleaf | ILA | 0.00 | 0.00 | 216 | 5.19 | < 0.001 | 0.17 | *** |
| Dwleaf | CA | 0.00 | 0.00 | 216 | 13.41 | < 0.001 | 0.61 | *** |
| Dwleaf | TLN | 0.02 | 0.00 | 216 | 10.55 | < 0.001 | 0.43 | *** |
| Dwleaf | Dia | 0.06 | 0.02 | 216 | 2.86 | 0.005 | 0.08 | ** |
| Dwleaf | Ht | -0.01 | 0.00 | 216 | -3.61 | < 0.001 | -0.12 | *** |
| ~~ILA | ~~LI | 0.20 | - | 223 | 2.95 | 0.002 | 0.20 | ** |
| ~~CA | ~~TLN | 0.60 | - | 223 | 11.08 | < 0.001 | 0.60 | *** |
| ~~LI | ~~CA | 0.16 | - | 223 | 2.40 | 0.009 | 0.16 | ** |
| ~~ILA | ~~CA | 0.38 | - | 223 | 6.16 | < 0.001 | 0.38 | *** |
| ~~ILA | ~~TLN | -0.23 | - | 223 | -3.52 | < 0.001 | -0.23 | *** |
| ~~Dwroot | ~~Dwstem | 0.38 | - | 223 | 6.03 | < 0.001 | 0.38 | *** |
| ~~Dwroot | ~~Dwleaf | 0.27 | - | 223 | 4.13 | < 0.001 | 0.27 | *** |

**Table S10. Admixture-mapping outliers and their effects on phenotypic principal components.** For each SNP: chromosome, position, nearest gene and distance, effect sizes (β_PC1–β_PC4), SE, raw and adjusted P-values, and AIM/Δ p metadata.

| Chromosome | Position | P-value | adjP | beta_PC1 | beta_PC2 | beta_PC3 | beta_PC4 |
|---|---|---|---|---|---|---|---|
| 6 | 11740457 | 5.24E-18 | 1.90E-11 | -1.06 | 0.90 | 0.85 | -1.01 |
| 6 | 35536448 | 1.07E-14 | 8.53E-09 | 0.43 | -0.78 | -0.65 | -0.43 |
| 6 | 35539387 | 1.07E-14 | 8.53E-09 | 0.43 | -0.78 | -0.65 | -0.43 |
| 6 | 35541709 | 1.07E-14 | 8.53E-09 | 0.43 | -0.78 | -0.65 | -0.43 |
| 6 | 35540141 | 1.18E-14 | 8.53E-09 | 0.43 | -0.64 | -0.68 | -0.49 |
| 6 | 35535831 | 6.16E-13 | 3.72E-07 | 0.44 | -0.71 | -0.60 | -0.38 |
| 6 | 35541893 | 7.16E-12 | 3.51E-06 | 0.52 | -0.83 | -0.61 | -0.32 |
| 6 | 35535927 | 7.75E-12 | 3.51E-06 | 0.46 | -0.71 | -0.54 | -0.38 |
| 6 | 15366169 | 1.17E-11 | 4.71E-06 | 0.58 | -0.62 | -0.71 | 0.28 |
| 5 | 15419533 | 1.89E-11 | 6.84E-06 | 1.08 | -0.52 | -0.69 | 0.36 |
| 5 | 15426263 | 2.91E-11 | 7.44E-06 | 1.12 | -0.52 | -0.65 | 0.35 |
| 5 | 15422584 | 3.49E-11 | 7.44E-06 | 1.12 | -0.53 | -0.64 | 0.36 |
| 5 | 15424574 | 3.49E-11 | 7.44E-06 | 1.12 | -0.53 | -0.64 | 0.36 |
| 5 | 15426581 | 3.49E-11 | 7.44E-06 | 1.12 | -0.53 | -0.64 | 0.36 |
| 5 | 15429168 | 3.49E-11 | 7.44E-06 | 1.12 | -0.53 | -0.64 | 0.36 |
| 5 | 15430309 | 3.49E-11 | 7.44E-06 | 1.12 | -0.53 | -0.64 | 0.36 |
| 5 | 15430773 | 3.49E-11 | 7.44E-06 | 1.12 | -0.53 | -0.64 | 0.36 |
| 6 | 35546716 | 4.49E-11 | 9.04E-06 | 0.06 | -0.87 | -0.62 | -0.38 |
| 6 | 15528826 | 5.88E-11 | 1.12E-05 | -0.45 | -1.03 | -0.77 | 0.31 |
| 6 | 34960106 | 6.78E-11 | 1.20E-05 | 1.56 | -1.13 | -0.56 | -0.69 |
| 2 | 21687220 | 6.92E-11 | 1.20E-05 | -0.27 | -1.10 | -0.68 | 0.26 |
| 11 | 22078682 | 8.57E-11 | 1.41E-05 | 0.53 | -0.61 | -0.46 | -0.61 |
| 6 | 35536219 | 1.12E-10 | 1.76E-05 | 0.40 | -0.66 | -0.54 | -0.36 |
| 6 | 35532093 | 1.20E-10 | 1.81E-05 | 0.50 | -0.66 | -0.52 | -0.37 |
| 5 | 25777454 | 1.53E-10 | 2.23E-05 | -0.73 | -0.73 | 0.09 | 0.25 |
| 6 | 44822795 | 1.96E-10 | 2.40E-05 | 0.85 | -0.88 | -0.81 | -0.01 |
| 6 | 44823661 | 1.96E-10 | 2.40E-05 | 0.85 | -0.88 | -0.81 | -0.01 |
| 6 | 44824354 | 1.96E-10 | 2.40E-05 | 0.85 | -0.88 | -0.81 | -0.01 |
| 6 | 44827841 | 1.96E-10 | 2.40E-05 | 0.85 | -0.88 | -0.81 | -0.01 |
| 5 | 38453386 | 1.98E-10 | 2.40E-05 | 1.01 | -0.56 | 0.11 | -0.55 |
| 5 | 38452628 | 2.23E-10 | 2.60E-05 | 1.11 | -0.50 | 0.18 | -0.61 |
| 11 | 52568109 | 2.98E-10 | 3.37E-05 | 1.24 | -0.17 | -0.35 | -0.20 |
| 6 | 44824050 | 3.62E-10 | 3.98E-05 | 0.81 | -0.88 | -0.79 | -0.03 |
| 8 | 65016216 | 5.48E-10 | 5.85E-05 | -0.85 | -0.60 | -0.10 | 0.27 |
| 6 | 44826105 | 5.67E-10 | 5.87E-05 | 0.84 | -0.81 | -0.84 | -0.05 |
| 8 | 56396065 | 6.77E-10 | 6.82E-05 | 0.30 | -0.52 | -0.65 | -0.35 |
| 1 | 17397098 | 7.49E-10 | 7.33E-05 | -0.84 | -0.70 | 0.09 | 0.05 |
| 6 | 17278852 | 7.98E-10 | 7.33E-05 | 0.37 | -0.39 | -0.90 | -0.14 |
| 9 | 27308111 | 7.99E-10 | 7.33E-05 | 0.37 | 0.23 | -0.41 | 0.55 |
| 11 | 52568011 | 8.09E-10 | 7.33E-05 | 1.17 | -0.28 | -0.35 | -0.13 |
| 10 | 20455066 | 8.36E-10 | 7.40E-05 | 0.79 | -0.76 | -0.05 | 0.31 |
| 4 | 50749128 | 1.13E-09 | 9.77E-05 | 0.05 | -0.47 | -0.67 | 0.27 |
| 8 | 21359259 | 1.22E-09 | 1.03E-04 | 1.64 | -0.77 | -0.02 | 0.54 |
| 12 | 455035 | 1.85E-09 | 1.47E-04 | 0.18 | -0.05 | -0.74 | -0.41 |
| 6 | 44819224 | 1.86E-09 | 1.47E-04 | 0.95 | -0.94 | -0.73 | -0.02 |
| 6 | 44819465 | 1.86E-09 | 1.47E-04 | 0.95 | -0.94 | -0.73 | -0.02 |
| 6 | 44822088 | 2.24E-09 | 1.73E-04 | 0.77 | -0.83 | -0.81 | -0.17 |

| | | | | | | | |
|---|---|---|---|---|---|---|---|
| 2 | 21348415 | 3.27E-09 | 2.47E-04 | 0.66 | -0.54 | -0.09 | -1.01 |
| 6 | 28426135 | 3.76E-09 | 2.73E-04 | -0.74 | -0.23 | 0.19 | 0.59 |
| 6 | 28428087 | 3.76E-09 | 2.73E-04 | -0.74 | -0.23 | 0.19 | 0.59 |
| 5 | 15437233 | 3.95E-09 | 2.81E-04 | 0.97 | -0.47 | -0.61 | 0.34 |
| 6 | 10836347 | 4.08E-09 | 2.85E-04 | -0.44 | -0.20 | -0.76 | -0.21 |
| 4 | 53013211 | 4.22E-09 | 2.89E-04 | 0.82 | -0.49 | -0.54 | -0.18 |
| 11 | 42093191 | 4.91E-09 | 3.29E-04 | 1.18 | -0.40 | -0.55 | -0.53 |
| 7 | 30507049 | 4.99E-09 | 3.29E-04 | 0.38 | -0.06 | 0.21 | 0.71 |
| 10 | 20456952 | 5.24E-09 | 3.39E-04 | 0.90 | -0.69 | -0.08 | 0.16 |
| 6 | 36371408 | 5.74E-09 | 3.42E-04 | 1.33 | -0.38 | 0.08 | -0.21 |
| 6 | 36371739 | 5.74E-09 | 3.42E-04 | 1.33 | -0.38 | 0.08 | -0.21 |
| 6 | 36375311 | 5.74E-09 | 3.42E-04 | 1.33 | -0.38 | 0.08 | -0.21 |
| 6 | 36376093 | 5.74E-09 | 3.42E-04 | 1.33 | -0.38 | 0.08 | -0.21 |
| 8 | 21413526 | 5.78E-09 | 3.42E-04 | 0.81 | -0.47 | 0.34 | 0.29 |
| 12 | 460072 | 5.85E-09 | 3.42E-04 | 0.31 | -0.18 | -0.66 | -0.37 |
| 5 | 15430089 | 6.40E-09 | 3.68E-04 | 0.99 | -0.43 | -0.65 | 0.24 |
| 5 | 38441970 | 6.93E-09 | 3.93E-04 | 0.77 | -0.76 | 0.08 | -0.35 |
| 5 | 25775843 | 7.45E-09 | 4.16E-04 | -0.68 | -0.67 | 0.00 | 0.29 |
| 5 | 20581274 | 7.60E-09 | 4.18E-04 | 1.00 | -0.36 | -0.41 | -0.64 |
| 1 | 7046665 | 7.79E-09 | 4.22E-04 | -0.39 | 0.04 | -0.50 | 0.59 |
| 6 | 14707773 | 9.04E-09 | 4.82E-04 | 0.40 | -0.68 | -0.34 | -0.42 |
| 8 | 47804209 | 9.34E-09 | 4.91E-04 | 0.81 | -0.47 | 0.72 | -0.13 |
| 2 | 21541564 | 9.57E-09 | 4.96E-04 | -1.09 | -0.18 | 0.20 | 0.06 |
| 2 | 10864656 | 1.06E-08 | 5.43E-04 | -0.61 | -0.57 | 0.18 | -0.29 |
| 6 | 44819029 | 1.18E-08 | 5.92E-04 | 0.66 | -0.81 | -0.77 | 0.08 |
| 12 | 16628834 | 1.19E-08 | 5.92E-04 | -0.33 | -0.77 | 0.43 | -0.12 |
| 11 | 22084982 | 1.24E-08 | 6.09E-04 | 0.54 | -0.62 | -0.41 | -0.35 |
| 1 | 7604547 | 1.31E-08 | 6.35E-04 | -0.40 | 0.24 | -0.99 | 0.83 |
| 6 | 28428136 | 1.37E-08 | 6.49E-04 | -0.65 | -0.18 | 0.23 | 0.63 |
| 6 | 6368586 | 1.38E-08 | 6.49E-04 | 0.64 | -0.57 | -0.58 | -0.20 |
| 11 | 22079270 | 1.41E-08 | 6.57E-04 | 0.58 | -0.49 | -0.40 | -0.52 |
| 1 | 39718777 | 1.54E-08 | 7.06E-04 | 0.89 | -1.24 | 0.55 | 0.74 |
| 8 | 56013898 | 1.56E-08 | 7.07E-04 | 0.11 | 0.32 | -0.80 | 0.04 |
| 5 | 38441924 | 1.75E-08 | 7.82E-04 | 0.73 | -0.74 | 0.11 | -0.36 |
| 7 | 26081948 | 1.93E-08 | 8.53E-04 | -0.22 | -1.06 | -0.75 | 0.35 |
| 11 | 51865677 | 2.06E-08 | 8.68E-04 | 0.02 | 0.29 | -0.38 | 0.59 |
| 2 | 26700636 | 2.10E-08 | 8.68E-04 | -0.22 | -0.17 | -0.63 | -0.36 |
| 12 | 16248480 | 2.11E-08 | 8.68E-04 | -0.79 | -0.39 | 0.42 | 0.21 |
| 12 | 453897 | 2.12E-08 | 8.68E-04 | 0.27 | -0.16 | -0.65 | -0.38 |
| 12 | 456299 | 2.12E-08 | 8.68E-04 | 0.27 | -0.16 | -0.65 | -0.38 |
| 12 | 456656 | 2.12E-08 | 8.68E-04 | 0.27 | -0.16 | -0.65 | -0.38 |
| 2 | 26660703 | 2.13E-08 | 8.68E-04 | 1.02 | -0.05 | -0.56 | -0.31 |
| 12 | 11986094 | 2.29E-08 | 9.23E-04 | 0.76 | 0.13 | -0.55 | -0.34 |
| 6 | 36372679 | 2.32E-08 | 9.23E-04 | 1.26 | -0.38 | 0.08 | -0.20 |
| 5 | 25760628 | 2.52E-08 | 9.90E-04 | -0.55 | -0.79 | -0.19 | 0.28 |
| 11 | 52549682 | 2.54E-08 | 9.90E-04 | 1.05 | -0.42 | -0.45 | -0.13 |
| 2 | 6721135 | 2.64E-08 | 1.02E-03 | 0.67 | 0.63 | -0.36 | -0.18 |
| 2 | 6721168 | 2.70E-08 | 1.03E-03 | 0.69 | 0.62 | -0.35 | -0.19 |
| 1 | 39718783 | 2.76E-08 | 1.04E-03 | 0.84 | -1.10 | 0.54 | 0.84 |
| 1 | 13328475 | 2.80E-08 | 1.04E-03 | 0.38 | -0.61 | -0.05 | 0.45 |
| 5 | 25748513 | 2.81E-08 | 1.04E-03 | -0.60 | -0.80 | -0.16 | 0.21 |
| 4 | 73006761 | 2.94E-08 | 1.08E-03 | 1.08 | -0.32 | 0.04 | 0.18 |

| | | | | | | | |
|---|---|---|---|---|---|---|---|
| 7 | 31085399 | 3.01E-08 | 1.09E-03 | -0.14 | -0.29 | 0.05 | 0.82 |
| 7 | 31086567 | 3.30E-08 | 1.19E-03 | -0.14 | -0.30 | 0.03 | 0.80 |
| 6 | 36370018 | 3.44E-08 | 1.22E-03 | 1.09 | -0.50 | 0.07 | -0.17 |
| 6 | 27593897 | 3.54E-08 | 1.25E-03 | 0.23 | 0.12 | -0.09 | 0.89 |
| 6 | 15393348 | 3.77E-08 | 1.31E-03 | 0.29 | -0.39 | -0.28 | 0.54 |
| 1 | 37449579 | 4.12E-08 | 1.41E-03 | -0.16 | -0.70 | -0.36 | -0.33 |
| 2 | 10864928 | 4.12E-08 | 1.41E-03 | -0.69 | -0.55 | 0.08 | -0.22 |
| 1 | 13318361 | 4.35E-08 | 1.48E-03 | 0.71 | -0.44 | 0.08 | 0.46 |
| 12 | 16246698 | 4.69E-08 | 1.57E-03 | -0.83 | -0.33 | 0.36 | 0.25 |
| 5 | 25778633 | 4.73E-08 | 1.57E-03 | -0.51 | -0.83 | -0.12 | 0.21 |
| 6 | 14706496 | 4.91E-08 | 1.60E-03 | 0.41 | -0.66 | -0.37 | -0.37 |
| 6 | 14708728 | 4.91E-08 | 1.60E-03 | 0.41 | -0.66 | -0.37 | -0.37 |
| 8 | 21413778 | 5.12E-08 | 1.65E-03 | 0.70 | -0.51 | 0.20 | 0.27 |
| 12 | 16250851 | 5.16E-08 | 1.65E-03 | -0.74 | -0.38 | 0.38 | 0.29 |
| 7 | 16657439 | 5.22E-08 | 1.66E-03 | 0.33 | 0.70 | -0.48 | -0.06 |
| 10 | 6698239 | 5.40E-08 | 1.70E-03 | 1.13 | 0.13 | -0.35 | 0.03 |
| 6 | 44815176 | 5.48E-08 | 1.71E-03 | 1.20 | -0.89 | -0.75 | -0.08 |
| 12 | 452523 | 5.74E-08 | 1.75E-03 | 0.30 | -0.12 | -0.66 | -0.37 |
| 6 | 28429452 | 5.75E-08 | 1.75E-03 | -0.93 | -0.10 | 0.38 | 0.79 |
| 6 | 28429564 | 5.75E-08 | 1.75E-03 | -0.93 | -0.10 | 0.38 | 0.79 |
| 6 | 45022050 | 5.80E-08 | 1.75E-03 | 0.60 | -0.64 | -0.74 | -0.30 |
| 11 | 22089680 | 6.02E-08 | 1.81E-03 | 0.40 | -0.59 | -0.41 | -0.39 |
| 8 | 21413525 | 7.07E-08 | 2.08E-03 | 0.75 | -0.46 | 0.24 | 0.26 |
| 8 | 21414689 | 7.07E-08 | 2.08E-03 | 0.75 | -0.46 | 0.24 | 0.26 |
| 11 | 22090250 | 7.10E-08 | 2.08E-03 | 0.62 | -0.52 | -0.38 | -0.39 |
| 2 | 9520676 | 7.22E-08 | 2.10E-03 | 0.19 | -0.46 | -0.70 | 0.10 |
| 11 | 22079232 | 7.44E-08 | 2.14E-03 | 0.63 | -0.47 | -0.39 | -0.46 |
| 6 | 36373564 | 7.85E-08 | 2.23E-03 | 1.24 | -0.34 | 0.08 | -0.20 |
| 2 | 9512244 | 7.89E-08 | 2.23E-03 | 0.06 | -0.56 | -0.73 | 0.14 |
| 6 | 35523013 | 8.26E-08 | 2.32E-03 | 0.56 | -1.02 | -0.22 | -0.18 |
| 11 | 22090621 | 8.46E-08 | 2.35E-03 | 0.56 | -0.58 | -0.34 | -0.43 |
| 1 | 13307385 | 8.60E-08 | 2.35E-03 | 0.68 | -0.54 | 0.00 | 0.36 |
| 1 | 13315644 | 8.60E-08 | 2.35E-03 | 0.68 | -0.54 | 0.00 | 0.36 |
| 1 | 13316191 | 8.60E-08 | 2.35E-03 | 0.68 | -0.54 | 0.00 | 0.36 |
| 4 | 11860146 | 8.73E-08 | 2.36E-03 | -0.37 | -0.38 | -0.07 | -0.58 |
| 11 | 22090200 | 8.86E-08 | 2.38E-03 | 0.54 | -0.52 | -0.45 | -0.31 |
| 5 | 15420257 | 8.98E-08 | 2.39E-03 | 1.05 | -0.52 | -0.60 | 0.27 |
| 12 | 16257780 | 9.08E-08 | 2.40E-03 | -0.85 | -0.29 | 0.20 | 0.32 |
| 7 | 28701458 | 9.25E-08 | 2.43E-03 | 0.69 | -1.17 | -0.62 | -0.65 |
| 5 | 58087753 | 9.53E-08 | 2.48E-03 | -1.03 | 0.10 | 0.03 | 0.48 |
| 11 | 22084126 | 9.77E-08 | 2.48E-03 | 0.51 | -0.60 | -0.42 | -0.32 |
| 11 | 22090512 | 9.77E-08 | 2.48E-03 | 0.51 | -0.60 | -0.42 | -0.32 |
| 11 | 22090952 | 9.77E-08 | 2.48E-03 | 0.51 | -0.60 | -0.42 | -0.32 |
| 11 | 22091188 | 9.77E-08 | 2.48E-03 | 0.51 | -0.60 | -0.42 | -0.32 |
| 11 | 34818472 | 1.01E-07 | 2.54E-03 | -0.12 | -0.39 | -0.28 | 0.51 |
| 10 | 2364561 | 1.07E-07 | 2.68E-03 | 0.25 | -0.82 | -0.57 | 0.03 |
| 12 | 456474 | 1.10E-07 | 2.72E-03 | 0.26 | -0.13 | -0.63 | -0.37 |
| 7 | 31083357 | 1.15E-07 | 2.83E-03 | -0.08 | -0.25 | -0.04 | 0.77 |
| 1 | 13318367 | 1.16E-07 | 2.84E-03 | 0.71 | -0.37 | 0.19 | 0.48 |
| 7 | 31081232 | 1.20E-07 | 2.92E-03 | -0.16 | -0.25 | -0.07 | 0.78 |
| 6 | 45018126 | 1.29E-07 | 3.11E-03 | 0.54 | -0.62 | -0.75 | -0.29 |
| 6 | 44825264 | 1.31E-07 | 3.14E-03 | 0.56 | -0.78 | -0.66 | 0.02 |

| | | | | | | | |
|---|---|---|---|---|---|---|---|
| 2 | 17822264 | 1.35E-07 | 3.22E-03 | 0.05 | -0.62 | -0.87 | -0.74 |
| 4 | 73067497 | 1.40E-07 | 3.32E-03 | 1.62 | -1.30 | 0.22 | -0.88 |
| 2 | 15108070 | 1.45E-07 | 3.41E-03 | 0.62 | -0.97 | -0.25 | -0.25 |
| 12 | 448309 | 1.47E-07 | 3.44E-03 | 0.32 | -0.03 | -0.64 | -0.40 |
| 8 | 22402286 | 1.51E-07 | 3.48E-03 | -0.75 | 0.34 | -0.17 | 0.43 |
| 8 | 22402288 | 1.51E-07 | 3.48E-03 | -0.75 | 0.34 | -0.17 | 0.43 |
| 6 | 21651174 | 1.61E-07 | 3.68E-03 | 1.14 | -0.59 | -0.40 | -0.28 |
| 6 | 42332342 | 1.62E-07 | 3.68E-03 | 0.25 | -0.72 | -0.41 | -0.76 |
| 6 | 21651176 | 1.62E-07 | 3.68E-03 | 1.14 | -0.59 | -0.39 | -0.28 |
| 12 | 454041 | 1.69E-07 | 3.75E-03 | 0.19 | -0.17 | -0.63 | -0.34 |
| 6 | 44803510 | 1.70E-07 | 3.75E-03 | 1.15 | -0.87 | -0.66 | -0.15 |
| 6 | 44806928 | 1.70E-07 | 3.75E-03 | 1.15 | -0.87 | -0.66 | -0.15 |
| 12 | 16255212 | 1.71E-07 | 3.75E-03 | -0.67 | -0.37 | 0.38 | 0.28 |
| 12 | 16255665 | 1.71E-07 | 3.75E-03 | -0.67 | -0.37 | 0.38 | 0.28 |
| 2 | 6718124 | 1.82E-07 | 3.94E-03 | 0.69 | 0.62 | -0.29 | -0.18 |
| 2 | 6718178 | 1.82E-07 | 3.94E-03 | 0.69 | 0.62 | -0.29 | -0.18 |
| 5 | 20581210 | 1.82E-07 | 3.94E-03 | 0.98 | -0.35 | -0.45 | -0.53 |
| 5 | 25752528 | 1.87E-07 | 4.01E-03 | -0.65 | -0.57 | 0.06 | 0.21 |
| 1 | 40308195 | 1.90E-07 | 4.06E-03 | 0.12 | -0.08 | 0.24 | 0.86 |
| 6 | 35551456 | 1.99E-07 | 4.17E-03 | 0.30 | -0.82 | -0.52 | -0.16 |
| 6 | 35552296 | 1.99E-07 | 4.17E-03 | 0.30 | -0.82 | -0.52 | -0.16 |
| 12 | 13530893 | 1.99E-07 | 4.17E-03 | -1.11 | 0.21 | 0.55 | 0.46 |
| 12 | 16246554 | 2.08E-07 | 4.33E-03 | -0.65 | -0.38 | 0.37 | 0.29 |
| 6 | 36356963 | 2.16E-07 | 4.48E-03 | 0.95 | -0.46 | -0.27 | -0.42 |
| 10 | 8071647 | 2.19E-07 | 4.51E-03 | 0.33 | -0.46 | 0.05 | 0.56 |
| 1 | 37438209 | 2.22E-07 | 4.55E-03 | 0.23 | -0.54 | -0.27 | -0.52 |
| 6 | 15215091 | 2.25E-07 | 4.56E-03 | -0.24 | -1.05 | -0.72 | 0.30 |
| 6 | 44760681 | 2.25E-07 | 4.56E-03 | 0.48 | -0.88 | -0.50 | 0.19 |
| 4 | 19537314 | 2.30E-07 | 4.63E-03 | 2.08 | 0.17 | -0.84 | -0.18 |
| 5 | 58088839 | 2.32E-07 | 4.64E-03 | -1.00 | -0.10 | 0.03 | 0.47 |
| 6 | 15302841 | 2.41E-07 | 4.75E-03 | 0.53 | -0.55 | -0.39 | 0.32 |
| 8 | 16388685 | 2.41E-07 | 4.75E-03 | -0.67 | -0.23 | -0.04 | 0.50 |
| 8 | 16390057 | 2.41E-07 | 4.75E-03 | -0.67 | -0.23 | -0.04 | 0.50 |
| 11 | 22088211 | 2.42E-07 | 4.75E-03 | 0.42 | -0.56 | -0.39 | -0.37 |
| 2 | 6725695 | 2.53E-07 | 4.90E-03 | 0.70 | 0.58 | -0.29 | -0.19 |
| 12 | 16249430 | 2.57E-07 | 4.90E-03 | -0.67 | -0.35 | 0.37 | 0.28 |
| 12 | 16252243 | 2.57E-07 | 4.90E-03 | -0.67 | -0.35 | 0.37 | 0.28 |
| 12 | 16252391 | 2.57E-07 | 4.90E-03 | -0.67 | -0.35 | 0.37 | 0.28 |
| 12 | 16252408 | 2.57E-07 | 4.90E-03 | -0.67 | -0.35 | 0.37 | 0.28 |
| 6 | 15276953 | 2.58E-07 | 4.90E-03 | 0.63 | -0.54 | -0.23 | 0.33 |
| 10 | 10595127 | 2.80E-07 | 5.22E-03 | 0.32 | -0.79 | -0.24 | -0.34 |
| 11 | 52549034 | 2.81E-07 | 5.22E-03 | 1.06 | -0.36 | -0.40 | -0.16 |
| 11 | 52549733 | 2.81E-07 | 5.22E-03 | 1.06 | -0.36 | -0.40 | -0.16 |
| 11 | 52549987 | 2.81E-07 | 5.22E-03 | 1.06 | -0.36 | -0.40 | -0.16 |
| 6 | 49814450 | 2.90E-07 | 5.36E-03 | 1.08 | -0.09 | -0.12 | 0.14 |
| 6 | 17278714 | 2.95E-07 | 5.43E-03 | 0.43 | -0.39 | -0.68 | -0.21 |
| 5 | 38370662 | 2.98E-07 | 5.45E-03 | 0.86 | -0.41 | 0.04 | -0.41 |
| 2 | 18181622 | 3.02E-07 | 5.51E-03 | -0.65 | -0.10 | -0.73 | -0.88 |
| 9 | 36323123 | 3.07E-07 | 5.55E-03 | 0.99 | -0.09 | -0.43 | 0.84 |
| 6 | 44487306 | 3.16E-07 | 5.55E-03 | 1.40 | -0.72 | -0.71 | -0.63 |
| 6 | 14313935 | 3.17E-07 | 5.55E-03 | 0.12 | -0.23 | -0.50 | -0.59 |
| 2 | 18188992 | 3.20E-07 | 5.55E-03 | -0.74 | -0.09 | -0.79 | -0.85 |

| | | | | | | | |
|---|---|---|---|---|---|---|---|
| 2 | 18189817 | 3.20E-07 | 5.55E-03 | -0.74 | -0.09 | -0.79 | -0.85 |
| 2 | 18189922 | 3.20E-07 | 5.55E-03 | -0.74 | -0.09 | -0.79 | -0.85 |
| 2 | 18190175 | 3.20E-07 | 5.55E-03 | -0.74 | -0.09 | -0.79 | -0.85 |
| 2 | 18190390 | 3.20E-07 | 5.55E-03 | -0.74 | -0.09 | -0.79 | -0.85 |
| 2 | 18195470 | 3.20E-07 | 5.55E-03 | -0.74 | -0.09 | -0.79 | -0.85 |
| 2 | 18195632 | 3.20E-07 | 5.55E-03 | -0.74 | -0.09 | -0.79 | -0.85 |
| 6 | 18822127 | 3.24E-07 | 5.60E-03 | 0.09 | -0.76 | -0.56 | 0.29 |
| 2 | 6720784 | 3.36E-07 | 5.78E-03 | 0.68 | 0.58 | -0.32 | -0.18 |
| 4 | 32841519 | 3.48E-07 | 5.96E-03 | -0.89 | -0.34 | -0.04 | 0.26 |
| 9 | 30511904 | 3.53E-07 | 5.98E-03 | 0.84 | -1.19 | -0.15 | -0.09 |
| 9 | 24641064 | 3.53E-07 | 5.98E-03 | 0.92 | -0.41 | -0.16 | 0.01 |
| 10 | 24306173 | 3.55E-07 | 5.99E-03 | -0.78 | -0.43 | 0.33 | 0.28 |
| 7 | 16614036 | 3.58E-07 | 6.02E-03 | -0.55 | -0.53 | 0.14 | 0.27 |
| 12 | 16249770 | 3.66E-07 | 6.11E-03 | -0.67 | -0.30 | 0.36 | 0.33 |
| 1 | 13311449 | 3.72E-07 | 6.17E-03 | 0.73 | -0.38 | 0.17 | 0.41 |
| 10 | 2396133 | 3.75E-07 | 6.17E-03 | 0.19 | -0.78 | 0.11 | -0.16 |
| 10 | 2396136 | 3.75E-07 | 6.17E-03 | 0.19 | -0.78 | 0.11 | -0.16 |
| 6 | 28450833 | 3.76E-07 | 6.17E-03 | 0.11 | 0.24 | -0.37 | 0.48 |
| 11 | 51859763 | 3.82E-07 | 6.21E-03 | 0.09 | 0.29 | -0.57 | 0.41 |
| 12 | 16247621 | 3.82E-07 | 6.21E-03 | -0.71 | -0.37 | 0.37 | 0.21 |
| 5 | 66743794 | 3.86E-07 | 6.26E-03 | -0.34 | -1.04 | -0.71 | 0.31 |
| 9 | 20436749 | 3.95E-07 | 6.33E-03 | -1.02 | -0.10 | -0.01 | 0.37 |
| 2 | 6692073 | 3.97E-07 | 6.33E-03 | 0.66 | 0.60 | -0.35 | -0.09 |
| 8 | 56013925 | 3.98E-07 | 6.33E-03 | 0.17 | 0.23 | -0.72 | 0.07 |
| 9 | 45523800 | 3.98E-07 | 6.33E-03 | 0.12 | -0.82 | -0.60 | 0.03 |
| 2 | 6683640 | 4.06E-07 | 6.42E-03 | 0.71 | 0.58 | -0.33 | -0.06 |
| 2 | 25604254 | 4.07E-07 | 6.42E-03 | -0.95 | 0.19 | -0.36 | -0.29 |
| 12 | 16254101 | 4.40E-07 | 6.90E-03 | -0.66 | -0.39 | 0.37 | 0.25 |
| 6 | 36484967 | 4.48E-07 | 7.00E-03 | 0.76 | -0.55 | -0.36 | -0.29 |
| 4 | 30460221 | 4.53E-07 | 7.01E-03 | 1.99 | 0.44 | -0.03 | -0.83 |
| 6 | 36360632 | 4.54E-07 | 7.01E-03 | 0.87 | -0.38 | -0.03 | -0.55 |
| 6 | 36363630 | 4.54E-07 | 7.01E-03 | 0.87 | -0.38 | -0.03 | -0.55 |
| 10 | 16079961 | 4.67E-07 | 7.18E-03 | -0.17 | -0.08 | -0.97 | -0.98 |
| 11 | 52546200 | 4.88E-07 | 7.40E-03 | 1.07 | -0.25 | -0.45 | -0.14 |
| 11 | 52546284 | 4.88E-07 | 7.40E-03 | 1.07 | -0.25 | -0.45 | -0.14 |
| 11 | 52566633 | 4.88E-07 | 7.40E-03 | 1.07 | -0.25 | -0.45 | -0.14 |
| 9 | 7860839 | 4.92E-07 | 7.44E-03 | -0.68 | 0.10 | -0.42 | -0.44 |
| 8 | 37692269 | 4.96E-07 | 7.46E-03 | 0.23 | -0.12 | -0.34 | 0.49 |
| 11 | 41931333 | 5.00E-07 | 7.49E-03 | 1.41 | 0.03 | 0.30 | -0.11 |
| 6 | 36371259 | 5.16E-07 | 7.69E-03 | 1.10 | -0.40 | 0.17 | -0.26 |
| 8 | 10689674 | 5.24E-07 | 7.78E-03 | 0.10 | 0.19 | -0.57 | -0.53 |
| 5 | 25872417 | 5.29E-07 | 7.83E-03 | 1.14 | 0.34 | -0.19 | 0.00 |
| 1 | 22001064 | 5.36E-07 | 7.87E-03 | 0.60 | -0.38 | -0.55 | -0.41 |
| 1 | 22001285 | 5.36E-07 | 7.87E-03 | 0.60 | -0.38 | -0.55 | -0.41 |
| 1 | 8467126 | 5.44E-07 | 7.96E-03 | 1.46 | 0.40 | -1.26 | 0.54 |
| 10 | 8692835 | 5.53E-07 | 8.05E-03 | 0.80 | 1.08 | -0.96 | 0.41 |
| 8 | 49052349 | 5.64E-07 | 8.19E-03 | 0.41 | -0.30 | -0.68 | -0.14 |
| 6 | 35543933 | 5.71E-07 | 8.25E-03 | 0.21 | -0.72 | -0.47 | -0.29 |
| 9 | 39334998 | 5.74E-07 | 8.25E-03 | 1.17 | -0.40 | -0.47 | -0.16 |
| 8 | 16388871 | 5.80E-07 | 8.28E-03 | -0.48 | -0.33 | -0.05 | 0.54 |
| 1 | 37429066 | 5.80E-07 | 8.28E-03 | -0.12 | -0.61 | -0.33 | -0.37 |
| 7 | 49253245 | 5.91E-07 | 8.39E-03 | 0.94 | -0.59 | -0.39 | -0.40 |

| | | | | | | | |
|---|---|---|---|---|---|---|---|
| 2 | 26648099 | 5.93E-07 | 8.39E-03 | 0.88 | -0.01 | -0.46 | -0.37 |
| 7 | 27143144 | 6.01E-07 | 8.48E-03 | 0.23 | -0.60 | -0.60 | 0.18 |
| 5 | 38447677 | 6.13E-07 | 8.61E-03 | 0.80 | -0.47 | 0.18 | -0.43 |
| 6 | 44488595 | 6.15E-07 | 8.61E-03 | 0.82 | -0.82 | -0.32 | -0.54 |
| 8 | 22403961 | 6.33E-07 | 8.83E-03 | -0.20 | 0.15 | -0.16 | 0.76 |
| 12 | 16255022 | 6.45E-07 | 8.97E-03 | -0.70 | -0.38 | 0.34 | 0.24 |
| 2 | 10147473 | 6.63E-07 | 9.17E-03 | -1.06 | 0.18 | 0.19 | 0.62 |
| 9 | 29169195 | 6.71E-07 | 9.21E-03 | 0.90 | -0.31 | 0.00 | 0.17 |
| 9 | 29169206 | 6.71E-07 | 9.21E-03 | 0.90 | -0.31 | 0.00 | 0.17 |
| 4 | 22380900 | 6.81E-07 | 9.32E-03 | 0.12 | -0.42 | 0.20 | 1.46 |
| 1 | 40312803 | 6.85E-07 | 9.33E-03 | 0.21 | -0.13 | 0.19 | 0.75 |
| 6 | 380367 | 7.15E-07 | 9.71E-03 | 0.87 | -0.49 | -0.31 | 0.87 |
| 7 | 16610781 | 7.22E-07 | 9.73E-03 | -0.53 | -0.63 | 0.02 | 0.22 |
| 12 | 454258 | 7.23E-07 | 9.73E-03 | 0.28 | 0.05 | -0.61 | -0.42 |
| 1 | 37436922 | 7.24E-07 | 9.73E-03 | 0.00 | -0.67 | -0.31 | -0.31 |
| 2 | 10147013 | 7.39E-07 | 9.79E-03 | -1.14 | 0.14 | 0.20 | 0.57 |
| 2 | 21670608 | 7.41E-07 | 9.79E-03 | 0.05 | -0.59 | -0.60 | 0.27 |
| 2 | 15709538 | 7.43E-07 | 9.79E-03 | -0.32 | -0.60 | 0.11 | -0.46 |
| 10 | 6794984 | 7.45E-07 | 9.79E-03 | 0.17 | -0.59 | -0.53 | 0.30 |
| 10 | 6794993 | 7.45E-07 | 9.79E-03 | 0.17 | -0.59 | -0.53 | 0.30 |
| 6 | 11593747 | 7.45E-07 | 9.79E-03 | 0.73 | -0.16 | -0.25 | 0.57 |
| 1 | 38318860 | 7.57E-07 | 9.87E-03 | 0.59 | -0.78 | -0.02 | -0.10 |
| 5 | 897615 | 7.57E-07 | 9.87E-03 | -0.01 | -0.82 | -0.41 | -1.28 |
| 1 | 40302374 | 7.66E-07 | 9.95E-03 | 0.07 | -0.07 | 0.24 | 0.82 |
| 6 | 14705106 | 7.73E-07 | 9.95E-03 | 0.40 | -0.63 | -0.48 | -0.12 |
| 6 | 14713585 | 7.73E-07 | 9.95E-03 | 0.40 | -0.63 | -0.48 | -0.12 |
| 6 | 28194590 | 7.74E-07 | 9.95E-03 | 0.87 | -0.59 | -0.50 | -0.29 |
| 10 | 5935824 | 7.84E-07 | 9.96E-03 | 0.65 | -0.39 | -0.40 | -0.06 |
| 8 | 9271944 | 7.85E-07 | 9.96E-03 | 0.19 | -0.73 | 0.37 | 1.17 |
| 8 | 9272959 | 7.85E-07 | 9.96E-03 | 0.19 | -0.73 | 0.37 | 1.17 |
| 8 | 9293718 | 7.85E-07 | 9.96E-03 | 0.19 | -0.73 | 0.37 | 1.17 |
| 1 | 5109904 | 7.88E-07 | 9.96E-03 | 0.32 | -0.12 | -0.02 | 0.61 |
| 8 | 27307412 | 8.01E-07 | 1.00E-02 | 0.41 | 0.37 | -0.39 | -0.55 |
| 4 | 41243751 | 8.01E-07 | 1.00E-02 | -0.15 | -0.23 | -0.86 | -0.91 |
| 4 | 34902021 | 8.04E-07 | 1.01E-02 | 0.80 | -0.39 | -0.46 | -0.32 |
| 2 | 10864510 | 8.10E-07 | 1.01E-02 | -0.51 | -0.52 | 0.12 | -0.24 |
| 10 | 10574480 | 8.18E-07 | 1.02E-02 | 0.38 | -0.72 | -0.17 | -0.44 |
| 12 | 15705039 | 8.31E-07 | 1.03E-02 | -1.19 | 0.75 | 0.15 | 0.24 |
| 5 | 25746155 | 8.32E-07 | 1.03E-02 | -0.51 | -0.73 | -0.08 | 0.25 |
| 11 | 51859762 | 8.34E-07 | 1.03E-02 | 0.17 | 0.35 | -0.40 | 0.45 |
| 5 | 40589031 | 8.37E-07 | 1.03E-02 | 0.99 | -0.73 | -1.16 | -0.18 |
| 9 | 30309465 | 8.40E-07 | 1.03E-02 | -0.64 | 0.12 | -0.63 | -0.34 |
| 6 | 14708419 | 8.78E-07 | 1.06E-02 | 0.16 | -0.66 | -0.30 | -0.37 |
| 12 | 16249698 | 8.78E-07 | 1.06E-02 | -0.62 | -0.34 | 0.38 | 0.28 |
| 6 | 16370605 | 8.79E-07 | 1.06E-02 | 0.65 | -0.60 | -0.13 | -0.17 |
| 12 | 16253381 | 8.89E-07 | 1.07E-02 | -0.68 | -0.30 | 0.36 | 0.29 |
| 6 | 15266146 | 9.13E-07 | 1.08E-02 | 0.63 | -0.51 | -0.32 | 0.25 |
| 5 | 38457874 | 9.15E-07 | 1.08E-02 | 0.79 | -0.51 | 0.05 | -0.54 |
| 6 | 44487185 | 9.16E-07 | 1.08E-02 | 1.59 | -0.87 | -0.34 | -0.45 |
| 1 | 40295089 | 9.16E-07 | 1.08E-02 | 0.14 | -0.09 | 0.22 | 0.80 |
| 1 | 40300388 | 9.16E-07 | 1.08E-02 | 0.14 | -0.09 | 0.22 | 0.80 |
| 1 | 40304382 | 9.16E-07 | 1.08E-02 | 0.14 | -0.09 | 0.22 | 0.80 |

| | | | | | | | |
|---|---|---|---|---|---|---|---|
| 1 | 40337221 | 9.16E-07 | 1.08E-02 | 0.14 | -0.09 | 0.22 | 0.80 |
| 6 | 9920519 | 9.54E-07 | 1.12E-02 | 0.55 | -0.56 | -0.26 | 0.19 |
| 2 | 6719892 | 9.66E-07 | 1.13E-02 | 0.63 | 0.60 | -0.27 | -0.18 |
| 6 | 15275914 | 9.72E-07 | 1.13E-02 | 0.68 | -0.41 | -0.30 | 0.29 |
| 8 | 49051116 | 9.77E-07 | 1.14E-02 | 0.51 | -0.22 | -0.64 | -0.16 |
| 11 | 22079125 | 9.85E-07 | 1.14E-02 | 0.41 | -0.43 | -0.45 | -0.38 |
| 6 | 15198411 | 9.90E-07 | 1.14E-02 | -0.28 | -1.03 | -0.71 | 0.31 |
| 6 | 15203435 | 9.90E-07 | 1.14E-02 | -0.28 | -1.03 | -0.71 | 0.31 |
| 5 | 38370630 | 1.00E-06 | 1.15E-02 | 0.85 | -0.41 | 0.04 | -0.40 |
| 5 | 38370751 | 1.00E-06 | 1.15E-02 | 0.85 | -0.41 | 0.04 | -0.40 |
| 1 | 22013557 | 1.01E-06 | 1.15E-02 | 0.56 | -0.38 | -0.46 | -0.53 |
| 5 | 15416811 | 1.01E-06 | 1.15E-02 | 0.76 | -0.58 | -0.46 | 0.25 |
| 7 | 26104183 | 1.03E-06 | 1.16E-02 | 0.38 | -0.66 | -0.28 | 0.35 |
| 8 | 21410787 | 1.03E-06 | 1.16E-02 | 0.81 | -0.30 | 0.26 | 0.27 |
| 6 | 44508533 | 1.04E-06 | 1.16E-02 | 0.65 | -0.58 | -0.39 | -0.39 |
| 11 | 52551026 | 1.04E-06 | 1.16E-02 | 0.95 | -0.33 | -0.43 | -0.12 |
| 6 | 6370051 | 1.05E-06 | 1.17E-02 | 0.67 | -0.66 | -0.33 | -0.03 |
| 1 | 37453025 | 1.05E-06 | 1.17E-02 | -0.04 | -0.57 | -0.37 | -0.36 |
| 6 | 6361277 | 1.05E-06 | 1.17E-02 | 0.74 | -0.58 | -0.29 | -0.04 |
| 6 | 42350064 | 1.06E-06 | 1.17E-02 | 0.42 | -0.73 | -0.37 | -0.65 |
| 2 | 13505760 | 1.07E-06 | 1.17E-02 | -0.34 | -1.08 | -0.73 | 0.31 |
| 2 | 6693408 | 1.07E-06 | 1.17E-02 | -0.76 | -0.74 | 0.58 | 0.05 |
| 5 | 25750184 | 1.07E-06 | 1.17E-02 | -0.51 | -0.69 | -0.07 | 0.28 |
| 1 | 7726440 | 1.08E-06 | 1.17E-02 | -0.16 | 0.38 | -0.46 | 0.48 |
| 4 | 46851094 | 1.08E-06 | 1.17E-02 | 0.15 | -0.51 | -0.72 | 0.21 |
| 2 | 6720055 | 1.08E-06 | 1.17E-02 | 0.63 | 0.56 | -0.33 | -0.20 |
| 2 | 6720079 | 1.08E-06 | 1.17E-02 | 0.63 | 0.56 | -0.33 | -0.20 |
| 5 | 3944794 | 1.09E-06 | 1.17E-02 | -0.90 | -0.42 | 0.02 | -0.19 |
| 5 | 47705450 | 1.10E-06 | 1.17E-02 | 0.69 | -0.61 | -0.13 | -0.40 |
| 5 | 47705726 | 1.10E-06 | 1.17E-02 | 0.69 | -0.61 | -0.13 | -0.40 |
| 5 | 47705873 | 1.10E-06 | 1.17E-02 | 0.69 | -0.61 | -0.13 | -0.40 |
| 5 | 47706041 | 1.10E-06 | 1.17E-02 | 0.69 | -0.61 | -0.13 | -0.40 |
| 6 | 15399324 | 1.12E-06 | 1.17E-02 | 0.32 | -0.50 | -0.18 | 0.46 |
| 5 | 898764 | 1.12E-06 | 1.17E-02 | -0.04 | -0.89 | -0.39 | -1.26 |
| 5 | 900406 | 1.12E-06 | 1.17E-02 | -0.04 | -0.89 | -0.39 | -1.26 |
| 5 | 901258 | 1.12E-06 | 1.17E-02 | -0.04 | -0.89 | -0.39 | -1.26 |
| 5 | 902415 | 1.12E-06 | 1.17E-02 | -0.04 | -0.89 | -0.39 | -1.26 |
| 5 | 902570 | 1.12E-06 | 1.17E-02 | -0.04 | -0.89 | -0.39 | -1.26 |
| 6 | 42325884 | 1.13E-06 | 1.18E-02 | 0.41 | -0.69 | -0.46 | -0.68 |
| 12 | 16247701 | 1.13E-06 | 1.18E-02 | -0.70 | -0.32 | 0.36 | 0.25 |
| 5 | 3945418 | 1.13E-06 | 1.18E-02 | -0.91 | -0.39 | -0.03 | -0.19 |
| 1 | 37448789 | 1.16E-06 | 1.19E-02 | -0.05 | -0.57 | -0.32 | -0.39 |
| 9 | 17888197 | 1.16E-06 | 1.19E-02 | 0.08 | -0.25 | -0.33 | 0.48 |
| 2 | 21671039 | 1.16E-06 | 1.19E-02 | 0.07 | -0.53 | -0.62 | 0.25 |
| 2 | 21671148 | 1.16E-06 | 1.19E-02 | 0.07 | -0.53 | -0.62 | 0.25 |
| 8 | 21416760 | 1.17E-06 | 1.19E-02 | 0.75 | -0.37 | 0.24 | 0.25 |
| 2 | 18192583 | 1.18E-06 | 1.19E-02 | -0.73 | -0.07 | -0.75 | -0.82 |
| 12 | 13943802 | 1.18E-06 | 1.19E-02 | 1.43 | -0.29 | -0.25 | 0.18 |
| 5 | 38370435 | 1.18E-06 | 1.19E-02 | 0.75 | -0.42 | 0.00 | -0.42 |
| 5 | 38370438 | 1.18E-06 | 1.19E-02 | 0.75 | -0.42 | 0.00 | -0.42 |
| 9 | 30508861 | 1.18E-06 | 1.19E-02 | 0.82 | -1.04 | -0.10 | -0.07 |
| 9 | 37053106 | 1.18E-06 | 1.19E-02 | 0.09 | -0.40 | 0.12 | 0.73 |

| | | | | | | | |
|---|---|---|---|---|---|---|---|
| 6 | 10410368 | 1.18E-06 | 1.19E-02 | 0.67 | -0.16 | -0.65 | -1.03 |
| 6 | 10410405 | 1.18E-06 | 1.19E-02 | 0.67 | -0.16 | -0.65 | -1.03 |
| 6 | 16953114 | 1.20E-06 | 1.20E-02 | 0.16 | -0.13 | 0.72 | -0.12 |
| 10 | 15916647 | 1.20E-06 | 1.20E-02 | -0.02 | -0.32 | 0.27 | 0.56 |
| 2 | 6709580 | 1.21E-06 | 1.20E-02 | 0.67 | 0.54 | -0.30 | -0.16 |
| 6 | 6693142 | 1.22E-06 | 1.21E-02 | 0.49 | -0.53 | -0.49 | 0.07 |
| 6 | 31508559 | 1.24E-06 | 1.23E-02 | -0.21 | -0.49 | -0.54 | -0.30 |
| 8 | 20935269 | 1.25E-06 | 1.23E-02 | 0.47 | 0.02 | -0.54 | 0.22 |
| 6 | 15329392 | 1.25E-06 | 1.23E-02 | 0.55 | -0.54 | -0.35 | 0.27 |
| 6 | 9997138 | 1.26E-06 | 1.23E-02 | 0.55 | -0.50 | -0.41 | 0.01 |
| 5 | 25754604 | 1.27E-06 | 1.24E-02 | -0.59 | -0.61 | -0.05 | 0.23 |
| 10 | 6794758 | 1.27E-06 | 1.24E-02 | 0.18 | -0.60 | -0.55 | 0.28 |
| 6 | 9906618 | 1.28E-06 | 1.24E-02 | 0.39 | -0.71 | -0.22 | 0.18 |
| 6 | 9918091 | 1.28E-06 | 1.24E-02 | 0.39 | -0.71 | -0.22 | 0.18 |
| 5 | 25811788 | 1.30E-06 | 1.25E-02 | -0.76 | -0.48 | 0.06 | -0.09 |
| 5 | 25811906 | 1.30E-06 | 1.25E-02 | -0.76 | -0.48 | 0.06 | -0.09 |
| 11 | 25634771 | 1.30E-06 | 1.25E-02 | -0.55 | 0.12 | -0.69 | -0.48 |
| 6 | 9969865 | 1.31E-06 | 1.25E-02 | -0.24 | -0.42 | -0.75 | -0.96 |
| 6 | 28702912 | 1.31E-06 | 1.25E-02 | 1.60 | -0.74 | -0.91 | -0.45 |
| 9 | 27186705 | 1.31E-06 | 1.25E-02 | 0.95 | -0.24 | 0.03 | 0.36 |
| 6 | 39786947 | 1.34E-06 | 1.28E-02 | 1.74 | -0.49 | -0.94 | 0.25 |
| 10 | 20455283 | 1.35E-06 | 1.28E-02 | 0.74 | -0.58 | 0.06 | 0.27 |
| 1 | 37402195 | 1.35E-06 | 1.28E-02 | -0.10 | -0.58 | -0.33 | -0.35 |
| 2 | 14148852 | 1.35E-06 | 1.28E-02 | 0.53 | -0.29 | 0.25 | 1.14 |
| 1 | 7774757 | 1.36E-06 | 1.28E-02 | 0.09 | 0.57 | -0.28 | 0.47 |
| 4 | 50733233 | 1.36E-06 | 1.28E-02 | -1.03 | 1.95 | -0.53 | -1.08 |
| 8 | 51624037 | 1.38E-06 | 1.29E-02 | -0.11 | -0.12 | 0.08 | 0.73 |
| 2 | 21782416 | 1.39E-06 | 1.29E-02 | 0.43 | -0.52 | -0.35 | 0.39 |
| 10 | 2376484 | 1.39E-06 | 1.29E-02 | 0.42 | -0.61 | -0.48 | -0.06 |
| 5 | 38439839 | 1.40E-06 | 1.29E-02 | 0.89 | -0.42 | 0.09 | -0.34 |
| 2 | 18848735 | 1.40E-06 | 1.29E-02 | -0.71 | 0.14 | -0.52 | -0.43 |
| 2 | 18848780 | 1.40E-06 | 1.29E-02 | -0.71 | 0.14 | -0.52 | -0.43 |
| 2 | 18849126 | 1.40E-06 | 1.29E-02 | -0.71 | 0.14 | -0.52 | -0.43 |
| 2 | 18849172 | 1.40E-06 | 1.29E-02 | -0.71 | 0.14 | -0.52 | -0.43 |
| 2 | 21745534 | 1.41E-06 | 1.29E-02 | 0.30 | -0.70 | -0.39 | 0.23 |
| 9 | 27178282 | 1.41E-06 | 1.30E-02 | 1.28 | 0.05 | 0.20 | 0.15 |
| 6 | 9917740 | 1.43E-06 | 1.31E-02 | 0.47 | -0.66 | -0.19 | 0.18 |
| 11 | 22083165 | 1.44E-06 | 1.31E-02 | 0.28 | -0.49 | -0.41 | -0.47 |
| 11 | 22084380 | 1.44E-06 | 1.31E-02 | 0.28 | -0.49 | -0.41 | -0.47 |
| 11 | 22087893 | 1.44E-06 | 1.31E-02 | 0.28 | -0.49 | -0.41 | -0.47 |
| 4 | 53683159 | 1.46E-06 | 1.32E-02 | 0.93 | -0.36 | -0.25 | -0.35 |
| 9 | 30509024 | 1.46E-06 | 1.32E-02 | 0.88 | -1.04 | -0.06 | 0.00 |
| 9 | 20739186 | 1.47E-06 | 1.32E-02 | 1.07 | -0.67 | -0.01 | -0.72 |
| 2 | 26704511 | 1.47E-06 | 1.32E-02 | -0.17 | -0.28 | -0.73 | -0.22 |
| 8 | 21387385 | 1.47E-06 | 1.32E-02 | 0.83 | -0.19 | 0.12 | 0.33 |
| 5 | 38437768 | 1.49E-06 | 1.33E-02 | 0.99 | -0.23 | 0.10 | -0.53 |
| 1 | 37441405 | 1.49E-06 | 1.33E-02 | -0.16 | -0.57 | -0.21 | -0.38 |
| 6 | 44530603 | 1.49E-06 | 1.33E-02 | 0.68 | -0.64 | -0.55 | -0.21 |
| 6 | 16952519 | 1.51E-06 | 1.34E-02 | 0.07 | -0.15 | 0.72 | -0.11 |
| 6 | 28884409 | 1.52E-06 | 1.35E-02 | -1.10 | 0.61 | 0.03 | -0.83 |
| 9 | 38800617 | 1.52E-06 | 1.35E-02 | 1.49 | -0.63 | -0.06 | 0.73 |
| 6 | 32124893 | 1.53E-06 | 1.35E-02 | -0.19 | -0.52 | 0.55 | -0.23 |

| | | | | | | | |
|---|---|---|---|---|---|---|---|
| 7 | 40562114 | 1.54E-06 | 1.35E-02 | 0.31 | -0.21 | -0.43 | 0.52 |
| 8 | 44912075 | 1.54E-06 | 1.35E-02 | 0.11 | 0.48 | -0.66 | -0.01 |
| 12 | 457347 | 1.54E-06 | 1.35E-02 | 0.27 | 0.01 | -0.61 | -0.39 |
| 5 | 15424154 | 1.54E-06 | 1.35E-02 | 0.90 | -0.26 | -0.58 | 0.21 |
| 2 | 17343676 | 1.55E-06 | 1.35E-02 | -0.60 | -0.51 | 0.23 | -0.35 |
| 6 | 15307223 | 1.55E-06 | 1.35E-02 | -0.42 | -1.34 | -0.90 | 0.35 |
| 6 | 10414325 | 1.56E-06 | 1.35E-02 | 0.26 | -0.22 | -0.72 | -1.03 |
| 1 | 13310558 | 1.56E-06 | 1.35E-02 | 0.59 | -0.53 | 0.01 | 0.34 |
| 10 | 24307676 | 1.57E-06 | 1.36E-02 | -0.81 | -0.39 | 0.30 | 0.19 |
| 11 | 54742321 | 1.57E-06 | 1.36E-02 | -1.23 | 0.57 | 0.03 | 0.23 |
| 8 | 27349256 | 1.58E-06 | 1.36E-02 | 0.35 | 0.36 | -0.40 | -0.54 |
| 6 | 15086198 | 1.58E-06 | 1.36E-02 | -0.46 | -1.09 | -0.76 | 0.26 |
| 2 | 21685913 | 1.60E-06 | 1.37E-02 | 0.78 | -0.55 | -0.27 | -0.26 |
| 9 | 49402699 | 1.61E-06 | 1.37E-02 | 1.56 | 0.99 | -1.06 | -0.51 |
| 2 | 10147447 | 1.63E-06 | 1.38E-02 | -1.07 | 0.32 | 0.16 | 0.68 |
| 9 | 17258472 | 1.63E-06 | 1.38E-02 | -0.03 | -0.73 | 0.20 | 0.16 |
| 12 | 10686573 | 1.65E-06 | 1.38E-02 | -0.72 | 0.24 | 0.31 | 0.51 |
| 6 | 5478130 | 1.65E-06 | 1.38E-02 | 0.24 | -0.07 | -0.93 | -0.78 |
| 6 | 5479474 | 1.65E-06 | 1.38E-02 | 0.24 | -0.07 | -0.93 | -0.78 |
| 6 | 5479799 | 1.65E-06 | 1.38E-02 | 0.24 | -0.07 | -0.93 | -0.78 |
| 6 | 5481772 | 1.65E-06 | 1.38E-02 | 0.24 | -0.07 | -0.93 | -0.78 |
| 8 | 21414151 | 1.66E-06 | 1.39E-02 | 0.72 | -0.40 | 0.22 | 0.20 |
| 9 | 9879767 | 1.66E-06 | 1.39E-02 | 2.38 | 0.58 | -1.47 | -0.16 |
| 2 | 27105444 | 1.67E-06 | 1.39E-02 | -1.08 | -0.01 | 0.37 | 0.20 |
| 8 | 35818559 | 1.67E-06 | 1.39E-02 | -0.36 | -0.53 | -0.31 | 0.42 |
| 9 | 20739303 | 1.68E-06 | 1.39E-02 | 0.98 | -0.23 | 0.01 | -0.82 |
| 6 | 5481442 | 1.68E-06 | 1.39E-02 | 0.35 | -0.10 | -0.93 | -0.82 |
| 1 | 5105456 | 1.69E-06 | 1.39E-02 | 0.36 | -0.11 | 0.00 | 0.58 |
| 8 | 21387570 | 1.69E-06 | 1.39E-02 | 0.86 | -0.23 | 0.11 | 0.29 |
| 6 | 15372156 | 1.69E-06 | 1.39E-02 | 0.57 | -0.48 | -0.35 | 0.30 |
| 6 | 15373234 | 1.69E-06 | 1.39E-02 | 0.57 | -0.48 | -0.35 | 0.30 |
| 8 | 21411345 | 1.72E-06 | 1.41E-02 | 0.99 | -0.22 | 0.10 | 0.18 |
| 8 | 21411346 | 1.72E-06 | 1.41E-02 | 0.99 | -0.22 | 0.10 | 0.18 |
| 7 | 26100594 | 1.73E-06 | 1.41E-02 | 0.28 | -0.60 | -0.40 | 0.29 |
| 8 | 22436564 | 1.75E-06 | 1.42E-02 | -0.37 | 0.43 | -0.52 | 0.54 |
| 4 | 27795427 | 1.75E-06 | 1.42E-02 | -0.23 | 0.32 | -0.62 | 1.36 |
| 6 | 12585776 | 1.76E-06 | 1.43E-02 | -0.29 | -0.65 | -0.69 | -0.26 |
| 12 | 16247948 | 1.80E-06 | 1.45E-02 | -0.67 | -0.31 | 0.33 | 0.27 |
| 4 | 52713330 | 1.80E-06 | 1.45E-02 | -0.09 | -0.08 | -0.80 | -0.31 |
| 4 | 52878140 | 1.80E-06 | 1.45E-02 | 0.74 | -0.38 | -0.40 | -0.20 |
| 1 | 1411194 | 1.83E-06 | 1.47E-02 | 0.11 | -0.32 | -0.38 | -0.48 |
| 5 | 38449745 | 1.83E-06 | 1.47E-02 | 0.92 | -0.43 | 0.12 | -0.18 |
| 4 | 65108415 | 1.85E-06 | 1.47E-02 | 0.73 | -0.27 | -0.29 | -0.58 |
| 6 | 36373131 | 1.88E-06 | 1.50E-02 | 1.38 | -0.52 | 0.08 | -0.30 |
| 2 | 15031901 | 1.89E-06 | 1.50E-02 | 0.77 | -0.58 | -0.33 | 0.29 |
| 6 | 15091614 | 1.89E-06 | 1.50E-02 | 1.11 | -1.51 | -0.21 | -0.54 |
| 10 | 24307423 | 1.90E-06 | 1.50E-02 | -0.78 | -0.40 | 0.25 | 0.25 |
| 9 | 45523850 | 1.91E-06 | 1.50E-02 | 0.09 | -0.74 | -0.65 | -0.05 |
| 9 | 45523873 | 1.91E-06 | 1.50E-02 | 0.09 | -0.74 | -0.65 | -0.05 |
| 9 | 42328868 | 1.93E-06 | 1.50E-02 | 0.17 | 0.25 | -1.35 | -0.06 |
| 8 | 49048590 | 1.93E-06 | 1.50E-02 | 0.38 | -0.29 | -0.67 | -0.11 |
| 8 | 49049081 | 1.93E-06 | 1.50E-02 | 0.38 | -0.29 | -0.67 | -0.11 |

| | | | | | | | |
|---|---|---|---|---|---|---|---|
| 8 | 49049715 | 1.93E-06 | 1.50E-02 | 0.38 | -0.29 | -0.67 | -0.11 |
| 8 | 49050751 | 1.93E-06 | 1.50E-02 | 0.38 | -0.29 | -0.67 | -0.11 |
| 8 | 49050865 | 1.93E-06 | 1.50E-02 | 0.38 | -0.29 | -0.67 | -0.11 |
| 8 | 49057614 | 1.93E-06 | 1.50E-02 | 0.38 | -0.29 | -0.67 | -0.11 |
| 8 | 49057929 | 1.93E-06 | 1.50E-02 | 0.38 | -0.29 | -0.67 | -0.11 |
| 8 | 36608782 | 1.94E-06 | 1.50E-02 | 0.85 | -0.20 | -0.27 | 0.17 |
| 10 | 15774297 | 1.95E-06 | 1.50E-02 | 1.03 | -0.01 | -0.43 | 0.15 |
| 10 | 16112183 | 1.95E-06 | 1.50E-02 | 0.86 | -0.18 | 0.46 | 0.28 |
| 9 | 17887105 | 1.96E-06 | 1.50E-02 | 0.08 | -0.53 | -0.38 | 0.40 |
| 11 | 12862573 | 1.96E-06 | 1.50E-02 | 0.88 | -0.04 | -0.61 | 0.11 |
| 6 | 44825341 | 1.97E-06 | 1.51E-02 | 0.72 | -0.62 | -0.71 | 0.01 |
| 9 | 30509221 | 1.98E-06 | 1.51E-02 | 0.84 | -1.07 | -0.09 | 0.01 |
| 2 | 26608743 | 1.98E-06 | 1.51E-02 | 0.92 | -0.10 | -0.41 | -0.23 |
| 5 | 38439726 | 2.01E-06 | 1.52E-02 | 0.91 | -0.40 | 0.21 | -0.37 |
| 6 | 6361249 | 2.01E-06 | 1.52E-02 | 0.69 | -0.58 | -0.29 | -0.05 |
| 6 | 45040591 | 2.01E-06 | 1.52E-02 | 0.17 | -0.42 | -0.83 | -0.38 |
| 6 | 31500901 | 2.02E-06 | 1.52E-02 | 0.35 | -0.67 | -0.42 | -0.13 |
| 12 | 13943748 | 2.03E-06 | 1.53E-02 | 1.48 | -0.25 | -0.24 | 0.15 |
| 1 | 40312715 | 2.04E-06 | 1.53E-02 | 0.10 | 0.00 | 0.29 | 0.78 |
| 1 | 26606984 | 2.05E-06 | 1.54E-02 | 0.56 | -0.50 | -0.39 | -0.26 |
| 2 | 18188913 | 2.05E-06 | 1.54E-02 | 0.45 | -0.55 | -0.06 | 0.82 |
| 6 | 44516685 | 2.05E-06 | 1.54E-02 | 0.64 | -0.76 | -0.51 | -0.18 |
| 6 | 44061420 | 2.06E-06 | 1.54E-02 | -0.20 | -0.34 | 0.09 | 0.49 |
| 2 | 10146202 | 2.07E-06 | 1.54E-02 | -1.15 | 0.13 | 0.19 | 0.53 |
| 6 | 28466274 | 2.08E-06 | 1.55E-02 | 0.38 | 0.14 | 0.10 | 0.69 |
| 5 | 66744270 | 2.11E-06 | 1.56E-02 | -0.33 | -1.01 | -0.67 | 0.26 |
| 7 | 30945595 | 2.12E-06 | 1.57E-02 | -0.51 | -0.48 | 0.19 | 0.35 |
| 6 | 36355197 | 2.13E-06 | 1.57E-02 | 1.15 | -0.60 | -0.30 | -0.27 |
| 2 | 6720141 | 2.16E-06 | 1.59E-02 | 0.54 | 0.54 | -0.35 | -0.22 |
| 8 | 26431829 | 2.18E-06 | 1.61E-02 | 0.95 | -0.22 | 0.00 | 0.17 |
| 7 | 26096720 | 2.21E-06 | 1.62E-02 | 0.30 | -0.53 | -0.26 | 0.34 |
| 9 | 20761300 | 2.22E-06 | 1.63E-02 | -0.93 | 0.50 | 0.66 | -0.14 |
| 9 | 29162530 | 2.23E-06 | 1.63E-02 | 0.83 | -0.42 | -0.21 | 0.03 |
| 1 | 13017528 | 2.24E-06 | 1.63E-02 | 0.54 | -0.56 | -0.56 | -0.22 |
| 6 | 15324065 | 2.24E-06 | 1.63E-02 | 0.59 | -0.51 | -0.31 | 0.27 |
| 6 | 29803384 | 2.25E-06 | 1.63E-02 | 0.26 | -0.42 | 0.16 | 0.60 |
| 1 | 15380928 | 2.25E-06 | 1.63E-02 | 0.82 | -0.34 | -0.29 | -0.12 |
| 6 | 17723539 | 2.31E-06 | 1.67E-02 | 0.54 | -0.41 | -0.45 | -0.24 |
| 6 | 9906612 | 2.33E-06 | 1.68E-02 | 0.50 | -0.65 | -0.22 | 0.11 |
| 10 | 20433773 | 2.33E-06 | 1.68E-02 | 0.82 | -0.70 | -0.12 | 0.16 |
| 5 | 47704437 | 2.36E-06 | 1.70E-02 | 0.60 | -0.59 | -0.11 | -0.43 |
| 6 | 44826442 | 2.37E-06 | 1.70E-02 | 0.59 | -1.04 | -0.56 | -0.19 |
| 11 | 51866782 | 2.37E-06 | 1.70E-02 | -0.11 | 0.27 | -0.38 | 0.54 |
| 2 | 6719445 | 2.38E-06 | 1.70E-02 | 0.53 | 0.58 | -0.36 | -0.14 |
| 1 | 47626889 | 2.40E-06 | 1.71E-02 | -2.27 | -1.18 | -1.03 | -0.77 |
| 5 | 25001334 | 2.40E-06 | 1.71E-02 | -0.22 | -0.34 | -0.17 | 0.71 |
| 12 | 10689812 | 2.41E-06 | 1.71E-02 | -0.66 | 0.05 | 0.12 | 0.49 |
| 2 | 18189049 | 2.43E-06 | 1.71E-02 | 0.47 | -0.58 | -0.04 | 0.79 |
| 10 | 10597856 | 2.43E-06 | 1.71E-02 | 0.26 | -0.69 | -0.22 | -0.41 |
| 10 | 10606035 | 2.43E-06 | 1.71E-02 | 0.26 | -0.69 | -0.22 | -0.41 |
| 2 | 6715378 | 2.43E-06 | 1.71E-02 | 0.61 | 0.55 | -0.33 | -0.16 |
| 2 | 6716026 | 2.43E-06 | 1.71E-02 | 0.61 | 0.55 | -0.33 | -0.16 |

| | | | | | | | |
|---|---|---|---|---|---|---|---|
| 1 | 13330653 | 2.43E-06 | 1.71E-02 | -0.08 | -0.36 | 0.10 | 0.53 |
| 4 | 19405339 | 2.43E-06 | 1.71E-02 | 0.99 | -0.15 | -0.02 | -0.47 |
| 6 | 44507633 | 2.45E-06 | 1.71E-02 | 0.66 | -0.51 | -0.30 | -0.51 |
| 1 | 37399254 | 2.46E-06 | 1.72E-02 | -0.08 | -0.61 | -0.34 | -0.31 |
| 1 | 13316515 | 2.49E-06 | 1.74E-02 | 0.56 | -0.51 | 0.09 | 0.36 |
| 6 | 15327993 | 2.50E-06 | 1.74E-02 | 0.60 | -0.48 | -0.32 | 0.27 |
| 5 | 65806532 | 2.51E-06 | 1.74E-02 | 0.22 | -0.63 | -0.52 | -0.28 |
| 5 | 65808280 | 2.51E-06 | 1.74E-02 | 0.22 | -0.63 | -0.52 | -0.28 |
| 5 | 65810458 | 2.51E-06 | 1.74E-02 | 0.22 | -0.63 | -0.52 | -0.28 |
| 6 | 44528298 | 2.51E-06 | 1.74E-02 | 0.63 | -0.60 | -0.50 | -0.23 |
| 11 | 30920336 | 2.52E-06 | 1.74E-02 | -0.20 | -0.68 | 0.22 | 0.04 |
| 5 | 25753227 | 2.52E-06 | 1.74E-02 | -0.63 | -0.53 | 0.09 | 0.16 |
| 11 | 32462236 | 2.55E-06 | 1.74E-02 | -0.88 | -0.15 | 0.28 | 0.25 |
| 6 | 53079043 | 2.57E-06 | 1.74E-02 | 1.12 | -0.23 | -0.34 | 0.07 |
| 2 | 6710292 | 2.58E-06 | 1.74E-02 | 0.66 | 0.52 | -0.29 | -0.17 |
| 10 | 8069875 | 2.59E-06 | 1.74E-02 | 0.30 | -0.32 | 0.19 | 0.60 |
| 2 | 6718886 | 2.60E-06 | 1.74E-02 | 0.60 | 0.56 | -0.31 | -0.18 |
| 7 | 31022136 | 2.60E-06 | 1.74E-02 | -0.55 | -0.41 | -0.13 | 0.67 |
| 7 | 31022231 | 2.60E-06 | 1.74E-02 | -0.55 | -0.41 | -0.13 | 0.67 |
| 6 | 44528270 | 2.61E-06 | 1.74E-02 | 0.73 | -0.64 | -0.51 | -0.17 |
| 11 | 22081025 | 2.61E-06 | 1.74E-02 | 0.27 | -0.52 | -0.36 | -0.47 |
| 6 | 15322209 | 2.64E-06 | 1.74E-02 | 0.55 | -0.55 | -0.33 | 0.25 |
| 7 | 26087974 | 2.64E-06 | 1.74E-02 | 0.36 | -0.64 | -0.37 | 0.28 |
| 6 | 44017711 | 2.64E-06 | 1.74E-02 | -0.63 | -0.48 | 0.18 | 0.24 |
| 6 | 44019346 | 2.64E-06 | 1.74E-02 | -0.63 | -0.48 | 0.18 | 0.24 |
| 6 | 44019757 | 2.64E-06 | 1.74E-02 | -0.63 | -0.48 | 0.18 | 0.24 |
| 6 | 44020433 | 2.64E-06 | 1.74E-02 | -0.63 | -0.48 | 0.18 | 0.24 |
| 6 | 44021758 | 2.64E-06 | 1.74E-02 | -0.63 | -0.48 | 0.18 | 0.24 |
| 2 | 26671842 | 2.64E-06 | 1.74E-02 | 0.80 | -0.05 | -0.43 | -0.39 |
| 11 | 46753797 | 2.65E-06 | 1.74E-02 | 1.33 | -0.31 | -0.77 | -0.31 |
| 2 | 10147364 | 2.65E-06 | 1.74E-02 | -1.12 | 0.14 | 0.19 | 0.54 |
| 6 | 45049339 | 2.66E-06 | 1.74E-02 | 0.31 | -0.53 | -0.79 | -0.39 |
| 6 | 45050874 | 2.66E-06 | 1.74E-02 | 0.31 | -0.53 | -0.79 | -0.39 |
| 12 | 13530841 | 2.68E-06 | 1.74E-02 | -1.08 | 0.01 | 0.33 | 0.36 |
| 6 | 45568774 | 2.68E-06 | 1.74E-02 | -0.55 | -0.10 | 0.47 | -0.37 |
| 2 | 26608138 | 2.70E-06 | 1.74E-02 | 0.83 | -0.11 | -0.48 | -0.21 |
| 6 | 15355308 | 2.70E-06 | 1.74E-02 | 0.50 | -0.56 | -0.38 | 0.24 |
| 10 | 40578055 | 2.72E-06 | 1.74E-02 | 0.53 | -0.66 | -0.30 | 0.03 |
| 6 | 15211315 | 2.72E-06 | 1.74E-02 | -0.18 | -0.92 | -0.65 | 0.27 |
| 8 | 36608892 | 2.73E-06 | 1.74E-02 | 0.82 | -0.20 | -0.28 | 0.17 |
| 5 | 3944171 | 2.74E-06 | 1.74E-02 | -0.87 | -0.44 | -0.03 | -0.13 |
| 5 | 25751219 | 2.75E-06 | 1.74E-02 | -0.53 | -0.60 | -0.05 | 0.31 |
| 10 | 24306490 | 2.75E-06 | 1.74E-02 | -0.84 | -0.28 | 0.36 | 0.23 |
| 10 | 24306524 | 2.75E-06 | 1.74E-02 | -0.84 | -0.28 | 0.36 | 0.23 |
| 10 | 24306557 | 2.75E-06 | 1.74E-02 | -0.84 | -0.28 | 0.36 | 0.23 |
| 6 | 10399177 | 2.75E-06 | 1.74E-02 | 0.56 | -0.15 | -0.70 | -0.97 |
| 6 | 10400036 | 2.75E-06 | 1.74E-02 | 0.56 | -0.15 | -0.70 | -0.97 |
| 6 | 10400159 | 2.75E-06 | 1.74E-02 | 0.56 | -0.15 | -0.70 | -0.97 |
| 6 | 10401672 | 2.75E-06 | 1.74E-02 | 0.56 | -0.15 | -0.70 | -0.97 |
| 6 | 10402511 | 2.75E-06 | 1.74E-02 | 0.56 | -0.15 | -0.70 | -0.97 |
| 6 | 10408258 | 2.75E-06 | 1.74E-02 | 0.56 | -0.15 | -0.70 | -0.97 |
| 6 | 10408830 | 2.75E-06 | 1.74E-02 | 0.56 | -0.15 | -0.70 | -0.97 |

| | | | | | | | |
|---|---|---|---|---|---|---|---|
| 6 | 10408963 | 2.75E-06 | 1.74E-02 | 0.56 | -0.15 | -0.70 | -0.97 |
| 6 | 10409223 | 2.75E-06 | 1.74E-02 | 0.56 | -0.15 | -0.70 | -0.97 |
| 6 | 10410043 | 2.75E-06 | 1.74E-02 | 0.56 | -0.15 | -0.70 | -0.97 |
| 6 | 10410079 | 2.75E-06 | 1.74E-02 | 0.56 | -0.15 | -0.70 | -0.97 |
| 6 | 10410421 | 2.75E-06 | 1.74E-02 | 0.56 | -0.15 | -0.70 | -0.97 |
| 6 | 10412202 | 2.75E-06 | 1.74E-02 | 0.56 | -0.15 | -0.70 | -0.97 |
| 6 | 10412230 | 2.75E-06 | 1.74E-02 | 0.56 | -0.15 | -0.70 | -0.97 |
| 6 | 10414004 | 2.75E-06 | 1.74E-02 | 0.56 | -0.15 | -0.70 | -0.97 |
| 1 | 14561344 | 2.77E-06 | 1.74E-02 | 0.72 | 0.70 | -0.20 | 0.05 |
| 6 | 15412607 | 2.78E-06 | 1.74E-02 | 0.40 | -0.46 | -0.23 | 0.39 |
| 6 | 15412614 | 2.78E-06 | 1.74E-02 | 0.40 | -0.46 | -0.23 | 0.39 |
| 6 | 15412654 | 2.78E-06 | 1.74E-02 | 0.40 | -0.46 | -0.23 | 0.39 |
| 5 | 60658137 | 2.79E-06 | 1.74E-02 | -0.50 | -0.56 | 0.24 | 0.24 |
| 1 | 22016275 | 2.80E-06 | 1.75E-02 | 0.65 | -0.44 | -0.43 | -0.48 |
| 6 | 19629038 | 2.80E-06 | 1.75E-02 | 1.81 | -0.58 | -0.82 | -0.70 |
| 6 | 9997211 | 2.82E-06 | 1.75E-02 | 0.85 | -0.61 | -0.26 | -0.46 |
| 10 | 32982270 | 2.83E-06 | 1.76E-02 | 1.74 | -0.66 | -0.12 | 0.02 |
| 1 | 40307232 | 2.84E-06 | 1.76E-02 | -0.01 | -0.19 | -0.02 | 0.68 |
| 6 | 6365481 | 2.85E-06 | 1.76E-02 | 0.72 | -0.60 | -0.50 | -0.48 |
| 2 | 18848099 | 2.85E-06 | 1.76E-02 | -0.76 | 0.08 | -0.53 | -0.39 |
| 5 | 71560470 | 2.87E-06 | 1.77E-02 | -0.39 | 0.55 | -0.18 | 0.58 |
| 6 | 12591366 | 2.89E-06 | 1.78E-02 | -0.09 | -0.41 | -0.59 | -0.48 |
| 6 | 47223895 | 2.89E-06 | 1.78E-02 | 0.01 | 1.04 | 0.37 | -0.02 |
| 4 | 33724703 | 2.90E-06 | 1.78E-02 | 0.42 | -0.07 | 0.24 | 0.58 |
| 2 | 18181804 | 2.90E-06 | 1.78E-02 | -0.78 | 0.08 | -0.72 | -0.89 |
| 5 | 27726344 | 2.93E-06 | 1.79E-02 | -0.51 | 0.40 | 0.55 | -0.05 |
| 2 | 4505994 | 2.94E-06 | 1.79E-02 | -0.98 | -0.24 | 0.88 | -0.41 |
| 6 | 45567186 | 2.94E-06 | 1.79E-02 | -0.46 | -0.13 | 0.49 | -0.38 |
| 9 | 21442785 | 2.95E-06 | 1.80E-02 | -0.69 | 0.73 | -0.72 | 0.25 |
| 10 | 40109644 | 2.98E-06 | 1.81E-02 | 0.40 | -0.68 | -0.05 | -0.44 |
| 1 | 25330139 | 2.99E-06 | 1.81E-02 | 1.64 | 0.08 | 0.17 | -0.05 |
| 5 | 25777725 | 3.03E-06 | 1.84E-02 | -0.70 | -0.48 | 0.07 | -0.08 |
| 6 | 15410757 | 3.04E-06 | 1.84E-02 | 0.46 | -0.40 | -0.28 | 0.40 |
| 6 | 15399323 | 3.04E-06 | 1.84E-02 | -0.62 | -0.66 | -0.19 | 0.08 |
| 11 | 51866174 | 3.05E-06 | 1.84E-02 | 0.02 | 0.37 | -0.35 | 0.48 |
| 6 | 32517549 | 3.06E-06 | 1.84E-02 | 0.97 | -0.26 | -0.77 | -0.74 |
| 9 | 35421054 | 3.07E-06 | 1.84E-02 | -0.58 | -0.38 | 0.52 | 0.14 |
| 9 | 35816758 | 3.09E-06 | 1.84E-02 | -0.37 | -0.88 | 0.16 | 0.36 |
| 8 | 20945701 | 3.10E-06 | 1.84E-02 | 0.41 | -0.19 | -0.42 | 0.32 |
| 2 | 29400467 | 3.11E-06 | 1.84E-02 | -0.12 | -0.22 | 0.57 | 0.46 |
| 6 | 44377864 | 3.11E-06 | 1.84E-02 | -0.16 | 0.92 | -0.80 | -0.28 |
| 1 | 40312055 | 3.11E-06 | 1.84E-02 | 0.25 | -0.21 | 0.21 | 0.65 |
| 5 | 38441234 | 3.11E-06 | 1.84E-02 | 0.86 | -0.34 | 0.16 | -0.45 |
| 5 | 38441337 | 3.11E-06 | 1.84E-02 | 0.86 | -0.34 | 0.16 | -0.45 |
| 5 | 38441369 | 3.11E-06 | 1.84E-02 | 0.86 | -0.34 | 0.16 | -0.45 |
| 6 | 35543767 | 3.12E-06 | 1.85E-02 | -1.05 | 0.86 | 0.66 | 0.71 |
| 6 | 35532854 | 3.14E-06 | 1.85E-02 | -1.05 | 1.10 | 0.81 | 0.76 |
| 6 | 15360657 | 3.17E-06 | 1.86E-02 | 0.50 | -0.56 | -0.35 | 0.27 |
| 9 | 28000748 | 3.17E-06 | 1.86E-02 | 1.48 | -0.57 | -0.51 | 0.08 |
| 5 | 25872318 | 3.17E-06 | 1.86E-02 | 1.06 | 0.31 | -0.06 | 0.04 |
| 8 | 44087772 | 3.18E-06 | 1.86E-02 | 0.73 | -0.20 | 0.37 | -0.30 |
| 3 | 272822 | 3.19E-06 | 1.87E-02 | -0.49 | -0.18 | 0.59 | -0.40 |

| | | | | | | | |
|---|---|---|---|---|---|---|---|
| 2 | 6693372 | 3.24E-06 | 1.89E-02 | -0.72 | -0.72 | 0.57 | 0.08 |
| 8 | 9304529 | 3.24E-06 | 1.89E-02 | 0.29 | -0.80 | 0.36 | 1.22 |
| 8 | 21357817 | 3.24E-06 | 1.89E-02 | 1.42 | -0.60 | 0.25 | 0.48 |
| 5 | 25749730 | 3.26E-06 | 1.89E-02 | -0.52 | -0.67 | -0.10 | 0.27 |
| 6 | 15299004 | 3.28E-06 | 1.90E-02 | 0.39 | -0.63 | -0.56 | 0.17 |
| 4 | 54034100 | 3.28E-06 | 1.90E-02 | 0.97 | -0.42 | 0.32 | -0.34 |
| 2 | 16367805 | 3.29E-06 | 1.90E-02 | -0.31 | 0.24 | -0.08 | 0.87 |
| 6 | 15328586 | 3.29E-06 | 1.90E-02 | 0.51 | -0.54 | -0.32 | 0.29 |
| 12 | 16254760 | 3.29E-06 | 1.90E-02 | -0.66 | -0.35 | 0.33 | 0.23 |
| 5 | 80075375 | 3.33E-06 | 1.92E-02 | -1.18 | 0.54 | 1.28 | 0.63 |
| 7 | 42264267 | 3.34E-06 | 1.93E-02 | 1.16 | -0.24 | 0.30 | -0.32 |
| 1 | 43825473 | 3.36E-06 | 1.93E-02 | 1.76 | -0.55 | -1.11 | -0.75 |
| 4 | 30260939 | 3.38E-06 | 1.94E-02 | 0.87 | 0.20 | -0.62 | -0.35 |
| 8 | 27308105 | 3.41E-06 | 1.95E-02 | 0.07 | 0.25 | -0.62 | -0.37 |
| 6 | 44512976 | 3.42E-06 | 1.96E-02 | 0.62 | -0.70 | -0.44 | -0.20 |
| 4 | 65108257 | 3.42E-06 | 1.96E-02 | 0.78 | -0.38 | -0.28 | -0.50 |
| 1 | 22016372 | 3.43E-06 | 1.96E-02 | 0.59 | -0.44 | -0.43 | -0.48 |
| 9 | 39336948 | 3.44E-06 | 1.96E-02 | 1.04 | -0.48 | -0.33 | -0.09 |
| 10 | 5943132 | 3.46E-06 | 1.97E-02 | 0.69 | -0.37 | -0.34 | -0.11 |
| 7 | 30800134 | 3.48E-06 | 1.97E-02 | 0.56 | -0.16 | -0.57 | 0.41 |
| 8 | 57626511 | 3.48E-06 | 1.97E-02 | -0.10 | -0.02 | -0.68 | -0.12 |
| 6 | 44060436 | 3.50E-06 | 1.98E-02 | -0.21 | -0.28 | 0.14 | 0.49 |
| 2 | 10147501 | 3.50E-06 | 1.98E-02 | -1.08 | 0.30 | 0.19 | 0.70 |
| 9 | 39340201 | 3.53E-06 | 1.99E-02 | 1.08 | -0.48 | -0.42 | -0.18 |
| 8 | 21385964 | 3.58E-06 | 2.01E-02 | 0.86 | -0.14 | -0.04 | 0.30 |
| 2 | 10864371 | 3.59E-06 | 2.01E-02 | -0.52 | -0.48 | 0.11 | -0.24 |
| 10 | 10580632 | 3.59E-06 | 2.01E-02 | 0.35 | -0.69 | -0.21 | -0.43 |
| 12 | 16250221 | 3.68E-06 | 2.06E-02 | -0.60 | -0.33 | 0.29 | 0.29 |
| 5 | 25773280 | 3.69E-06 | 2.06E-02 | -0.76 | -0.48 | 0.09 | -0.06 |
| 5 | 25773305 | 3.69E-06 | 2.06E-02 | -0.76 | -0.48 | 0.09 | -0.06 |
| 5 | 25754656 | 3.70E-06 | 2.06E-02 | -0.51 | -0.64 | -0.05 | 0.26 |
| 8 | 44077645 | 3.71E-06 | 2.06E-02 | 0.72 | -0.15 | 0.41 | -0.28 |
| 8 | 44077713 | 3.71E-06 | 2.06E-02 | 0.72 | -0.15 | 0.41 | -0.28 |
| 5 | 38458735 | 3.72E-06 | 2.06E-02 | 0.83 | -0.44 | 0.03 | -0.32 |
| 5 | 38458828 | 3.72E-06 | 2.06E-02 | 0.83 | -0.44 | 0.03 | -0.32 |
| 10 | 2373605 | 3.73E-06 | 2.06E-02 | 0.26 | -0.73 | -0.50 | 0.02 |
| 6 | 15260349 | 3.73E-06 | 2.06E-02 | -0.32 | -1.06 | -0.75 | 0.30 |
| 6 | 15261187 | 3.73E-06 | 2.06E-02 | -0.32 | -1.06 | -0.75 | 0.30 |
| 10 | 15918138 | 3.74E-06 | 2.06E-02 | -0.01 | -0.42 | 0.18 | 0.51 |
| 2 | 18180456 | 3.75E-06 | 2.06E-02 | -0.60 | -0.21 | -0.64 | -0.80 |
| 6 | 35363243 | 3.77E-06 | 2.07E-02 | -0.14 | -0.58 | -0.56 | -0.19 |
| 10 | 10577765 | 3.78E-06 | 2.08E-02 | 0.36 | -0.69 | -0.20 | -0.43 |
| 2 | 261568 | 3.81E-06 | 2.08E-02 | 1.15 | -0.06 | 0.28 | 0.22 |
| 9 | 38808166 | 3.87E-06 | 2.12E-02 | 1.17 | -0.51 | -0.23 | 0.23 |
| 12 | 16254867 | 3.91E-06 | 2.13E-02 | -0.68 | -0.37 | 0.29 | 0.22 |
| 5 | 13845001 | 3.92E-06 | 2.13E-02 | 1.18 | 0.18 | -0.99 | -0.41 |
| 8 | 44912335 | 3.93E-06 | 2.13E-02 | 0.17 | 0.47 | -0.65 | -0.02 |
| 2 | 28983502 | 3.94E-06 | 2.13E-02 | -0.67 | -0.53 | -0.10 | 1.13 |
| 2 | 28983561 | 3.94E-06 | 2.13E-02 | -0.67 | -0.53 | -0.10 | 1.13 |
| 2 | 28983716 | 3.94E-06 | 2.13E-02 | -0.67 | -0.53 | -0.10 | 1.13 |
| 2 | 28984722 | 3.94E-06 | 2.13E-02 | -0.67 | -0.53 | -0.10 | 1.13 |
| 12 | 10825650 | 3.95E-06 | 2.13E-02 | -0.35 | 0.49 | 0.21 | 0.59 |

| | | | | | | | |
|---|---|---|---|---|---|---|---|
| 6 | 44527239 | 3.99E-06 | 2.15E-02 | 0.60 | -0.69 | -0.52 | -0.14 |
| 11 | 31783479 | 4.00E-06 | 2.15E-02 | 0.22 | -0.36 | -0.12 | 0.50 |
| 6 | 6369155 | 4.01E-06 | 2.15E-02 | 0.69 | -0.53 | -0.04 | 0.12 |
| 6 | 6369176 | 4.01E-06 | 2.15E-02 | 0.69 | -0.53 | -0.04 | 0.12 |
| 6 | 6369215 | 4.01E-06 | 2.15E-02 | 0.69 | -0.53 | -0.04 | 0.12 |
| 6 | 6365165 | 4.03E-06 | 2.15E-02 | 0.65 | -0.60 | -0.31 | -0.06 |
| 6 | 9997273 | 4.04E-06 | 2.15E-02 | 0.70 | -0.48 | -0.32 | -0.66 |
| 6 | 9997284 | 4.04E-06 | 2.15E-02 | 0.70 | -0.48 | -0.32 | -0.66 |
| 10 | 10576016 | 4.04E-06 | 2.15E-02 | 0.27 | -0.71 | -0.23 | -0.42 |
| 6 | 15353096 | 4.05E-06 | 2.15E-02 | 0.50 | -0.54 | -0.36 | 0.27 |
| 6 | 15353119 | 4.05E-06 | 2.15E-02 | 0.50 | -0.54 | -0.36 | 0.27 |
| 9 | 21442607 | 4.06E-06 | 2.15E-02 | -0.71 | 0.68 | -0.91 | 0.11 |
| 8 | 37692490 | 4.06E-06 | 2.15E-02 | 0.21 | -0.48 | -0.12 | 0.46 |
| 1 | 37429100 | 4.07E-06 | 2.15E-02 | -0.12 | -0.58 | -0.34 | -0.32 |
| 3 | 8083924 | 4.08E-06 | 2.16E-02 | 1.25 | -0.57 | -0.52 | -0.22 |
| 6 | 27976932 | 4.09E-06 | 2.16E-02 | -0.08 | -0.19 | -0.11 | -0.62 |
| 1 | 34679063 | 4.12E-06 | 2.16E-02 | 0.61 | 0.45 | -0.54 | -0.71 |
| 6 | 9919569 | 4.15E-06 | 2.16E-02 | 0.53 | -0.62 | -0.13 | 0.14 |
| 7 | 26108361 | 4.15E-06 | 2.16E-02 | 0.33 | -0.57 | -0.30 | 0.31 |
| 7 | 26108417 | 4.15E-06 | 2.16E-02 | 0.33 | -0.57 | -0.30 | 0.31 |
| 7 | 26109165 | 4.15E-06 | 2.16E-02 | 0.33 | -0.57 | -0.30 | 0.31 |
| 7 | 26109250 | 4.15E-06 | 2.16E-02 | 0.33 | -0.57 | -0.30 | 0.31 |
| 7 | 26109285 | 4.15E-06 | 2.16E-02 | 0.33 | -0.57 | -0.30 | 0.31 |
| 2 | 18186052 | 4.15E-06 | 2.16E-02 | 0.41 | -0.54 | -0.10 | 0.80 |
| 5 | 15433232 | 4.16E-06 | 2.17E-02 | 0.94 | -0.48 | -0.51 | 0.31 |
| 9 | 38803433 | 4.17E-06 | 2.17E-02 | 1.48 | -0.73 | -0.19 | 0.30 |
| 6 | 11689886 | 4.18E-06 | 2.17E-02 | 1.11 | -0.38 | -0.67 | -0.33 |
| 9 | 41019868 | 4.18E-06 | 2.17E-02 | 1.63 | -0.08 | -0.92 | 0.13 |
| 1 | 37436841 | 4.24E-06 | 2.19E-02 | -0.04 | -0.62 | -0.33 | -0.30 |
| 10 | 17184223 | 4.24E-06 | 2.19E-02 | -0.46 | -0.59 | -0.05 | -0.23 |
| 6 | 12591456 | 4.25E-06 | 2.20E-02 | -0.27 | -0.31 | -0.61 | -0.46 |
| 10 | 2882599 | 4.27E-06 | 2.20E-02 | 0.06 | -0.30 | 0.68 | 0.18 |
| 5 | 20428114 | 4.28E-06 | 2.20E-02 | 1.10 | -0.51 | -0.18 | -0.33 |
| 6 | 14712630 | 4.29E-06 | 2.20E-02 | 0.33 | -0.62 | -0.44 | -0.18 |
| 9 | 38801711 | 4.29E-06 | 2.20E-02 | 1.88 | -0.77 | -0.13 | -0.07 |
| 9 | 38801721 | 4.29E-06 | 2.20E-02 | 1.88 | -0.77 | -0.13 | -0.07 |
| 6 | 16304871 | 4.32E-06 | 2.21E-02 | 0.57 | -0.52 | -0.20 | -0.23 |
| 6 | 16900673 | 4.33E-06 | 2.21E-02 | -0.67 | -0.28 | -0.16 | -0.40 |
| 1 | 40313335 | 4.33E-06 | 2.21E-02 | 0.09 | -0.03 | 0.27 | 0.75 |
| 2 | 26702532 | 4.35E-06 | 2.22E-02 | -0.18 | -0.24 | -0.57 | -0.28 |
| 2 | 6695148 | 4.37E-06 | 2.23E-02 | -0.86 | -0.68 | 0.42 | 0.12 |
| 6 | 44557095 | 4.39E-06 | 2.23E-02 | 0.78 | -0.43 | -0.38 | -0.20 |
| 12 | 20352887 | 4.41E-06 | 2.24E-02 | 0.24 | -0.72 | -0.44 | 0.12 |
| 11 | 54746240 | 4.41E-06 | 2.24E-02 | -1.15 | 0.55 | 0.05 | 0.23 |
| 7 | 26107826 | 4.42E-06 | 2.24E-02 | 0.35 | -0.59 | -0.32 | 0.34 |
| 9 | 39233058 | 4.42E-06 | 2.24E-02 | 0.85 | -0.33 | -0.14 | 0.17 |
| 6 | 16304223 | 4.46E-06 | 2.25E-02 | 0.38 | -0.64 | -0.13 | -0.19 |
| 1 | 40275379 | 4.47E-06 | 2.25E-02 | -0.20 | -0.05 | 0.34 | 0.73 |
| 6 | 15279725 | 4.49E-06 | 2.25E-02 | 0.71 | -0.41 | -0.20 | 0.27 |
| 1 | 37396073 | 4.49E-06 | 2.25E-02 | -0.06 | -0.58 | -0.31 | -0.31 |
| 1 | 37398853 | 4.49E-06 | 2.25E-02 | -0.06 | -0.58 | -0.31 | -0.31 |
| 2 | 21751226 | 4.49E-06 | 2.25E-02 | -0.59 | -0.04 | -0.58 | -0.02 |

| | | | | | | | |
|---|---|---|---|---|---|---|---|
| 4 | 72956482 | 4.49E-06 | 2.25E-02 | 0.88 | -0.33 | 0.42 | 0.28 |
| 10 | 6795053 | 4.50E-06 | 2.25E-02 | 0.06 | -0.61 | -0.55 | 0.29 |
| 6 | 42349610 | 4.50E-06 | 2.25E-02 | 0.26 | -0.85 | -0.31 | -0.51 |
| 8 | 1646095 | 4.54E-06 | 2.25E-02 | -0.01 | 0.12 | 0.40 | 0.61 |
| 6 | 15255866 | 4.55E-06 | 2.25E-02 | 0.39 | -0.63 | -0.56 | 0.18 |
| 6 | 15261288 | 4.55E-06 | 2.25E-02 | 0.39 | -0.63 | -0.56 | 0.18 |
| 6 | 15268205 | 4.55E-06 | 2.25E-02 | 0.39 | -0.63 | -0.56 | 0.18 |
| 6 | 15268853 | 4.55E-06 | 2.25E-02 | 0.39 | -0.63 | -0.56 | 0.18 |
| 6 | 15270462 | 4.55E-06 | 2.25E-02 | 0.39 | -0.63 | -0.56 | 0.18 |
| 6 | 15274099 | 4.55E-06 | 2.25E-02 | 0.39 | -0.63 | -0.56 | 0.18 |
| 6 | 15282219 | 4.55E-06 | 2.25E-02 | 0.39 | -0.63 | -0.56 | 0.18 |
| 6 | 15301008 | 4.55E-06 | 2.25E-02 | 0.39 | -0.63 | -0.56 | 0.18 |
| 8 | 20932146 | 4.56E-06 | 2.25E-02 | 0.35 | -0.04 | -0.54 | 0.23 |
| 6 | 16917419 | 4.64E-06 | 2.28E-02 | -0.72 | -0.37 | -0.29 | -0.30 |
| 5 | 58053655 | 4.64E-06 | 2.28E-02 | -0.79 | 0.26 | 0.11 | 0.46 |
| 7 | 31022488 | 4.66E-06 | 2.29E-02 | -0.50 | -0.41 | -0.16 | 0.66 |
| 6 | 15328107 | 4.68E-06 | 2.29E-02 | -0.06 | -0.93 | -0.64 | 0.25 |
| 5 | 65811093 | 4.72E-06 | 2.29E-02 | 0.18 | -0.65 | -0.50 | -0.27 |
| 5 | 25842623 | 4.73E-06 | 2.29E-02 | -0.74 | -0.46 | -0.10 | -0.14 |
| 12 | 16251499 | 4.73E-06 | 2.29E-02 | -0.59 | -0.35 | 0.30 | 0.28 |
| 10 | 29835689 | 4.73E-06 | 2.29E-02 | 0.06 | -0.16 | -0.61 | -0.38 |
| 1 | 37429234 | 4.75E-06 | 2.29E-02 | -0.10 | -0.60 | -0.32 | -0.32 |
| 1 | 37447838 | 4.75E-06 | 2.29E-02 | -0.10 | -0.60 | -0.32 | -0.32 |
| 1 | 37447937 | 4.75E-06 | 2.29E-02 | -0.10 | -0.60 | -0.32 | -0.32 |
| 1 | 37450182 | 4.75E-06 | 2.29E-02 | -0.10 | -0.60 | -0.32 | -0.32 |
| 1 | 37453175 | 4.75E-06 | 2.29E-02 | -0.10 | -0.60 | -0.32 | -0.32 |
| 1 | 37456593 | 4.75E-06 | 2.29E-02 | -0.10 | -0.60 | -0.32 | -0.32 |
| 1 | 37457093 | 4.75E-06 | 2.29E-02 | -0.10 | -0.60 | -0.32 | -0.32 |
| 1 | 37457405 | 4.75E-06 | 2.29E-02 | -0.10 | -0.60 | -0.32 | -0.32 |
| 6 | 15276904 | 4.76E-06 | 2.29E-02 | 0.56 | -0.54 | -0.29 | 0.25 |
| 6 | 15370352 | 4.76E-06 | 2.29E-02 | 0.47 | -0.53 | -0.38 | 0.27 |
| 5 | 44482264 | 4.78E-06 | 2.30E-02 | -1.18 | 0.24 | 0.06 | 0.64 |
| 8 | 9302341 | 4.81E-06 | 2.30E-02 | 0.18 | -0.73 | 0.42 | 1.26 |
| 8 | 9304183 | 4.81E-06 | 2.30E-02 | 0.18 | -0.73 | 0.42 | 1.26 |
| 8 | 9304560 | 4.81E-06 | 2.30E-02 | 0.18 | -0.73 | 0.42 | 1.26 |
| 7 | 30799468 | 4.82E-06 | 2.30E-02 | 0.30 | -0.44 | -0.05 | 0.41 |
| 1 | 34659957 | 4.87E-06 | 2.31E-02 | 0.50 | 0.21 | -0.49 | -0.87 |
| 1 | 13328039 | 4.87E-06 | 2.31E-02 | 0.59 | -0.53 | -0.05 | 0.32 |
| 1 | 26935179 | 4.88E-06 | 2.31E-02 | -0.18 | -0.16 | -0.67 | 0.03 |
| 3 | 11177623 | 4.91E-06 | 2.31E-02 | 1.07 | -0.16 | -0.28 | 0.01 |
| 12 | 16244447 | 4.92E-06 | 2.31E-02 | 0.71 | 0.42 | -0.25 | -0.13 |
| 4 | 73067430 | 4.97E-06 | 2.31E-02 | 1.31 | -1.14 | 0.04 | -0.81 |
| 12 | 10679516 | 4.98E-06 | 2.31E-02 | -1.00 | 0.25 | 0.49 | 0.35 |
| 9 | 38834184 | 4.99E-06 | 2.31E-02 | 0.39 | -0.67 | -0.33 | 0.12 |
| 6 | 44526760 | 5.00E-06 | 2.31E-02 | 0.56 | -0.65 | -0.50 | -0.26 |
| 8 | 21386035 | 5.00E-06 | 2.31E-02 | 0.80 | -0.27 | 0.03 | 0.24 |
| 10 | 18085808 | 5.05E-06 | 2.31E-02 | 0.79 | -0.16 | 0.01 | 0.33 |
| 7 | 26097030 | 5.05E-06 | 2.31E-02 | 0.40 | -0.64 | -0.22 | 0.31 |
| 8 | 9302238 | 5.09E-06 | 2.31E-02 | 0.14 | -0.75 | 0.41 | 1.22 |
| 6 | 10413857 | 5.10E-06 | 2.31E-02 | 0.73 | -0.19 | -0.73 | -0.87 |
| 6 | 10413907 | 5.10E-06 | 2.31E-02 | 0.73 | -0.19 | -0.73 | -0.87 |
| 5 | 25842273 | 5.12E-06 | 2.31E-02 | -0.67 | -0.50 | -0.14 | -0.13 |

| | | | | | | | |
|---|---|---|---|---|---|---|---|
| 1 | 40302720 | 5.12E-06 | 2.31E-02 | 0.11 | -0.09 | 0.18 | 0.75 |
| 6 | 16894520 | 5.14E-06 | 2.31E-02 | -0.79 | -0.16 | -0.15 | -0.42 |
| 5 | 25896144 | 5.17E-06 | 2.31E-02 | -0.72 | -0.44 | 0.13 | -0.06 |
| 1 | 13338736 | 5.18E-06 | 2.31E-02 | 0.53 | -0.52 | 0.02 | 0.35 |
| 1 | 13338754 | 5.18E-06 | 2.31E-02 | 0.53 | -0.52 | 0.02 | 0.35 |
| 6 | 36360191 | 5.19E-06 | 2.31E-02 | 0.82 | -0.51 | 0.18 | -0.18 |
| 6 | 36360227 | 5.19E-06 | 2.31E-02 | 0.82 | -0.51 | 0.18 | -0.18 |
| 8 | 9310670 | 5.19E-06 | 2.31E-02 | 0.03 | -0.52 | 0.50 | 1.29 |
| 8 | 9310827 | 5.19E-06 | 2.31E-02 | 0.03 | -0.52 | 0.50 | 1.29 |
| 8 | 9311250 | 5.19E-06 | 2.31E-02 | 0.03 | -0.52 | 0.50 | 1.29 |
| 8 | 9311510 | 5.19E-06 | 2.31E-02 | 0.03 | -0.52 | 0.50 | 1.29 |
| 4 | 72958518 | 5.20E-06 | 2.31E-02 | 0.79 | -0.27 | 0.55 | 0.30 |
| 4 | 72958608 | 5.20E-06 | 2.31E-02 | 0.79 | -0.27 | 0.55 | 0.30 |
| 8 | 18991593 | 5.22E-06 | 2.31E-02 | -0.26 | 1.10 | 0.77 | -0.12 |
| 2 | 29390804 | 5.22E-06 | 2.31E-02 | -0.36 | -0.17 | 0.47 | 0.44 |
| 8 | 20998819 | 5.23E-06 | 2.31E-02 | 0.94 | -0.23 | -0.48 | -0.07 |
| 11 | 33895633 | 5.25E-06 | 2.31E-02 | 1.19 | 0.09 | 0.11 | 0.14 |
| 6 | 15302899 | 5.26E-06 | 2.31E-02 | 0.51 | -0.51 | -0.34 | 0.29 |
| 6 | 15303423 | 5.26E-06 | 2.31E-02 | 0.51 | -0.51 | -0.34 | 0.29 |
| 6 | 15305567 | 5.26E-06 | 2.31E-02 | 0.51 | -0.51 | -0.34 | 0.29 |
| 6 | 15308949 | 5.26E-06 | 2.31E-02 | 0.51 | -0.51 | -0.34 | 0.29 |
| 6 | 15311459 | 5.26E-06 | 2.31E-02 | 0.51 | -0.51 | -0.34 | 0.29 |
| 6 | 15312145 | 5.26E-06 | 2.31E-02 | 0.51 | -0.51 | -0.34 | 0.29 |
| 6 | 15312367 | 5.26E-06 | 2.31E-02 | 0.51 | -0.51 | -0.34 | 0.29 |
| 6 | 15313721 | 5.26E-06 | 2.31E-02 | 0.51 | -0.51 | -0.34 | 0.29 |
| 6 | 15316979 | 5.26E-06 | 2.31E-02 | 0.51 | -0.51 | -0.34 | 0.29 |
| 6 | 15318186 | 5.26E-06 | 2.31E-02 | 0.51 | -0.51 | -0.34 | 0.29 |
| 6 | 15318614 | 5.26E-06 | 2.31E-02 | 0.51 | -0.51 | -0.34 | 0.29 |
| 6 | 15322515 | 5.26E-06 | 2.31E-02 | 0.51 | -0.51 | -0.34 | 0.29 |
| 6 | 15322867 | 5.26E-06 | 2.31E-02 | 0.51 | -0.51 | -0.34 | 0.29 |
| 6 | 15323211 | 5.26E-06 | 2.31E-02 | 0.51 | -0.51 | -0.34 | 0.29 |
| 6 | 15323624 | 5.26E-06 | 2.31E-02 | 0.51 | -0.51 | -0.34 | 0.29 |
| 6 | 15326500 | 5.26E-06 | 2.31E-02 | 0.51 | -0.51 | -0.34 | 0.29 |
| 6 | 15327994 | 5.26E-06 | 2.31E-02 | 0.51 | -0.51 | -0.34 | 0.29 |
| 6 | 15329704 | 5.26E-06 | 2.31E-02 | 0.51 | -0.51 | -0.34 | 0.29 |
| 6 | 15329757 | 5.26E-06 | 2.31E-02 | 0.51 | -0.51 | -0.34 | 0.29 |
| 6 | 15331417 | 5.26E-06 | 2.31E-02 | 0.51 | -0.51 | -0.34 | 0.29 |
| 6 | 15331780 | 5.26E-06 | 2.31E-02 | 0.51 | -0.51 | -0.34 | 0.29 |
| 6 | 15343673 | 5.26E-06 | 2.31E-02 | 0.51 | -0.51 | -0.34 | 0.29 |
| 6 | 15345543 | 5.26E-06 | 2.31E-02 | 0.51 | -0.51 | -0.34 | 0.29 |
| 6 | 15345601 | 5.26E-06 | 2.31E-02 | 0.51 | -0.51 | -0.34 | 0.29 |
| 6 | 15346587 | 5.26E-06 | 2.31E-02 | 0.51 | -0.51 | -0.34 | 0.29 |
| 6 | 15351731 | 5.26E-06 | 2.31E-02 | 0.51 | -0.51 | -0.34 | 0.29 |
| 6 | 15356473 | 5.26E-06 | 2.31E-02 | 0.51 | -0.51 | -0.34 | 0.29 |
| 6 | 15357867 | 5.26E-06 | 2.31E-02 | 0.51 | -0.51 | -0.34 | 0.29 |
| 6 | 15363692 | 5.26E-06 | 2.31E-02 | 0.51 | -0.51 | -0.34 | 0.29 |
| 6 | 15364105 | 5.26E-06 | 2.31E-02 | 0.51 | -0.51 | -0.34 | 0.29 |
| 6 | 15368320 | 5.26E-06 | 2.31E-02 | 0.51 | -0.51 | -0.34 | 0.29 |
| 6 | 15372669 | 5.26E-06 | 2.31E-02 | 0.51 | -0.51 | -0.34 | 0.29 |
| 6 | 15270903 | 5.30E-06 | 2.33E-02 | 0.67 | -0.30 | -0.40 | 0.21 |
| 6 | 45045104 | 5.33E-06 | 2.33E-02 | 0.20 | -0.46 | -0.78 | -0.40 |
| 6 | 45045282 | 5.33E-06 | 2.33E-02 | 0.20 | -0.46 | -0.78 | -0.40 |

| | | | | | | | |
|---|---|---|---|---|---|---|---|
| 6 | 45045534 | 5.33E-06 | 2.33E-02 | 0.20 | -0.46 | -0.78 | -0.40 |
| 7 | 31017086 | 5.38E-06 | 2.35E-02 | -0.56 | -0.44 | -0.04 | 0.63 |
| 6 | 15432375 | 5.39E-06 | 2.35E-02 | 0.35 | -0.38 | -0.29 | 0.43 |
| 4 | 52208859 | 5.41E-06 | 2.36E-02 | -0.48 | -0.53 | -0.11 | -0.27 |
| 5 | 38458745 | 5.43E-06 | 2.36E-02 | 0.81 | -0.51 | 0.02 | -0.42 |
| 1 | 26939948 | 5.43E-06 | 2.36E-02 | -1.41 | 0.59 | 1.29 | 0.75 |
| 10 | 5992955 | 5.45E-06 | 2.37E-02 | 0.35 | -0.60 | -0.46 | 0.07 |
| 1 | 37459159 | 5.45E-06 | 2.37E-02 | -0.12 | -0.59 | -0.31 | -0.32 |
| 9 | 5680822 | 5.46E-06 | 2.37E-02 | -0.10 | -0.68 | -0.04 | -0.28 |
| 5 | 4023915 | 5.47E-06 | 2.37E-02 | -1.41 | 0.06 | 1.16 | 1.03 |
| 4 | 19388772 | 5.48E-06 | 2.37E-02 | 1.07 | 0.66 | -0.21 | -0.08 |
| 4 | 72959054 | 5.49E-06 | 2.37E-02 | 0.64 | -0.32 | 0.60 | 0.37 |
| 8 | 47758935 | 5.50E-06 | 2.37E-02 | 0.27 | -0.22 | 0.64 | -0.11 |
| 9 | 29145259 | 5.55E-06 | 2.39E-02 | 0.85 | -0.31 | -0.13 | 0.14 |
| 6 | 47665162 | 5.55E-06 | 2.39E-02 | 0.62 | -0.51 | -0.61 | -0.37 |
| 9 | 29148400 | 5.56E-06 | 2.39E-02 | 0.71 | -0.34 | -0.17 | 0.18 |
| 5 | 38437675 | 5.62E-06 | 2.41E-02 | 0.85 | -0.13 | 0.06 | -0.58 |
| 5 | 38437692 | 5.62E-06 | 2.41E-02 | 0.85 | -0.13 | 0.06 | -0.58 |
| 12 | 16251966 | 5.62E-06 | 2.41E-02 | -0.60 | -0.31 | 0.32 | 0.29 |
| 1 | 40336422 | 5.67E-06 | 2.43E-02 | 0.13 | -0.16 | 0.00 | 0.69 |
| 6 | 53078258 | 5.73E-06 | 2.45E-02 | 1.13 | -0.08 | -0.46 | 0.03 |
| 6 | 14850139 | 5.74E-06 | 2.45E-02 | -0.86 | 0.04 | 0.24 | 0.56 |
| 8 | 21387602 | 5.77E-06 | 2.45E-02 | 0.80 | -0.23 | 0.03 | 0.27 |
| 8 | 21408082 | 5.77E-06 | 2.45E-02 | 0.80 | -0.23 | 0.03 | 0.27 |
| 8 | 21408167 | 5.77E-06 | 2.45E-02 | 0.80 | -0.23 | 0.03 | 0.27 |
| 8 | 21439335 | 5.79E-06 | 2.45E-02 | 1.42 | -0.49 | 0.06 | 0.96 |
| 6 | 53077371 | 5.79E-06 | 2.45E-02 | 1.13 | -0.09 | -0.45 | 0.02 |
| 6 | 53077604 | 5.79E-06 | 2.45E-02 | 1.13 | -0.09 | -0.45 | 0.02 |
| 6 | 53077666 | 5.79E-06 | 2.45E-02 | 1.13 | -0.09 | -0.45 | 0.02 |
| 6 | 53079875 | 5.79E-06 | 2.45E-02 | 1.13 | -0.09 | -0.45 | 0.02 |
| 8 | 56013020 | 5.80E-06 | 2.45E-02 | 0.24 | 0.29 | -0.61 | 0.16 |
| 8 | 45430552 | 5.84E-06 | 2.47E-02 | 0.16 | -0.21 | -0.67 | 0.09 |
| 7 | 26094599 | 5.85E-06 | 2.47E-02 | 0.41 | -0.57 | -0.25 | 0.36 |
| 7 | 26097156 | 5.85E-06 | 2.47E-02 | 0.28 | -0.66 | -0.30 | 0.32 |
| 6 | 5835091 | 5.87E-06 | 2.47E-02 | 0.25 | -0.83 | -0.44 | 0.23 |
| 2 | 6699209 | 5.87E-06 | 2.47E-02 | 0.61 | 0.58 | -0.21 | -0.13 |
| 5 | 42631472 | 5.89E-06 | 2.47E-02 | -0.67 | 0.19 | 0.55 | 1.16 |
| 7 | 40542119 | 5.91E-06 | 2.48E-02 | 0.13 | -0.04 | -0.39 | 0.57 |
| 4 | 35894188 | 5.94E-06 | 2.49E-02 | 0.42 | -0.24 | 0.54 | 0.87 |
| 10 | 36120612 | 5.96E-06 | 2.49E-02 | 1.21 | -0.87 | -0.44 | 0.53 |
| 6 | 380645 | 5.96E-06 | 2.49E-02 | 0.88 | -0.49 | -0.32 | 0.93 |
| 6 | 380881 | 5.96E-06 | 2.49E-02 | 0.88 | -0.49 | -0.32 | 0.93 |
| 6 | 16304427 | 5.97E-06 | 2.49E-02 | 0.49 | -0.56 | -0.21 | -0.23 |
| 11 | 43596582 | 6.02E-06 | 2.51E-02 | -0.58 | 0.76 | -1.32 | 0.56 |
| 6 | 15363896 | 6.05E-06 | 2.52E-02 | 0.55 | -0.47 | -0.33 | 0.28 |
| 7 | 16630727 | 6.08E-06 | 2.53E-02 | -0.33 | -0.61 | 0.12 | 0.24 |
| 9 | 38804357 | 6.11E-06 | 2.53E-02 | 1.67 | -0.87 | -0.25 | -0.24 |
| 4 | 80005597 | 6.14E-06 | 2.54E-02 | -1.36 | -0.22 | -0.56 | -0.11 |
| 5 | 38454663 | 6.16E-06 | 2.55E-02 | 0.73 | -0.50 | 0.04 | -0.27 |
| 5 | 38443410 | 6.17E-06 | 2.55E-02 | 0.41 | -0.67 | -0.04 | -0.13 |
| 5 | 38443413 | 6.17E-06 | 2.55E-02 | 0.41 | -0.67 | -0.04 | -0.13 |
| 6 | 15970858 | 6.17E-06 | 2.55E-02 | -0.15 | -0.22 | -0.63 | 0.31 |

| | | | | | | | |
|---|---|---|---|---|---|---|---|
| 7 | 30803089 | 6.21E-06 | 2.56E-02 | 0.40 | -0.47 | -0.17 | 0.33 |
| 12 | 16258322 | 6.24E-06 | 2.57E-02 | -0.63 | -0.31 | 0.33 | 0.26 |
| 4 | 53628882 | 6.28E-06 | 2.58E-02 | 0.68 | -0.39 | -0.42 | -0.16 |
| 10 | 40109124 | 6.30E-06 | 2.58E-02 | 0.38 | -0.67 | -0.06 | -0.44 |
| 5 | 25754136 | 6.30E-06 | 2.58E-02 | -0.56 | -0.59 | 0.03 | 0.13 |
| 5 | 25754224 | 6.30E-06 | 2.58E-02 | -0.56 | -0.59 | 0.03 | 0.13 |
| 7 | 16610698 | 6.30E-06 | 2.58E-02 | -0.36 | -0.64 | 0.04 | 0.23 |
| 1 | 13333197 | 6.37E-06 | 2.60E-02 | 0.49 | -0.61 | -0.13 | 0.29 |
| 1 | 13333227 | 6.37E-06 | 2.60E-02 | 0.49 | -0.61 | -0.13 | 0.29 |
| 1 | 13366932 | 6.37E-06 | 2.60E-02 | 0.49 | -0.61 | -0.13 | 0.29 |
| 10 | 5892862 | 6.41E-06 | 2.61E-02 | 0.38 | -0.62 | -0.39 | 0.13 |
| 4 | 19499130 | 6.48E-06 | 2.64E-02 | 1.78 | 0.20 | -0.69 | -0.28 |
| 6 | 9908101 | 6.53E-06 | 2.66E-02 | 0.43 | -0.66 | -0.25 | 0.09 |
| 7 | 16620243 | 6.54E-06 | 2.66E-02 | -0.35 | -0.54 | 0.30 | 0.19 |
| 2 | 24720597 | 6.58E-06 | 2.67E-02 | -0.17 | 0.17 | -0.73 | 0.77 |
| 6 | 15398548 | 6.62E-06 | 2.67E-02 | 0.43 | -0.36 | -0.22 | 0.44 |
| 6 | 15399114 | 6.62E-06 | 2.67E-02 | 0.43 | -0.36 | -0.22 | 0.44 |
| 6 | 15399276 | 6.62E-06 | 2.67E-02 | 0.43 | -0.36 | -0.22 | 0.44 |
| 6 | 10409049 | 6.62E-06 | 2.67E-02 | 0.55 | -0.11 | -0.73 | -0.92 |
| 6 | 10409057 | 6.62E-06 | 2.67E-02 | 0.55 | -0.11 | -0.73 | -0.92 |
| 7 | 40261307 | 6.64E-06 | 2.67E-02 | 0.53 | -0.38 | -0.51 | -0.23 |
| 6 | 15410177 | 6.65E-06 | 2.67E-02 | 0.31 | -0.43 | -0.34 | 0.42 |
| 6 | 15373923 | 6.68E-06 | 2.68E-02 | 0.54 | -0.46 | -0.31 | 0.29 |
| 9 | 35488434 | 6.69E-06 | 2.68E-02 | -0.19 | -0.80 | 0.04 | 0.19 |
| 6 | 15299055 | 6.69E-06 | 2.68E-02 | 0.37 | -0.62 | -0.58 | 0.14 |
| 1 | 37437934 | 6.72E-06 | 2.69E-02 | -0.12 | -0.58 | -0.30 | -0.32 |
| 2 | 15012585 | 6.73E-06 | 2.69E-02 | 1.18 | 0.04 | -0.30 | 0.22 |
| 5 | 2887703 | 6.73E-06 | 2.69E-02 | -0.93 | -0.29 | 0.06 | 0.04 |
| 11 | 51865543 | 6.73E-06 | 2.69E-02 | -0.07 | 0.21 | -0.44 | 0.43 |
| 6 | 15351136 | 6.76E-06 | 2.70E-02 | 0.49 | -0.52 | -0.36 | 0.27 |
| 7 | 41711019 | 6.77E-06 | 2.70E-02 | -1.10 | -0.01 | -0.18 | 0.57 |
| 6 | 16293711 | 6.80E-06 | 2.71E-02 | 0.73 | -0.54 | 0.02 | -0.17 |
| 4 | 41434369 | 6.81E-06 | 2.71E-02 | -0.98 | 0.18 | -0.87 | -0.75 |
| 11 | 33877244 | 6.83E-06 | 2.71E-02 | 0.96 | 0.21 | 0.45 | 0.37 |
| 1 | 34659953 | 6.89E-06 | 2.73E-02 | 0.64 | 0.26 | -0.53 | -0.82 |
| 7 | 26096969 | 6.90E-06 | 2.73E-02 | 0.36 | -0.61 | -0.20 | 0.30 |
| 7 | 26096988 | 6.90E-06 | 2.73E-02 | 0.36 | -0.61 | -0.20 | 0.30 |
| 2 | 18185957 | 7.01E-06 | 2.77E-02 | -0.58 | -0.09 | -0.61 | -0.77 |
| 11 | 16704561 | 7.06E-06 | 2.79E-02 | 1.69 | 0.36 | -1.06 | -0.12 |
| 10 | 29834498 | 7.07E-06 | 2.79E-02 | -0.05 | -0.12 | -0.55 | -0.43 |
| 5 | 25898578 | 7.09E-06 | 2.79E-02 | -0.67 | -0.47 | 0.07 | -0.06 |
| 5 | 25898596 | 7.09E-06 | 2.79E-02 | -0.67 | -0.47 | 0.07 | -0.06 |
| 11 | 32460943 | 7.10E-06 | 2.79E-02 | -0.69 | -0.26 | 0.34 | 0.26 |
| 5 | 73045541 | 7.20E-06 | 2.79E-02 | -0.22 | -0.29 | 0.69 | -0.42 |
| 6 | 44512372 | 7.23E-06 | 2.79E-02 | 0.66 | -0.62 | -0.46 | -0.20 |
| 8 | 27309516 | 7.24E-06 | 2.79E-02 | 0.46 | 0.22 | -0.43 | -0.48 |
| 8 | 27309517 | 7.24E-06 | 2.79E-02 | 0.46 | 0.22 | -0.43 | -0.48 |
| 8 | 36506336 | 7.28E-06 | 2.79E-02 | 0.67 | -0.20 | -0.17 | 0.30 |
| 6 | 15561999 | 7.29E-06 | 2.79E-02 | -0.24 | -1.01 | -0.72 | 0.28 |
| 11 | 34527090 | 7.29E-06 | 2.79E-02 | -0.12 | 0.36 | 0.19 | 0.61 |
| 7 | 31019979 | 7.30E-06 | 2.79E-02 | -0.57 | -0.40 | -0.04 | 0.65 |
| 7 | 31021709 | 7.30E-06 | 2.79E-02 | -0.57 | -0.40 | -0.04 | 0.65 |

| | | | | | | | |
|---|---|---|---|---|---|---|---|
| 6 | 9906020 | 7.33E-06 | 2.79E-02 | 0.43 | -0.65 | -0.23 | 0.12 |
| 6 | 9910868 | 7.33E-06 | 2.79E-02 | 0.43 | -0.65 | -0.23 | 0.12 |
| 6 | 9916901 | 7.33E-06 | 2.79E-02 | 0.43 | -0.65 | -0.23 | 0.12 |
| 6 | 9916906 | 7.33E-06 | 2.79E-02 | 0.43 | -0.65 | -0.23 | 0.12 |
| 6 | 9920377 | 7.33E-06 | 2.79E-02 | 0.43 | -0.65 | -0.23 | 0.12 |
| 5 | 34052064 | 7.37E-06 | 2.79E-02 | 0.56 | -0.26 | -0.40 | -0.28 |
| 6 | 15253437 | 7.37E-06 | 2.79E-02 | 0.56 | -0.49 | -0.35 | 0.22 |
| 6 | 15254165 | 7.37E-06 | 2.79E-02 | 0.56 | -0.49 | -0.35 | 0.22 |
| 6 | 15261411 | 7.37E-06 | 2.79E-02 | 0.56 | -0.49 | -0.35 | 0.22 |
| 6 | 15263056 | 7.37E-06 | 2.79E-02 | 0.56 | -0.49 | -0.35 | 0.22 |
| 6 | 15264841 | 7.37E-06 | 2.79E-02 | 0.56 | -0.49 | -0.35 | 0.22 |
| 6 | 15273653 | 7.37E-06 | 2.79E-02 | 0.56 | -0.49 | -0.35 | 0.22 |
| 6 | 15273973 | 7.37E-06 | 2.79E-02 | 0.56 | -0.49 | -0.35 | 0.22 |
| 6 | 15278934 | 7.37E-06 | 2.79E-02 | 0.56 | -0.49 | -0.35 | 0.22 |
| 6 | 15279229 | 7.37E-06 | 2.79E-02 | 0.56 | -0.49 | -0.35 | 0.22 |
| 6 | 15279440 | 7.37E-06 | 2.79E-02 | 0.56 | -0.49 | -0.35 | 0.22 |
| 6 | 15280239 | 7.37E-06 | 2.79E-02 | 0.56 | -0.49 | -0.35 | 0.22 |
| 6 | 15281370 | 7.37E-06 | 2.79E-02 | 0.56 | -0.49 | -0.35 | 0.22 |
| 6 | 15282242 | 7.37E-06 | 2.79E-02 | 0.56 | -0.49 | -0.35 | 0.22 |
| 6 | 15282940 | 7.37E-06 | 2.79E-02 | 0.56 | -0.49 | -0.35 | 0.22 |
| 6 | 15284902 | 7.37E-06 | 2.79E-02 | 0.56 | -0.49 | -0.35 | 0.22 |
| 6 | 15285939 | 7.37E-06 | 2.79E-02 | 0.56 | -0.49 | -0.35 | 0.22 |
| 6 | 15285993 | 7.37E-06 | 2.79E-02 | 0.56 | -0.49 | -0.35 | 0.22 |
| 6 | 15289194 | 7.37E-06 | 2.79E-02 | 0.56 | -0.49 | -0.35 | 0.22 |
| 6 | 15298793 | 7.37E-06 | 2.79E-02 | 0.56 | -0.49 | -0.35 | 0.22 |
| 6 | 15300297 | 7.37E-06 | 2.79E-02 | 0.56 | -0.49 | -0.35 | 0.22 |
| 6 | 15300719 | 7.37E-06 | 2.79E-02 | 0.56 | -0.49 | -0.35 | 0.22 |
| 1 | 7692801 | 7.40E-06 | 2.80E-02 | 0.21 | 0.54 | -0.25 | 0.41 |
| 12 | 4503472 | 7.43E-06 | 2.81E-02 | 0.13 | -0.33 | -0.66 | 0.14 |
| 8 | 21427145 | 7.44E-06 | 2.81E-02 | 1.25 | -0.48 | 0.11 | 0.83 |
| 6 | 9675191 | 7.46E-06 | 2.81E-02 | 0.16 | -0.17 | 0.31 | 0.76 |
| 9 | 41007917 | 7.46E-06 | 2.81E-02 | 1.40 | -0.06 | -1.16 | 0.01 |
| 9 | 41007963 | 7.46E-06 | 2.81E-02 | 1.40 | -0.06 | -1.16 | 0.01 |
| 8 | 9309603 | 7.50E-06 | 2.82E-02 | 0.07 | -0.76 | 0.48 | 1.20 |
| 9 | 7434706 | 7.58E-06 | 2.84E-02 | 0.33 | -0.30 | -0.09 | -0.62 |
| 4 | 27839876 | 7.58E-06 | 2.84E-02 | -0.67 | -0.59 | -0.15 | 1.52 |
| 4 | 20957575 | 7.64E-06 | 2.86E-02 | -0.98 | -0.32 | -0.24 | -0.14 |
| 6 | 16370529 | 7.65E-06 | 2.86E-02 | 0.64 | -0.53 | 0.07 | -0.16 |
| 8 | 56016594 | 7.65E-06 | 2.86E-02 | 0.18 | 0.38 | -0.57 | 0.20 |
| 6 | 44824328 | 7.67E-06 | 2.86E-02 | 0.56 | -1.01 | -0.52 | -0.19 |
| 9 | 39337208 | 7.68E-06 | 2.86E-02 | 1.04 | -0.37 | -0.51 | -0.21 |
| 2 | 6695603 | 7.69E-06 | 2.87E-02 | 0.61 | 0.54 | -0.23 | -0.16 |
| 4 | 50787177 | 7.77E-06 | 2.89E-02 | -0.21 | -0.66 | -0.62 | 0.23 |
| 6 | 13682456 | 7.81E-06 | 2.90E-02 | 0.12 | -0.41 | -0.82 | -0.48 |
| 6 | 391869 | 7.82E-06 | 2.90E-02 | 0.88 | -0.56 | 0.05 | 0.86 |
| 6 | 392062 | 7.82E-06 | 2.90E-02 | 0.88 | -0.56 | 0.05 | 0.86 |
| 9 | 7865154 | 7.83E-06 | 2.90E-02 | -0.94 | 0.19 | -0.32 | -0.18 |
| 9 | 29086854 | 7.84E-06 | 2.90E-02 | 0.97 | -0.32 | 0.15 | -0.23 |
| 6 | 27616353 | 7.87E-06 | 2.91E-02 | 1.38 | -0.70 | -0.63 | 0.03 |
| 3 | 5363454 | 7.88E-06 | 2.91E-02 | 2.19 | 0.42 | -0.59 | 0.70 |
| 11 | 41408642 | 7.88E-06 | 2.91E-02 | -0.99 | 0.34 | -0.09 | 0.80 |
| 1 | 8447688 | 7.88E-06 | 2.91E-02 | 1.37 | -0.26 | -0.77 | 0.07 |

| | | | | | | | |
|---|---|---|---|---|---|---|---|
| 1 | 26945775 | 7.89E-06 | 2.91E-02 | -0.23 | -0.16 | -0.64 | -0.06 |
| 7 | 26104292 | 7.89E-06 | 2.91E-02 | 0.35 | -0.59 | -0.30 | 0.34 |
| 10 | 29838225 | 7.94E-06 | 2.91E-02 | 0.01 | -0.17 | -0.57 | -0.39 |
| 10 | 29839935 | 7.94E-06 | 2.91E-02 | 0.01 | -0.17 | -0.57 | -0.39 |
| 9 | 24630863 | 7.95E-06 | 2.91E-02 | 0.78 | -0.34 | -0.26 | -0.09 |
| 7 | 36325771 | 7.95E-06 | 2.91E-02 | -1.16 | 0.54 | 0.16 | 0.58 |
| 7 | 36325786 | 7.95E-06 | 2.91E-02 | -1.16 | 0.54 | 0.16 | 0.58 |
| 9 | 22998134 | 7.99E-06 | 2.92E-02 | 0.72 | -0.49 | -0.15 | -0.24 |
| 9 | 51384086 | 7.99E-06 | 2.92E-02 | -0.30 | 0.60 | 0.01 | 0.46 |
| 6 | 15351666 | 7.99E-06 | 2.92E-02 | 0.55 | -0.49 | -0.33 | 0.25 |
| 6 | 42349607 | 7.99E-06 | 2.92E-02 | 0.27 | -0.79 | -0.28 | -0.57 |
| 2 | 17326614 | 8.01E-06 | 2.92E-02 | -0.63 | -0.07 | -0.55 | -0.03 |
| 6 | 15282811 | 8.05E-06 | 2.93E-02 | 0.63 | -0.46 | -0.34 | 0.19 |
| 2 | 9511519 | 8.07E-06 | 2.93E-02 | -0.13 | -0.69 | -0.70 | 0.18 |
| 2 | 9511708 | 8.07E-06 | 2.93E-02 | -0.13 | -0.69 | -0.70 | 0.18 |
| 10 | 40566044 | 8.08E-06 | 2.93E-02 | 0.71 | -0.53 | -0.30 | 0.01 |
| 2 | 11109562 | 8.10E-06 | 2.94E-02 | 2.23 | -0.48 | -0.83 | -0.36 |
| 4 | 65109984 | 8.17E-06 | 2.96E-02 | 0.57 | -0.24 | -0.29 | -0.62 |
| 6 | 49254906 | 8.18E-06 | 2.96E-02 | 0.46 | -0.84 | -0.77 | 0.04 |
| 8 | 21718711 | 8.19E-06 | 2.96E-02 | 0.67 | -0.45 | 0.50 | 0.28 |
| 6 | 15279701 | 8.21E-06 | 2.96E-02 | 0.71 | -0.36 | -0.30 | 0.21 |
| 1 | 13352430 | 8.22E-06 | 2.96E-02 | 0.55 | -0.59 | -0.02 | 0.26 |
| 1 | 13352431 | 8.22E-06 | 2.96E-02 | 0.55 | -0.59 | -0.02 | 0.26 |
| 2 | 17417846 | 8.24E-06 | 2.97E-02 | -0.12 | -0.82 | -0.19 | 0.19 |
| 5 | 44481871 | 8.28E-06 | 2.98E-02 | -1.17 | 0.25 | -0.11 | 0.62 |
| 5 | 34375527 | 8.38E-06 | 2.98E-02 | 0.19 | 0.50 | -0.51 | 0.25 |
| 1 | 37421838 | 8.38E-06 | 2.98E-02 | -0.15 | -0.56 | -0.34 | -0.31 |
| 1 | 42157587 | 8.39E-06 | 2.98E-02 | 0.69 | -0.64 | -0.35 | 0.05 |
| 7 | 26088209 | 8.39E-06 | 2.98E-02 | 0.39 | -0.59 | -0.30 | 0.32 |
| 7 | 26088460 | 8.39E-06 | 2.98E-02 | 0.39 | -0.59 | -0.30 | 0.32 |
| 7 | 26092931 | 8.39E-06 | 2.98E-02 | 0.39 | -0.59 | -0.30 | 0.32 |
| 7 | 26093380 | 8.39E-06 | 2.98E-02 | 0.39 | -0.59 | -0.30 | 0.32 |
| 7 | 26098016 | 8.39E-06 | 2.98E-02 | 0.39 | -0.59 | -0.30 | 0.32 |
| 7 | 26103731 | 8.39E-06 | 2.98E-02 | 0.39 | -0.59 | -0.30 | 0.32 |
| 5 | 58085223 | 8.39E-06 | 2.98E-02 | -0.82 | 0.00 | 0.21 | 0.43 |
| 2 | 17167027 | 8.40E-06 | 2.98E-02 | -0.56 | -0.30 | -0.22 | -0.69 |
| 2 | 17175740 | 8.40E-06 | 2.98E-02 | -0.56 | -0.30 | -0.22 | -0.69 |
| 2 | 17200408 | 8.40E-06 | 2.98E-02 | -0.56 | -0.30 | -0.22 | -0.69 |
| 6 | 44530692 | 8.42E-06 | 2.98E-02 | 0.66 | -0.62 | -0.49 | -0.17 |
| 6 | 29628523 | 8.42E-06 | 2.98E-02 | -0.57 | -0.10 | -1.19 | -0.71 |
| 12 | 4498270 | 8.42E-06 | 2.98E-02 | 0.06 | -0.27 | -0.67 | 0.17 |
| 2 | 4801257 | 8.46E-06 | 2.99E-02 | -0.77 | 0.33 | 0.06 | 0.92 |
| 6 | 44520451 | 8.47E-06 | 2.99E-02 | 0.65 | -0.64 | -0.45 | -0.19 |
| 6 | 15154166 | 8.48E-06 | 2.99E-02 | -0.31 | -1.08 | -0.73 | 0.27 |
| 12 | 4498051 | 8.48E-06 | 2.99E-02 | 0.04 | -0.32 | -0.69 | 0.08 |
| 6 | 32517118 | 8.49E-06 | 2.99E-02 | -0.50 | 0.13 | 0.53 | 0.53 |
| 1 | 29710560 | 8.49E-06 | 2.99E-02 | -1.02 | 0.85 | -0.45 | 0.26 |
| 6 | 10037823 | 8.55E-06 | 3.00E-02 | 0.39 | -0.69 | -0.65 | -0.73 |
| 6 | 10404304 | 8.56E-06 | 3.00E-02 | 0.48 | -0.12 | -0.69 | -0.92 |
| 6 | 15402115 | 8.57E-06 | 3.00E-02 | 0.33 | -0.41 | -0.30 | 0.41 |
| 1 | 7144808 | 8.57E-06 | 3.00E-02 | 0.14 | -0.19 | -0.68 | 0.47 |
| 6 | 10048730 | 8.57E-06 | 3.00E-02 | 0.61 | -0.70 | -0.66 | -0.73 |

| | | | | | | | |
|---|---|---|---|---|---|---|---|
| 6 | 10049847 | 8.57E-06 | 3.00E-02 | 0.61 | -0.70 | -0.66 | -0.73 |
| 6 | 21649319 | 8.60E-06 | 3.00E-02 | 1.18 | -0.48 | -0.46 | -0.17 |
| 8 | 62052088 | 8.60E-06 | 3.00E-02 | 0.01 | 0.13 | -0.63 | -0.32 |
| 9 | 32367102 | 8.65E-06 | 3.02E-02 | -0.07 | 0.01 | 0.01 | 0.66 |
| 5 | 38445001 | 8.65E-06 | 3.02E-02 | 0.83 | -0.39 | 0.08 | -0.34 |
| 4 | 33968317 | 8.69E-06 | 3.02E-02 | 2.28 | -0.12 | 0.08 | 0.52 |
| 5 | 25756896 | 8.70E-06 | 3.02E-02 | -0.47 | -0.66 | -0.12 | 0.29 |
| 12 | 4424398 | 8.73E-06 | 3.02E-02 | 0.64 | 0.23 | 0.09 | 0.48 |
| 6 | 44510087 | 8.73E-06 | 3.02E-02 | 0.61 | -0.51 | -0.47 | -0.22 |
| 1 | 13324328 | 8.75E-06 | 3.02E-02 | 0.52 | -0.58 | -0.05 | 0.30 |
| 1 | 13331080 | 8.75E-06 | 3.02E-02 | 0.52 | -0.58 | -0.05 | 0.30 |
| 1 | 13331291 | 8.75E-06 | 3.02E-02 | 0.52 | -0.58 | -0.05 | 0.30 |
| 1 | 13331798 | 8.75E-06 | 3.02E-02 | 0.52 | -0.58 | -0.05 | 0.30 |
| 1 | 13340643 | 8.75E-06 | 3.02E-02 | 0.52 | -0.58 | -0.05 | 0.30 |
| 6 | 42412185 | 8.86E-06 | 3.06E-02 | 0.26 | -0.77 | -0.26 | -0.67 |
| 2 | 6694788 | 8.89E-06 | 3.06E-02 | -0.93 | -0.61 | 0.40 | 0.12 |
| 2 | 6694825 | 8.89E-06 | 3.06E-02 | -0.93 | -0.61 | 0.40 | 0.12 |
| 6 | 53305113 | 8.93E-06 | 3.08E-02 | -0.41 | 0.23 | -0.62 | -0.42 |
| 7 | 16611770 | 8.96E-06 | 3.08E-02 | -0.52 | -0.55 | 0.15 | 0.16 |
| 11 | 34818490 | 8.99E-06 | 3.09E-02 | -0.17 | -0.31 | -0.26 | 0.47 |
| 11 | 47326645 | 8.99E-06 | 3.09E-02 | 1.06 | -0.53 | -0.41 | -0.12 |
| 9 | 6404151 | 9.00E-06 | 3.09E-02 | 0.10 | 0.32 | 0.59 | 0.24 |
| 12 | 4500608 | 9.01E-06 | 3.09E-02 | 0.32 | -0.30 | -0.72 | 0.07 |
| 7 | 40539340 | 9.07E-06 | 3.09E-02 | 0.08 | -0.06 | -0.39 | 0.58 |
| 7 | 40540273 | 9.07E-06 | 3.09E-02 | 0.08 | -0.06 | -0.39 | 0.58 |
| 7 | 40540988 | 9.07E-06 | 3.09E-02 | 0.08 | -0.06 | -0.39 | 0.58 |
| 7 | 40543866 | 9.07E-06 | 3.09E-02 | 0.08 | -0.06 | -0.39 | 0.58 |
| 7 | 40544985 | 9.07E-06 | 3.09E-02 | 0.08 | -0.06 | -0.39 | 0.58 |
| 8 | 28173592 | 9.09E-06 | 3.10E-02 | 0.57 | -0.39 | -0.13 | 0.72 |
| 6 | 16374824 | 9.14E-06 | 3.11E-02 | 0.67 | -0.49 | -0.16 | -0.21 |
| 5 | 59108084 | 9.15E-06 | 3.11E-02 | -0.03 | -0.05 | -0.46 | -0.45 |
| 4 | 51826021 | 9.17E-06 | 3.12E-02 | 0.88 | -0.26 | -0.52 | 0.24 |
| 6 | 6374231 | 9.22E-06 | 3.13E-02 | 0.68 | -0.56 | -0.25 | -0.06 |
| 6 | 10414158 | 9.23E-06 | 3.13E-02 | 0.41 | -0.25 | -0.76 | -0.88 |
| 6 | 10414160 | 9.23E-06 | 3.13E-02 | 0.41 | -0.25 | -0.76 | -0.88 |
| 4 | 46220041 | 9.24E-06 | 3.13E-02 | 0.89 | -0.25 | -0.33 | 0.38 |
| 10 | 15339138 | 9.27E-06 | 3.14E-02 | 0.71 | -0.13 | 0.62 | 0.37 |
| 9 | 27985598 | 9.30E-06 | 3.14E-02 | 1.39 | -0.66 | -0.44 | 0.07 |
| 4 | 30260933 | 9.31E-06 | 3.14E-02 | 0.83 | 0.20 | -0.62 | -0.33 |
| 11 | 51862426 | 9.39E-06 | 3.17E-02 | 0.33 | 0.25 | -0.29 | 0.46 |
| 2 | 6696166 | 9.42E-06 | 3.17E-02 | 0.64 | 0.52 | -0.26 | -0.13 |
| 10 | 6556866 | 9.44E-06 | 3.18E-02 | 0.10 | 0.05 | 0.56 | -0.84 |
| 6 | 15305220 | 9.44E-06 | 3.18E-02 | 0.57 | -0.49 | -0.28 | 0.26 |
| 7 | 26797325 | 9.45E-06 | 3.18E-02 | 0.87 | -0.51 | 0.17 | 0.26 |
| 6 | 16952325 | 9.47E-06 | 3.18E-02 | 0.15 | -0.12 | 0.67 | -0.10 |
| 6 | 53045230 | 9.51E-06 | 3.19E-02 | 1.54 | -0.16 | -0.23 | 0.16 |
| 6 | 15342470 | 9.51E-06 | 3.19E-02 | 0.53 | -0.49 | -0.33 | 0.26 |
| 9 | 53501410 | 9.52E-06 | 3.19E-02 | 0.96 | -0.26 | 0.33 | -0.23 |
| 6 | 15303344 | 9.53E-06 | 3.19E-02 | 0.52 | -0.44 | -0.33 | 0.31 |
| 6 | 10388402 | 9.54E-06 | 3.19E-02 | 0.48 | -0.38 | -0.53 | -0.82 |
| 6 | 42347730 | 9.55E-06 | 3.19E-02 | 0.22 | -0.71 | -0.35 | -0.58 |
| 7 | 36325837 | 9.57E-06 | 3.19E-02 | -1.14 | 0.55 | 0.17 | 0.57 |

| | | | | | | | |
|---|---|---|---|---|---|---|---|
| 6 | 27580991 | 9.59E-06 | 3.20E-02 | -0.30 | 0.25 | 0.40 | 0.86 |
| 1 | 24577230 | 9.62E-06 | 3.20E-02 | 0.88 | -0.68 | -0.13 | 0.60 |
| 8 | 47731295 | 9.63E-06 | 3.20E-02 | 0.39 | -0.24 | 0.56 | -0.18 |
| 8 | 47731301 | 9.63E-06 | 3.20E-02 | 0.39 | -0.24 | 0.56 | -0.18 |
| 9 | 38798928 | 9.64E-06 | 3.20E-02 | 1.08 | -0.38 | -0.10 | 0.18 |
| 3 | 8021072 | 9.67E-06 | 3.21E-02 | -0.72 | -0.16 | 0.66 | 0.16 |
| 8 | 43920274 | 9.68E-06 | 3.21E-02 | 0.46 | -0.27 | 0.51 | -0.16 |
| 11 | 22616898 | 9.69E-06 | 3.21E-02 | -0.80 | 0.13 | 0.58 | 0.34 |
| 8 | 21410663 | 9.74E-06 | 3.22E-02 | 0.81 | -0.29 | 0.16 | 0.20 |
| 8 | 54880020 | 9.76E-06 | 3.23E-02 | 1.14 | 0.82 | -0.22 | 0.10 |
| 2 | 14149176 | 9.76E-06 | 3.23E-02 | 0.09 | -0.26 | 0.32 | 1.09 |
| 12 | 15373474 | 9.88E-06 | 3.23E-02 | -0.38 | -0.32 | 0.57 | 0.26 |
| 12 | 20295830 | 9.90E-06 | 3.23E-02 | -1.04 | 0.26 | 0.74 | 1.07 |
| 11 | 52558033 | 9.92E-06 | 3.23E-02 | 0.97 | -0.20 | -0.32 | -0.20 |
| 10 | 54841407 | 9.97E-06 | 3.23E-02 | -0.25 | 0.83 | -0.65 | 0.96 |
| 6 | 45040062 | 9.99E-06 | 3.23E-02 | 0.20 | -0.46 | -0.73 | -0.37 |
| 6 | 44510663 | 1.00E-05 | 3.23E-02 | 0.62 | -0.63 | -0.49 | -0.20 |
| 6 | 44510739 | 1.00E-05 | 3.23E-02 | 0.62 | -0.63 | -0.49 | -0.20 |
| 6 | 44510785 | 1.00E-05 | 3.23E-02 | 0.62 | -0.63 | -0.49 | -0.20 |
| 6 | 44512870 | 1.00E-05 | 3.23E-02 | 0.62 | -0.63 | -0.49 | -0.20 |
| 6 | 44513187 | 1.00E-05 | 3.23E-02 | 0.62 | -0.63 | -0.49 | -0.20 |
| 6 | 44513987 | 1.00E-05 | 3.23E-02 | 0.62 | -0.63 | -0.49 | -0.20 |
| 6 | 44514022 | 1.00E-05 | 3.23E-02 | 0.62 | -0.63 | -0.49 | -0.20 |
| 6 | 44515060 | 1.00E-05 | 3.23E-02 | 0.62 | -0.63 | -0.49 | -0.20 |
| 6 | 44519914 | 1.00E-05 | 3.23E-02 | 0.62 | -0.63 | -0.49 | -0.20 |
| 6 | 44520176 | 1.00E-05 | 3.23E-02 | 0.62 | -0.63 | -0.49 | -0.20 |
| 6 | 44522982 | 1.00E-05 | 3.23E-02 | 0.62 | -0.63 | -0.49 | -0.20 |
| 6 | 44524092 | 1.00E-05 | 3.23E-02 | 0.62 | -0.63 | -0.49 | -0.20 |
| 6 | 44526324 | 1.00E-05 | 3.23E-02 | 0.62 | -0.63 | -0.49 | -0.20 |
| 6 | 44526437 | 1.00E-05 | 3.23E-02 | 0.62 | -0.63 | -0.49 | -0.20 |
| 6 | 44529576 | 1.00E-05 | 3.23E-02 | 0.62 | -0.63 | -0.49 | -0.20 |
| 6 | 44529684 | 1.00E-05 | 3.23E-02 | 0.62 | -0.63 | -0.49 | -0.20 |
| 6 | 44529799 | 1.00E-05 | 3.23E-02 | 0.62 | -0.63 | -0.49 | -0.20 |
| 6 | 44530676 | 1.00E-05 | 3.23E-02 | 0.62 | -0.63 | -0.49 | -0.20 |
| 6 | 44532081 | 1.00E-05 | 3.23E-02 | 0.62 | -0.63 | -0.49 | -0.20 |
| 12 | 16240762 | 1.00E-05 | 3.24E-02 | -0.63 | -0.29 | 0.32 | 0.26 |
| 6 | 15286037 | 1.01E-05 | 3.24E-02 | -0.35 | -1.13 | -0.76 | 0.30 |
| 6 | 16926012 | 1.01E-05 | 3.24E-02 | 0.27 | -0.11 | 0.65 | 0.02 |
| 8 | 47680233 | 1.01E-05 | 3.24E-02 | 0.51 | -0.21 | 0.60 | 0.00 |
| 5 | 11342053 | 1.01E-05 | 3.24E-02 | -1.54 | -0.07 | 0.19 | -0.25 |
| 1 | 50197196 | 1.01E-05 | 3.24E-02 | 1.60 | -0.01 | -0.23 | 0.41 |
| 8 | 27410019 | 1.01E-05 | 3.25E-02 | 0.28 | 0.59 | -0.61 | -0.25 |
| 2 | 17194813 | 1.01E-05 | 3.25E-02 | -0.26 | -0.32 | -0.36 | -0.49 |
| 9 | 30667089 | 1.02E-05 | 3.26E-02 | 0.56 | 0.84 | -1.20 | -0.50 |
| 9 | 50950919 | 1.02E-05 | 3.26E-02 | 0.68 | -0.84 | 0.09 | -0.08 |
| 5 | 44480732 | 1.02E-05 | 3.26E-02 | -1.14 | 0.22 | 0.06 | 0.61 |
| 5 | 58047500 | 1.02E-05 | 3.27E-02 | 0.02 | -0.41 | 0.05 | 0.44 |
| 11 | 33782435 | 1.03E-05 | 3.28E-02 | 0.88 | -0.07 | 0.49 | 0.15 |
| 10 | 40601289 | 1.03E-05 | 3.28E-02 | 0.49 | -0.63 | -0.17 | 0.08 |
| 6 | 49814651 | 1.03E-05 | 3.28E-02 | 0.95 | -0.05 | -0.13 | 0.14 |
| 2 | 25596757 | 1.03E-05 | 3.29E-02 | -0.57 | 0.80 | 0.48 | 1.14 |
| 12 | 16258502 | 1.03E-05 | 3.29E-02 | -0.63 | -0.32 | 0.32 | 0.23 |

| | | | | | | | |
|---|---|---|---|---|---|---|---|
| 6 | 18817728 | 1.04E-05 | 3.29E-02 | 0.27 | -0.66 | -0.39 | 0.26 |
| 5 | 25750731 | 1.04E-05 | 3.30E-02 | -0.36 | -0.75 | -0.22 | 0.27 |
| 5 | 25750745 | 1.04E-05 | 3.30E-02 | -0.36 | -0.75 | -0.22 | 0.27 |
| 5 | 25750768 | 1.04E-05 | 3.30E-02 | -0.36 | -0.75 | -0.22 | 0.27 |
| 6 | 34160521 | 1.04E-05 | 3.31E-02 | 1.01 | 0.51 | -0.47 | 0.02 |
| 2 | 8433809 | 1.05E-05 | 3.32E-02 | 0.76 | -0.68 | -0.28 | -0.18 |
| 9 | 24640218 | 1.05E-05 | 3.32E-02 | 0.78 | -0.35 | -0.23 | -0.07 |
| 10 | 29838558 | 1.05E-05 | 3.32E-02 | 0.05 | -0.14 | -0.58 | -0.37 |
| 10 | 29838626 | 1.05E-05 | 3.32E-02 | 0.05 | -0.14 | -0.58 | -0.37 |
| 11 | 51964732 | 1.05E-05 | 3.32E-02 | -0.14 | 0.24 | -0.34 | 0.41 |
| 1 | 37440222 | 1.05E-05 | 3.32E-02 | -0.18 | -0.56 | -0.22 | -0.33 |
| 5 | 66165224 | 1.06E-05 | 3.32E-02 | 2.09 | 0.03 | -0.51 | -0.03 |
| 8 | 36636253 | 1.06E-05 | 3.32E-02 | 0.61 | -0.33 | -0.20 | 0.26 |
| 5 | 25742660 | 1.07E-05 | 3.32E-02 | -0.47 | -0.71 | -0.19 | 0.27 |
| 5 | 25742741 | 1.07E-05 | 3.32E-02 | -0.47 | -0.71 | -0.19 | 0.27 |
| 5 | 25742983 | 1.07E-05 | 3.32E-02 | -0.47 | -0.71 | -0.19 | 0.27 |
| 5 | 25744119 | 1.07E-05 | 3.32E-02 | -0.47 | -0.71 | -0.19 | 0.27 |
| 5 | 25745374 | 1.07E-05 | 3.32E-02 | -0.47 | -0.71 | -0.19 | 0.27 |
| 5 | 25746709 | 1.07E-05 | 3.32E-02 | -0.47 | -0.71 | -0.19 | 0.27 |
| 5 | 25746741 | 1.07E-05 | 3.32E-02 | -0.47 | -0.71 | -0.19 | 0.27 |
| 5 | 25749328 | 1.07E-05 | 3.32E-02 | -0.47 | -0.71 | -0.19 | 0.27 |
| 5 | 25749607 | 1.07E-05 | 3.32E-02 | -0.47 | -0.71 | -0.19 | 0.27 |
| 5 | 25757166 | 1.07E-05 | 3.32E-02 | -0.47 | -0.71 | -0.19 | 0.27 |
| 5 | 25757515 | 1.07E-05 | 3.32E-02 | -0.47 | -0.71 | -0.19 | 0.27 |
| 5 | 25758476 | 1.07E-05 | 3.32E-02 | -0.47 | -0.71 | -0.19 | 0.27 |
| 1 | 42359109 | 1.07E-05 | 3.32E-02 | -0.69 | 0.50 | -1.02 | -0.96 |
| 6 | 42325331 | 1.07E-05 | 3.32E-02 | 0.39 | -0.69 | -0.45 | -0.57 |
| 1 | 28056459 | 1.07E-05 | 3.33E-02 | -0.29 | 0.40 | -0.64 | -0.48 |
| 2 | 30190998 | 1.07E-05 | 3.33E-02 | 0.43 | -0.11 | -0.54 | 0.19 |
| 5 | 64527542 | 1.08E-05 | 3.33E-02 | -0.59 | -0.49 | -0.20 | -0.19 |
| 9 | 39510710 | 1.08E-05 | 3.33E-02 | 1.12 | -0.80 | -0.97 | -0.51 |
| 6 | 45041880 | 1.08E-05 | 3.34E-02 | 0.15 | -0.48 | -0.75 | -0.33 |
| 7 | 37175599 | 1.08E-05 | 3.35E-02 | -0.01 | -0.27 | 0.92 | 0.07 |
| 7 | 26082448 | 1.09E-05 | 3.35E-02 | 0.37 | -0.59 | -0.32 | 0.30 |
| 7 | 26082490 | 1.09E-05 | 3.35E-02 | 0.37 | -0.59 | -0.32 | 0.30 |
| 6 | 34140073 | 1.09E-05 | 3.36E-02 | 1.02 | 0.44 | -0.49 | 0.12 |
| 8 | 21507987 | 1.09E-05 | 3.36E-02 | 1.25 | -0.38 | 0.11 | 0.25 |
| 4 | 72956734 | 1.09E-05 | 3.36E-02 | 0.91 | -0.32 | 0.42 | 0.25 |
| 7 | 30966820 | 1.09E-05 | 3.36E-02 | -0.26 | -0.26 | -0.12 | 0.52 |
| 1 | 14358212 | 1.10E-05 | 3.37E-02 | -0.12 | 0.04 | -0.03 | 0.62 |
| 8 | 36521615 | 1.10E-05 | 3.37E-02 | 0.57 | -0.30 | -0.28 | 0.27 |
| 2 | 9110118 | 1.10E-05 | 3.37E-02 | -1.24 | 0.27 | 0.36 | -0.49 |
| 5 | 65813745 | 1.10E-05 | 3.37E-02 | 0.27 | -0.56 | -0.50 | -0.32 |
| 8 | 44075188 | 1.10E-05 | 3.37E-02 | 0.82 | -0.26 | 0.29 | -0.29 |
| 6 | 41078303 | 1.10E-05 | 3.37E-02 | 0.08 | -0.16 | -0.12 | -0.78 |
| 6 | 15353920 | 1.10E-05 | 3.38E-02 | 0.44 | -0.53 | -0.39 | 0.28 |
| 8 | 20932183 | 1.10E-05 | 3.38E-02 | 0.37 | -0.02 | -0.51 | 0.23 |
| 6 | 28499584 | 1.10E-05 | 3.38E-02 | 0.09 | 0.66 | -0.50 | 0.48 |
| 8 | 35942530 | 1.11E-05 | 3.39E-02 | 0.59 | -0.25 | -0.01 | 0.37 |
| 3 | 3608570 | 1.12E-05 | 3.40E-02 | 0.46 | 0.10 | -0.56 | -0.88 |
| 10 | 27759240 | 1.12E-05 | 3.40E-02 | 0.68 | 0.25 | -0.49 | -0.19 |
| 6 | 6373339 | 1.12E-05 | 3.40E-02 | 0.62 | -0.60 | -0.27 | -0.08 |

| | | | | | | | |
|---|---|---|---|---|---|---|---|
| 6 | 6373659 | 1.12E-05 | 3.40E-02 | 0.62 | -0.60 | -0.27 | -0.08 |
| 6 | 6376526 | 1.12E-05 | 3.40E-02 | 0.62 | -0.60 | -0.27 | -0.08 |
| 6 | 6376616 | 1.12E-05 | 3.40E-02 | 0.62 | -0.60 | -0.27 | -0.08 |
| 12 | 4497834 | 1.12E-05 | 3.40E-02 | 0.05 | -0.31 | -0.68 | 0.08 |
| 11 | 34465340 | 1.13E-05 | 3.41E-02 | 0.41 | 0.49 | 0.52 | 0.56 |
| 4 | 18693213 | 1.13E-05 | 3.42E-02 | -0.95 | 0.42 | 0.85 | -0.31 |
| 9 | 2030676 | 1.13E-05 | 3.42E-02 | 0.14 | 0.77 | 0.25 | -0.29 |
| 1 | 37438204 | 1.13E-05 | 3.42E-02 | 0.23 | -0.55 | -0.32 | -0.37 |
| 11 | 19533584 | 1.14E-05 | 3.43E-02 | 0.79 | -0.58 | -1.24 | -0.78 |
| 6 | 44514107 | 1.14E-05 | 3.44E-02 | 0.85 | -0.45 | -0.27 | -0.15 |
| 6 | 7148277 | 1.14E-05 | 3.44E-02 | 0.49 | -0.37 | -0.35 | 0.28 |
| 8 | 46238187 | 1.14E-05 | 3.44E-02 | -0.21 | -0.48 | -0.40 | -0.45 |
| 12 | 457480 | 1.14E-05 | 3.44E-02 | 0.19 | -0.10 | -0.57 | -0.31 |
| 9 | 37651757 | 1.14E-05 | 3.44E-02 | -1.12 | 0.44 | 0.36 | 0.58 |
| 7 | 30944108 | 1.15E-05 | 3.44E-02 | 0.24 | -0.30 | 0.05 | 0.45 |
| 7 | 30944880 | 1.15E-05 | 3.44E-02 | 0.24 | -0.30 | 0.05 | 0.45 |
| 6 | 44524793 | 1.15E-05 | 3.46E-02 | 0.58 | -0.64 | -0.48 | -0.20 |
| 8 | 49079543 | 1.16E-05 | 3.47E-02 | 0.50 | -0.27 | -0.56 | 0.03 |
| 8 | 21358951 | 1.16E-05 | 3.47E-02 | 1.54 | -0.69 | 0.34 | 0.51 |
| 2 | 1880078 | 1.16E-05 | 3.47E-02 | 0.33 | -0.59 | -0.58 | 0.27 |
| 6 | 16266719 | 1.16E-05 | 3.47E-02 | 0.25 | -0.55 | -0.36 | -0.38 |
| 6 | 15321656 | 1.16E-05 | 3.48E-02 | -0.41 | -1.22 | -0.87 | 0.33 |
| 9 | 26985103 | 1.17E-05 | 3.49E-02 | 1.58 | 0.47 | -1.36 | -0.38 |
| 4 | 83135853 | 1.17E-05 | 3.49E-02 | -0.31 | 0.35 | -1.02 | -0.39 |
| 6 | 40708517 | 1.17E-05 | 3.50E-02 | -0.23 | 0.41 | 0.64 | 0.97 |
| 6 | 40708617 | 1.17E-05 | 3.50E-02 | -0.23 | 0.41 | 0.64 | 0.97 |
| 6 | 28499940 | 1.17E-05 | 3.50E-02 | 0.03 | 0.64 | -0.47 | 0.46 |
| 6 | 10414644 | 1.18E-05 | 3.50E-02 | 0.59 | -0.15 | -0.62 | -0.94 |
| 6 | 15354604 | 1.18E-05 | 3.50E-02 | 0.46 | -0.51 | -0.35 | 0.29 |
| 6 | 15372332 | 1.18E-05 | 3.50E-02 | 0.46 | -0.51 | -0.35 | 0.29 |
| 6 | 6368995 | 1.18E-05 | 3.51E-02 | 0.69 | -0.56 | -0.32 | -0.03 |
| 6 | 5004655 | 1.20E-05 | 3.54E-02 | 0.12 | -0.64 | 0.02 | 1.22 |
| 11 | 39777999 | 1.20E-05 | 3.54E-02 | 0.80 | 0.06 | -0.22 | 0.23 |
| 6 | 34306985 | 1.21E-05 | 3.54E-02 | 0.37 | 1.57 | -0.97 | -0.18 |
| 5 | 25774544 | 1.21E-05 | 3.54E-02 | -0.71 | -0.45 | 0.07 | -0.07 |
| 5 | 25775787 | 1.21E-05 | 3.54E-02 | -0.71 | -0.45 | 0.07 | -0.07 |
| 5 | 25788309 | 1.21E-05 | 3.54E-02 | -0.71 | -0.45 | 0.07 | -0.07 |
| 5 | 25798909 | 1.21E-05 | 3.54E-02 | -0.71 | -0.45 | 0.07 | -0.07 |
| 5 | 25803384 | 1.21E-05 | 3.54E-02 | -0.71 | -0.45 | 0.07 | -0.07 |
| 5 | 25804392 | 1.21E-05 | 3.54E-02 | -0.71 | -0.45 | 0.07 | -0.07 |
| 5 | 25805203 | 1.21E-05 | 3.54E-02 | -0.71 | -0.45 | 0.07 | -0.07 |
| 5 | 25805357 | 1.21E-05 | 3.54E-02 | -0.71 | -0.45 | 0.07 | -0.07 |
| 5 | 25812152 | 1.21E-05 | 3.54E-02 | -0.71 | -0.45 | 0.07 | -0.07 |
| 5 | 25812325 | 1.21E-05 | 3.54E-02 | -0.71 | -0.45 | 0.07 | -0.07 |
| 5 | 25897576 | 1.21E-05 | 3.54E-02 | -0.71 | -0.45 | 0.07 | -0.07 |
| 5 | 25897700 | 1.21E-05 | 3.54E-02 | -0.71 | -0.45 | 0.07 | -0.07 |
| 10 | 44291880 | 1.21E-05 | 3.55E-02 | -0.65 | 0.61 | -0.77 | -0.18 |
| 10 | 44292678 | 1.21E-05 | 3.55E-02 | -0.65 | 0.61 | -0.77 | -0.18 |
| 10 | 6709881 | 1.21E-05 | 3.55E-02 | 0.81 | -0.05 | -0.25 | -0.36 |
| 8 | 44912058 | 1.22E-05 | 3.55E-02 | 0.18 | 0.45 | -0.61 | -0.04 |
| 8 | 21411576 | 1.22E-05 | 3.55E-02 | 0.79 | -0.28 | 0.19 | 0.21 |
| 9 | 20739161 | 1.22E-05 | 3.56E-02 | 1.01 | -0.52 | 0.04 | -0.72 |

| | | | | | | | |
|---|---|---|---|---|---|---|---|
| 9 | 20739169 | 1.22E-05 | 3.56E-02 | 1.01 | -0.52 | 0.04 | -0.72 |
| 5 | 25753459 | 1.22E-05 | 3.56E-02 | -0.58 | -0.55 | 0.03 | 0.15 |
| 6 | 15331946 | 1.23E-05 | 3.57E-02 | 0.41 | -0.52 | -0.42 | 0.29 |
| 8 | 52864158 | 1.23E-05 | 3.58E-02 | -0.14 | 0.88 | -0.78 | 0.08 |
| 6 | 11689710 | 1.23E-05 | 3.58E-02 | 1.08 | -0.31 | -0.64 | -0.32 |
| 6 | 11689786 | 1.23E-05 | 3.58E-02 | 1.08 | -0.31 | -0.64 | -0.32 |
| 11 | 51897615 | 1.23E-05 | 3.58E-02 | 0.27 | 0.20 | -0.55 | 0.23 |
| 5 | 38440044 | 1.24E-05 | 3.58E-02 | 0.78 | -0.39 | 0.15 | -0.38 |
| 8 | 27308090 | 1.25E-05 | 3.61E-02 | 0.31 | 0.34 | -0.42 | -0.47 |
| 8 | 33382441 | 1.25E-05 | 3.62E-02 | -0.90 | 0.23 | -1.56 | 1.28 |
| 5 | 66743106 | 1.25E-05 | 3.62E-02 | -0.36 | -0.99 | -0.64 | 0.28 |
| 2 | 4255473 | 1.26E-05 | 3.63E-02 | -0.29 | -0.08 | -0.28 | -0.54 |
| 6 | 20458228 | 1.26E-05 | 3.63E-02 | 0.81 | -0.34 | 0.37 | 0.52 |
| 9 | 20754756 | 1.26E-05 | 3.63E-02 | -1.01 | 0.53 | 0.59 | -0.19 |
| 9 | 41018174 | 1.26E-05 | 3.63E-02 | 1.26 | -0.02 | -0.95 | 0.22 |
| 10 | 40562184 | 1.26E-05 | 3.63E-02 | 0.64 | -0.54 | -0.23 | 0.03 |
| 12 | 1845396 | 1.26E-05 | 3.63E-02 | 0.47 | 0.27 | -0.21 | 0.50 |
| 6 | 15408605 | 1.26E-05 | 3.63E-02 | 0.30 | -0.42 | -0.28 | 0.43 |
| 6 | 15408623 | 1.26E-05 | 3.63E-02 | 0.30 | -0.42 | -0.28 | 0.43 |
| 5 | 25754569 | 1.26E-05 | 3.63E-02 | -0.55 | -0.54 | 0.00 | 0.24 |
| 6 | 12603385 | 1.27E-05 | 3.63E-02 | -0.16 | -0.37 | -0.57 | -0.44 |
| 2 | 6690590 | 1.27E-05 | 3.63E-02 | -0.79 | -0.65 | 0.50 | 0.10 |
| 1 | 34679127 | 1.27E-05 | 3.64E-02 | 0.46 | 0.34 | -0.49 | -0.74 |
| 6 | 44532431 | 1.27E-05 | 3.65E-02 | 0.61 | -0.64 | -0.47 | -0.19 |
| 6 | 44532438 | 1.27E-05 | 3.65E-02 | 0.61 | -0.64 | -0.47 | -0.19 |
| 6 | 18527757 | 1.28E-05 | 3.66E-02 | 1.10 | -0.41 | -0.20 | 0.27 |
| 11 | 58815989 | 1.29E-05 | 3.68E-02 | -0.13 | -0.60 | -0.56 | 0.29 |
| 9 | 29216884 | 1.29E-05 | 3.68E-02 | 1.03 | -0.15 | 0.19 | -0.10 |
| 9 | 29216931 | 1.29E-05 | 3.68E-02 | 1.03 | -0.15 | 0.19 | -0.10 |
| 9 | 38083027 | 1.30E-05 | 3.70E-02 | -1.67 | 0.40 | 0.33 | 0.48 |
| 8 | 22753657 | 1.30E-05 | 3.70E-02 | 0.37 | -0.59 | 0.14 | 0.39 |
| 8 | 20927267 | 1.31E-05 | 3.72E-02 | 0.54 | -0.08 | -0.43 | 0.22 |
| 4 | 21388718 | 1.31E-05 | 3.73E-02 | -1.20 | 0.55 | -0.08 | 0.99 |
| 2 | 14110930 | 1.31E-05 | 3.73E-02 | 0.19 | -0.68 | -0.51 | 0.29 |
| 7 | 31038387 | 1.32E-05 | 3.73E-02 | -0.13 | -0.67 | -0.32 | 0.51 |
| 9 | 29567007 | 1.32E-05 | 3.73E-02 | 0.71 | -0.19 | 0.18 | -0.63 |
| 6 | 15326306 | 1.32E-05 | 3.73E-02 | 0.49 | -0.50 | -0.40 | 0.22 |
| 1 | 8464914 | 1.32E-05 | 3.73E-02 | 1.04 | 0.26 | -0.26 | -0.37 |
| 4 | 35894860 | 1.32E-05 | 3.74E-02 | 0.44 | -0.41 | 0.49 | 0.81 |
| 10 | 40600102 | 1.32E-05 | 3.74E-02 | 0.47 | -0.64 | -0.15 | 0.08 |
| 6 | 34157607 | 1.33E-05 | 3.76E-02 | 0.99 | 0.51 | -0.48 | 0.05 |
| 11 | 41189580 | 1.33E-05 | 3.76E-02 | -0.13 | 0.49 | -0.54 | 0.34 |
| 6 | 42342306 | 1.33E-05 | 3.76E-02 | 0.20 | -0.82 | -0.39 | -0.64 |
| 6 | 42342342 | 1.33E-05 | 3.76E-02 | 0.20 | -0.82 | -0.39 | -0.64 |
| 6 | 15413435 | 1.34E-05 | 3.76E-02 | 0.46 | -0.31 | -0.31 | 0.38 |
| 6 | 15413471 | 1.34E-05 | 3.76E-02 | 0.46 | -0.31 | -0.31 | 0.38 |
| 5 | 25142153 | 1.34E-05 | 3.76E-02 | 0.41 | -0.51 | -0.09 | 0.81 |
| 1 | 13328658 | 1.34E-05 | 3.77E-02 | 0.60 | -0.48 | 0.13 | 0.30 |
| 5 | 25896780 | 1.35E-05 | 3.78E-02 | -0.71 | -0.43 | 0.06 | -0.08 |
| 2 | 26070955 | 1.35E-05 | 3.78E-02 | 0.30 | -0.26 | -0.44 | -0.51 |
| 10 | 36120762 | 1.36E-05 | 3.80E-02 | 1.10 | -0.83 | -0.43 | 0.49 |
| 8 | 16392608 | 1.36E-05 | 3.81E-02 | -0.43 | -0.14 | -0.08 | 0.49 |

| | | | | | | | |
|---|---|---|---|---|---|---|---|
| 6 | 15267134 | 1.36E-05 | 3.81E-02 | 0.34 | -0.65 | -0.54 | 0.17 |
| 8 | 44078832 | 1.36E-05 | 3.81E-02 | 0.67 | -0.12 | 0.41 | -0.27 |
| 10 | 1892004 | 1.37E-05 | 3.83E-02 | 0.38 | -0.29 | 0.10 | 0.41 |
| 6 | 15264509 | 1.37E-05 | 3.83E-02 | -0.32 | -1.03 | -0.74 | 0.30 |
| 6 | 15264532 | 1.37E-05 | 3.83E-02 | -0.32 | -1.03 | -0.74 | 0.30 |
| 6 | 15264547 | 1.37E-05 | 3.83E-02 | -0.32 | -1.03 | -0.74 | 0.30 |
| 12 | 9880574 | 1.38E-05 | 3.84E-02 | -0.77 | 0.61 | 0.97 | 0.71 |
| 4 | 19517626 | 1.38E-05 | 3.85E-02 | 1.94 | 0.09 | -0.67 | -0.36 |
| 9 | 30504840 | 1.39E-05 | 3.85E-02 | 0.56 | -0.92 | -0.07 | -0.19 |
| 9 | 30504847 | 1.39E-05 | 3.85E-02 | 0.56 | -0.92 | -0.07 | -0.19 |
| 9 | 30505172 | 1.39E-05 | 3.85E-02 | 0.56 | -0.92 | -0.07 | -0.19 |
| 9 | 28031743 | 1.39E-05 | 3.85E-02 | 1.34 | -0.57 | -0.52 | 0.07 |
| 2 | 24120130 | 1.39E-05 | 3.85E-02 | 0.64 | 0.06 | -0.50 | -0.10 |
| 6 | 6366700 | 1.39E-05 | 3.85E-02 | 0.61 | -0.51 | -0.35 | -0.10 |
| 5 | 24759597 | 1.39E-05 | 3.85E-02 | -0.13 | 0.38 | 0.00 | 0.91 |
| 5 | 38450748 | 1.39E-05 | 3.85E-02 | 0.96 | -0.29 | 0.34 | -0.30 |
| 5 | 38450783 | 1.39E-05 | 3.85E-02 | 0.96 | -0.29 | 0.34 | -0.30 |
| 8 | 43921390 | 1.39E-05 | 3.85E-02 | 0.35 | -0.28 | 0.49 | -0.19 |
| 10 | 20432858 | 1.40E-05 | 3.86E-02 | 0.68 | -0.70 | -0.17 | 0.17 |
| 3 | 8026671 | 1.40E-05 | 3.86E-02 | 1.34 | -0.40 | -0.40 | -0.25 |
| 6 | 15424346 | 1.40E-05 | 3.86E-02 | 0.50 | -0.30 | -0.14 | 0.42 |
| 5 | 6353438 | 1.40E-05 | 3.86E-02 | 0.32 | -0.46 | 0.12 | 0.36 |
| 4 | 73010244 | 1.40E-05 | 3.86E-02 | 0.54 | -0.22 | 0.25 | 0.45 |
| 7 | 19139334 | 1.41E-05 | 3.87E-02 | 1.04 | -0.54 | -0.24 | -0.16 |
| 6 | 16218182 | 1.41E-05 | 3.87E-02 | -0.64 | -0.09 | -0.18 | 0.41 |
| 1 | 13322283 | 1.41E-05 | 3.88E-02 | 0.54 | -0.55 | 0.02 | 0.27 |
| 6 | 53054251 | 1.42E-05 | 3.89E-02 | 1.53 | -0.17 | -0.23 | 0.10 |
| 6 | 53054324 | 1.42E-05 | 3.89E-02 | 1.53 | -0.17 | -0.23 | 0.10 |
| 6 | 53054451 | 1.42E-05 | 3.89E-02 | 1.53 | -0.17 | -0.23 | 0.10 |
| 5 | 25760709 | 1.42E-05 | 3.89E-02 | -0.43 | -0.68 | -0.15 | 0.25 |
| 5 | 25760710 | 1.42E-05 | 3.89E-02 | -0.43 | -0.68 | -0.15 | 0.25 |
| 1 | 23002475 | 1.42E-05 | 3.89E-02 | 0.46 | 0.60 | -0.23 | -0.10 |
| 8 | 21507924 | 1.43E-05 | 3.90E-02 | 1.30 | -0.33 | 0.10 | 0.19 |
| 5 | 38455012 | 1.43E-05 | 3.90E-02 | 0.81 | -0.37 | 0.14 | -0.35 |
| 1 | 23222290 | 1.43E-05 | 3.90E-02 | 0.76 | -0.10 | -0.31 | 0.25 |
| 1 | 8447892 | 1.43E-05 | 3.90E-02 | 1.32 | -0.28 | -0.75 | 0.08 |
| 6 | 15266132 | 1.44E-05 | 3.91E-02 | 0.61 | -0.43 | -0.29 | 0.24 |
| 1 | 26952719 | 1.44E-05 | 3.91E-02 | -0.29 | -0.07 | -0.67 | -0.02 |
| 4 | 66200575 | 1.44E-05 | 3.91E-02 | -0.34 | 0.36 | 0.53 | -0.60 |
| 6 | 42326881 | 1.45E-05 | 3.91E-02 | 0.31 | -0.67 | -0.33 | -0.64 |
| 6 | 42326937 | 1.45E-05 | 3.91E-02 | 0.31 | -0.67 | -0.33 | -0.64 |
| 6 | 9919655 | 1.45E-05 | 3.91E-02 | 0.51 | -0.58 | -0.15 | 0.14 |
| 6 | 9919665 | 1.45E-05 | 3.91E-02 | 0.51 | -0.58 | -0.15 | 0.14 |
| 10 | 16153430 | 1.45E-05 | 3.91E-02 | 0.64 | -0.10 | 0.40 | 0.41 |
| 12 | 4502594 | 1.45E-05 | 3.91E-02 | 0.12 | -0.34 | -0.67 | 0.07 |
| 12 | 4503034 | 1.45E-05 | 3.91E-02 | 0.12 | -0.34 | -0.67 | 0.07 |
| 5 | 78934499 | 1.45E-05 | 3.91E-02 | -1.54 | 0.90 | 0.65 | 0.21 |
| 4 | 33550846 | 1.45E-05 | 3.91E-02 | 0.84 | 0.53 | 0.83 | 0.51 |
| 4 | 33551725 | 1.45E-05 | 3.91E-02 | 0.84 | 0.53 | 0.83 | 0.51 |
| 4 | 33551768 | 1.45E-05 | 3.91E-02 | 0.84 | 0.53 | 0.83 | 0.51 |
| 12 | 15357833 | 1.46E-05 | 3.92E-02 | -0.67 | -0.36 | 0.42 | 0.13 |
| 6 | 16378344 | 1.46E-05 | 3.94E-02 | 0.81 | -0.43 | -0.04 | -0.19 |

| | | | | | | | |
|---|---|---|---|---|---|---|---|
| 6 | 34151442 | 1.46E-05 | 3.94E-02 | 1.04 | 0.49 | -0.43 | 0.04 |
| 12 | 4500626 | 1.47E-05 | 3.94E-02 | 0.27 | -0.33 | -0.68 | 0.06 |
| 5 | 38460797 | 1.47E-05 | 3.94E-02 | 0.67 | -0.49 | -0.08 | -0.27 |
| 5 | 38460798 | 1.47E-05 | 3.94E-02 | 0.67 | -0.49 | -0.08 | -0.27 |
| 1 | 13352402 | 1.48E-05 | 3.95E-02 | 0.58 | -0.53 | 0.04 | 0.27 |
| 2 | 28900890 | 1.48E-05 | 3.95E-02 | -0.17 | -0.65 | -0.11 | 0.91 |
| 10 | 33060464 | 1.48E-05 | 3.95E-02 | 1.34 | -0.60 | -0.11 | 0.21 |
| 10 | 33060852 | 1.48E-05 | 3.95E-02 | 1.34 | -0.60 | -0.11 | 0.21 |
| 10 | 33062219 | 1.48E-05 | 3.95E-02 | 1.34 | -0.60 | -0.11 | 0.21 |
| 6 | 16370474 | 1.48E-05 | 3.96E-02 | 0.53 | -0.56 | 0.04 | -0.16 |
| 6 | 2958376 | 1.48E-05 | 3.96E-02 | 0.66 | -0.34 | -0.62 | -0.11 |
| 5 | 25759027 | 1.49E-05 | 3.97E-02 | -0.49 | -0.68 | -0.15 | 0.26 |
| 2 | 10148008 | 1.50E-05 | 3.98E-02 | -1.12 | 0.26 | 0.17 | 0.54 |
| 4 | 22836958 | 1.50E-05 | 3.98E-02 | -0.78 | -0.12 | -0.74 | -0.93 |
| 10 | 8974241 | 1.50E-05 | 3.98E-02 | -1.09 | 0.41 | -0.61 | -0.26 |
| 8 | 27224660 | 1.50E-05 | 3.98E-02 | -0.58 | 0.53 | -0.30 | 0.23 |
| 10 | 30713161 | 1.51E-05 | 3.98E-02 | 0.07 | 0.10 | 0.12 | -0.55 |
| 8 | 47783572 | 1.51E-05 | 3.98E-02 | 0.29 | -0.28 | 0.57 | -0.20 |
| 4 | 57631305 | 1.51E-05 | 3.98E-02 | -0.12 | 0.42 | -0.85 | -0.51 |
| 6 | 15299369 | 1.51E-05 | 3.98E-02 | 0.56 | -0.46 | -0.33 | 0.23 |
| 4 | 53024444 | 1.52E-05 | 3.98E-02 | -0.29 | -0.93 | -0.68 | 0.26 |
| 5 | 38437062 | 1.53E-05 | 3.98E-02 | 0.79 | -0.38 | 0.13 | -0.36 |
| 5 | 38437273 | 1.53E-05 | 3.98E-02 | 0.79 | -0.38 | 0.13 | -0.36 |
| 5 | 38437532 | 1.53E-05 | 3.98E-02 | 0.79 | -0.38 | 0.13 | -0.36 |
| 5 | 38439576 | 1.53E-05 | 3.98E-02 | 0.79 | -0.38 | 0.13 | -0.36 |
| 5 | 38440513 | 1.53E-05 | 3.98E-02 | 0.79 | -0.38 | 0.13 | -0.36 |
| 5 | 38441003 | 1.53E-05 | 3.98E-02 | 0.79 | -0.38 | 0.13 | -0.36 |
| 5 | 38441152 | 1.53E-05 | 3.98E-02 | 0.79 | -0.38 | 0.13 | -0.36 |
| 5 | 38442065 | 1.53E-05 | 3.98E-02 | 0.79 | -0.38 | 0.13 | -0.36 |
| 5 | 38442289 | 1.53E-05 | 3.98E-02 | 0.79 | -0.38 | 0.13 | -0.36 |
| 5 | 38442369 | 1.53E-05 | 3.98E-02 | 0.79 | -0.38 | 0.13 | -0.36 |
| 5 | 38445687 | 1.53E-05 | 3.98E-02 | 0.79 | -0.38 | 0.13 | -0.36 |
| 5 | 38446584 | 1.53E-05 | 3.98E-02 | 0.79 | -0.38 | 0.13 | -0.36 |
| 5 | 38447524 | 1.53E-05 | 3.98E-02 | 0.79 | -0.38 | 0.13 | -0.36 |
| 5 | 38447551 | 1.53E-05 | 3.98E-02 | 0.79 | -0.38 | 0.13 | -0.36 |
| 5 | 38448369 | 1.53E-05 | 3.98E-02 | 0.79 | -0.38 | 0.13 | -0.36 |
| 5 | 38448391 | 1.53E-05 | 3.98E-02 | 0.79 | -0.38 | 0.13 | -0.36 |
| 5 | 38449157 | 1.53E-05 | 3.98E-02 | 0.79 | -0.38 | 0.13 | -0.36 |
| 5 | 38451264 | 1.53E-05 | 3.98E-02 | 0.79 | -0.38 | 0.13 | -0.36 |
| 5 | 38454395 | 1.53E-05 | 3.98E-02 | 0.79 | -0.38 | 0.13 | -0.36 |
| 5 | 38454459 | 1.53E-05 | 3.98E-02 | 0.79 | -0.38 | 0.13 | -0.36 |
| 5 | 38456536 | 1.53E-05 | 3.98E-02 | 0.79 | -0.38 | 0.13 | -0.36 |
| 5 | 38457020 | 1.53E-05 | 3.98E-02 | 0.79 | -0.38 | 0.13 | -0.36 |
| 5 | 38457245 | 1.53E-05 | 3.98E-02 | 0.79 | -0.38 | 0.13 | -0.36 |
| 10 | 40568001 | 1.53E-05 | 3.98E-02 | 0.50 | -0.59 | -0.31 | 0.01 |
| 6 | 36371724 | 1.53E-05 | 3.98E-02 | 0.76 | -0.34 | 0.02 | -0.48 |
| 1 | 37449432 | 1.53E-05 | 3.98E-02 | 0.00 | -0.55 | -0.32 | -0.34 |
| 12 | 4509941 | 1.53E-05 | 3.98E-02 | 0.15 | -0.34 | -0.66 | 0.08 |
| 2 | 29434098 | 1.54E-05 | 3.98E-02 | -0.22 | -0.33 | -0.28 | 0.78 |
| 9 | 34516370 | 1.54E-05 | 3.98E-02 | 1.71 | -0.69 | -0.47 | -0.53 |
| 4 | 45030841 | 1.54E-05 | 3.98E-02 | -0.02 | 0.71 | -0.97 | -0.72 |
| 6 | 28491337 | 1.55E-05 | 3.98E-02 | 0.03 | 0.78 | -0.57 | 0.55 |

| | | | | | | | |
|---|---|---|---|---|---|---|---|
| 7 | 23560999 | 1.55E-05 | 3.98E-02 | 0.91 | -0.53 | -0.33 | 0.15 |
| 6 | 44800429 | 1.55E-05 | 3.98E-02 | 0.80 | -0.68 | -0.46 | -0.11 |
| 6 | 6373318 | 1.55E-05 | 3.98E-02 | 0.64 | -0.61 | -0.48 | -0.35 |
| 7 | 31020470 | 1.55E-05 | 3.98E-02 | -0.56 | -0.42 | -0.05 | 0.61 |
| 2 | 9514717 | 1.55E-05 | 3.98E-02 | 0.16 | -0.46 | -0.62 | 0.11 |
| 6 | 44511974 | 1.55E-05 | 3.98E-02 | 0.66 | -0.67 | -0.37 | -0.30 |
| 1 | 37405706 | 1.56E-05 | 3.98E-02 | -0.21 | -0.57 | -0.30 | -0.31 |
| 1 | 13320936 | 1.57E-05 | 3.98E-02 | 0.55 | -0.51 | 0.02 | 0.31 |
| 1 | 13321674 | 1.57E-05 | 3.98E-02 | 0.55 | -0.51 | 0.02 | 0.31 |
| 1 | 13326493 | 1.57E-05 | 3.98E-02 | 0.55 | -0.51 | 0.02 | 0.31 |
| 1 | 13327483 | 1.57E-05 | 3.98E-02 | 0.55 | -0.51 | 0.02 | 0.31 |
| 1 | 13328561 | 1.57E-05 | 3.98E-02 | 0.55 | -0.51 | 0.02 | 0.31 |
| 1 | 13329469 | 1.57E-05 | 3.98E-02 | 0.55 | -0.51 | 0.02 | 0.31 |
| 1 | 13332260 | 1.57E-05 | 3.98E-02 | 0.55 | -0.51 | 0.02 | 0.31 |
| 1 | 13334575 | 1.57E-05 | 3.98E-02 | 0.55 | -0.51 | 0.02 | 0.31 |
| 1 | 13335173 | 1.57E-05 | 3.98E-02 | 0.55 | -0.51 | 0.02 | 0.31 |
| 1 | 13335783 | 1.57E-05 | 3.98E-02 | 0.55 | -0.51 | 0.02 | 0.31 |
| 1 | 13338148 | 1.57E-05 | 3.98E-02 | 0.55 | -0.51 | 0.02 | 0.31 |
| 1 | 13339784 | 1.57E-05 | 3.98E-02 | 0.55 | -0.51 | 0.02 | 0.31 |
| 1 | 13340030 | 1.57E-05 | 3.98E-02 | 0.55 | -0.51 | 0.02 | 0.31 |
| 1 | 13340797 | 1.57E-05 | 3.98E-02 | 0.55 | -0.51 | 0.02 | 0.31 |
| 1 | 13340939 | 1.57E-05 | 3.98E-02 | 0.55 | -0.51 | 0.02 | 0.31 |
| 1 | 13342131 | 1.57E-05 | 3.98E-02 | 0.55 | -0.51 | 0.02 | 0.31 |
| 1 | 13345260 | 1.57E-05 | 3.98E-02 | 0.55 | -0.51 | 0.02 | 0.31 |
| 1 | 13346026 | 1.57E-05 | 3.98E-02 | 0.55 | -0.51 | 0.02 | 0.31 |
| 1 | 13347976 | 1.57E-05 | 3.98E-02 | 0.55 | -0.51 | 0.02 | 0.31 |
| 1 | 15419988 | 1.57E-05 | 3.98E-02 | 1.02 | -0.28 | -0.29 | 0.15 |
| 12 | 18582249 | 1.57E-05 | 3.99E-02 | 0.85 | -0.47 | 0.04 | -0.08 |
| 7 | 35965565 | 1.57E-05 | 3.99E-02 | 1.15 | -0.69 | -0.53 | -0.29 |
| 12 | 21573005 | 1.58E-05 | 4.00E-02 | 1.84 | -1.52 | 0.17 | 0.63 |
| 11 | 33954398 | 1.58E-05 | 4.00E-02 | 1.12 | 0.11 | 0.43 | 0.22 |
| 10 | 43965618 | 1.58E-05 | 4.00E-02 | 2.62 | 0.06 | -1.31 | -0.36 |
| 6 | 36342299 | 1.58E-05 | 4.00E-02 | 1.49 | -0.30 | -0.18 | -0.43 |
| 5 | 65813727 | 1.58E-05 | 4.00E-02 | 0.23 | -0.59 | -0.51 | -0.28 |
| 10 | 16137248 | 1.58E-05 | 4.00E-02 | 0.90 | -0.21 | 0.28 | 0.58 |
| 2 | 30230874 | 1.59E-05 | 4.01E-02 | 1.69 | -0.59 | -0.28 | 0.56 |
| 4 | 19389027 | 1.59E-05 | 4.03E-02 | 0.82 | 0.53 | -0.35 | -0.12 |
| 1 | 37445015 | 1.60E-05 | 4.03E-02 | 0.00 | -0.51 | -0.19 | -0.38 |
| 1 | 13324664 | 1.60E-05 | 4.03E-02 | 0.63 | -0.40 | 0.21 | 0.33 |
| 4 | 19377729 | 1.60E-05 | 4.03E-02 | 1.23 | 0.80 | 0.12 | 0.23 |
| 4 | 35688459 | 1.60E-05 | 4.04E-02 | 0.36 | -0.52 | -0.95 | -0.36 |
| 7 | 29819764 | 1.61E-05 | 4.05E-02 | 0.72 | -0.21 | 0.24 | 0.45 |
| 4 | 33298596 | 1.61E-05 | 4.05E-02 | 1.60 | 0.00 | 0.89 | 0.00 |
| 1 | 16235294 | 1.61E-05 | 4.05E-02 | -1.00 | 0.01 | 0.34 | 0.92 |
| 4 | 41455049 | 1.62E-05 | 4.06E-02 | -0.42 | 0.29 | -0.82 | -0.95 |
| 5 | 74753702 | 1.62E-05 | 4.06E-02 | -0.39 | 0.63 | 0.43 | 1.04 |
| 5 | 74756006 | 1.62E-05 | 4.06E-02 | -0.39 | 0.63 | 0.43 | 1.04 |
| 9 | 27941845 | 1.62E-05 | 4.06E-02 | 1.41 | -0.44 | -0.28 | 0.19 |
| 9 | 41007084 | 1.62E-05 | 4.06E-02 | 1.74 | -0.09 | -0.99 | -0.04 |
| 9 | 41007174 | 1.62E-05 | 4.06E-02 | 1.74 | -0.09 | -0.99 | -0.04 |
| 9 | 41028835 | 1.62E-05 | 4.06E-02 | 1.74 | -0.09 | -0.99 | -0.04 |
| 9 | 41029522 | 1.62E-05 | 4.06E-02 | 1.74 | -0.09 | -0.99 | -0.04 |

| | | | | | | | |
|---|---|---|---|---|---|---|---|
| 1 | 33334772 | 1.62E-05 | 4.06E-02 | 0.97 | -1.14 | -0.41 | 0.38 |
| 6 | 25881981 | 1.63E-05 | 4.06E-02 | 0.50 | -0.28 | -0.79 | -0.79 |
| 12 | 5346830 | 1.64E-05 | 4.06E-02 | 0.03 | -0.34 | -0.64 | 0.22 |
| 10 | 29834636 | 1.64E-05 | 4.06E-02 | -0.05 | -0.11 | -0.55 | -0.41 |
| 4 | 19503731 | 1.64E-05 | 4.06E-02 | 1.92 | 0.13 | -0.71 | -0.32 |
| 8 | 36501420 | 1.64E-05 | 4.06E-02 | 0.60 | -0.20 | -0.24 | 0.28 |
| 8 | 36504456 | 1.64E-05 | 4.06E-02 | 0.60 | -0.20 | -0.24 | 0.28 |
| 8 | 36504538 | 1.64E-05 | 4.06E-02 | 0.60 | -0.20 | -0.24 | 0.28 |
| 8 | 36505700 | 1.64E-05 | 4.06E-02 | 0.60 | -0.20 | -0.24 | 0.28 |
| 8 | 36509643 | 1.64E-05 | 4.06E-02 | 0.60 | -0.20 | -0.24 | 0.28 |
| 8 | 36511896 | 1.64E-05 | 4.06E-02 | 0.60 | -0.20 | -0.24 | 0.28 |
| 8 | 36523439 | 1.64E-05 | 4.06E-02 | 0.60 | -0.20 | -0.24 | 0.28 |
| 8 | 36524129 | 1.64E-05 | 4.06E-02 | 0.60 | -0.20 | -0.24 | 0.28 |
| 8 | 36524593 | 1.64E-05 | 4.06E-02 | 0.60 | -0.20 | -0.24 | 0.28 |
| 1 | 14867061 | 1.64E-05 | 4.06E-02 | 2.14 | -0.58 | 0.45 | 0.55 |
| 7 | 40529278 | 1.64E-05 | 4.06E-02 | 0.41 | 0.14 | -0.34 | 0.62 |
| 4 | 81139782 | 1.64E-05 | 4.06E-02 | 0.05 | 1.09 | -0.14 | 0.18 |
| 2 | 10864294 | 1.64E-05 | 4.06E-02 | -0.50 | -0.48 | 0.10 | -0.22 |
| 8 | 29175166 | 1.65E-05 | 4.06E-02 | 0.41 | -0.53 | 0.13 | -0.38 |
| 8 | 29175181 | 1.65E-05 | 4.06E-02 | 0.41 | -0.53 | 0.13 | -0.38 |
| 4 | 34668747 | 1.65E-05 | 4.07E-02 | 0.24 | -0.82 | 0.34 | 0.21 |
| 6 | 6369266 | 1.65E-05 | 4.07E-02 | 0.53 | -0.63 | -0.37 | 0.00 |
| 9 | 20759343 | 1.66E-05 | 4.09E-02 | -0.84 | 0.46 | 0.63 | -0.17 |
| 6 | 16899048 | 1.66E-05 | 4.09E-02 | 0.09 | -0.18 | 0.64 | -0.13 |
| 10 | 24307517 | 1.67E-05 | 4.09E-02 | -0.72 | -0.39 | 0.21 | 0.22 |
| 10 | 23658192 | 1.67E-05 | 4.09E-02 | 0.67 | -0.41 | -0.94 | -0.14 |
| 4 | 8020089 | 1.68E-05 | 4.11E-02 | 1.26 | 1.52 | -0.99 | -0.46 |
| 10 | 24291873 | 1.69E-05 | 4.13E-02 | 0.80 | -0.52 | 0.08 | 0.20 |
| 7 | 30940200 | 1.69E-05 | 4.13E-02 | -0.19 | -0.48 | 0.01 | 0.42 |
| 8 | 27221733 | 1.69E-05 | 4.13E-02 | 1.47 | -0.57 | -0.35 | -0.45 |
| 4 | 46815711 | 1.69E-05 | 4.13E-02 | -0.07 | -0.71 | -0.62 | 0.27 |
| 7 | 35964250 | 1.69E-05 | 4.14E-02 | 1.21 | -0.67 | -0.55 | -0.32 |
| 1 | 37444889 | 1.69E-05 | 4.14E-02 | -0.12 | -0.54 | -0.17 | -0.34 |
| 6 | 10414355 | 1.70E-05 | 4.15E-02 | 0.61 | -0.15 | -0.59 | -0.85 |
| 2 | 21687757 | 1.71E-05 | 4.17E-02 | 0.70 | -0.52 | -0.28 | -0.23 |
| 7 | 10654060 | 1.71E-05 | 4.17E-02 | 1.14 | 0.53 | -0.36 | 0.16 |
| 5 | 25777392 | 1.71E-05 | 4.17E-02 | -0.67 | -0.45 | 0.08 | -0.05 |
| 5 | 25801617 | 1.71E-05 | 4.17E-02 | -0.67 | -0.45 | 0.08 | -0.05 |
| 7 | 35969106 | 1.71E-05 | 4.17E-02 | 1.26 | -0.77 | -0.35 | -0.18 |
| 5 | 60930483 | 1.71E-05 | 4.17E-02 | -1.35 | 0.29 | 0.25 | 0.73 |
| 1 | 42299529 | 1.72E-05 | 4.19E-02 | 0.45 | -0.61 | -0.49 | -0.02 |
| 6 | 45870965 | 1.73E-05 | 4.19E-02 | 0.42 | 0.10 | -0.51 | 0.24 |
| 10 | 5990132 | 1.73E-05 | 4.19E-02 | 0.48 | -0.51 | -0.34 | 0.00 |
| 9 | 35346405 | 1.73E-05 | 4.21E-02 | -0.36 | -0.18 | 0.48 | 0.86 |
| 12 | 13947492 | 1.74E-05 | 4.21E-02 | 1.15 | -0.10 | -0.10 | 0.24 |
| 8 | 44912268 | 1.74E-05 | 4.21E-02 | 0.11 | 0.48 | -0.59 | -0.04 |
| 8 | 383577 | 1.74E-05 | 4.21E-02 | 0.56 | -0.41 | -0.49 | 0.25 |
| 5 | 36746352 | 1.75E-05 | 4.22E-02 | 1.62 | 0.91 | -1.32 | 0.32 |
| 5 | 36747210 | 1.75E-05 | 4.22E-02 | 1.62 | 0.91 | -1.32 | 0.32 |
| 5 | 36747992 | 1.75E-05 | 4.22E-02 | 1.62 | 0.91 | -1.32 | 0.32 |
| 5 | 36748679 | 1.75E-05 | 4.22E-02 | 1.62 | 0.91 | -1.32 | 0.32 |
| 5 | 36749400 | 1.75E-05 | 4.22E-02 | 1.62 | 0.91 | -1.32 | 0.32 |

| | | | | | | | |
|---|---|---|---|---|---|---|---|
| 12 | 16259301 | 1.75E-05 | 4.22E-02 | -0.66 | -0.31 | 0.31 | 0.21 |
| 12 | 16259390 | 1.75E-05 | 4.22E-02 | -0.66 | -0.31 | 0.31 | 0.21 |
| 1 | 48735286 | 1.76E-05 | 4.22E-02 | -1.33 | 1.47 | 0.60 | 0.74 |
| 11 | 33834615 | 1.76E-05 | 4.22E-02 | 1.09 | 0.23 | 0.20 | 0.25 |
| 2 | 17417790 | 1.76E-05 | 4.22E-02 | -0.15 | -0.77 | -0.26 | 0.19 |
| 7 | 16627935 | 1.76E-05 | 4.22E-02 | -0.36 | -0.57 | 0.10 | 0.25 |
| 7 | 16630870 | 1.76E-05 | 4.22E-02 | -0.36 | -0.57 | 0.10 | 0.25 |
| 7 | 16631140 | 1.76E-05 | 4.22E-02 | -0.36 | -0.57 | 0.10 | 0.25 |
| 7 | 16631213 | 1.76E-05 | 4.22E-02 | -0.36 | -0.57 | 0.10 | 0.25 |
| 7 | 16640865 | 1.76E-05 | 4.22E-02 | -0.36 | -0.57 | 0.10 | 0.25 |
| 7 | 16644525 | 1.76E-05 | 4.22E-02 | -0.36 | -0.57 | 0.10 | 0.25 |
| 7 | 16646177 | 1.76E-05 | 4.22E-02 | -0.36 | -0.57 | 0.10 | 0.25 |
| 11 | 54744353 | 1.77E-05 | 4.23E-02 | -1.07 | 0.58 | 0.02 | 0.20 |
| 9 | 17896003 | 1.77E-05 | 4.24E-02 | -0.25 | -0.89 | -0.62 | 0.30 |
| 6 | 15085166 | 1.78E-05 | 4.24E-02 | -0.30 | -0.96 | -0.70 | 0.25 |
| 4 | 32834749 | 1.78E-05 | 4.24E-02 | -0.81 | -0.33 | -0.11 | 0.15 |
| 7 | 35962893 | 1.78E-05 | 4.25E-02 | 1.12 | -0.69 | -0.58 | -0.31 |
| 6 | 9906948 | 1.78E-05 | 4.25E-02 | 0.44 | -0.60 | -0.19 | 0.17 |
| 11 | 57506960 | 1.79E-05 | 4.25E-02 | 0.98 | -0.62 | -0.27 | -0.12 |
| 6 | 16291306 | 1.79E-05 | 4.25E-02 | 0.64 | -0.52 | -0.13 | -0.16 |
| 6 | 6366887 | 1.80E-05 | 4.26E-02 | 0.58 | -0.50 | -0.38 | -0.11 |
| 6 | 6366905 | 1.80E-05 | 4.26E-02 | 0.58 | -0.50 | -0.38 | -0.11 |
| 6 | 6366906 | 1.80E-05 | 4.26E-02 | 0.58 | -0.50 | -0.38 | -0.11 |
| 9 | 27176321 | 1.80E-05 | 4.26E-02 | 1.04 | 0.01 | 0.22 | 0.22 |
| 6 | 30890911 | 1.80E-05 | 4.26E-02 | -0.54 | 0.21 | 0.43 | 0.68 |
| 4 | 54034128 | 1.80E-05 | 4.26E-02 | 0.95 | -0.37 | 0.27 | -0.31 |
| 5 | 25779957 | 1.80E-05 | 4.26E-02 | -0.69 | -0.39 | 0.25 | -0.06 |
| 11 | 52565815 | 1.80E-05 | 4.27E-02 | 0.96 | -0.22 | -0.29 | -0.18 |
| 10 | 30712910 | 1.81E-05 | 4.28E-02 | 0.11 | 0.13 | 0.11 | -0.55 |
| 8 | 44077056 | 1.81E-05 | 4.29E-02 | 0.75 | -0.06 | 0.35 | -0.24 |
| 12 | 4502373 | 1.82E-05 | 4.29E-02 | 0.13 | -0.32 | -0.66 | 0.11 |
| 12 | 4507891 | 1.82E-05 | 4.29E-02 | 0.13 | -0.32 | -0.66 | 0.11 |
| 10 | 6554706 | 1.82E-05 | 4.30E-02 | 0.60 | -0.36 | -0.33 | 0.23 |
| 1 | 40260106 | 1.83E-05 | 4.31E-02 | 0.09 | -0.08 | 0.31 | 0.68 |
| 2 | 8435280 | 1.83E-05 | 4.31E-02 | 0.78 | -0.68 | -0.24 | -0.18 |
| 6 | 44533660 | 1.83E-05 | 4.31E-02 | 0.64 | -0.56 | -0.49 | -0.23 |
| 7 | 16616614 | 1.84E-05 | 4.32E-02 | -0.38 | -0.57 | 0.12 | 0.24 |
| 8 | 47697278 | 1.84E-05 | 4.32E-02 | 0.29 | -0.21 | 0.60 | -0.16 |
| 8 | 47697343 | 1.84E-05 | 4.32E-02 | 0.29 | -0.21 | 0.60 | -0.16 |
| 9 | 29166864 | 1.84E-05 | 4.32E-02 | 0.57 | -0.44 | -0.18 | 0.23 |
| 7 | 16635730 | 1.85E-05 | 4.34E-02 | -0.22 | -0.62 | 0.12 | 0.23 |
| 8 | 20729094 | 1.85E-05 | 4.34E-02 | -0.61 | 0.05 | 0.43 | 0.78 |
| 6 | 44521337 | 1.85E-05 | 4.34E-02 | 0.62 | -0.68 | -0.42 | -0.18 |
| 6 | 15401820 | 1.85E-05 | 4.34E-02 | 0.41 | -0.33 | -0.25 | 0.42 |
| 8 | 22403994 | 1.86E-05 | 4.35E-02 | -0.56 | 0.24 | -0.05 | 0.59 |
| 6 | 15327937 | 1.86E-05 | 4.35E-02 | -0.33 | -1.08 | -0.78 | 0.30 |
| 9 | 30508001 | 1.86E-05 | 4.35E-02 | 0.54 | -0.85 | -0.16 | -0.26 |
| 1 | 24679668 | 1.86E-05 | 4.35E-02 | -0.01 | 0.13 | 0.09 | -0.65 |
| 8 | 49044296 | 1.86E-05 | 4.35E-02 | 0.31 | -0.12 | -0.65 | -0.16 |
| 6 | 44520090 | 1.86E-05 | 4.35E-02 | 0.36 | -0.50 | -0.57 | -0.24 |
| 5 | 52793525 | 1.86E-05 | 4.35E-02 | 1.49 | 0.30 | -0.83 | -0.60 |
| 4 | 81162805 | 1.87E-05 | 4.36E-02 | 0.21 | 0.88 | -0.37 | -0.16 |

| | | | | | | | |
|---|---|---|---|---|---|---|---|
| 7 | 39742474 | 1.87E-05 | 4.36E-02 | 0.22 | -0.77 | -0.14 | 0.59 |
| 5 | 59091005 | 1.87E-05 | 4.36E-02 | 0.61 | -0.11 | -0.45 | 0.11 |
| 12 | 4500439 | 1.89E-05 | 4.39E-02 | 0.17 | -0.34 | -0.66 | 0.07 |
| 2 | 17213524 | 1.89E-05 | 4.39E-02 | -0.50 | -0.22 | -0.20 | -0.53 |
| 6 | 2088544 | 1.90E-05 | 4.41E-02 | 0.02 | -0.61 | -0.54 | -0.62 |
| 1 | 27290152 | 1.90E-05 | 4.41E-02 | -0.49 | -0.76 | -0.02 | 0.56 |
| 5 | 25752746 | 1.90E-05 | 4.41E-02 | -0.58 | -0.52 | 0.05 | 0.16 |
| 7 | 40549368 | 1.90E-05 | 4.41E-02 | 0.01 | -0.21 | -0.45 | 0.50 |
| 2 | 29015156 | 1.90E-05 | 4.41E-02 | -0.36 | -0.50 | -0.05 | 0.93 |
| 4 | 11812226 | 1.90E-05 | 4.41E-02 | 1.03 | 0.44 | -0.08 | 0.14 |
| 6 | 45055278 | 1.91E-05 | 4.42E-02 | 0.23 | -0.73 | -0.58 | -0.46 |
| 5 | 25812022 | 1.91E-05 | 4.42E-02 | -0.68 | -0.50 | 0.05 | -0.08 |
| 4 | 49486627 | 1.92E-05 | 4.44E-02 | 1.23 | 0.45 | -0.52 | -0.52 |
| 2 | 17217593 | 1.92E-05 | 4.44E-02 | -0.10 | -0.25 | -0.32 | -0.53 |
| 6 | 6374361 | 1.92E-05 | 4.44E-02 | 0.67 | -0.56 | -0.20 | -0.01 |
| 8 | 20933468 | 1.93E-05 | 4.45E-02 | 0.40 | -0.08 | -0.50 | 0.21 |
| 6 | 43820296 | 1.93E-05 | 4.45E-02 | 0.64 | -0.27 | 0.03 | 0.41 |
| 6 | 45186505 | 1.93E-05 | 4.45E-02 | -0.52 | 1.20 | 0.72 | 0.47 |
| 7 | 16632883 | 1.94E-05 | 4.47E-02 | -0.24 | -0.55 | 0.40 | 0.11 |
| 8 | 27385051 | 1.95E-05 | 4.48E-02 | 0.39 | 0.57 | -0.59 | -0.24 |
| 6 | 34163118 | 1.95E-05 | 4.48E-02 | 0.99 | 0.51 | -0.45 | 0.01 |
| 7 | 37017891 | 1.95E-05 | 4.48E-02 | -0.68 | 0.33 | -0.99 | -1.19 |
| 7 | 31989945 | 1.95E-05 | 4.48E-02 | -0.01 | -0.51 | -0.14 | -0.44 |
| 7 | 31990008 | 1.95E-05 | 4.48E-02 | -0.01 | -0.51 | -0.14 | -0.44 |
| 12 | 4497791 | 1.95E-05 | 4.49E-02 | 0.09 | -0.32 | -0.65 | 0.10 |
| 9 | 30507479 | 1.96E-05 | 4.49E-02 | 0.54 | -0.84 | -0.17 | -0.26 |
| 9 | 30508127 | 1.96E-05 | 4.49E-02 | 0.54 | -0.84 | -0.17 | -0.26 |
| 1 | 13345967 | 1.96E-05 | 4.49E-02 | 0.55 | -0.53 | -0.01 | 0.29 |
| 8 | 51922373 | 1.97E-05 | 4.50E-02 | -0.54 | -0.38 | 0.51 | -0.12 |
| 7 | 15577236 | 1.97E-05 | 4.50E-02 | 0.36 | 0.17 | -0.37 | 0.51 |
| 6 | 13406034 | 1.98E-05 | 4.52E-02 | 0.07 | -0.33 | -0.67 | -1.03 |
| 5 | 66698844 | 1.98E-05 | 4.52E-02 | -0.58 | -0.31 | -0.66 | 1.27 |
| 6 | 9996977 | 1.98E-05 | 4.52E-02 | 0.50 | -0.51 | -0.31 | -0.01 |
| 6 | 24658910 | 1.98E-05 | 4.52E-02 | 1.20 | -0.47 | -0.42 | -0.54 |
| 1 | 40303529 | 1.98E-05 | 4.52E-02 | 0.03 | -0.17 | -0.03 | 0.64 |
| 1 | 40339410 | 1.98E-05 | 4.52E-02 | 0.03 | -0.17 | -0.03 | 0.64 |
| 7 | 30806060 | 1.99E-05 | 4.53E-02 | 0.82 | -0.68 | -0.30 | -0.43 |
| 7 | 30806090 | 1.99E-05 | 4.53E-02 | 0.82 | -0.68 | -0.30 | -0.43 |
| 4 | 69670056 | 1.99E-05 | 4.53E-02 | -1.55 | 0.38 | -1.31 | -0.32 |
| 4 | 11833791 | 2.00E-05 | 4.54E-02 | 0.99 | 0.41 | -0.08 | 0.18 |
| 5 | 38372180 | 2.00E-05 | 4.54E-02 | 0.69 | -0.43 | 0.06 | -0.32 |
| 1 | 18324729 | 2.00E-05 | 4.54E-02 | -1.31 | 1.03 | -1.05 | -0.70 |
| 7 | 19272725 | 2.00E-05 | 4.54E-02 | 1.54 | -0.35 | -0.66 | 0.07 |
| 4 | 51063779 | 2.00E-05 | 4.54E-02 | 1.31 | 0.44 | -0.19 | 0.32 |
| 5 | 48564030 | 2.01E-05 | 4.54E-02 | -0.76 | -0.04 | 0.29 | -0.25 |
| 9 | 36299381 | 2.01E-05 | 4.54E-02 | 0.99 | 0.31 | -0.96 | 0.82 |
| 1 | 7692810 | 2.01E-05 | 4.55E-02 | 0.20 | 0.52 | -0.26 | 0.38 |
| 1 | 7692984 | 2.01E-05 | 4.55E-02 | 0.20 | 0.52 | -0.26 | 0.38 |
| 4 | 54033137 | 2.02E-05 | 4.56E-02 | 0.99 | -0.24 | 0.26 | -0.34 |
| 4 | 54033205 | 2.02E-05 | 4.56E-02 | 0.99 | -0.24 | 0.26 | -0.34 |
| 4 | 54033212 | 2.02E-05 | 4.56E-02 | 0.99 | -0.24 | 0.26 | -0.34 |
| 12 | 16238523 | 2.02E-05 | 4.56E-02 | -0.64 | -0.29 | 0.30 | 0.24 |

| | | | | | | | |
|---|---|---|---|---|---|---|---|
| 12 | 16239139 | 2.02E-05 | 4.56E-02 | -0.64 | -0.29 | 0.30 | 0.24 |
| 12 | 16241629 | 2.02E-05 | 4.56E-02 | -0.64 | -0.29 | 0.30 | 0.24 |
| 6 | 15198707 | 2.02E-05 | 4.56E-02 | -0.26 | -0.93 | -0.62 | 0.30 |
| 8 | 45888388 | 2.03E-05 | 4.58E-02 | 0.68 | -0.17 | -0.34 | -0.38 |
| 6 | 14588868 | 2.04E-05 | 4.58E-02 | 0.01 | 0.39 | 0.49 | -0.20 |
| 5 | 50215260 | 2.04E-05 | 4.59E-02 | 0.94 | 0.21 | 0.61 | -0.07 |
| 5 | 20418425 | 2.04E-05 | 4.59E-02 | -0.23 | -0.58 | 0.21 | -0.49 |
| 5 | 25772512 | 2.04E-05 | 4.59E-02 | -0.71 | -0.40 | 0.10 | -0.03 |
| 1 | 40284606 | 2.05E-05 | 4.59E-02 | -0.24 | -0.01 | 0.29 | 0.69 |
| 6 | 15412001 | 2.05E-05 | 4.59E-02 | 0.39 | -0.37 | -0.29 | 0.39 |
| 1 | 42326594 | 2.05E-05 | 4.59E-02 | 0.57 | -0.65 | -0.43 | 0.06 |
| 6 | 15349094 | 2.05E-05 | 4.59E-02 | 0.47 | -0.51 | -0.31 | 0.28 |
| 6 | 29877849 | 2.05E-05 | 4.59E-02 | 1.09 | -0.18 | 0.05 | 0.17 |
| 6 | 42345774 | 2.06E-05 | 4.59E-02 | 0.51 | -0.75 | -0.24 | -0.55 |
| 6 | 44495267 | 2.06E-05 | 4.59E-02 | 0.95 | -0.52 | -0.34 | -0.10 |
| 8 | 9306959 | 2.06E-05 | 4.59E-02 | 0.24 | -0.70 | 0.41 | 1.11 |
| 4 | 34897794 | 2.06E-05 | 4.59E-02 | 1.11 | -0.24 | -0.41 | -0.29 |
| 4 | 34900458 | 2.06E-05 | 4.59E-02 | 1.11 | -0.24 | -0.41 | -0.29 |
| 4 | 34901056 | 2.06E-05 | 4.59E-02 | 1.11 | -0.24 | -0.41 | -0.29 |
| 6 | 16706791 | 2.06E-05 | 4.59E-02 | 3.05 | 0.36 | -1.50 | 1.17 |
| 1 | 37441226 | 2.06E-05 | 4.59E-02 | -0.20 | -0.55 | -0.24 | -0.30 |
| 9 | 6404349 | 2.06E-05 | 4.59E-02 | 0.09 | 0.30 | 0.57 | 0.23 |
| 12 | 20299820 | 2.06E-05 | 4.59E-02 | 1.18 | -0.44 | -0.17 | -0.46 |
| 4 | 50767588 | 2.06E-05 | 4.59E-02 | 0.02 | -0.51 | -0.57 | 0.21 |
| 6 | 41402410 | 2.07E-05 | 4.59E-02 | 0.60 | -0.74 | -0.35 | 0.18 |
| 7 | 27680173 | 2.07E-05 | 4.60E-02 | 0.61 | -1.13 | -0.13 | -0.53 |
| 6 | 44524170 | 2.07E-05 | 4.60E-02 | 0.58 | -0.58 | -0.52 | -0.20 |
| 8 | 43910576 | 2.08E-05 | 4.60E-02 | 0.56 | -0.27 | 0.42 | -0.19 |
| 12 | 4251475 | 2.08E-05 | 4.60E-02 | 0.20 | 0.13 | 0.56 | 0.50 |
| 12 | 4251476 | 2.08E-05 | 4.60E-02 | 0.20 | 0.13 | 0.56 | 0.50 |
| 7 | 16616719 | 2.08E-05 | 4.60E-02 | -0.58 | -0.42 | 0.34 | 0.19 |
| 10 | 15339025 | 2.08E-05 | 4.60E-02 | 0.75 | -0.05 | 0.51 | 0.37 |
| 12 | 15705941 | 2.08E-05 | 4.61E-02 | -1.04 | 0.61 | 0.27 | 0.27 |
| 10 | 30716222 | 2.09E-05 | 4.61E-02 | 0.11 | 0.04 | -0.01 | -0.56 |
| 6 | 44604573 | 2.09E-05 | 4.62E-02 | 0.91 | -0.05 | 0.45 | 0.09 |
| 10 | 40583863 | 2.10E-05 | 4.62E-02 | 0.54 | -0.53 | -0.30 | 0.02 |
| 4 | 11820875 | 2.10E-05 | 4.63E-02 | 1.05 | 0.40 | -0.05 | 0.19 |
| 8 | 49066418 | 2.10E-05 | 4.63E-02 | 0.46 | -0.28 | -0.58 | -0.01 |
| 7 | 45027476 | 2.10E-05 | 4.63E-02 | 0.37 | -0.56 | -0.93 | -0.61 |
| 2 | 6135697 | 2.10E-05 | 4.63E-02 | -2.24 | 0.42 | 0.46 | 0.76 |
| 6 | 44570957 | 2.10E-05 | 4.63E-02 | 0.33 | -0.85 | -0.51 | -0.40 |
| 6 | 9908689 | 2.11E-05 | 4.63E-02 | 0.38 | -0.63 | -0.29 | 0.15 |
| 10 | 5992419 | 2.11E-05 | 4.63E-02 | 0.56 | -0.41 | -0.36 | -0.01 |
| 6 | 28499181 | 2.11E-05 | 4.64E-02 | 0.06 | 0.63 | -0.47 | 0.46 |
| 5 | 11342188 | 2.11E-05 | 4.64E-02 | -1.47 | 0.14 | -0.04 | 0.10 |
| 5 | 2142722 | 2.12E-05 | 4.65E-02 | -2.29 | -0.18 | -0.74 | 0.64 |
| 5 | 44485831 | 2.12E-05 | 4.65E-02 | -1.08 | 0.21 | 0.03 | 0.59 |
| 11 | 54739805 | 2.12E-05 | 4.65E-02 | -1.07 | 0.51 | 0.04 | 0.19 |
| 8 | 46728481 | 2.12E-05 | 4.65E-02 | 0.00 | 0.56 | -0.58 | -0.14 |
| 4 | 54269030 | 2.13E-05 | 4.65E-02 | -0.09 | -0.10 | -0.87 | -0.25 |
| 5 | 36294962 | 2.13E-05 | 4.65E-02 | 0.68 | -0.58 | 0.13 | 0.25 |
| 5 | 44658601 | 2.13E-05 | 4.65E-02 | 2.22 | 0.16 | -0.87 | 0.42 |

| | | | | | | | |
|---|---|---|---|---|---|---|---|
| 1 | 5195994 | 2.13E-05 | 4.65E-02 | -0.55 | -0.29 | 0.38 | -0.26 |
| 1 | 5197194 | 2.13E-05 | 4.65E-02 | -0.55 | -0.29 | 0.38 | -0.26 |
| 3 | 14371222 | 2.13E-05 | 4.65E-02 | 1.47 | 0.70 | -1.09 | 0.29 |
| 4 | 65059995 | 2.13E-05 | 4.65E-02 | 0.36 | -0.26 | -0.51 | -0.35 |
| 6 | 23660618 | 2.14E-05 | 4.66E-02 | -0.40 | 0.48 | 0.19 | 0.51 |
| 7 | 40644847 | 2.14E-05 | 4.66E-02 | 0.17 | -0.35 | -0.48 | 0.31 |
| 4 | 35903976 | 2.14E-05 | 4.66E-02 | 0.68 | -0.28 | 0.27 | 1.02 |
| 12 | 16238110 | 2.14E-05 | 4.66E-02 | -0.64 | -0.29 | 0.29 | 0.23 |
| 8 | 49295201 | 2.15E-05 | 4.66E-02 | -0.55 | -0.61 | 0.13 | -0.01 |
| 8 | 20936914 | 2.15E-05 | 4.66E-02 | 0.41 | -0.07 | -0.47 | 0.22 |
| 8 | 20936932 | 2.15E-05 | 4.66E-02 | 0.41 | -0.07 | -0.47 | 0.22 |
| 1 | 33547027 | 2.15E-05 | 4.66E-02 | -0.04 | -0.53 | -0.65 | -0.04 |
| 5 | 25745178 | 2.15E-05 | 4.66E-02 | -0.60 | -0.55 | -0.03 | 0.23 |
| 11 | 11254398 | 2.15E-05 | 4.67E-02 | -0.71 | -0.41 | -0.46 | -1.04 |
| 8 | 19763242 | 2.15E-05 | 4.67E-02 | -1.49 | 0.51 | 0.13 | 0.13 |
| 6 | 34152911 | 2.16E-05 | 4.67E-02 | 0.97 | 0.54 | -0.46 | 0.00 |
| 6 | 15410768 | 2.16E-05 | 4.67E-02 | 0.37 | -0.43 | -0.33 | 0.37 |
| 2 | 21677313 | 2.16E-05 | 4.67E-02 | -0.30 | -0.73 | -0.61 | 0.24 |
| 6 | 14590493 | 2.16E-05 | 4.68E-02 | 0.01 | 0.39 | 0.47 | -0.16 |
| 1 | 23283648 | 2.17E-05 | 4.68E-02 | -0.77 | -0.33 | -0.16 | -0.23 |
| 6 | 44921036 | 2.17E-05 | 4.69E-02 | -0.64 | -0.25 | -0.29 | -0.56 |
| 11 | 58384763 | 2.18E-05 | 4.70E-02 | 0.47 | -0.34 | -0.67 | 0.07 |
| 4 | 53732228 | 2.19E-05 | 4.71E-02 | 0.81 | -0.69 | -0.18 | -0.21 |
| 2 | 25607373 | 2.19E-05 | 4.71E-02 | -0.23 | -0.97 | -0.65 | 0.30 |
| 8 | 50572400 | 2.21E-05 | 4.74E-02 | -0.63 | 0.08 | 0.39 | 0.52 |
| 6 | 32226930 | 2.21E-05 | 4.74E-02 | 0.91 | -0.26 | -0.38 | -0.56 |
| 5 | 5193183 | 2.21E-05 | 4.74E-02 | 1.54 | -0.29 | 0.14 | -0.24 |
| 10 | 15958030 | 2.21E-05 | 4.74E-02 | -0.16 | -0.49 | 0.26 | 0.41 |
| 9 | 17267477 | 2.21E-05 | 4.74E-02 | -0.28 | -1.01 | -0.63 | 0.28 |
| 6 | 28490362 | 2.21E-05 | 4.74E-02 | 0.02 | 0.81 | -0.56 | 0.52 |
| 4 | 34897242 | 2.21E-05 | 4.74E-02 | 1.07 | -0.25 | -0.41 | -0.33 |
| 4 | 34898329 | 2.21E-05 | 4.74E-02 | 1.07 | -0.25 | -0.41 | -0.33 |
| 4 | 34898706 | 2.21E-05 | 4.74E-02 | 1.07 | -0.25 | -0.41 | -0.33 |
| 4 | 34900242 | 2.21E-05 | 4.74E-02 | 1.07 | -0.25 | -0.41 | -0.33 |
| 4 | 34901420 | 2.21E-05 | 4.74E-02 | 1.07 | -0.25 | -0.41 | -0.33 |
| 6 | 15345235 | 2.22E-05 | 4.74E-02 | 0.47 | -0.50 | -0.29 | 0.30 |
| 6 | 27579204 | 2.22E-05 | 4.76E-02 | -0.24 | 0.06 | 0.17 | 0.93 |
| 6 | 15538265 | 2.23E-05 | 4.76E-02 | -0.23 | -0.99 | -0.68 | 0.27 |
| 12 | 16251783 | 2.23E-05 | 4.76E-02 | -0.48 | -0.41 | 0.23 | 0.29 |
| 12 | 14732694 | 2.23E-05 | 4.76E-02 | 0.23 | -0.91 | -0.36 | 0.08 |
| 12 | 11413585 | 2.23E-05 | 4.76E-02 | 0.16 | -0.49 | -0.62 | 0.17 |
| 12 | 11413588 | 2.23E-05 | 4.76E-02 | 0.16 | -0.49 | -0.62 | 0.17 |
| 1 | 29660981 | 2.24E-05 | 4.77E-02 | -2.41 | 2.41 | 1.21 | 0.93 |
| 8 | 47665036 | 2.24E-05 | 4.78E-02 | 0.40 | 0.02 | 0.57 | -0.21 |
| 1 | 13318681 | 2.24E-05 | 4.78E-02 | 0.61 | -0.43 | 0.10 | 0.33 |
| 6 | 46015786 | 2.25E-05 | 4.78E-02 | 0.18 | -0.68 | -0.59 | -0.95 |
| 6 | 15354622 | 2.25E-05 | 4.78E-02 | 0.37 | -0.51 | -0.52 | 0.30 |
| 4 | 20443705 | 2.25E-05 | 4.78E-02 | -1.23 | 0.47 | -0.40 | 1.02 |
| 1 | 40475066 | 2.25E-05 | 4.78E-02 | -0.52 | 0.09 | 0.15 | 0.92 |
| 9 | 41028125 | 2.25E-05 | 4.78E-02 | 1.60 | -0.06 | -0.94 | 0.05 |
| 1 | 9695557 | 2.26E-05 | 4.79E-02 | -1.23 | 0.86 | 0.09 | 0.48 |
| 6 | 7161353 | 2.26E-05 | 4.79E-02 | 0.23 | -0.54 | -0.68 | -0.23 |

| | | | | | | | |
|---|---|---|---|---|---|---|---|
| 2 | 614224 | 2.27E-05 | 4.81E-02 | 0.19 | 0.32 | 0.40 | 0.39 |
| 11 | 51966541 | 2.27E-05 | 4.81E-02 | 0.07 | 0.21 | -0.41 | 0.36 |
| 6 | 35387410 | 2.27E-05 | 4.81E-02 | -0.03 | -0.53 | -0.52 | -0.19 |
| 12 | 4498420 | 2.28E-05 | 4.83E-02 | 0.03 | -0.31 | -0.65 | 0.12 |
| 8 | 36743916 | 2.29E-05 | 4.83E-02 | 0.35 | -0.29 | -0.63 | 0.27 |
| 8 | 36743964 | 2.29E-05 | 4.83E-02 | 0.35 | -0.29 | -0.63 | 0.27 |
| 6 | 45366087 | 2.29E-05 | 4.83E-02 | 0.94 | -0.15 | -0.50 | -0.17 |
| 8 | 30668822 | 2.29E-05 | 4.84E-02 | 2.25 | -0.85 | -0.71 | 0.26 |
| 8 | 35436593 | 2.30E-05 | 4.85E-02 | -0.10 | -0.82 | -0.55 | -0.11 |
| 6 | 21755709 | 2.31E-05 | 4.86E-02 | -0.42 | -0.21 | -0.59 | 0.04 |
| 5 | 44536035 | 2.31E-05 | 4.86E-02 | -0.89 | 0.02 | -0.15 | 0.51 |
| 11 | 35218579 | 2.31E-05 | 4.86E-02 | -0.12 | -0.07 | 0.44 | 0.80 |
| 7 | 32001555 | 2.32E-05 | 4.87E-02 | -0.22 | -0.30 | -0.14 | -0.46 |
| 1 | 43882204 | 2.32E-05 | 4.87E-02 | -0.64 | 0.43 | -0.34 | -1.19 |
| 6 | 42324700 | 2.32E-05 | 4.87E-02 | 0.35 | -0.74 | -0.38 | -0.57 |
| 12 | 16257993 | 2.32E-05 | 4.87E-02 | -0.56 | -0.31 | 0.31 | 0.28 |
| 6 | 10433335 | 2.33E-05 | 4.89E-02 | 0.25 | 0.08 | -0.66 | -0.38 |
| 4 | 54034021 | 2.33E-05 | 4.89E-02 | 0.90 | -0.37 | 0.32 | -0.31 |
| 6 | 50242287 | 2.33E-05 | 4.89E-02 | 0.30 | -1.29 | -0.22 | -0.15 |
| 11 | 33812833 | 2.34E-05 | 4.90E-02 | 1.08 | 0.04 | 0.32 | 0.11 |
| 10 | 6578893 | 2.34E-05 | 4.90E-02 | 0.97 | 0.05 | -0.20 | -0.05 |
| 6 | 10115585 | 2.34E-05 | 4.90E-02 | 1.09 | -0.45 | -0.84 | -0.68 |
| 9 | 26957067 | 2.34E-05 | 4.90E-02 | 1.64 | 0.43 | -1.18 | -0.49 |
| 1 | 15398677 | 2.35E-05 | 4.90E-02 | 1.19 | -0.38 | -0.20 | 0.05 |
| 9 | 38843678 | 2.35E-05 | 4.90E-02 | 0.22 | -0.74 | -0.33 | 0.14 |
| 11 | 11315358 | 2.35E-05 | 4.91E-02 | 0.34 | -0.08 | -1.04 | -0.32 |
| 1 | 28104356 | 2.36E-05 | 4.92E-02 | -0.40 | 1.40 | -0.54 | 0.56 |
| 7 | 37107743 | 2.36E-05 | 4.92E-02 | 0.18 | -0.51 | -0.49 | 0.23 |
| 4 | 81144054 | 2.36E-05 | 4.92E-02 | -0.07 | 0.94 | -0.63 | 0.02 |
| 6 | 15093202 | 2.36E-05 | 4.92E-02 | 0.00 | -1.17 | -0.05 | -0.56 |
| 11 | 54737820 | 2.37E-05 | 4.93E-02 | -1.05 | 0.52 | 0.00 | 0.22 |
| 6 | 14848968 | 2.37E-05 | 4.93E-02 | 1.09 | -0.69 | -0.18 | -0.61 |
| 12 | 15704617 | 2.37E-05 | 4.93E-02 | -1.05 | 0.59 | 0.17 | 0.27 |
| 12 | 15705267 | 2.37E-05 | 4.93E-02 | -1.05 | 0.59 | 0.17 | 0.27 |
| 10 | 8071254 | 2.37E-05 | 4.93E-02 | 0.26 | -0.41 | -0.06 | 0.47 |
| 8 | 43925117 | 2.38E-05 | 4.93E-02 | 0.30 | -0.30 | 0.46 | -0.23 |
| 12 | 4505313 | 2.38E-05 | 4.93E-02 | 0.19 | -0.37 | -0.64 | 0.01 |
| 5 | 31793530 | 2.38E-05 | 4.93E-02 | 0.45 | 0.44 | -0.53 | 0.27 |
| 1 | 40276143 | 2.38E-05 | 4.93E-02 | -0.21 | -0.01 | 0.27 | 0.69 |
| 1 | 40278290 | 2.38E-05 | 4.93E-02 | -0.21 | -0.01 | 0.27 | 0.69 |
| 1 | 40281902 | 2.38E-05 | 4.93E-02 | -0.21 | -0.01 | 0.27 | 0.69 |
| 1 | 40288812 | 2.38E-05 | 4.93E-02 | -0.21 | -0.01 | 0.27 | 0.69 |
| 6 | 14624394 | 2.38E-05 | 4.93E-02 | -0.24 | -0.22 | -0.16 | -0.70 |
| 6 | 16851798 | 2.39E-05 | 4.93E-02 | -0.77 | -0.08 | -0.02 | -0.54 |
| 8 | 23950469 | 2.39E-05 | 4.93E-02 | -0.99 | 0.05 | -0.09 | 0.43 |
| 8 | 23950500 | 2.39E-05 | 4.93E-02 | -0.99 | 0.05 | -0.09 | 0.43 |
| 7 | 29052983 | 2.39E-05 | 4.94E-02 | -0.49 | -0.68 | 0.67 | -0.82 |
| 10 | 30728015 | 2.40E-05 | 4.94E-02 | 0.09 | 0.09 | 0.04 | -0.55 |
| 2 | 14605591 | 2.41E-05 | 4.95E-02 | 0.33 | -0.76 | -0.33 | 0.09 |
| 2 | 14605875 | 2.41E-05 | 4.95E-02 | 0.33 | -0.76 | -0.33 | 0.09 |
| 2 | 14610124 | 2.41E-05 | 4.95E-02 | 0.33 | -0.76 | -0.33 | 0.09 |
| 2 | 14610227 | 2.41E-05 | 4.95E-02 | 0.33 | -0.76 | -0.33 | 0.09 |

| | | | | | | | |
|---|---|---|---|---|---|---|---|
| 2 | 14611641 | 2.41E-05 | 4.95E-02 | 0.33 | -0.76 | -0.33 | 0.09 |
| 2 | 14614385 | 2.41E-05 | 4.95E-02 | 0.33 | -0.76 | -0.33 | 0.09 |
| 2 | 14614692 | 2.41E-05 | 4.95E-02 | 0.33 | -0.76 | -0.33 | 0.09 |
| 4 | 41455238 | 2.41E-05 | 4.95E-02 | -0.29 | 0.11 | -0.78 | -0.91 |
| 9 | 38799258 | 2.41E-05 | 4.95E-02 | 1.54 | -0.76 | -0.40 | -0.24 |
| 10 | 44288656 | 2.41E-05 | 4.95E-02 | -1.16 | 0.82 | -0.65 | 0.08 |
| 5 | 25109546 | 2.42E-05 | 4.95E-02 | 0.20 | -0.89 | 0.31 | 0.64 |
| 6 | 6375375 | 2.42E-05 | 4.95E-02 | 0.65 | -0.54 | -0.28 | -0.15 |
| 6 | 53059683 | 2.43E-05 | 4.97E-02 | 1.40 | -0.12 | -0.26 | 0.23 |
| 12 | 16258110 | 2.43E-05 | 4.97E-02 | -0.62 | -0.33 | 0.29 | 0.23 |
| 12 | 13530094 | 2.44E-05 | 4.98E-02 | -0.60 | 0.46 | 0.54 | 0.47 |
| 8 | 31031548 | 2.44E-05 | 4.99E-02 | 0.03 | 0.14 | -0.63 | 0.16 |
| 6 | 30977585 | 2.44E-05 | 4.99E-02 | -1.04 | 0.20 | -0.09 | -0.19 |
| 6 | 7153154 | 2.45E-05 | 4.99E-02 | 0.20 | -0.44 | -0.65 | -0.22 |
| 6 | 8071358 | 2.45E-05 | 4.99E-02 | 0.72 | 0.06 | -0.06 | 0.46 |
| 6 | 25925404 | 2.45E-05 | 4.99E-02 | 0.59 | -0.48 | -0.69 | -0.93 |
| 5 | 20423101 | 2.46E-05 | 4.99E-02 | 0.95 | -0.47 | -0.17 | -0.39 |
| 1 | 13321516 | 2.47E-05 | 4.99E-02 | 0.53 | -0.46 | 0.01 | 0.35 |
| 1 | 13345378 | 2.47E-05 | 4.99E-02 | 0.53 | -0.46 | 0.01 | 0.35 |
| 1 | 13350300 | 2.47E-05 | 4.99E-02 | 0.53 | -0.46 | 0.01 | 0.35 |
| 9 | 29148500 | 2.47E-05 | 4.99E-02 | 0.67 | -0.34 | -0.13 | 0.19 |
| 9 | 29148572 | 2.47E-05 | 4.99E-02 | 0.67 | -0.34 | -0.13 | 0.19 |
| 9 | 29149204 | 2.47E-05 | 4.99E-02 | 0.67 | -0.34 | -0.13 | 0.19 |
| 9 | 29164797 | 2.47E-05 | 4.99E-02 | 0.67 | -0.34 | -0.13 | 0.19 |
| 9 | 29168778 | 2.47E-05 | 4.99E-02 | 0.67 | -0.34 | -0.13 | 0.19 |
| 9 | 29169360 | 2.47E-05 | 4.99E-02 | 0.67 | -0.34 | -0.13 | 0.19 |
| 9 | 29170215 | 2.47E-05 | 4.99E-02 | 0.67 | -0.34 | -0.13 | 0.19 |
| 6 | 28524684 | 2.47E-05 | 4.99E-02 | -0.05 | 0.61 | -0.47 | 0.42 |
| 6 | 41740114 | 2.47E-05 | 4.99E-02 | 1.06 | -0.48 | -0.25 | 0.33 |
| 10 | 18009784 | 2.47E-05 | 5.00E-02 | -0.51 | -0.22 | 0.41 | -0.28 |
| 6 | 15154485 | 2.47E-05 | 5.00E-02 | 0.06 | -0.09 | -0.65 | -0.10 |
| 4 | 81142284 | 2.47E-05 | 5.00E-02 | -0.41 | 0.88 | -0.54 | 0.32 |

**Table S11. Within-chromosomal linkage disequilibrium at candidate maximum distances used for gene assignment.**

| Population | No. individuals | No. SNPs | Mean r2 at 5 kb | Mean r2 at 10 kb | Mean r2 at 25 kb | Mean r2 at 50 kb |
|---|---|---|---|---|---|---|
| *Q. mongolica* | 56 | 2,162,224 | 0.185 | 0.156 | 0.108 | 0.071 |
| *Q. serrata* | 60 | 2,701,087 | 0.170 | 0.144 | 0.101 | 0.066 |

**Table S12. Enriched GO molecular function pathways from ShinyGO analyses.** Shown separately for *Quercus lobata* and *Arabidopsis thaliana* annotations: term ID/name, gene count, gene ratio, background size, and adjusted P-values (Benjamini–Hochberg FDR, as specified in the Methods).

| **Reference: *Quercus lobata* (ValleyOak3.0)** | | | | |
|---|---|---|---|---|
| **Pathway name** | **Fold enrichment** | **adjP** | **nGenes** | **Genes** |
| **GO:0035251 Udp-glucosyltransferase activity** | 17.3 | 0.002 | 5 | QL04P068884 QL06P016165 QL06P016377 QL08P044680 QL09P031258 |
| **GO:0046527 Glucosyltransferase activity** | 12.4 | 0.005 | 5 | QL04P068884 QL06P016165 QL06P016377 QL08P044680 QL09P031258 |

| **Reference: *Arabidopsis thaliana* (TAIR10)** | | | | |
|---|---|---|---|---|
| **Pathway name** | **Fold enrichment** | **adjP** | **nGenes** | **Genes** |
| GO:0016759 Cellulose synthase activity | 22.4 | 0.020 | 3 | CSLG3 CSLG2 CSLG1 |
| GO:0016760 Cellulose synthase (udp-forming) activity | 22.4 | 0.020 | 3 | CSLG3 CSLG2 CSLG1 |
| GO:0019187 Beta-1,4-mannosyltransferase activity | 27.2 | 0.020 | 3 | CSLG3 CSLG2 CSLG1 |
| GO:0051753 Mannan synthase activity | 27.2 | 0.020 | 3 | CSLG3 CSLG2 CSLG1 |
| **GO:0035251 Udp-glucosyltransferase activity** | 7.6 | 0.027 | 5 | CalS7 TPS1 CSLG3 CSLG2 CSLG1 |
| GO:0008194 Udp-glycosyltransferase activity | 4.7 | 0.032 | 7 | CalS7 AT5G49690 TPS1 CSLG3 CSLG2 CSLG1 AT5G12890 |
| GO:0000030 Mannosyltransferase activity | 14.6 | 0.039 | 3 | CSLG3 CSLG2 CSLG1 |
| **GO:0046527 Glucosyltransferase activity** | 6.3 | 0.039 | 5 | CalS7 TPS1 CSLG3 CSLG2 CSLG1 |

**Table S13. Inter-chromosomal linkage disequilibrium (ICLD) across Δp bins: summary statistics and comparisons between trait-associated loci and genomic background.** For each Δp bin: mean, median, 95% CI of r², Wilcoxon test P-values, and effect sizes such as Cliff's delta.

| Group | Bin | Category | n | mean | median | sd | se | CI_lower | CI_upper | P-value | diff_median |
|---|---|---|---|---|---|---|---|---|---|---|---|
| Hybrid | [0.20,0.25] | trait loci | 92 | 0.0098 | 0.0048 | 0.0172 | 0.0018 | 0.0063 | 0.0133 | 0.4317 | 0.0014 |
| | [0.20,0.25] | background | 193 | 0.0067 | 0.0034 | 0.0086 | 0.0006 | 0.0055 | 0.0079 | | |
| | [0.25,0.30] | trait loci | 70 | 0.0233 | 0.0099 | 0.0369 | 0.0044 | 0.0147 | 0.0320 | **0.0075** | **0.0038** |
| | [0.25,0.30] | background | 241 | 0.0097 | 0.0061 | 0.0113 | 0.0007 | 0.0083 | 0.0111 | | |
| | [0.30,0.35] | trait loci | 171 | 0.0442 | 0.0206 | 0.0525 | 0.0040 | 0.0364 | 0.0521 | **0.0000** | **0.0104** |
| | [0.30,0.35] | background | 213 | 0.0163 | 0.0102 | 0.0201 | 0.0014 | 0.0136 | 0.0190 | | |
| | [0.35,0.40] | trait loci | 299 | 0.0819 | 0.0669 | 0.0712 | 0.0041 | 0.0738 | 0.0899 | **0.0000** | **0.0524** |
| | [0.35,0.40] | background | 344 | 0.0216 | 0.0145 | 0.0277 | 0.0015 | 0.0187 | 0.0246 | | |
| | [0.40,0.45] | trait loci | 246 | 0.1010 | 0.0880 | 0.0772 | 0.0049 | 0.0914 | 0.1107 | **0.0000** | **0.0657** |
| | [0.40,0.45] | background | 273 | 0.0312 | 0.0223 | 0.0348 | 0.0021 | 0.0270 | 0.0353 | | |
| | [0.45,0.50] | trait loci | 295 | 0.1363 | 0.1346 | 0.0754 | 0.0044 | 0.1277 | 0.1449 | **0.0000** | **0.1092** |
| | [0.45,0.50] | background | 319 | 0.0355 | 0.0254 | 0.0387 | 0.0022 | 0.0313 | 0.0398 | | |
| | [0.50,0.55] | trait loci | 235 | 0.1158 | 0.0938 | 0.0814 | 0.0053 | 0.1054 | 0.1262 | **0.0000** | **0.0590** |
| | [0.50,0.55] | background | 270 | 0.0467 | 0.0348 | 0.0435 | 0.0026 | 0.0415 | 0.0519 | | |
| | [0.55,0.60] | trait loci | 277 | 0.1489 | 0.1235 | 0.0995 | 0.0060 | 0.1371 | 0.1606 | **0.0000** | **0.0773** |
| | [0.55,0.60] | background | 320 | 0.0625 | 0.0462 | 0.0524 | 0.0029 | 0.0568 | 0.0682 | | |
| | [0.60,0.65] | trait loci | 414 | 0.1446 | 0.1202 | 0.0954 | 0.0047 | 0.1354 | 0.1537 | **0.0000** | **0.0532** |
| | [0.60,0.65] | background | 460 | 0.0755 | 0.0670 | 0.0505 | 0.0024 | 0.0709 | 0.0801 | | |
| *Quercus mongolica* | [0.20,0.25] | trait loci | 92 | 0.0177 | 0.0078 | 0.0240 | 0.0025 | 0.0128 | 0.0226 | 0.0866 | 0.0033 |
| | [0.20,0.25] | background | 193 | 0.0104 | 0.0045 | 0.0162 | 0.0012 | 0.0082 | 0.0127 | | |
| | [0.25,0.30] | trait loci | 70 | 0.0101 | 0.0064 | 0.0115 | 0.0014 | 0.0074 | 0.0128 | 0.4750 | 0.0004 |
| | [0.25,0.30] | background | 241 | 0.0120 | 0.0060 | 0.0149 | 0.0010 | 0.0102 | 0.0139 | | |
| | [0.30,0.35] | trait loci | 171 | 0.0115 | 0.0056 | 0.0151 | 0.0012 | 0.0093 | 0.0138 | 0.3558 | 0.0012 |
| | [0.30,0.35] | background | 213 | 0.0093 | 0.0044 | 0.0120 | 0.0008 | 0.0077 | 0.0109 | | |
| | [0.35,0.40] | trait loci | 299 | 0.0117 | 0.0064 | 0.0142 | 0.0008 | 0.0100 | 0.0133 | 0.1099 | 0.0008 |
| | [0.35,0.40] | background | 344 | 0.0095 | 0.0056 | 0.0151 | 0.0008 | 0.0079 | 0.0111 | | |
| | [0.40,0.45] | trait loci | 246 | 0.0134 | 0.0063 | 0.0179 | 0.0011 | 0.0111 | 0.0156 | **0.0177** | **0.0021** |
| | [0.40,0.45] | background | 273 | 0.0089 | 0.0042 | 0.0132 | 0.0008 | 0.0073 | 0.0105 | | |
| | [0.45,0.50] | trait loci | 295 | 0.0109 | 0.0057 | 0.0126 | 0.0007 | 0.0095 | 0.0124 | 0.2487 | 0.0003 |
| | [0.45,0.50] | background | 319 | 0.0104 | 0.0054 | 0.0138 | 0.0008 | 0.0089 | 0.0119 | | |
| | [0.50,0.55] | trait loci | 235 | 0.0123 | 0.0068 | 0.0149 | 0.0010 | 0.0104 | 0.0142 | 0.1083 | 0.0020 |
| | [0.50,0.55] | background | 270 | 0.0112 | 0.0048 | 0.0182 | 0.0011 | 0.0090 | 0.0134 | | |
| | [0.55,0.60] | trait loci | 277 | 0.0148 | 0.0074 | 0.0189 | 0.0011 | 0.0126 | 0.0170 | **0.0088** | **0.0026** |
| | [0.55,0.60] | background | 320 | 0.0100 | 0.0048 | 0.0132 | 0.0007 | 0.0086 | 0.0115 | | |
| | [0.60,0.65] | trait loci | 414 | 0.0137 | 0.0096 | 0.0150 | 0.0007 | 0.0123 | 0.0152 | **0.0000** | **0.0056** |
| | [0.60,0.65] | background | 460 | 0.0100 | 0.0040 | 0.0185 | 0.0009 | 0.0083 | 0.0117 | | |
| *Q. serrata* | [0.20,0.25] | trait loci | 92 | 0.0170 | 0.0057 | 0.0197 | 0.0021 | 0.0130 | 0.0211 | 0.1156 | 0.0010 |
| | [0.20,0.25] | background | 193 | 0.0093 | 0.0047 | 0.0127 | 0.0009 | 0.0075 | 0.0111 | | |
| | [0.25,0.30] | trait loci | 70 | 0.0088 | 0.0029 | 0.0165 | 0.0020 | 0.0049 | 0.0126 | 0.9261 | -0.0010 |
| | [0.25,0.30] | background | 241 | 0.0088 | 0.0039 | 0.0131 | 0.0008 | 0.0071 | 0.0104 | | |
| | [0.30,0.35] | trait loci | 171 | 0.0094 | 0.0024 | 0.0245 | 0.0019 | 0.0057 | 0.0131 | 0.0424 | -0.0015 |
| | [0.30,0.35] | background | 213 | 0.0083 | 0.0039 | 0.0105 | 0.0007 | 0.0069 | 0.0097 | | |
| | [0.35,0.40] | trait loci | 299 | 0.0128 | 0.0012 | 0.0423 | 0.0024 | 0.0080 | 0.0176 | **0.0085** | **-0.0033** |
| | [0.35,0.40] | background | 344 | 0.0092 | 0.0045 | 0.0157 | 0.0008 | 0.0075 | 0.0109 | | |
| | [0.40,0.45] | trait loci | 246 | 0.0052 | 0.0007 | 0.0099 | 0.0006 | 0.0040 | 0.0065 | **0.0000** | **-0.0054** |
| | [0.40,0.45] | background | 273 | 0.0099 | 0.0061 | 0.0140 | 0.0008 | 0.0082 | 0.0115 | | |
| | [0.45,0.50] | trait loci | 295 | 0.0053 | 0.0017 | 0.0110 | 0.0006 | 0.0040 | 0.0065 | **0.0000** | **-0.0034** |
| | [0.45,0.50] | background | 319 | 0.0092 | 0.0050 | 0.0119 | 0.0007 | 0.0079 | 0.0105 | | |
| | [0.50,0.55] | trait loci | 235 | 0.0089 | 0.0010 | 0.0234 | 0.0015 | 0.0059 | 0.0119 | **0.0000** | **-0.0035** |
| | [0.50,0.55] | background | 270 | 0.0096 | 0.0045 | 0.0127 | 0.0008 | 0.0081 | 0.0111 | | |
| | [0.55,0.60] | trait loci | 277 | 0.0130 | 0.0033 | 0.0260 | 0.0016 | 0.0099 | 0.0161 | 0.3630 | -0.0021 |
| | [0.55,0.60] | background | 320 | 0.0093 | 0.0053 | 0.0116 | 0.0006 | 0.0080 | 0.0105 | | |
| | [0.60,0.65] | trait loci | 414 | 0.0072 | 0.0028 | 0.0105 | 0.0005 | 0.0062 | 0.0083 | **0.0000** | **-0.0026** |
| | [0.60,0.65] | background | 460 | 0.0100 | 0.0054 | 0.0128 | 0.0006 | 0.0088 | 0.0112 | | |

**Table S14. Among-group comparisons of ICLD at trait-associated loci (Hybrid, *Q. mongolica* , *Q. serrata* ).** Δp-binned pairwise tests with summary statistics, P-values, effect sizes, and sample sizes.

| bin | Hybrid-Qmon | | Hybrid-Qser | | Qmon-Qser | |
|---|---|---|---|---|---|---|
| | Diff_median | P-value | Diff_median | P-value | Diff_median | P-value |
| [0.20,0.25] | -0.003 | 0.022 | -0.001 | 0.043 | 0.002 | 0.864 |
| [0.25,0.30] | 0.003 | 0.036 | 0.007 | 0.003 | 0.003 | 0.458 |
| [0.30,0.35] | 0.015 | 0.000 | 0.018 | 0.000 | 0.003 | 0.002 |
| [0.35,0.40] | 0.060 | 0.000 | 0.066 | 0.000 | 0.005 | 0.000 |
| [0.40,0.45] | 0.082 | 0.000 | 0.087 | 0.000 | 0.006 | 0.000 |
| [0.45,0.50] | 0.129 | 0.000 | 0.133 | 0.000 | 0.004 | 0.000 |
| [0.50,0.55] | 0.087 | 0.000 | 0.093 | 0.000 | 0.006 | 0.000 |
| [0.55,0.60] | 0.116 | 0.000 | 0.120 | 0.000 | 0.004 | 0.004 |
| [0.60,0.65] | 0.111 | 0.000 | 0.117 | 0.000 | 0.007 | 0.000 |

**Table S15. Between-group comparisons of local cline slope parameters.** Median and 95% CI of v from bgc-hm, with trait-loci vs background contrasts and bootstrap estimates of Pr(diff > 0) based on 5,000 resamples.

| Metric | Mean diff | 95%CI (lower) | 95%CI (upper) | Pr (diff >0) |
|---|---|---|---|---|
| Median | 0.052 | -0.100 | 0.232 | 0.732 |
| 95%CI (lower) | 0.064 | 0.009 | 0.114 | **0.987** |
| 95%CI (upper) | 0.236 | 0.053 | 0.426 | **0.996** |
